# Feature Reuse and Scaling: Understanding Transfer Learning with Protein Language Models

**DOI:** 10.1101/2024.02.05.578959

**Authors:** Francesca-Zhoufan Li, Ava P. Amini, Yisong Yue, Kevin K. Yang, Alex X. Lu

## Abstract

Large pretrained protein language models (PLMs) have improved protein property and structure prediction from sequences via transfer learning, in which weights and representations from PLMs are repurposed for downstream tasks. Although PLMs have shown great promise, currently there is little understanding of how the features learned by pretraining relate to and are useful for downstream tasks. We perform a systematic analysis of transfer learning using PLMs, conducting 370 experiments across a comprehensive suite of factors including different downstream tasks, architectures, model sizes, model depths, and pretraining time. We observe that while almost all down-stream tasks do benefit from pretrained models compared to naive sequence representations, for the majority of tasks performance does not scale with pretraining, and instead relies on low-level features learned early in pretraining. Our results point to a mismatch between current PLM pretraining paradigms and most applications of these models, indicating a need for better pretraining methods.

## 1. Introduction

Proteins perform a myriad of critical biological functions, and thus the ability to design proteins has vast impacts on healthcare, environment, and industry (Lutz & Iamurri, 2018). Since a protein’s function is largely determined by its amino acid sequence, specifying a sequence that will yield a desired function is feasible in principle. However, the relationship between amino acid sequence and function remains poorly understood, and most experimental methods for measuring function are costly and low-throughput (Maynard Smith, 1970; Romero & Arnold, 2009). To overcome the challenge presented by limited labelled data, researchers have sought to use transfer learning, in which models are pretrained in a self-supervised fashion on large public datasets in the hope that the pretrained features or model weights will improve performance on downstream tasks where supervised data is limited (Fig. 1a-b).

**Figure 1.**
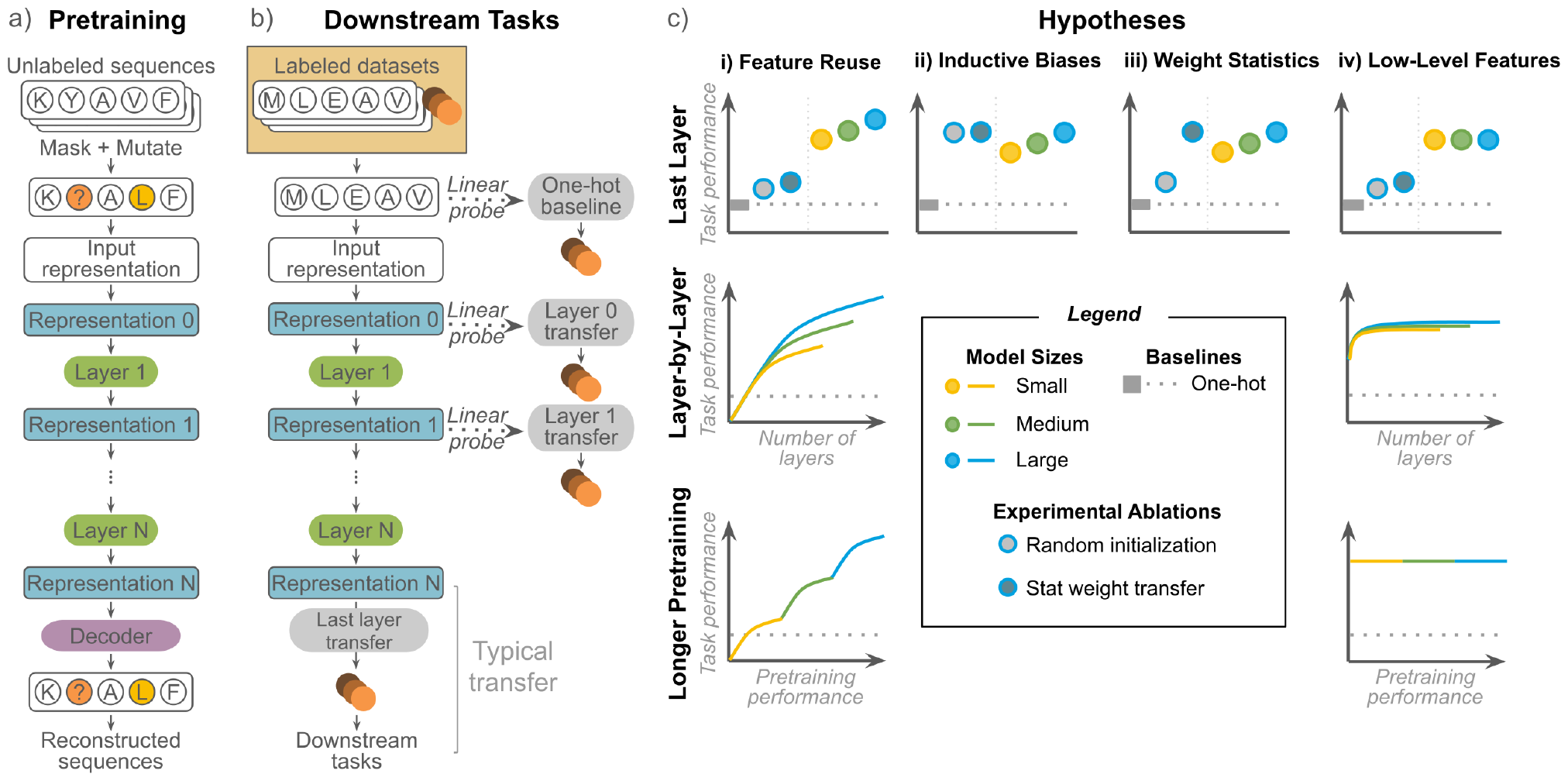
Summary of the transfer learning procedure and our analyses. a) PLMs are pretrained using masked language modeling. b) Typically, transfer learning uses respresentations from the last layer of the PLM for downstream tasks. We evaluate downstream task performance at every layer in the model. c) We compare to baselines and ablations and evaluate the effects of PLM size, model depth, and pretraining time. These experiments characterize behavior consistent with either feature reuse (i), or an alternative hypothesis (inductive biases/overparameterization - ii, weight statistics - iii, or reuse of low-level features only - iv).

Protein language models (PLMs) have emerged as the most popular framework for transfer learning for proteins (Rives et al., 2021; Yang et al., 2022; Elnaggar et al., 2022; Brandes et al., 2022; Alley et al., 2019; Elnaggar et al., 2023; Lin et al., 2023). Most PLMs pretrain using the masked language modeling (MLM) task, in which the model is trained to predict the original identity of masked or corrupted amino acids. PLMs have been effective at improving performance on many protein function prediction tasks, and some are now integrated into bioinformatics and structure prediction tools (Teufel et al., 2022; Thumuluri et al., 2022; Wu et al., 2022; Flamholz et al., 2024). Despite their widespread adoption, it is not understood how or why PLMs improve performance on downstream tasks.

Drawing from other domains like computer vision where investigations of transfer learning are more established, we synthesize a set of possible hypotheses to explain improvement in downstream tasks, and design and conduct a comprehensive series of experiments to test them. We structure our study around the following hypotheses:

### Feature reuse (Fig. 1c-i)

One popular hypothesis is that MLM pretraining learns general features of protein biology, and that these features can be re-used across tasks. Previous work has shown that transfer learning improves performance across diverse downstream tasks (Rao et al., 2019; Dallago et al., 2021). However, the degree of feature reuse is also important: ideally, the pretrain and downstream tasks should be *aligned*, such that transferring PLM representations improves downstream function prediction accuracy and that this improvement increases with larger model sizes, deeper layers, and better pretraining performance.

If this does not occur, it suggests that pretraining primarily learns features that cannot be reused on downstream tasks. To determine whether or not this is the case for PLMs, we explore three alternative hypotheses.

### Inductive biases and overparameterization (Fig. 1c-ii)

The large number of parameters in pretrained models may lead to some alignment with useful signal by chance (Raghu et al., 2019). If inductive biases are sufficient, then transferring from randomly-initialized version of the same model architecture should provide similar performance.

### Statistics of pretrained weights (Fig. 1c-iii)

The primary benefit of pretraining may be initializing weights to a sensible scale (Raghu et al., 2019; Matsoukas et al., 2022). If pretraining primarily provides better weight initialization, resampling weights from the empirical distribution after pretraining should provide similar performance.

### Reuse of low-level features (Fig. 1c-iv)

It is possible for only less complex features learned early in pretraining to contribute to transfer learning (Neyshabur et al., 2021). If low-level features are sufficient, then features extracted from earlier layers of the pretrained model may provide better or similar performance to those extracted from the last layer. Similarly, earlier pretraining checkpoints or smaller, less performant models should provide similar performance to the full-size, fully-pretrained model.

Critically, while all three alternative hypotheses can still lead to improvements in downstream task performance, they do not predict that downstream task performance can be improved by transferring representations from larger, better-trained models (Raghu et al., 2019; Abnar et al., 2022).

### Contributions

Our work evaluates the scalability of transfer learning for PLMs and makes the following contributions:

1. The most comprehensive evaluation, to date and to the best of our knowledge, of transfer learning with PLMs, spanning 370 experiments over a diverse suite of downstream tasks.

2. The discovery that current MLM pretraining paradigms underserve many aspects of protein biology, as supported empirically by evidence from both structure and function prediction tasks.

3. Systematic evidence that performance on many protein property prediction tasks does not scale with PLM size or pretraining. Our results uncouple improvements in downstream performance from scaling properties.

Together, our results predict that scaling PLMs under current pretraining paradigms may not improve performance on many protein function prediction tasks and charts a direction for identifying new, better-aligned pretraining tasks.

## 2. Related Work

### 2.1 Pretrained Protein Language Models

While numerous pretrained PLMs have been proposed in the past few years (Rives et al., 2021; Yang et al., 2022; Elnaggar et al., 2022; Brandes et al., 2022; Alley et al., 2019; Rao et al., 2019; Elnaggar et al., 2023; Lin et al., 2023), these works primarily focus on validating that pretraining improves performance on downstream tasks. In contrast, our work primarily seeks to understand the factors impacting transfer learning, which have not been rigorously studied to date for PLMs. Most PLM studies include comparisons to models with randomly initialized weights (Rives et al., 2021; Yang et al., 2022) to confirm that pretrained models do not improve downstream task performance due to overparameterization or inductive biases alone. Other studies show that under some circumstances, PLMs yield no detectable improvement over a simple one-hot representation of sequences (Wittmann et al., 2021; Hsu et al., 2021; Dallago et al., 2021). Compared to these individual baselines and benchmarks, our paper conducts a systematic analysis over many different factors impacting transfer learning.

The most similar work to ours is Detlefsen et al. (2022), which analyzes the effects of model architecture, fine-tuning, and different pooling schemes on transfer learning performance. However, we use MLMs trained on complete sequences instead of autoregressive models trained on Pfam domains. While they train proprietary, unreleased models for analysis, we use established models in the public domain. This makes our analysis more relevant to applications currently using these models and also improves documentation around these models. For example, neither their paper nor their released code describes the pretrained models in detail, so it is uncertain what the size of their model is, whereas we systematically vary the model size. More importantly, we evaluate a larger and more diverse set of downstream tasks with experiments designed to differentiate possible mechanisms by which transfer learning improves performance on downstream tasks. Critically, our systematic analysis identifies cases where transfer from PLMs is empirically effective in improving downstream task performance but the improvement is due to factors that are not expected to scale with further pretraining or larger models.

### 2.2 Understanding Transfer Learning in Computer Vision

While our analysis is differentiated as we focus on protein sequences, we take inspiration from computer vision studies that have sought to understand factors underlying successful transfer learning. Many are motivated by the observation that ImageNet-trained models are effective when transferred to medical images, raising the question of whether transfer performance is really due to reuse of features (given the extreme mismatch in domain), or due to more trivial factors. Raghu et al. (2019) compare pretrained models against random initialization to demonstrate that in some situations transfer performance is due to overparameterization. By randomly initializing models to match the weight statistics of pretrained models, the authors further demonstrate that improvements from pretraining may arise from good weight scalings rather than learning reusable features. By scrambling input images, Neyshabur et al. (2021) show that improvements from transfer learning can at least partially be attributed to the pretrained models learning low-level statistics of data rather than more sophisticated feature use. Matsoukas et al. (2022) further demonstrate that these factors vary depending upon downstream task dataset and model architecture.

Beyond models pretrained on ImageNet, some papers have looked at factors more specific to self-supervised pretraining. Abnar et al. (2022) show that improvements on the self-supervised pretraining task do not necessarily translate to improved performance on downstream tasks, and in some cases, are even anti-correlated. Pioneering work in generative self-supervised models also demonstrates that these models often saturate in downstream task performance in an intermediate layer of the model and degrade after (Jing & Tian, 2020). This is reinforced by empirical studies showing that the representations learned by self-supervised models versus supervised models rapidly diverge in the last few layers (Grigg et al., 2021), underscoring the importance of a layer-by-layer evaluation.

## 3. Datasets and Pretrained Models

To understand why and when transfer learning with PLMs improves downstream performance and how the improvements scale with increasingly large PLMs, we conducted 370 experiments on a diverse suite of downstream tasks with PLMs of different sizes, architectures, and at different check-points in training. The downstream tasks are summarized in Tables 1 and A1.

**Table 1.** Summary of downstream prediction tasks.

| Dataset | Description | Tasks | Task type |
| --- | --- | --- | --- |
| SS3 | Secondary structure | CB513, TS115, CASP12 | Residue-level classification |
| Thermostability | Melting temperature | Thermostability | Regression |
| Subcellular localization | Cellular location | Subcellular localization | Classification |
| GB1 | Immunoglobulin binding | Sampled, low vs. high, two vs. rest | Regression |
| AAV | Viral viability | Two vs. many, one vs. many | Regression |

### 3.1 Downstream Tasks

We test a diverse set of tasks covering both property and structure prediction, different types of distribution shift relevant to protein engineering, and global versus local variation over the sequence.

#### Structure prediction

We use the three-class secondary structure (SS3) task from TAPE with three independent test sets, SS3 – CB513 (Cuff & Barton, 1999), SS3 – TS115 (Yang et al., 2016), and SS3 – CASP12 (Moult et al., 2018), where the objective is to predict whether each residue belongs to an *α*-helix, *β*-strand, or coil (Rao et al., 2019).

#### Property prediction

We use the thermostability, subcellular localization, GB1, and AAV datasets from FLIP (Dallago et al., 2021).

Thermostability and subcellular localization are global protein properties measured for sequences spanning different functional families and domains of life. The thermostability dataset measures the melting temperature of 48,000 proteins across 13 species (Jarzab et al., 2020). Subcellular localization is a classification task predicting the cell compartment to which a eukaryotic protein localizes (Armenteros et al., 2017; Stärk et al., 2021).

In contrast, the GB1 and AAV datasets measure the effects of local sequence variation. GB1 is the 56 amino-acid B1 domain of protein G, an immunoglobulin-binding protein. The GB1 dataset covers binding measurements for simultaneous mutations of up to 4 interactive sites (Wu et al., 2016). VP1 is an adeno-associated virus (AAV) capsid protein, over 700 amino acids long (Bryant et al., 2021). The AAV dataset measures the effects of sparsely sampled mutations across a contiguous 28 amino-acid region over the binding interface on viral viability.

For GB1 and AAV, FLIP provides different train-test splits with different distribution shifts, including sampled (in-distribution) and out-of-distribution splits, as described in Table A1. Out-of-distribution splits more closely resemble protein engineering applications where a few low-functioning variants with a limited number of mutations are initially generated, but high-functioning variants across the larger sequence space are the engineering end goal. For GB1, we test three splits, in order of increasing difficulty:

- **Sampled:** sequences randomly partitioned between 80% training and 20% testing.
- **Low vs high:** models are trained on mutants with function worse than the parent and tested on those with better function.
- **Two vs rest:** Models are trained on single and double mutants and tested on triple and quadruple mutants.

For AAV, we test two splits, in order of increasing difficulty:

- **Two vs many:** Models are trained on single and double mutants and tested on variants with three or more mutations.
- **One vs many:** Models are trained on single mutants and tested on variants with more mutations.

### 3.2 Transfer Learning with Protein Language Models

While a number of pretraining tasks have been proposed for protein sequences, we focused on models trained using the popular BERT (Devlin et al., 2019) masked language modeling (MLM) task. During pretraining, 15% of tokens are randomly selected. Of the 15%, 10% are replaced with a special masking token, 2.5% are randomly changed to another token, and the remaining 2.5% are unperturbed to encourage the model to preserve the input sequence. The corrupted sequence is passed to the model, which is trained to maximize the probability of the original tokens at the selected locations.

To evaluate the effect of model architecture, we chose two families of protein MLMs with comparable model sizes trained on UniRef50 (Suzek et al., 2015): the ESM (Rives et al., 2021) family of transformer models and the Convolutional Autoencoding Representations of Proteins (CARP) (Yang et al., 2022) family of convolutional models. Due to the sequence length limit of the ESM-1b transformer model, the first and last 511 amino acids were taken for all sequences exceeding 1022 amino acids. This length restriction chiefly impacts the subcellular localization dataset: targeting signals often occur at the N-or C-terminal, and we reason that taking both terminals preserves biologically-relevant signals.

Following standard protein transfer learning practice when resources for full finetuning are not available (Dallago et al., 2021), we pass representations from each PLM layer to a linear model and compare the performance to a linear model on the one-hot encoding of the sequence for each task (Fig. 1b). For the SS3 and subcellular localization tasks, we train linear classifiers with mini-batches in PyTorch and perform early stopping based on the validation set. For the regression tasks, we train ridge regression models with Scikit-learn (Buitinck et al., 2013), using a grid search on the validation set to tune the regularization strength. For all tasks except secondary structure prediction, we mean pool the representations over the length dimension from each layer. Secondary structure prediction requires a representation for every residue, so no pooling is performed.

As protein engineers often seek to identify top-ranked mutants as opposed to predicting the absolute function of mutations, we use ranking metrics, Spearman’s rank correlation and Normalized Discounted Cumulative Gain (NDCG), as the primary metrics for the regression tasks. For concision, we report Spearman’s rank correlation for regression tasks and accuracy for classification tasks in the main text. Complete results, including mean square error, cross-entropy loss, NDCG, and ROC-AUC are provided in the Supplemental Materials.

## 4. Experimental Setup

### Baseline and ablations

We conduct baselines and model ablations to determine when transfer learning improves downstream task performance and whether improvements in downstream task performance can be attributed to mechanisms other than feature reuse (Fig. 1b-ii and 1b-iii).

- **One-hot baseline** 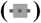. To determine whether transfer learning with PLMs improves performance, we test if representations from pretrained models perform better than a one-hot representation.
- **Random init** 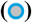. To evaluate whether the effect of transfer learning is due to overparameterization and/or the inductive biases of the PLM architecture, we test the impact of randomly initialized weights.
- **Stat transfer** 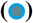. To evaluate whether the effect of transfer learning is due to weight statistics and/or initializing the weights to a sensible scale, we test the impact of randomly initialized weights matching the weight distribution of the pretrained PLM.

We consider transfer learning from a PLM to have improved performance over a baseline or ablation if it improves the metric by at least 10%.

### Scaling experiments

To further understand if the MLM pretraining task is aligned with downstream tasks, we sought to understand if improving PLM performance by scaling across three factors also improves transfer learning performance on downstream tasks (Fig. 1b-iv):

- **Model size**. For both CARP 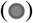 and ESM 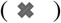, we test models with different numbers of layers and parameters (Table A2). For concision, we refer to CARP-38M and ESM-43M as the “small” models 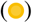, CARP-76M and ESM-85M as the “medium” models 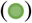, and CARP-640M and ESM-650M (ESM-1b) as the “large” models 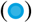.
- **Model depth**. For each architecture (CARP: 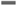, ESM: 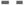 and model size 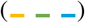, we test whether downstream task performance improves as we transfer deeper layers by determining whether the Spearman rank correlation between layer number and performance is greater than 0.9 (Table A6). This experiment allows us to understand if tasks primarily reuse low-level features early in the pretrained models, or if more complex features deeper in the models also contribute to downstream task performance. Convolutional neural networks (CNNs) induce a stronger correlation between the depth of the layer and the complexity of the features than transformers, leading to different patterns of feature reuse in previous transfer learning studies (Matsoukas et al., 2022). However, we find little empirical difference between CNNs (CARP) and transformers (ESM) in our analyses.
- **Model checkpoint**. For each model size 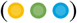, we test the effect of using checkpoints from earlier in pretraining. We order these checkpoints based upon their pretraining loss, as earlier checkpoints have higher losses on the MLM pretraining task (Table A7). We evaluate whether features from later in pretraining improve transfer learning by determining whether the Spearman rank correlation between the negative pretrain loss and downstream performance is greater than 0.9 (Table A8). Unfortunately, checkpoints are only publicly available for CARP, so we cannot run this analysis with ESM.

We define the MLM pretraining task to be *aligned* with a downstream task if transferring PLM representations improves downstream task performance over the baseline and ablations *and* this improvement scales with improvements to pretraining. Code for all experiments is available at https://github.com/microsoft/protein-transfer

## 5. Results

Overall, our analyses reveal three clusters of transfer learning behavior across downstream tasks (Figure 2). First, we find that secondary structure prediction tasks are the only tasks where pretraining improves downstream performance and the pretrain and downstream tasks are aligned. Second, we observe that transfer learning improves performance for many downstream tasks despite the pretrain and downstream tasks not being well-aligned, indicating that performance on these tasks will not improve as PLMs improve. Third, we observe that although transfer learning improves performance on almost all downstream tasks, for some tasks this improvement can be attributed to overparameterization, inductive biases, or sensible weight initialization. In subsequent sections, we expand on each of these clusters of observations in detail. The full results of our experiments are available in the Supplemental Materials.

**Figure 2.**
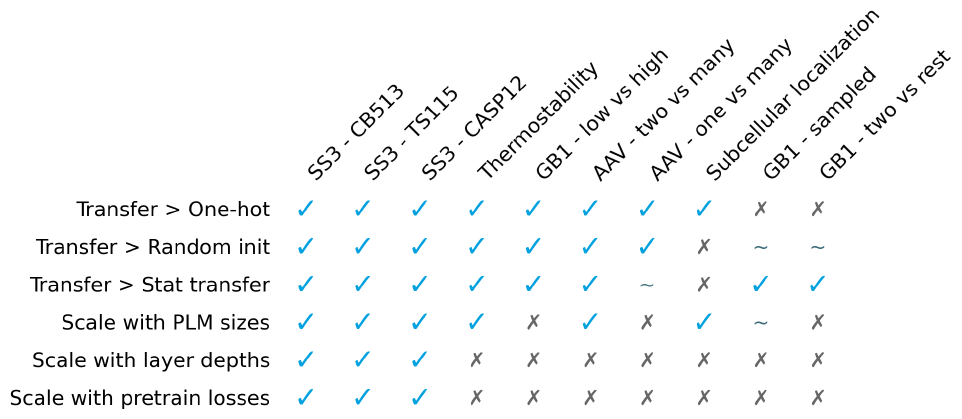
Downstream task result summary. 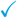 indicates true, *∼* indicates true for only one architecture, and 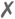 indicates false.

### 5.1 Structure Prediction Benefits from Transfer Learning Because It Is Well-Aligned with MLM pretraining

For all three residue-level secondary structure prediction tasks, Fig. 3a and Table A3 show that PLM embeddings out-perform the one-hot baseline as well as the random init and stat transfer ablations, demonstrating that transfer learning improves secondary structure prediction performance and that the improvement is not due to the inductive biases or weight statistics of the models. Secondary structure prediction performance improves when transferring deeper PLM features (Fig. 3b), indicating that more complex features from later layers continue to improve performance. Furthermore, transfer learning with features from larger models and from later in pretraining improve secondary structure prediction (Fig. 3a and 3c), as previously observed by Rives et al. (2021), Elnaggar et al. (2022), and Yang et al. (2022). We therefore conclude that MLM pretraining is well-aligned to structure prediction, allowing PLM features to be reused when predicting secondary structure from sequence.

### 5.2 Many Tasks Benefit from Transfer Learning Despite Lack of Alignment with MLM Pretraining

Next, we observe a cluster of four downstream tasks (thermostability, GB1 – low vs high, AAV – two vs many, and AAV – one vs many) where transfer learning improves performance over baselines even though the tasks do not align well with the pretraining task (Fig. 4). For these tasks, transfer learning improves performance over both the random init and stat transfer ablations, indicating that transfer learning confers at least some benefit over the inductive biases, parameterization, or weight statistics of the models alone (with the exception of the AAV – one vs many task, where weight statistics may still explain transfer learning performance) (Fig. 4a and Table A4). However, for all of these tasks, downstream task performance does not improve as features from deeper layers are transferred (Fig. 4b) or as the PLMs improve their pretraining loss over checkpoints (Fig. 4c), suggesting that these tasks may rely upon low-level features learned early in pretraining.

To supplement our quantitative cut-offs for alignment, we qualitatively assess trends in layer-by-layer performance across tasks. We observe that for all tasks where transfer learning improves performance over the baselines (including the secondary structure prediction tasks), the largest gains in performance occur in the first 3-5 layers of both the ESM and CARP models, across model sizes (Fig. 3b and 4b). However, unlike the secondary structure prediction tasks, which continue to improve in performance past this initial peak, improvement on the downstream tasks in this cluster generally plateaus (e.g. for the AAV – two vs many task), supporting our interpretation that features contributing to these tasks are already present within the first few layers of pretrained PLMs.

**Figure 3.**
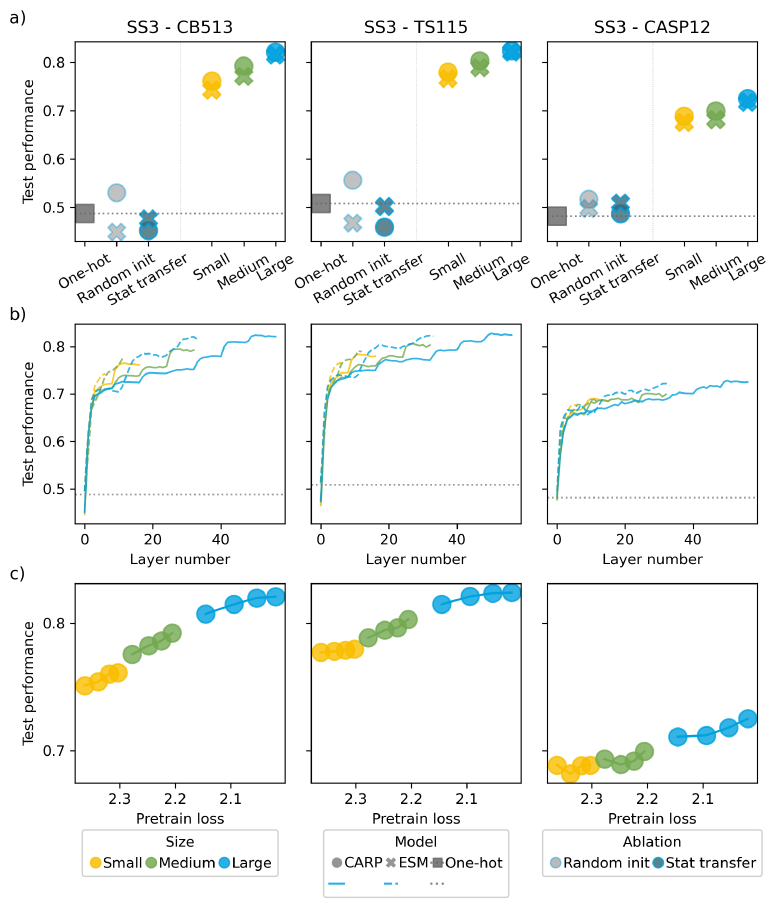
Results for secondary structure prediction. a) Performance on downstream tasks when transferring the final layer representation from various sizes of ESM and CARP compared to baselines and ablations. b) Downstream task performance by depth of layer transferred. c) Downstream task performance by pretraining loss. Each dot is a model checkpoint. For all subplots, downstream task test performance is quantified using accuracy.

Interestingly, although none of these tasks scale with model depth or pretraining loss, two downstream tasks (thermostability and AAV – two vs many) scale with PLM size (Fig. 4a). We reasoned that while our random init ablation rules out that improvements in downstream task performance is entirely due to parameterization, parameterization may still partially contribute to performance independently of feature reuse. To test this, we additionally evaluated the performance of small and medium randomly initialized models. Indeed, we observe that both types of randomly initialized models scale in performance with model size for both tasks, and in similar proportions to the improvements for the pretrained models (Table A4). Together, this suggests that observing that downstream task performance scales with model size alone is not sufficient to conclude that pretraining and downstream tasks are aligned, and that demonstrating scaling across other axes (such as model depth and checkpoints in training, as we propose here) is necessary.

**Figure 4.**
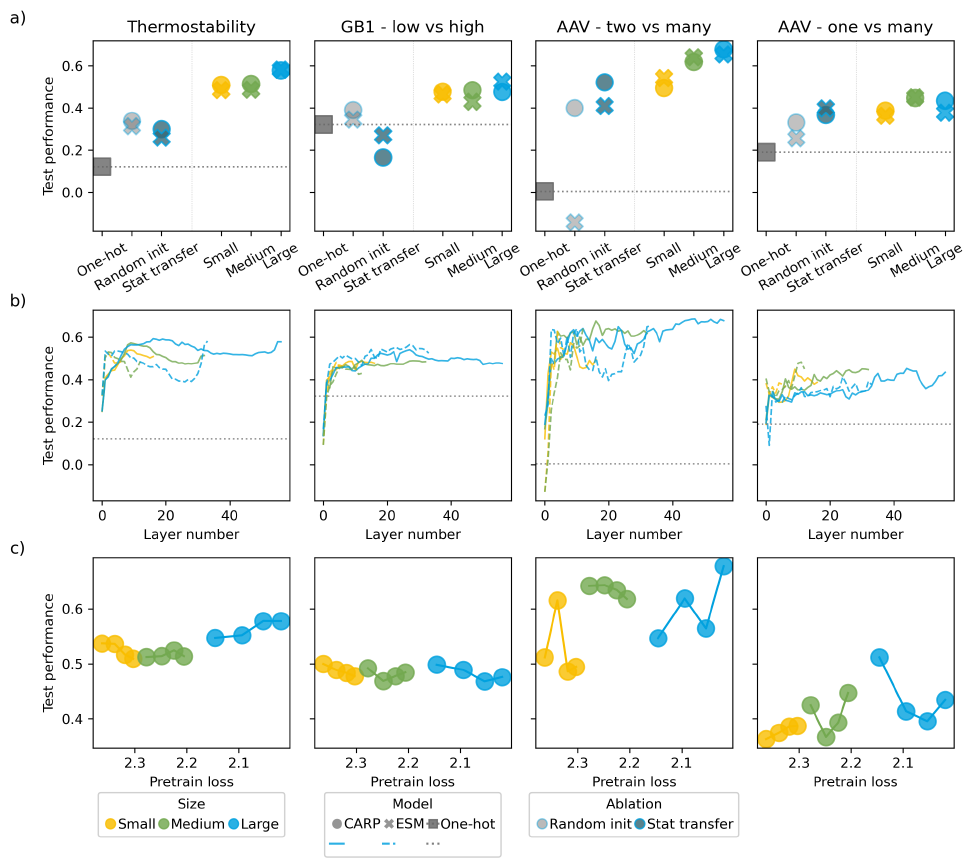
Results for tasks where transfer learning improves downstream task performance, but the pretrain and downstream tasks are not aligned. a) Performance on downstream tasks when transferring the final layer representation from various sizes of ESM and CARP compared to baselines and ablations. b) Downstream task performance by depth of layer transferred. c) Downstream task performance by pretraining loss. Each dot is a model checkpoint. For all subplots, downstream task test performance is quantified using Spearman’s rank correlation.

### 5.3 Some Tasks Do Not Benefit from MLM Pretraining

Finally, we observe a cluster of three downstream tasks (subcellular localization, GB1 – sampled, and GB1 – two vs rest) where pretraining does not improve transfer learning performance (Fig. 5). For subcellular localization, although transfer learning improves over a one-hot representation, pretrained models perform no better than randomly initialized models, suggesting that the improvement can be entirely attributed to inductive biases and parameterization. In contrast, the GB1 tasks in this cluster fail to outperform a one-hot representation by at least 10% (Fig. 5a and Table A5). We hypothesize the GB1 splits in this cluster are either too trivial, or too challenging for any representation. GB1 – sampled is an in-distribution task with a relatively large training set, and all models and baselines perform well. Meanwhile, GB1 – two vs rest is a challenging out-of-distribution split.

**Figure 5.**
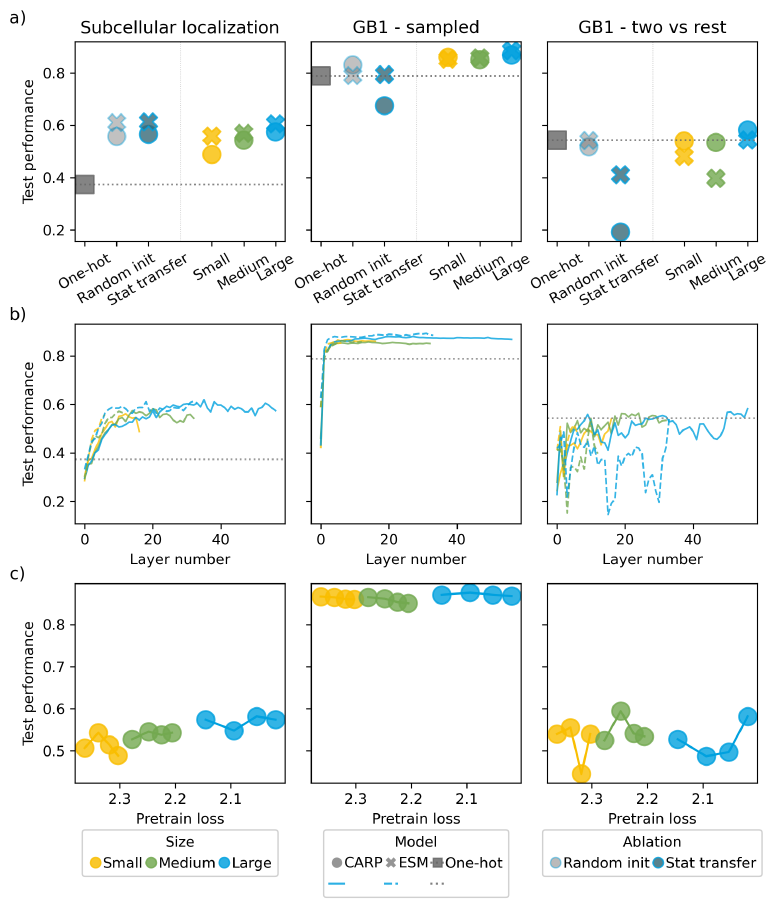
Results for task where pretraining does not improve downstream task performance. a) Performance on downstream tasks when transferring the final layer representation from various sizes of ESM and CARP compared to baselines and ablations. b) Downstream task performance by depth of layer transferred. c) Downstream task performance by pretraining loss. Each dot is a model checkpoint. For subcellular localization, the downstream classification task performance is quantified using accuracy. For other tasks, the downstream regression task performance is quantified using Spearman’s rank correlation.

Intriguingly, our stats transfer ablation decreases performance for all GB1 tasks, including the GB1 – low vs high task in the previous section, compared to the one-hot and random initialization baselines (Fig. 4b and 5b; Tables A4 and A5). We hypothesize that this is because the GB1 dataset is a highly local task, depending on finding interactions between just four mutated positions in a sequence.

## 6. Discussion

In this work, we systematically evaluate the mechanisms via which transfer learning from large pretrained protein language models improve performance on downstream protein function and structure prediction tasks. While most down-stream tasks benefit from transfer learning, structure prediction is the only task where we observe pretrain-downstream alignment. Our results are consistent with previous studies that show MLM pretraining imparts information about protein structure. Previous work has shown that the deepest attention matrices in pretrained PLMs recapitulate contact maps (Vig et al., 2020; Rao et al., 2021), and that it is possible to extract contact maps by perturbing the inputs to PLMs (Zhang et al., 2024). More recent work has argued that PLM representations contain similar co-evolution information to multiple sequence alignments (Chowdhury et al., 2022; Lin et al., 2023; Wu et al., 2022).

Our primary contribution is showing that scaling pretraining does not improve performance on prediction tasks that are less reliant on coevolutionary patterns, and that outperforming the one-hot and randomly initialized baselines does not imply that downstream task performance will scale with pretraining performance. Our results are most pronounced on the protein engineering datasets, where the sequence variation is introduced artificially. The engineering tasks in FLIP measure the effect of local sequence variation on the native function. We expect coevolutionary patterns learned during MLM pretraining to be even less helpful when designing proteins for non-natural functions.

### Limitations

There are factors known to impact transfer that we could not test for PLMs due to a lack of public models or to computational expense. First, pretraining dataset is important, both in terms of distance between the pretraining and downstream task data domains (Cherti & Jitsev, 2022) and data size (Abnar et al., 2022). PLMs pretrain on large databases of natural sequences. In principle, this means that some downstream tasks may be out-of-distribution (e.g. those involving artificial variation or non-natural function), or subject to biases in data collection (e.g. taxonomies less-represented in UniProt (Consortium, 2019)). Previous studies have shown differences in pretraining performance by taxonomy (Almagro Armenteros et al., 2020). However, Meier et al. (2021) trained versions of ESM on UniRef100 instead of UniRef50, and Dallago et al. (2021) show that they perform very similarly on function prediction tasks. Moreover, subsampling pretraining sequence datasets has not been explored beyond downsampling redundant sequences, making the impact of data difficult to evaluate for pretrained PLMs.

Second, while a variety of other pretraining tasks have been proposed for protein transfer learning, and different pretraining tasks could potentially learn different aspects of protein biology, we remain uncertain if they will result in significant differences from MLMs. Many pretraining tasks still aim to reconstruct natural sequences (He et al., 2021; Notin et al., 2022; Tan et al., 2023; Ma et al., 2023) and so are also likely to primarily learn coevolutionary patterns. Other tasks use structure as an additional input or target, but they generally make only modest improvements on function prediction tasks (Mansoor et al., 2021; Wang et al., 2022; Yang et al., 2023; Su et al., 2023). Supporting the assertion that learning to predict structure may not improve function prediction, Hu et al. (2022) show that transfer learning using the AlphaFold2 (Jumper et al., 2021) structure module is less effective for function prediction than transferring PLMs. Finally, Brandes et al. (2022) and Xu et al. (2023) reconstruct both sequence and functional annotations but also find that downstream performance does not always scale with pretraining time.

Finally, we only test linear probes on mean pooled representations to limit computational cost, but previous work shows that for many tasks finetuning the PLM end-to-end outperforms a linear probe or training a small neural network on top of the frozen pretrained weights (Dallago et al., 2021; Yang et al., 2022), and that mean-pooling is rarely optimal (Detlefsen et al., 2022; Goldman et al., 2022). In computer vision, models trained on different datasets (Cherti & Jitsev, 2022) and pretraining tasks (Grigg et al., 2021) exhibit different finetuning dynamics, and there is some evidence for this in proteins as well (Detlefsen et al., 2022).

### Implications for future work

First, our work emphasizes the need for improved evaluation standards for PLMs. We show that checking for improved performance over baselines may overestimate the generality of PLMs across applications in protein biology, as it does not rule out that improvement may be due to alternate hypotheses that do not scale. However, most current works rely on comparisons to baselines to argue that PLMs are widely applicable, and to the extent scaling has been studied, most only use scaling on structure prediction accuracy alone to justify training larger models (Rives et al., 2021; Elnaggar et al., 2022; Lin et al., 2023; Chen et al., 2024). Future PLM evaluation should therefore assess scaling on diverse downstream function prediction and engineering tasks, and not just structure alone, to validate the generality of models.

Second, synthesizing our empirical results with how the current landscape of protein sequence pretraining tasks primarily align with structure prediction, our work points to a need for new pretraining tasks. For many downstream tasks, the lack of alignment prevents transfer learning from taking full advantage of the pretrained model, as features from deep in the PLM perform no better than features from early layers in the PLM. Likewise, for these tasks, simply scaling to larger PLMs trained for more steps on more data will not improve performance. Our study suggests that the field needs to explore diversified pretraining strategies instead of further scaling existing strategies in order to reach aspects of protein biology that are not well-served by PLMs.

## 7. Impact Statement

This paper exposes current limitations in protein language models, which are routinely used in protein engineering and bioinformatics. Protein language models scale from year to year, with current models reaching hundreds of billions of parameters (Chen et al., 2024). By showing that current model pretraining paradigms fail to confer benefits on many aspects of protein biology, we caution against uncritically investing compute resources into scaling these models, which we hope will translate to impact through reduced carbon emissions. Additionally, by showing what kinds of tasks protein language models currently fail to scale on, we hope our work leads to the development of pretrained models that improve bioinformatics and protein design predictions beyond those currently well-served by protein language models. If so, we anticipate both positive and negative impacts from an expanded capability to design new proteins.

## 8. Acknowledgements and Funding

The authors thank Sarah Alamdari, Frances Arnold, Sébastien Bubeck, Kianoush Falahkheirkhah, Nicolo Fusi, Kadina Johnston, Alexander Lin, David Alvarez Melis, Ariane Mora, Ivan Dario Jimenez Rodriguez, Nitya Thakkar, Amy Wang, Sean Whitzell, Bruce Wittmann, Kevin Wu, and Wenjun Wu for ideas and discussions that helped us improve our work. F.Z.L was partially supported by the National Science Foundation Graduate Research Fellowship and Amazon AI4Science Fellowship at Caltech. This material is based upon work supported by the U.S. Department of Energy, Office of Science, Office of Basic Energy Sciences, under Award Number DE-SC0022218. This report was prepared as an account of work sponsored by an agency of the United States Government. Neither the United States Government nor any agency thereof, nor any of their employees, makes any warranty, express or implied, or assumes any legal liability or responsibility for the accuracy, completeness, or usefulness of any information, apparatus, product, or process disclosed, or represents that its use would not infringe privately owned rights. Reference herein to any specific commercial product, process, or service by trade name, trademark, manufacturer, or otherwise does not necessarily constitute or imply its endorsement, recommendation, or favoring by the United States Government or any agency thereof. The views and opinions of authors expressed herein do not necessarily state or reflect those of the United States Government or any agency thereof.

## A. Additional tables and figures

**Table A1.** Downstream functional and structural tasks.

| Task | Task type | Split type | $n$ Train | $n$ Val | $n$ Test | Model type ($n$ classes) |
| --- | --- | --- | --- | --- | --- | --- |
| SS3 – CB513 | Residue-level secondary structure | Minimal homology | 8678 | 2170 | 513 | PyTorch linear classifier (3) |
| SS3 – TS115 | Residue-level secondary structure | Released in 2016, from 43 to 1085 residues | 8678 | 2170 | 115 | PyTorch linear classifier (3) |
| SS3 – CASP12 | Residue-level secondary structure | CASP12 targets, mostly more than 400 residues | 8678 | 2170 | 21 | PyTorch linear classifier (3) |
| Thermostability | Global property | In-distribution | 22335 | 2482 | 3134 | Scikit-learn ridge regression |
| Subcellular localization (scl) | Global property | In-distribution | 9503 | 1678 | 385 | PyTorch linear classifier (10) |
| GB1 – sampled | Local property | In-distribution | 6289 | 699 | 1745 | Scikit-learn ridge regression |
| GB1 – low vs high | Local property | Out-of-distribution | 4580 | 509 | 3644 | Scikit-learn ridge regression |
| GB1 – two vs rest | Local property | Out-of-distribution (fewer training samples) | 381 | 43 | 8309 | Scikit-learn ridge regression |
| AAV – two vs many | Local property | Out-of-distribution | 28626 | 3181 | 50776 | Scikit-learn ridge regression |
| AAV – one vs many | Local property | Out-of-distribution (fewer training samples) | 1053 | 117 | 81413 | Scikit-learn ridge regression |

**Table A2.**
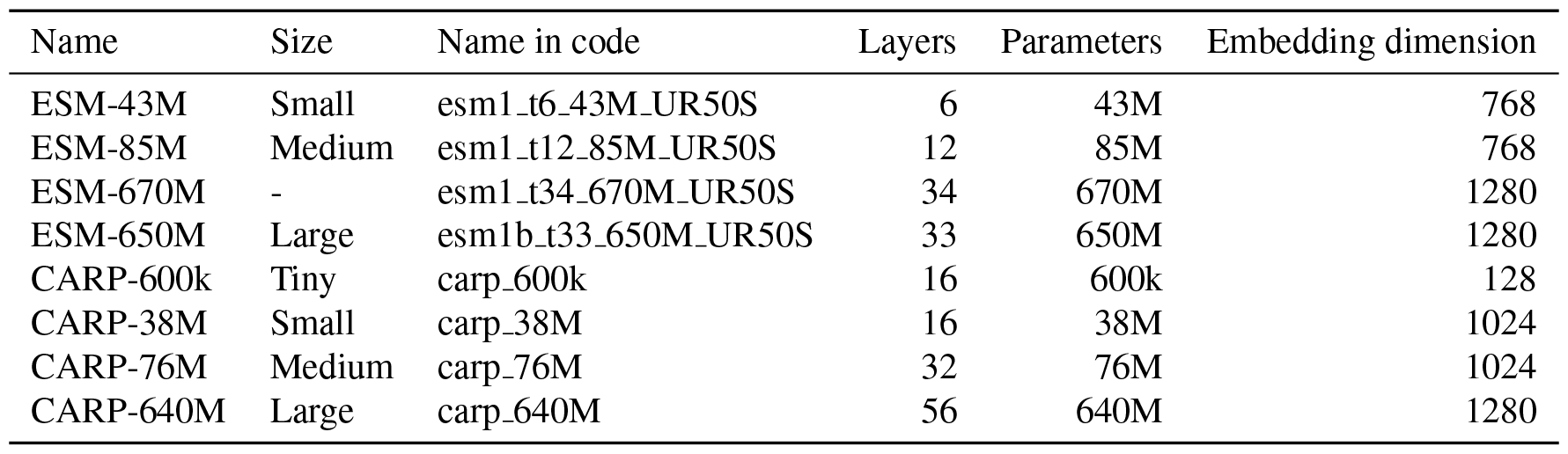
Pretrained models.

| Name | Size | Name in code | Layers | Parameters | Embedding dimension |
| --- | --- | --- | --- | --- | --- |
| ESM-43M | Small | esm1_t6_43M_UR50S | 6 | 43M | 768 |
| ESM-85M | Medium | esm1_t12_85M_UR50S | 12 | 85M | 768 |
| ESM-670M | - | esm1_t34_670M_UR50S | 34 | 670M | 1280 |
| ESM-650M | Large | esm1b_t33_650M_UR50S | 33 | 650M | 1280 |
| CARP-600k | Tiny | carp_600k | 16 | 600k | 128 |
| CARP-38M | Small | carp_38M | 16 | 38M | 1024 |
| CARP-76M | Medium | carp_76M | 32 | 76M | 1024 |
| CARP-640M | Large | carp_640M | 56 | 640M | 1280 |

**Table A3.**
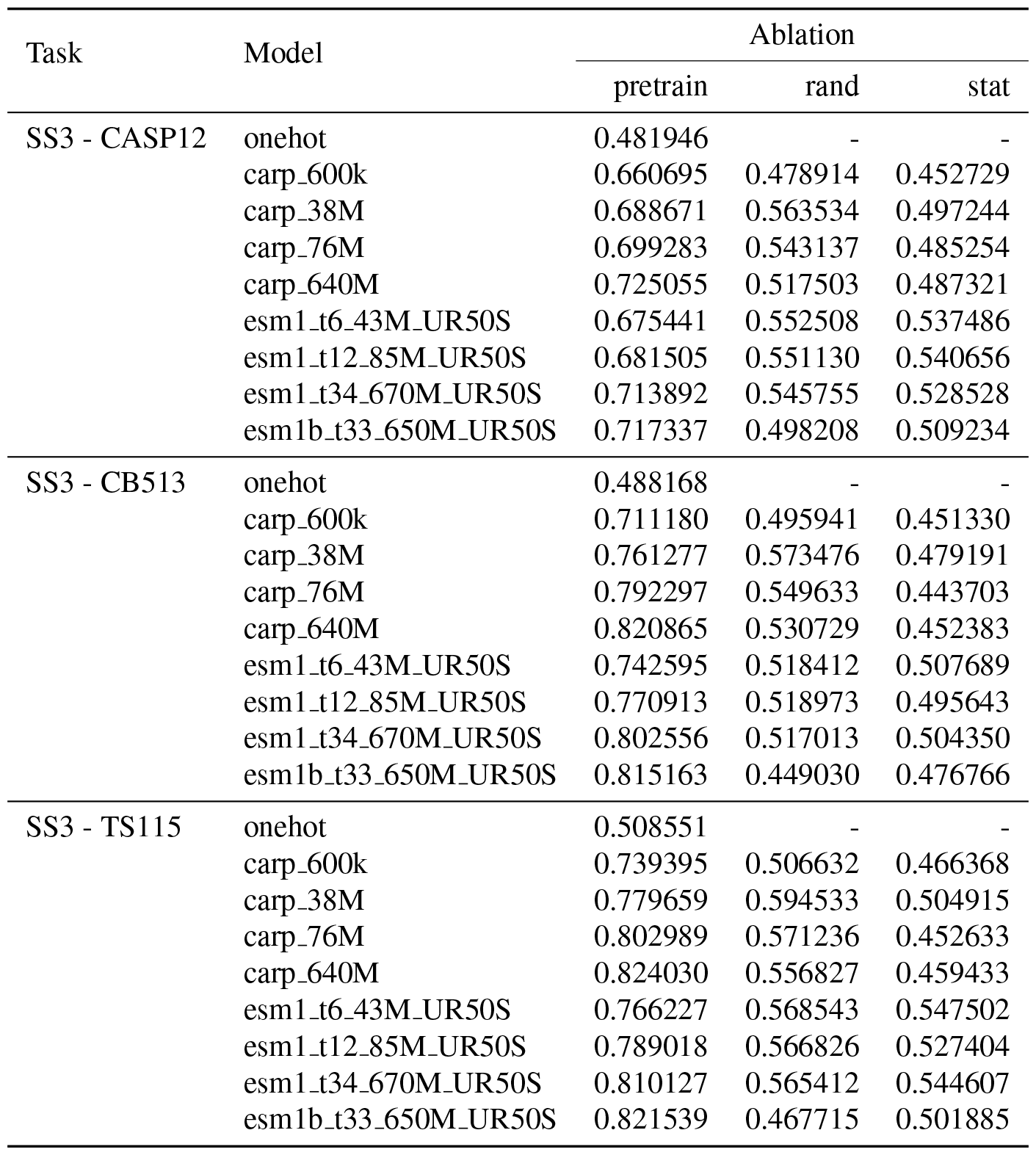
Last layer transfer learning performance for tasks that are aligned with MLM pretraining. Values are accuracy.

| Task | Model | Ablation |  |  |
| --- | --- | --- | --- | --- |
|  |  | pretrain | rand | stat |
| SS3 - CASP12 | onehot | 0.481946 | - | - |
|  | carp_600k | 0.660695 | 0.478914 | 0.452729 |
|  | carp_38M | 0.688671 | 0.563534 | 0.497244 |
|  | carp_76M | 0.699283 | 0.543137 | 0.485254 |
|  | carp_640M | 0.725055 | 0.517503 | 0.487321 |
|  | esm1_t6_43M_UR50S | 0.675441 | 0.552508 | 0.537486 |
|  | esm1_t12_85M_UR50S | 0.681505 | 0.551130 | 0.540656 |
|  | esm1_t34_670M_UR50S | 0.713892 | 0.545755 | 0.528528 |
|  | esm1b_t33_650M_UR50S | 0.717337 | 0.498208 | 0.509234 |
| SS3 - CB513 | onehot | 0.488168 | - | - |
|  | carp_600k | 0.711180 | 0.495941 | 0.451330 |
|  | carp_38M | 0.761277 | 0.573476 | 0.479191 |
|  | carp_76M | 0.792297 | 0.549633 | 0.443703 |
|  | carp_640M | 0.820865 | 0.530729 | 0.452383 |
|  | esm1_t6_43M_UR50S | 0.742595 | 0.518412 | 0.507689 |
|  | esm1_t12_85M_UR50S | 0.770913 | 0.518973 | 0.495643 |
|  | esm1_t34_670M_UR50S | 0.802556 | 0.517013 | 0.504350 |
|  | esm1b_t33_650M_UR50S | 0.815163 | 0.449030 | 0.476766 |
| SS3 - TS115 | onehot | 0.508551 | - | - |
|  | carp_600k | 0.739395 | 0.506632 | 0.466368 |
|  | carp_38M | 0.779659 | 0.594533 | 0.504915 |
|  | carp_76M | 0.802989 | 0.571236 | 0.452633 |
|  | carp_640M | 0.824030 | 0.556827 | 0.459433 |
|  | esm1_t6_43M_UR50S | 0.766227 | 0.568543 | 0.547502 |
|  | esm1_t12_85M_UR50S | 0.789018 | 0.566826 | 0.527404 |
|  | esm1_t34_670M_UR50S | 0.810127 | 0.565412 | 0.544607 |
|  | esm1b_t33_650M_UR50S | 0.821539 | 0.467715 | 0.501885 |

**Table A4.**
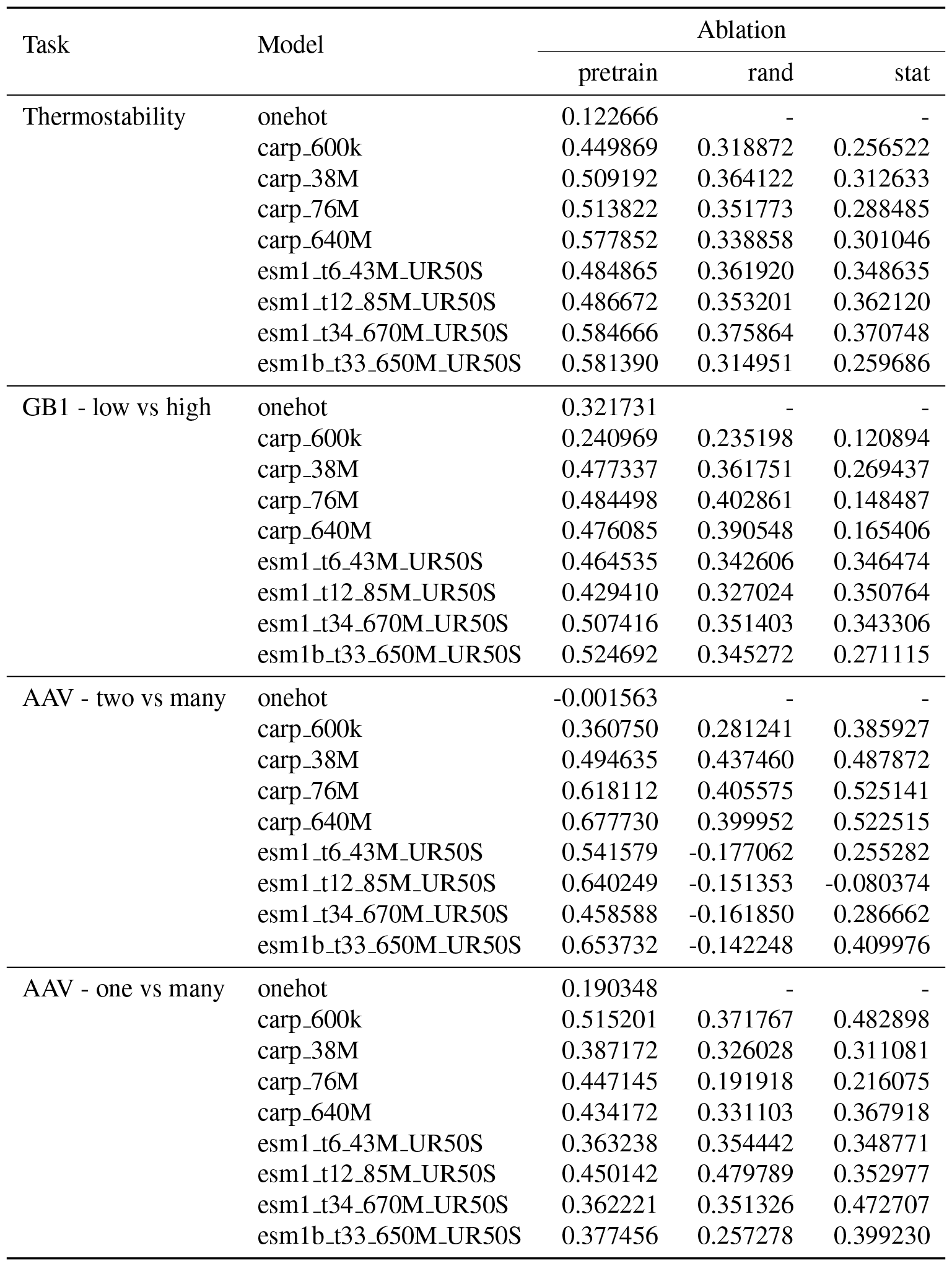
Last layer transfer learning performance for tasks where transfer learning improves performance but the pretrain and downstream tasks are not aligned. Values are Spearman rank correlation.

| Task | Model | Ablation |  |  |
| --- | --- | --- | --- | --- |
|  |  | pretrain | rand | stat |
| Thermostability | onehot | 0.122666 | - | - |
|  | carp_600k | 0.449869 | 0.318872 | 0.256522 |
|  | carp_38M | 0.509192 | 0.364122 | 0.312633 |
|  | carp_76M | 0.513822 | 0.351773 | 0.288485 |
|  | carp_640M | 0.577852 | 0.338858 | 0.301046 |
|  | esm1_t6_43M_UR50S | 0.484865 | 0.361920 | 0.348635 |
|  | esm1_t12_85M_UR50S | 0.486672 | 0.353201 | 0.362120 |
|  | esm1_t34_670M_UR50S | 0.584666 | 0.375864 | 0.370748 |
|  | esm1b_t33_650M_UR50S | 0.581390 | 0.314951 | 0.259686 |
| GB1 - low vs high | onehot | 0.321731 | - | - |
|  | carp_600k | 0.240969 | 0.235198 | 0.120894 |
|  | carp_38M | 0.477337 | 0.361751 | 0.269437 |
|  | carp_76M | 0.484498 | 0.402861 | 0.148487 |
|  | carp_640M | 0.476085 | 0.390548 | 0.165406 |
|  | esm1_t6_43M_UR50S | 0.464535 | 0.342606 | 0.346474 |
|  | esm1_t12_85M_UR50S | 0.429410 | 0.327024 | 0.350764 |
|  | esm1_t34_670M_UR50S | 0.507416 | 0.351403 | 0.343306 |
|  | esm1b_t33_650M_UR50S | 0.524692 | 0.345272 | 0.271115 |
| AAV - two vs many | onehot | -0.001563 | - | - |
|  | carp_600k | 0.360750 | 0.281241 | 0.385927 |
|  | carp_38M | 0.494635 | 0.437460 | 0.487872 |
|  | carp_76M | 0.618112 | 0.405575 | 0.525141 |
|  | carp_640M | 0.677730 | 0.399952 | 0.522515 |
|  | esm1_t6_43M_UR50S | 0.541579 | -0.177062 | 0.255282 |
|  | esm1_t12_85M_UR50S | 0.640249 | -0.151353 | -0.080374 |
|  | esm1_t34_670M_UR50S | 0.458588 | -0.161850 | 0.286662 |
|  | esm1b_t33_650M_UR50S | 0.653732 | -0.142248 | 0.409976 |
| AAV - one vs many | onehot | 0.190348 | - | - |
|  | carp_600k | 0.515201 | 0.371767 | 0.482898 |
|  | carp_38M | 0.387172 | 0.326028 | 0.311081 |
|  | carp_76M | 0.447145 | 0.191918 | 0.216075 |
|  | carp_640M | 0.434172 | 0.331103 | 0.367918 |
|  | esm1_t6_43M_UR50S | 0.363238 | 0.354442 | 0.348771 |
|  | esm1_t12_85M_UR50S | 0.450142 | 0.479789 | 0.352977 |
|  | esm1_t34_670M_UR50S | 0.362221 | 0.351326 | 0.472707 |
|  | esm1b_t33_650M_UR50S | 0.377456 | 0.257278 | 0.399230 |

**Table A5.** Last layer transfer learning performance for tasks where transfer learning does not improve performance. Values are Spearman rank correlation for the GB1 tasks and accuracy for subcellular localization.

| Task | Model | Ablation |  |  |
| --- | --- | --- | --- | --- |
|  |  | pretrain | rand | stat |
| GB1 - two vs rest | onehot | 0.542840 | - | - |
|  | carp_600k | 0.562338 | 0.515097 | 0.175361 |
|  | carp_38M | 0.539823 | 0.420235 | 0.413975 |
|  | carp_76M | 0.534144 | 0.444437 | 0.293423 |
|  | carp_640M | 0.581670 | 0.518491 | 0.191882 |
|  | esm1_t6_43M_UR50S | 0.480617 | 0.609323 | 0.629515 |
|  | esm1_t12_85M_UR50S | 0.396941 | 0.585249 | 0.597770 |
|  | esm1_t34_670M_UR50S | 0.506492 | 0.588215 | 0.566406 |
|  | esm1b_t33_650M_UR50S | 0.543660 | 0.542069 | 0.411151 |
| GB1 - sampled | onehot | 0.788505 | - | - |
|  | carp_600k | 0.785568 | 0.752780 | 0.655448 |
|  | carp_38M | 0.859780 | 0.825859 | 0.776026 |
|  | carp_76M | 0.850627 | 0.831454 | 0.715282 |
|  | carp_640M | 0.867894 | 0.831186 | 0.675240 |
|  | esm1_t6_43M_UR50S | 0.852435 | 0.797663 | 0.813344 |
|  | esm1_t12_85M_UR50S | 0.857460 | 0.793028 | 0.813187 |
|  | esm1_t34_670M_UR50S | 0.865990 | 0.800310 | 0.816400 |
|  | esm1b_t33_650M_UR50S | 0.884644 | 0.789944 | 0.793893 |
| Subcellular localization | onehot | 0.374026 | - | - |
|  | carp_600k | 0.449351 | 0.433766 | 0.431169 |
|  | carp_38M | 0.488312 | 0.548052 | 0.501299 |
|  | carp_76M | 0.542857 | 0.537662 | 0.550649 |
|  | carp_640M | 0.574026 | 0.558442 | 0.566234 |
|  | esm1_t6_43M_UR50S | 0.558442 | 0.555844 | 0.576623 |
|  | esm1_t12_85M_UR50S | 0.568831 | 0.571429 | 0.568831 |
|  | esm1_t34_670M_UR50S | 0.615584 | 0.610390 | 0.610390 |
|  | esm1b_t33_650M_UR50S | 0.605195 | 0.610390 | 0.612987 |

**Table A6.** Spearman’s rank correlation (*ρ*) between downstream task performance and layer depth.

| Task | CARP-640M |  | ESM-650M |  |
| --- | --- | --- | --- | --- |
| | $\rho$ | P | $\rho$ | P |
| SS3 - CB513 | 0.989 | $5.850 \times 10^{-47}$ | 0.954 | $2.511 \times 10^{-18}$ |
| SS3 - TS115 | 0.985 | $6.109 \times 10^{-44}$ | 0.953 | $4.197 \times 10^{-18}$ |
| SS3 - CASP12 | 0.991 | $2.136 \times 10^{-49}$ | 0.957 | $1.063 \times 10^{-18}$ |
| Thermostability | -0.090 | $5.042 \times 10^{-1}$ | -0.432 | $1.068 \times 10^{-2}$ |
| GB1 - low vs high | 0.289 | $2.922 \times 10^{-2}$ | 0.814 | $4.817 \times 10^{-9}$ |
| AAV - two vs many | 0.809 | $2.583 \times 10^{-14}$ | 0.014 | $9.378 \times 10^{-1}$ |
| AAV - one vs many | 0.853 | $3.966 \times 10^{-17}$ | 0.757 | $2.160 \times 10^{-7}$ |
| Subcellular localization | 0.694 | $2.085 \times 10^{-9}$ | 0.621 | $8.896 \times 10^{-5}$ |
| GB1 - sampled | 0.325 | $1.362 \times 10^{-2}$ | 0.850 | $1.961 \times 10^{-10}$ |
| GB1 - two vs rest | 0.436 | $7.023 \times 10^{-4}$ | -0.267 | $1.266 \times 10^{-1}$ |

**Table A7.** Pretrained CARP checkpoints.

| Name | Fraction | Loss | Accuracy | Step |
| --- | --- | --- | --- | --- |
| carp_600k | 1 | 2.505 | 0.240 | $4.889 \times 10^5$ |
| carp_600k | 0.5 | 2.512 | 0.239 | $2.393 \times 10^5$ |
| carp_600k | 0.25 | 2.518 | 0.237 | $1.143 \times 10^5$ |
| carp_600k | 0.125 | 2.527 | 0.234 | $5.204 \times 10^4$ |
| carp_38M | 1 | 2.303 | 0.300 | $1.027 \times 10^6$ |
| carp_38M | 0.5 | 2.319 | 0.295 | $5.176 \times 10^5$ |
| carp_38M | 0.25 | 2.339 | 0.289 | $2.569 \times 10^5$ |
| carp_38M | 0.125 | 2.363 | 0.282 | $1.296 \times 10^5$ |
| carp_76M | 1 | 2.206 | 0.328 | $6.545 \times 10^5$ |
| carp_76M | 0.5 | 2.225 | 0.322 | $3.280 \times 10^5$ |
| carp_76M | 0.25 | 2.248 | 0.315 | $1.630 \times 10^5$ |
| carp_76M | 0.125 | 2.278 | 0.307 | $8.318 \times 10^4$ |
| carp_640M | 1 | 2.019 | 0.382 | $6.220 \times 10^5$ |
| carp_640M | 0.5 | 2.054 | 0.372 | $3.118 \times 10^5$ |
| carp_640M | 0.25 | 2.094 | 0.360 | $1.547 \times 10^5$ |
| carp_640M | 0.125 | 2.146 | 0.345 | $7.881 \times 10^4$ |

**Table A8.** Spearman’s rank correlation (*ρ*) between downstream task performance and CARP pretrain loss.

| Task | $\rho$ | p |
| --- | --- | --- |
| SS3 - CB513 | 1.000 | 0.000 |
| SS3 - TS115 | 1.000 | 0.000 |
| SS3 - CASP12 | 0.949 | $2.000 \times 10^{-6}$ |
| Thermostability | 0.552 | $6.251 \times 10^{-2}$ |
| GB1 - low vs high | -0.392 | $2.081 \times 10^{-1}$ |
| AAV - two vs many | 0.483 | $1.121 \times 10^{-1}$ |
| AAV - one vs many | 0.727 | $7.355 \times 10^{-3}$ |
| Subcellular localization | 0.832 | $7.980 \times 10^{-4}$ |
| GB1 - sampled | 0.441 | $1.517 \times 10^{-1}$ |
| GB1 - two vs rest | -0.084 | $7.954 \times 10^{-1}$ |

