## Supplemental data and code for "Feature Reuse and Scaling: Understanding Transfer Learning with Protein Language Models": env-protein-transfer.yml

```
# Install or update using
# conda env update --file env-protein-transfer.yml --prune

name: protran

channels:
- pytorch
- conda-forge
- bioconda
- anaconda

dependencies:
- python=3.9.10
- anndata=0.7.5
- biopython=1.79
- bokeh=2.3.3
- colorcet=2.0.2
# - cudatoolkit=11.7
- cudatoolkit=11.6
- firefox=99.0
- geckodriver=0.30.0
- holoviews=1.14.6
- ipywidgets=7.6.5
- jinja2=3.0.3
- jupyterlab=3.3.3
- matplotlib=3.5.1
- numpy=1.22.3
- pandas=1.4.2
- panel=0.12.1
- pip=22.0.4
# - pytorch=1.11.0
- pytorch==1.12.1
- pyviz_comms=2.1.0
- scikit-learn=1.0.2
- scipy=1.8.0
- selenium=3.141.0
# - torchvision==0.12.0
- tqdm=4.64.0

- pip:
  - iqplot==0.3.0
#  - lmdb==0.97
  - scanpy==1.7.1
  - umap-learn==0.5.1
  - blackcellmagic==0.0.3
  - fair-esm
  - sequence-models
  - wget
```
