## Supplemental data and code for "Feature Reuse and Scaling: Understanding Transfer Learning with Protein Language Models": execute_gen_emb_carp.sh

```
# queue for generating and sorting all embs

# export CUDA_VISIBLE_DEVICES=""
# export CUDA_VISIBLE_DEVICES=1

### GB1 ###

# python run_pregen_emb.py --dataset_path="data/proeng/gb1/sampled.csv" --encoder_name="carp_600k" --checkpoint=1 --flatten_emb="mean" --seq_end_idx=56 --reset_param=True
# python run_pregen_emb.py --dataset_path="data/proeng/gb1/sampled.csv" --encoder_name="carp_600k" --checkpoint=1 --flatten_emb="mean" --seq_end_idx=56 --resample_param=True
# python run_pregen_emb.py --dataset_path="data/proeng/gb1/sampled.csv" --encoder_name="carp_600k" --checkpoint=1 --flatten_emb="mean" --seq_end_idx=56
# python run_pregen_emb.py --dataset_path="data/proeng/gb1/sampled.csv" --encoder_name="carp_600k" --checkpoint=0.5 --flatten_emb="mean" --seq_end_idx=56
# python run_pregen_emb.py --dataset_path="data/proeng/gb1/sampled.csv" --encoder_name="carp_600k" --checkpoint=0.25 --flatten_emb="mean" --seq_end_idx=56
# python run_pregen_emb.py --dataset_path="data/proeng/gb1/sampled.csv" --encoder_name="carp_600k" --checkpoint=0.125 --flatten_emb="mean" --seq_end_idx=56

# python run_pregen_emb.py --dataset_path="data/proeng/gb1/sampled.csv" --encoder_name="carp_38M" --checkpoint=1 --flatten_emb="mean" --seq_end_idx=56 --reset_param=True
# python run_pregen_emb.py --dataset_path="data/proeng/gb1/sampled.csv" --encoder_name="carp_38M" --checkpoint=1 --flatten_emb="mean" --seq_end_idx=56 --resample_param=True
# python run_pregen_emb.py --dataset_path="data/proeng/gb1/sampled.csv" --encoder_name="carp_38M" --checkpoint=1 --flatten_emb="mean" --seq_end_idx=56
# python run_pregen_emb.py --dataset_path="data/proeng/gb1/sampled.csv" --encoder_name="carp_38M" --checkpoint=0.5 --flatten_emb="mean" --seq_end_idx=56
# python run_pregen_emb.py --dataset_path="data/proeng/gb1/sampled.csv" --encoder_name="carp_38M" --checkpoint=0.25 --flatten_emb="mean" --seq_end_idx=56
# python run_pregen_emb.py --dataset_path="data/proeng/gb1/sampled.csv" --encoder_name="carp_38M" --checkpoint=0.125 --flatten_emb="mean" --seq_end_idx=56

# python run_pregen_emb.py --dataset_path="data/proeng/gb1/sampled.csv" --encoder_name="carp_76M" --checkpoint=1 --flatten_emb="mean" --seq_end_idx=56 --reset_param=True
# python run_pregen_emb.py --dataset_path="data/proeng/gb1/sampled.csv" --encoder_name="carp_76M" --checkpoint=1 --flatten_emb="mean" --seq_end_idx=56 --resample_param=True
# python run_pregen_emb.py --dataset_path="data/proeng/gb1/sampled.csv" --encoder_name="carp_76M" --checkpoint=1 --flatten_emb="mean" --seq_end_idx=56
# python run_pregen_emb.py --dataset_path="data/proeng/gb1/sampled.csv" --encoder_name="carp_76M" --checkpoint=0.5 --flatten_emb="mean" --seq_end_idx=56
# python run_pregen_emb.py --dataset_path="data/proeng/gb1/sampled.csv" --encoder_name="carp_76M" --checkpoint=0.25 --flatten_emb="mean" --seq_end_idx=56
# python run_pregen_emb.py --dataset_path="data/proeng/gb1/sampled.csv" --encoder_name="carp_76M" --checkpoint=0.125 --flatten_emb="mean" --seq_end_idx=56

# python run_pregen_emb.py --dataset_path="data/proeng/gb1/sampled.csv" --encoder_name="carp_640M" --checkpoint=1 --flatten_emb="mean" --seq_end_idx=56 --reset_param=True
# python run_pregen_emb.py --dataset_path="data/proeng/gb1/sampled.csv" --encoder_name="carp_640M" --checkpoint=1 --flatten_emb="mean" --seq_end_idx=56 --resample_param=True
# python run_pregen_emb.py --dataset_path="data/proeng/gb1/sampled.csv" --encoder_name="carp_640M" --checkpoint=1 --flatten_emb="mean" --seq_end_idx=56
# python run_pregen_emb.py --dataset_path="data/proeng/gb1/sampled.csv" --encoder_name="carp_640M" --checkpoint=0.5 --flatten_emb="mean" --seq_end_idx=56
# python run_pregen_emb.py --dataset_path="data/proeng/gb1/sampled.csv" --encoder_name="carp_640M" --checkpoint=0.25 --flatten_emb="mean" --seq_end_idx=56
# python run_pregen_emb.py --dataset_path="data/proeng/gb1/sampled.csv" --encoder_name="carp_640M" --checkpoint=0.125 --flatten_emb="mean" --seq_end_idx=56


# python run_pregen_emb.py --dataset_path="data/proeng/gb1/low_vs_high.csv" --encoder_name="carp_600k" --checkpoint=1 --flatten_emb="mean" --seq_end_idx=56 --reset_param=True
# python run_pregen_emb.py --dataset_path="data/proeng/gb1/low_vs_high.csv" --encoder_name="carp_600k" --checkpoint=1 --flatten_emb="mean" --seq_end_idx=56 --resample_param=True
# python run_pregen_emb.py --dataset_path="data/proeng/gb1/low_vs_high.csv" --encoder_name="carp_600k" --checkpoint=1 --flatten_emb="mean" --seq_end_idx=56
# python run_pregen_emb.py --dataset_path="data/proeng/gb1/low_vs_high.csv" --encoder_name="carp_600k" --checkpoint=0.5 --flatten_emb="mean" --seq_end_idx=56
# python run_pregen_emb.py --dataset_path="data/proeng/gb1/low_vs_high.csv" --encoder_name="carp_600k" --checkpoint=0.25 --flatten_emb="mean" --seq_end_idx=56
# python run_pregen_emb.py --dataset_path="data/proeng/gb1/low_vs_high.csv" --encoder_name="carp_600k" --checkpoint=0.125 --flatten_emb="mean" --seq_end_idx=56

# python run_pregen_emb.py --dataset_path="data/proeng/gb1/low_vs_high.csv" --encoder_name="carp_38M" --checkpoint=1 --flatten_emb="mean" --seq_end_idx=56 --reset_param=True
# python run_pregen_emb.py --dataset_path="data/proeng/gb1/low_vs_high.csv" --encoder_name="carp_38M" --checkpoint=1 --flatten_emb="mean" --seq_end_idx=56 --resample_param=True
# python run_pregen_emb.py --dataset_path="data/proeng/gb1/low_vs_high.csv" --encoder_name="carp_38M" --checkpoint=1 --flatten_emb="mean" --seq_end_idx=56
# python run_pregen_emb.py --dataset_path="data/proeng/gb1/low_vs_high.csv" --encoder_name="carp_38M" --checkpoint=0.5 --flatten_emb="mean" --seq_end_idx=56
# python run_pregen_emb.py --dataset_path="data/proeng/gb1/low_vs_high.csv" --encoder_name="carp_38M" --checkpoint=0.25 --flatten_emb="mean" --seq_end_idx=56
# python run_pregen_emb.py --dataset_path="data/proeng/gb1/low_vs_high.csv" --encoder_name="carp_38M" --checkpoint=0.125 --flatten_emb="mean" --seq_end_idx=56

# python run_pregen_emb.py --dataset_path="data/proeng/gb1/low_vs_high.csv" --encoder_name="carp_76M" --checkpoint=1 --flatten_emb="mean" --seq_end_idx=56 --reset_param=True
# python run_pregen_emb.py --dataset_path="data/proeng/gb1/low_vs_high.csv" --encoder_name="carp_76M" --checkpoint=1 --flatten_emb="mean" --seq_end_idx=56 --resample_param=True
# python run_pregen_emb.py --dataset_path="data/proeng/gb1/low_vs_high.csv" --encoder_name="carp_76M" --checkpoint=1 --flatten_emb="mean" --seq_end_idx=56
# python run_pregen_emb.py --dataset_path="data/proeng/gb1/low_vs_high.csv" --encoder_name="carp_76M" --checkpoint=0.5 --flatten_emb="mean" --seq_end_idx=56
# python run_pregen_emb.py --dataset_path="data/proeng/gb1/low_vs_high.csv" --encoder_name="carp_76M" --checkpoint=0.25 --flatten_emb="mean" --seq_end_idx=56
# python run_pregen_emb.py --dataset_path="data/proeng/gb1/low_vs_high.csv" --encoder_name="carp_76M" --checkpoint=0.125 --flatten_emb="mean" --seq_end_idx=56

# python run_pregen_emb.py --dataset_path="data/proeng/gb1/low_vs_high.csv" --encoder_name="carp_640M" --checkpoint=1 --flatten_emb="mean" --seq_end_idx=56 --reset_param=True
# python run_pregen_emb.py --dataset_path="data/proeng/gb1/low_vs_high.csv" --encoder_name="carp_640M" --checkpoint=1 --flatten_emb="mean" --seq_end_idx=56 --resample_param=True
# python run_pregen_emb.py --dataset_path="data/proeng/gb1/low_vs_high.csv" --encoder_name="carp_640M" --checkpoint=1 --flatten_emb="mean" --seq_end_idx=56
# python run_pregen_emb.py --dataset_path="data/proeng/gb1/low_vs_high.csv" --encoder_name="carp_640M" --checkpoint=0.5 --flatten_emb="mean" --seq_end_idx=56
# python run_pregen_emb.py --dataset_path="data/proeng/gb1/low_vs_high.csv" --encoder_name="carp_640M" --checkpoint=0.25 --flatten_emb="mean" --seq_end_idx=56
# python run_pregen_emb.py --dataset_path="data/proeng/gb1/low_vs_high.csv" --encoder_name="carp_640M" --checkpoint=0.125 --flatten_emb="mean" --seq_end_idx=56


# python run_pregen_emb.py --dataset_path="data/proeng/gb1/two_vs_rest.csv" --encoder_name="carp_600k" --checkpoint=1 --flatten_emb="mean" --seq_end_idx=56 --reset_param=True
# python run_pregen_emb.py --dataset_path="data/proeng/gb1/two_vs_rest.csv" --encoder_name="carp_600k" --checkpoint=1 --flatten_emb="mean" --seq_end_idx=56 --resample_param=True
# python run_pregen_emb.py --dataset_path="data/proeng/gb1/two_vs_rest.csv" --encoder_name="carp_600k" --checkpoint=1 --flatten_emb="mean" --seq_end_idx=56
# python run_pregen_emb.py --dataset_path="data/proeng/gb1/two_vs_rest.csv" --encoder_name="carp_600k" --checkpoint=0.5 --flatten_emb="mean" --seq_end_idx=56
# python run_pregen_emb.py --dataset_path="data/proeng/gb1/two_vs_rest.csv" --encoder_name="carp_600k" --checkpoint=0.25 --flatten_emb="mean" --seq_end_idx=56
# python run_pregen_emb.py --dataset_path="data/proeng/gb1/two_vs_rest.csv" --encoder_name="carp_600k" --checkpoint=0.125 --flatten_emb="mean" --seq_end_idx=56

# python run_pregen_emb.py --dataset_path="data/proeng/gb1/two_vs_rest.csv" --encoder_name="carp_38M" --checkpoint=1 --flatten_emb="mean" --seq_end_idx=56 --reset_param=True
# python run_pregen_emb.py --dataset_path="data/proeng/gb1/two_vs_rest.csv" --encoder_name="carp_38M" --checkpoint=1 --flatten_emb="mean" --seq_end_idx=56 --resample_param=True
# python run_pregen_emb.py --dataset_path="data/proeng/gb1/two_vs_rest.csv" --encoder_name="carp_38M" --checkpoint=1 --flatten_emb="mean" --seq_end_idx=56
# python run_pregen_emb.py --dataset_path="data/proeng/gb1/two_vs_rest.csv" --encoder_name="carp_38M" --checkpoint=0.5 --flatten_emb="mean" --seq_end_idx=56
# python run_pregen_emb.py --dataset_path="data/proeng/gb1/two_vs_rest.csv" --encoder_name="carp_38M" --checkpoint=0.25 --flatten_emb="mean" --seq_end_idx=56
# python run_pregen_emb.py --dataset_path="data/proeng/gb1/two_vs_rest.csv" --encoder_name="carp_38M" --checkpoint=0.125 --flatten_emb="mean" --seq_end_idx=56

# python run_pregen_emb.py --dataset_path="data/proeng/gb1/two_vs_rest.csv" --encoder_name="carp_76M" --checkpoint=1 --flatten_emb="mean" --seq_end_idx=56 --reset_param=True
# python run_pregen_emb.py --dataset_path="data/proeng/gb1/two_vs_rest.csv" --encoder_name="carp_76M" --checkpoint=1 --flatten_emb="mean" --seq_end_idx=56 --resample_param=True
# python run_pregen_emb.py --dataset_path="data/proeng/gb1/two_vs_rest.csv" --encoder_name="carp_76M" --checkpoint=1 --flatten_emb="mean" --seq_end_idx=56
# python run_pregen_emb.py --dataset_path="data/proeng/gb1/two_vs_rest.csv" --encoder_name="carp_76M" --checkpoint=0.5 --flatten_emb="mean" --seq_end_idx=56
# python run_pregen_emb.py --dataset_path="data/proeng/gb1/two_vs_rest.csv" --encoder_name="carp_76M" --checkpoint=0.25 --flatten_emb="mean" --seq_end_idx=56
# python run_pregen_emb.py --dataset_path="data/proeng/gb1/two_vs_rest.csv" --encoder_name="carp_76M" --checkpoint=0.125 --flatten_emb="mean" --seq_end_idx=56

# python run_pregen_emb.py --dataset_path="data/proeng/gb1/two_vs_rest.csv" --encoder_name="carp_640M" --checkpoint=1 --flatten_emb="mean" --seq_end_idx=56 --reset_param=True
# python run_pregen_emb.py --dataset_path="data/proeng/gb1/two_vs_rest.csv" --encoder_name="carp_640M" --checkpoint=1 --flatten_emb="mean" --seq_end_idx=56 --resample_param=True
# python run_pregen_emb.py --dataset_path="data/proeng/gb1/two_vs_rest.csv" --encoder_name="carp_640M" --checkpoint=1 --flatten_emb="mean" --seq_end_idx=56
# python run_pregen_emb.py --dataset_path="data/proeng/gb1/two_vs_rest.csv" --encoder_name="carp_640M" --checkpoint=0.5 --flatten_emb="mean" --seq_end_idx=56
# python run_pregen_emb.py --dataset_path="data/proeng/gb1/two_vs_rest.csv" --encoder_name="carp_640M" --checkpoint=0.25 --flatten_emb="mean" --seq_end_idx=56
# python run_pregen_emb.py --dataset_path="data/proeng/gb1/two_vs_rest.csv" --encoder_name="carp_640M" --checkpoint=0.125 --flatten_emb="mean" --seq_end_idx=56


### AAV ###
# python run_pregen_emb.py --dataset_path="data/proeng/aav/two_vs_many.csv" --encoder_name="carp_600k" --checkpoint=1 --flatten_emb="mean" --reset_param=True
# python run_pregen_emb.py --dataset_path="data/proeng/aav/two_vs_many.csv" --encoder_name="carp_600k" --checkpoint=1 --flatten_emb="mean" --resample_param=True
# python run_pregen_emb.py --dataset_path="data/proeng/aav/two_vs_many.csv" --encoder_name="carp_600k" --checkpoint=1 --flatten_emb="mean"
# python run_pregen_emb.py --dataset_path="data/proeng/aav/two_vs_many.csv" --encoder_name="carp_600k" --checkpoint=0.5 --flatten_emb="mean"
# python run_pregen_emb.py --dataset_path="data/proeng/aav/two_vs_many.csv" --encoder_name="carp_600k" --checkpoint=0.25 --flatten_emb="mean"
# python run_pregen_emb.py --dataset_path="data/proeng/aav/two_vs_many.csv" --encoder_name="carp_600k" --checkpoint=0.125 --flatten_emb="mean"

# python run_pregen_emb.py --dataset_path="data/proeng/aav/two_vs_many.csv" --encoder_name="carp_38M" --checkpoint=1 --flatten_emb="mean" --reset_param=True
# python run_pregen_emb.py --dataset_path="data/proeng/aav/two_vs_many.csv" --encoder_name="carp_38M" --checkpoint=1 --flatten_emb="mean" --resample_param=True
# python run_pregen_emb.py --dataset_path="data/proeng/aav/two_vs_many.csv" --encoder_name="carp_38M" --checkpoint=1 --flatten_emb="mean"
# python run_pregen_emb.py --dataset_path="data/proeng/aav/two_vs_many.csv" --encoder_name="carp_38M" --checkpoint=0.5 --flatten_emb="mean"
# python run_pregen_emb.py --dataset_path="data/proeng/aav/two_vs_many.csv" --encoder_name="carp_38M" --checkpoint=0.25 --flatten_emb="mean"
# python run_pregen_emb.py --dataset_path="data/proeng/aav/two_vs_many.csv" --encoder_name="carp_38M" --checkpoint=0.125 --flatten_emb="mean"

# python run_pregen_emb.py --dataset_path="data/proeng/aav/two_vs_many.csv" --encoder_name="carp_76M" --checkpoint=1 --flatten_emb="mean" --reset_param=True
# python run_pregen_emb.py --dataset_path="data/proeng/aav/two_vs_many.csv" --encoder_name="carp_76M" --checkpoint=1 --flatten_emb="mean" --resample_param=True
# python run_pregen_emb.py --dataset_path="data/proeng/aav/two_vs_many.csv" --encoder_name="carp_76M" --checkpoint=1 --flatten_emb="mean"
# python run_pregen_emb.py --dataset_path="data/proeng/aav/two_vs_many.csv" --encoder_name="carp_76M" --checkpoint=0.5 --flatten_emb="mean"
# python run_pregen_emb.py --dataset_path="data/proeng/aav/two_vs_many.csv" --encoder_name="carp_76M" --checkpoint=0.25 --flatten_emb="mean"
# python run_pregen_emb.py --dataset_path="data/proeng/aav/two_vs_many.csv" --encoder_name="carp_76M" --checkpoint=0.125 --flatten_emb="mean"

# python run_pregen_emb.py --dataset_path="data/proeng/aav/two_vs_many.csv" --encoder_name="carp_640M" --checkpoint=1 --flatten_emb="mean" --reset_param=True
# python run_pregen_emb.py --dataset_path="data/proeng/aav/two_vs_many.csv" --encoder_name="carp_640M" --checkpoint=1 --flatten_emb="mean" --resample_param=True
# python run_pregen_emb.py --dataset_path="data/proeng/aav/two_vs_many.csv" --encoder_name="carp_640M" --checkpoint=1 --flatten_emb="mean"
# python run_pregen_emb.py --dataset_path="data/proeng/aav/two_vs_many.csv" --encoder_name="carp_640M" --checkpoint=0.5 --flatten_emb="mean"
# python run_pregen_emb.py --dataset_path="data/proeng/aav/two_vs_many.csv" --encoder_name="carp_640M" --checkpoint=0.25 --flatten_emb="mean"
# python run_pregen_emb.py --dataset_path="data/proeng/aav/two_vs_many.csv" --encoder_name="carp_640M" --checkpoint=0.125 --flatten_emb="mean"


# python run_pregen_emb.py --dataset_path="data/proeng/aav/one_vs_many.csv" --encoder_name="carp_600k" --checkpoint=1 --flatten_emb="mean" --reset_param=True
# python run_pregen_emb.py --dataset_path="data/proeng/aav/one_vs_many.csv" --encoder_name="carp_600k" --checkpoint=1 --flatten_emb="mean" --resample_param=True
# python run_pregen_emb.py --dataset_path="data/proeng/aav/one_vs_many.csv" --encoder_name="carp_600k" --checkpoint=1 --flatten_emb="mean"
# python run_pregen_emb.py --dataset_path="data/proeng/aav/one_vs_many.csv" --encoder_name="carp_600k" --checkpoint=0.5 --flatten_emb="mean"
# python run_pregen_emb.py --dataset_path="data/proeng/aav/one_vs_many.csv" --encoder_name="carp_600k" --checkpoint=0.25 --flatten_emb="mean"
# python run_pregen_emb.py --dataset_path="data/proeng/aav/one_vs_many.csv" --encoder_name="carp_600k" --checkpoint=0.125 --flatten_emb="mean"

# python run_pregen_emb.py --dataset_path="data/proeng/aav/one_vs_many.csv" --encoder_name="carp_38M" --checkpoint=1 --flatten_emb="mean" --reset_param=True
# python run_pregen_emb.py --dataset_path="data/proeng/aav/one_vs_many.csv" --encoder_name="carp_38M" --checkpoint=1 --flatten_emb="mean" --resample_param=True
# python run_pregen_emb.py --dataset_path="data/proeng/aav/one_vs_many.csv" --encoder_name="carp_38M" --checkpoint=1 --flatten_emb="mean"
# python run_pregen_emb.py --dataset_path="data/proeng/aav/one_vs_many.csv" --encoder_name="carp_38M" --checkpoint=0.5 --flatten_emb="mean"
# python run_pregen_emb.py --dataset_path="data/proeng/aav/one_vs_many.csv" --encoder_name="carp_38M" --checkpoint=0.25 --flatten_emb="mean"
# python run_pregen_emb.py --dataset_path="data/proeng/aav/one_vs_many.csv" --encoder_name="carp_38M" --checkpoint=0.125 --flatten_emb="mean"

# python run_pregen_emb.py --dataset_path="data/proeng/aav/one_vs_many.csv" --encoder_name="carp_76M" --checkpoint=1 --flatten_emb="mean" --reset_param=True
# python run_pregen_emb.py --dataset_path="data/proeng/aav/one_vs_many.csv" --encoder_name="carp_76M" --checkpoint=1 --flatten_emb="mean" --resample_param=True
# python run_pregen_emb.py --dataset_path="data/proeng/aav/one_vs_many.csv" --encoder_name="carp_76M" --checkpoint=1 --flatten_emb="mean"
# python run_pregen_emb.py --dataset_path="data/proeng/aav/one_vs_many.csv" --encoder_name="carp_76M" --checkpoint=0.5 --flatten_emb="mean"
# python run_pregen_emb.py --dataset_path="data/proeng/aav/one_vs_many.csv" --encoder_name="carp_76M" --checkpoint=0.25 --flatten_emb="mean"
# python run_pregen_emb.py --dataset_path="data/proeng/aav/one_vs_many.csv" --encoder_name="carp_76M" --checkpoint=0.125 --flatten_emb="mean"

# python run_pregen_emb.py --dataset_path="data/proeng/aav/one_vs_many.csv" --encoder_name="carp_640M" --checkpoint=1 --flatten_emb="mean" --reset_param=True
# python run_pregen_emb.py --dataset_path="data/proeng/aav/one_vs_many.csv" --encoder_name="carp_640M" --checkpoint=1 --flatten_emb="mean" --resample_param=True
# python run_pregen_emb.py --dataset_path="data/proeng/aav/one_vs_many.csv" --encoder_name="carp_640M" --checkpoint=1 --flatten_emb="mean"
# python run_pregen_emb.py --dataset_path="data/proeng/aav/one_vs_many.csv" --encoder_name="carp_640M" --checkpoint=0.5 --flatten_emb="mean"
# python run_pregen_emb.py --dataset_path="data/proeng/aav/one_vs_many.csv" --encoder_name="carp_640M" --checkpoint=0.25 --flatten_emb="mean"
# python run_pregen_emb.py --dataset_path="data/proeng/aav/one_vs_many.csv" --encoder_name="carp_640M" --checkpoint=0.125 --flatten_emb="mean"


### thermo ###

# python run_pregen_emb.py --dataset_path="data/proeng/thermo/mixed_split.csv" --encoder_name="carp_600k" --checkpoint=1 --flatten_emb="mean" --reset_param=True
# python run_pregen_emb.py --dataset_path="data/proeng/thermo/mixed_split.csv" --encoder_name="carp_600k" --checkpoint=1 --flatten_emb="mean" --resample_param=True
# python run_pregen_emb.py --dataset_path="data/proeng/thermo/mixed_split.csv" --encoder_name="carp_600k" --checkpoint=1 --flatten_emb="mean"
# python run_pregen_emb.py --dataset_path="data/proeng/thermo/mixed_split.csv" --encoder_name="carp_600k" --checkpoint=0.5 --flatten_emb="mean"
# python run_pregen_emb.py --dataset_path="data/proeng/thermo/mixed_split.csv" --encoder_name="carp_600k" --checkpoint=0.25 --flatten_emb="mean"
# python run_pregen_emb.py --dataset_path="data/proeng/thermo/mixed_split.csv" --encoder_name="carp_600k" --checkpoint=0.125 --flatten_emb="mean"

# python run_pregen_emb.py --dataset_path="data/proeng/thermo/mixed_split.csv" --encoder_name="carp_38M" --checkpoint=1 --flatten_emb="mean" --reset_param=True
# python run_pregen_emb.py --dataset_path="data/proeng/thermo/mixed_split.csv" --encoder_name="carp_38M" --checkpoint=1 --flatten_emb="mean" --resample_param=True
# python run_pregen_emb.py --dataset_path="data/proeng/thermo/mixed_split.csv" --encoder_name="carp_38M" --checkpoint=1 --flatten_emb="mean"
# python run_pregen_emb.py --dataset_path="data/proeng/thermo/mixed_split.csv" --encoder_name="carp_38M" --checkpoint=0.5 --flatten_emb="mean"
# python run_pregen_emb.py --dataset_path="data/proeng/thermo/mixed_split.csv" --encoder_name="carp_38M" --checkpoint=0.25 --flatten_emb="mean"
# python run_pregen_emb.py --dataset_path="data/proeng/thermo/mixed_split.csv" --encoder_name="carp_38M" --checkpoint=0.125 --flatten_emb="mean"

# python run_pregen_emb.py --dataset_path="data/proeng/thermo/mixed_split.csv" --encoder_name="carp_76M" --checkpoint=1 --flatten_emb="mean" --reset_param=True
# python run_pregen_emb.py --dataset_path="data/proeng/thermo/mixed_split.csv" --encoder_name="carp_76M" --checkpoint=1 --flatten_emb="mean" --resample_param=True
# python run_pregen_emb.py --dataset_path="data/proeng/thermo/mixed_split.csv" --encoder_name="carp_76M" --checkpoint=1 --flatten_emb="mean"
# python run_pregen_emb.py --dataset_path="data/proeng/thermo/mixed_split.csv" --encoder_name="carp_76M" --checkpoint=0.5 --flatten_emb="mean"
# python run_pregen_emb.py --dataset_path="data/proeng/thermo/mixed_split.csv" --encoder_name="carp_76M" --checkpoint=0.25 --flatten_emb="mean"
# python run_pregen_emb.py --dataset_path="data/proeng/thermo/mixed_split.csv" --encoder_name="carp_76M" --checkpoint=0.125 --flatten_emb="mean"

# python run_pregen_emb.py --dataset_path="data/proeng/thermo/mixed_split.csv" --encoder_name="carp_640M" --checkpoint=1 --flatten_emb="mean" --reset_param=True
# python run_pregen_emb.py --dataset_path="data/proeng/thermo/mixed_split.csv" --encoder_name="carp_640M" --checkpoint=1 --flatten_emb="mean" --resample_param=True
# python run_pregen_emb.py --dataset_path="data/proeng/thermo/mixed_split.csv" --encoder_name="carp_640M" --checkpoint=1 --flatten_emb="mean"
# python run_pregen_emb.py --dataset_path="data/proeng/thermo/mixed_split.csv" --encoder_name="carp_640M" --checkpoint=0.5 --flatten_emb="mean"
# python run_pregen_emb.py --dataset_path="data/proeng/thermo/mixed_split.csv" --encoder_name="carp_640M" --checkpoint=0.25 --flatten_emb="mean"
# python run_pregen_emb.py --dataset_path="data/proeng/thermo/mixed_split.csv" --encoder_name="carp_640M" --checkpoint=0.125 --flatten_emb="mean"


### Annotation ### 

# python run_pregen_emb.py --dataset_path="data/annotation/scl/balanced.csv" --encoder_name="carp_600k" --checkpoint=1 --flatten_emb="mean" --reset_param=True
# python run_pregen_emb.py --dataset_path="data/annotation/scl/balanced.csv" --encoder_name="carp_600k" --checkpoint=1 --flatten_emb="mean" --resample_param=True
# python run_pregen_emb.py --dataset_path="data/annotation/scl/balanced.csv" --encoder_name="carp_600k" --checkpoint=1 --flatten_emb="mean"
# python run_pregen_emb.py --dataset_path="data/annotation/scl/balanced.csv" --encoder_name="carp_600k" --checkpoint=0.5 --flatten_emb="mean"
# python run_pregen_emb.py --dataset_path="data/annotation/scl/balanced.csv" --encoder_name="carp_600k" --checkpoint=0.25 --flatten_emb="mean"
# python run_pregen_emb.py --dataset_path="data/annotation/scl/balanced.csv" --encoder_name="carp_600k" --checkpoint=0.125 --flatten_emb="mean"

# python run_pregen_emb.py --dataset_path="data/annotation/scl/balanced.csv" --encoder_name="carp_38M" --checkpoint=1 --flatten_emb="mean" --reset_param=True
# python run_pregen_emb.py --dataset_path="data/annotation/scl/balanced.csv" --encoder_name="carp_38M" --checkpoint=1 --flatten_emb="mean" --resample_param=True
# python run_pregen_emb.py --dataset_path="data/annotation/scl/balanced.csv" --encoder_name="carp_38M" --checkpoint=1 --flatten_emb="mean"
# python run_pregen_emb.py --dataset_path="data/annotation/scl/balanced.csv" --encoder_name="carp_38M" --checkpoint=0.5 --flatten_emb="mean"
# python run_pregen_emb.py --dataset_path="data/annotation/scl/balanced.csv" --encoder_name="carp_38M" --checkpoint=0.25 --flatten_emb="mean"
# python run_pregen_emb.py --dataset_path="data/annotation/scl/balanced.csv" --encoder_name="carp_38M" --checkpoint=0.125 --flatten_emb="mean"

# python run_pregen_emb.py --dataset_path="data/annotation/scl/balanced.csv" --encoder_name="carp_76M" --checkpoint=1 --flatten_emb="mean" --reset_param=True
# python run_pregen_emb.py --dataset_path="data/annotation/scl/balanced.csv" --encoder_name="carp_76M" --checkpoint=1 --flatten_emb="mean" --resample_param=True
# python run_pregen_emb.py --dataset_path="data/annotation/scl/balanced.csv" --encoder_name="carp_76M" --checkpoint=1 --flatten_emb="mean"
# python run_pregen_emb.py --dataset_path="data/annotation/scl/balanced.csv" --encoder_name="carp_76M" --checkpoint=0.5 --flatten_emb="mean"
# python run_pregen_emb.py --dataset_path="data/annotation/scl/balanced.csv" --encoder_name="carp_76M" --checkpoint=0.25 --flatten_emb="mean"
# python run_pregen_emb.py --dataset_path="data/annotation/scl/balanced.csv" --encoder_name="carp_76M" --checkpoint=0.125 --flatten_emb="mean"

# python run_pregen_emb.py --dataset_path="data/annotation/scl/balanced.csv" --encoder_name="carp_640M" --checkpoint=1 --flatten_emb="mean" --reset_param=True
# python run_pregen_emb.py --dataset_path="data/annotation/scl/balanced.csv" --encoder_name="carp_640M" --checkpoint=1 --flatten_emb="mean" --resample_param=True
# python run_pregen_emb.py --dataset_path="data/annotation/scl/balanced.csv" --encoder_name="carp_640M" --checkpoint=1 --flatten_emb="mean"
# python run_pregen_emb.py --dataset_path="data/annotation/scl/balanced.csv" --encoder_name="carp_640M" --checkpoint=0.5 --flatten_emb="mean"
# python run_pregen_emb.py --dataset_path="data/annotation/scl/balanced.csv" --encoder_name="carp_640M" --checkpoint=0.25 --flatten_emb="mean"
# python run_pregen_emb.py --dataset_path="data/annotation/scl/balanced.csv" --encoder_name="carp_640M" --checkpoint=0.125 --flatten_emb="mean"

### ss3 ###

# python run_pregen_emb.py --dataset_path="data/structure/secondary_structure/tape_ss3_processed.csv" --encoder_name="carp_600k" --subset_list='["train","val","cb513","ts115","casp12"]' --checkpoint=1 --reset_param=True
# python run_pregen_emb.py --dataset_path="data/structure/secondary_structure/tape_ss3_processed.csv" --encoder_name="carp_600k" --subset_list='["train","val","cb513","ts115","casp12"]' --checkpoint=1 --resample_param=True
# python run_pregen_emb.py --dataset_path="data/structure/secondary_structure/tape_ss3_processed.csv" --encoder_name="carp_600k" --subset_list='["train","val","cb513","ts115","casp12"]' --checkpoint=1
# python run_pregen_emb.py --dataset_path="data/structure/secondary_structure/tape_ss3_processed.csv" --encoder_name="carp_600k" --subset_list='["train","val","cb513","ts115","casp12"]' --checkpoint=0.5
# python run_pregen_emb.py --dataset_path="data/structure/secondary_structure/tape_ss3_processed.csv" --encoder_name="carp_600k" --subset_list='["train","val","cb513","ts115","casp12"]' --checkpoint=0.25
# python run_pregen_emb.py --dataset_path="data/structure/secondary_structure/tape_ss3_processed.csv" --encoder_name="carp_600k" --subset_list='["train","val","cb513","ts115","casp12"]' --checkpoint=0.125

# python run_pregen_emb.py --dataset_path="data/structure/secondary_structure/tape_ss3_processed.csv" --encoder_name="carp_38M" --subset_list='["train","val","cb513","ts115","casp12"]' --checkpoint=1 --reset_param=True
# python run_pregen_emb.py --dataset_path="data/structure/secondary_structure/tape_ss3_processed.csv" --encoder_name="carp_38M" --subset_list='["train","val","cb513","ts115","casp12"]' --checkpoint=1 --resample_param=True
# python run_pregen_emb.py --dataset_path="data/structure/secondary_structure/tape_ss3_processed.csv" --encoder_name="carp_38M" --subset_list='["train","val","cb513","ts115","casp12"]' --checkpoint=1
# python run_pregen_emb.py --dataset_path="data/structure/secondary_structure/tape_ss3_processed.csv" --encoder_name="carp_38M" --subset_list='["train","val","cb513","ts115","casp12"]' --checkpoint=0.5
# python run_pregen_emb.py --dataset_path="data/structure/secondary_structure/tape_ss3_processed.csv" --encoder_name="carp_38M" --subset_list='["train","val","cb513","ts115","casp12"]' --checkpoint=0.25
# python run_pregen_emb.py --dataset_path="data/structure/secondary_structure/tape_ss3_processed.csv" --encoder_name="carp_38M" --subset_list='["train","val","cb513","ts115","casp12"]' --checkpoint=0.125

# python run_pregen_emb.py --dataset_path="data/structure/secondary_structure/tape_ss3_processed.csv" --encoder_name="carp_76M" --subset_list='["train","val","cb513","ts115","casp12"]' --checkpoint=1 --reset_param=True
# python run_pregen_emb.py --dataset_path="data/structure/secondary_structure/tape_ss3_processed.csv" --encoder_name="carp_76M" --subset_list='["train","val","cb513","ts115","casp12"]' --checkpoint=1 --resample_param=True
# python run_pregen_emb.py --dataset_path="data/structure/secondary_structure/tape_ss3_processed.csv" --encoder_name="carp_76M" --subset_list='["train","val","cb513","ts115","casp12"]' --checkpoint=1
# python run_pregen_emb.py --dataset_path="data/structure/secondary_structure/tape_ss3_processed.csv" --encoder_name="carp_76M" --subset_list='["train","val","cb513","ts115","casp12"]' --checkpoint=0.5
# python run_pregen_emb.py --dataset_path="data/structure/secondary_structure/tape_ss3_processed.csv" --encoder_name="carp_76M" --subset_list='["train","val","cb513","ts115","casp12"]' --checkpoint=0.25
# python run_pregen_emb.py --dataset_path="data/structure/secondary_structure/tape_ss3_processed.csv" --encoder_name="carp_76M" --subset_list='["train","val","cb513","ts115","casp12"]' --checkpoint=0.125

# python run_pregen_emb.py --dataset_path="data/structure/secondary_structure/tape_ss3_processed.csv" --encoder_name="carp_640M" --subset_list='["train","val","cb513","ts115","casp12"]' --checkpoint=1 --reset_param=True
# python run_pregen_emb.py --dataset_path="data/structure/secondary_structure/tape_ss3_processed.csv" --encoder_name="carp_640M" --subset_list='["train","val","cb513","ts115","casp12"]' --checkpoint=1 --resample_param=True
# python run_pregen_emb.py --dataset_path="data/structure/secondary_structure/tape_ss3_processed.csv" --encoder_name="carp_640M" --subset_list='["train","val","cb513","ts115","casp12"]' --checkpoint=1
# python run_pregen_emb.py --dataset_path="data/structure/secondary_structure/tape_ss3_processed.csv" --encoder_name="carp_640M" --subset_list='["train","val","cb513","ts115","casp12"]' --checkpoint=0.5
# python run_pregen_emb.py --dataset_path="data/structure/secondary_structure/tape_ss3_processed.csv" --encoder_name="carp_640M" --subset_list='["train","val","cb513","ts115","casp12"]' --checkpoint=0.25
# python run_pregen_emb.py --dataset_path="data/structure/secondary_structure/tape_ss3_processed.csv" --encoder_name="carp_640M" --subset_list='["train","val","cb513","ts115","casp12"]' --checkpoint=0.125
```
