## Supplemental data and code for "Feature Reuse and Scaling: Understanding Transfer Learning with Protein Language Models": execute_gen_emb_esm.sh

```
# queue for generating and sorting all embs

export CUDA_VISIBLE_DEVICES=1

### GB1 ###

# python run_pregen_emb.py --dataset_path="data/proeng/gb1/sampled.csv" --encoder_name="esm1_t6_43M_UR50S" --flatten_emb="mean" --seq_end_idx=56 --reset_param=True
# python run_pregen_emb.py --dataset_path="data/proeng/gb1/sampled.csv" --encoder_name="esm1_t6_43M_UR50S" --flatten_emb="mean" --seq_end_idx=56 --resample_param=True
# python run_pregen_emb.py --dataset_path="data/proeng/gb1/sampled.csv" --encoder_name="esm1_t6_43M_UR50S" --flatten_emb="mean" --seq_end_idx=56

# python run_pregen_emb.py --dataset_path="data/proeng/gb1/sampled.csv" --encoder_name="esm1_t12_85M_UR50S" --flatten_emb="mean" --seq_end_idx=56 --reset_param=True
# python run_pregen_emb.py --dataset_path="data/proeng/gb1/sampled.csv" --encoder_name="esm1_t12_85M_UR50S" --flatten_emb="mean" --seq_end_idx=56 --resample_param=True
# python run_pregen_emb.py --dataset_path="data/proeng/gb1/sampled.csv" --encoder_name="esm1_t12_85M_UR50S" --flatten_emb="mean" --seq_end_idx=56

# python run_pregen_emb.py --dataset_path="data/proeng/gb1/sampled.csv" --encoder_name="esm1_t34_670M_UR50S" --flatten_emb="mean" --seq_end_idx=56 --reset_param=True
# python run_pregen_emb.py --dataset_path="data/proeng/gb1/sampled.csv" --encoder_name="esm1_t34_670M_UR50S" --flatten_emb="mean" --seq_end_idx=56 --resample_param=True
# python run_pregen_emb.py --dataset_path="data/proeng/gb1/sampled.csv" --encoder_name="esm1_t34_670M_UR50S" --flatten_emb="mean" --seq_end_idx=56

# python run_pregen_emb.py --dataset_path="data/proeng/gb1/sampled.csv" --encoder_name="esm1b_t33_650M_UR50S" --flatten_emb="mean" --seq_end_idx=56 --reset_param=True
# python run_pregen_emb.py --dataset_path="data/proeng/gb1/sampled.csv" --encoder_name="esm1b_t33_650M_UR50S" --flatten_emb="mean" --seq_end_idx=56 --resample_param=True
# python run_pregen_emb.py --dataset_path="data/proeng/gb1/sampled.csv" --encoder_name="esm1b_t33_650M_UR50S" --flatten_emb="mean" --seq_end_idx=56


# python run_pregen_emb.py --dataset_path="data/proeng/gb1/low_vs_high.csv" --encoder_name="esm1_t6_43M_UR50S" --flatten_emb="mean" --seq_end_idx=56 --reset_param=True
# python run_pregen_emb.py --dataset_path="data/proeng/gb1/low_vs_high.csv" --encoder_name="esm1_t6_43M_UR50S" --flatten_emb="mean" --seq_end_idx=56 --resample_param=True
# python run_pregen_emb.py --dataset_path="data/proeng/gb1/low_vs_high.csv" --encoder_name="esm1_t6_43M_UR50S" --flatten_emb="mean" --seq_end_idx=56

# python run_pregen_emb.py --dataset_path="data/proeng/gb1/low_vs_high.csv" --encoder_name="esm1_t12_85M_UR50S" --flatten_emb="mean" --seq_end_idx=56 --reset_param=True
# python run_pregen_emb.py --dataset_path="data/proeng/gb1/low_vs_high.csv" --encoder_name="esm1_t12_85M_UR50S" --flatten_emb="mean" --seq_end_idx=56 --resample_param=True
# python run_pregen_emb.py --dataset_path="data/proeng/gb1/low_vs_high.csv" --encoder_name="esm1_t12_85M_UR50S" --flatten_emb="mean" --seq_end_idx=56

# python run_pregen_emb.py --dataset_path="data/proeng/gb1/low_vs_high.csv" --encoder_name="esm1_t34_670M_UR50S" --flatten_emb="mean" --seq_end_idx=56 --reset_param=True
# python run_pregen_emb.py --dataset_path="data/proeng/gb1/low_vs_high.csv" --encoder_name="esm1_t34_670M_UR50S" --flatten_emb="mean" --seq_end_idx=56 --resample_param=True
# python run_pregen_emb.py --dataset_path="data/proeng/gb1/low_vs_high.csv" --encoder_name="esm1_t34_670M_UR50S" --flatten_emb="mean" --seq_end_idx=56

# python run_pregen_emb.py --dataset_path="data/proeng/gb1/low_vs_high.csv" --encoder_name="esm1b_t33_650M_UR50S" --flatten_emb="mean" --seq_end_idx=56 --reset_param=True
# python run_pregen_emb.py --dataset_path="data/proeng/gb1/low_vs_high.csv" --encoder_name="esm1b_t33_650M_UR50S" --flatten_emb="mean" --seq_end_idx=56 --resample_param=True
# python run_pregen_emb.py --dataset_path="data/proeng/gb1/low_vs_high.csv" --encoder_name="esm1b_t33_650M_UR50S" --flatten_emb="mean" --seq_end_idx=56


# python run_pregen_emb.py --dataset_path="data/proeng/gb1/two_vs_rest.csv" --encoder_name="esm1_t6_43M_UR50S" --flatten_emb="mean" --seq_end_idx=56 --reset_param=True
# python run_pregen_emb.py --dataset_path="data/proeng/gb1/two_vs_rest.csv" --encoder_name="esm1_t6_43M_UR50S" --flatten_emb="mean" --seq_end_idx=56 --resample_param=True
# python run_pregen_emb.py --dataset_path="data/proeng/gb1/two_vs_rest.csv" --encoder_name="esm1_t6_43M_UR50S" --flatten_emb="mean" --seq_end_idx=56

# python run_pregen_emb.py --dataset_path="data/proeng/gb1/two_vs_rest.csv" --encoder_name="esm1_t12_85M_UR50S" --flatten_emb="mean" --seq_end_idx=56 --reset_param=True
# python run_pregen_emb.py --dataset_path="data/proeng/gb1/two_vs_rest.csv" --encoder_name="esm1_t12_85M_UR50S" --flatten_emb="mean" --seq_end_idx=56 --resample_param=True
# python run_pregen_emb.py --dataset_path="data/proeng/gb1/two_vs_rest.csv" --encoder_name="esm1_t12_85M_UR50S" --flatten_emb="mean" --seq_end_idx=56

# python run_pregen_emb.py --dataset_path="data/proeng/gb1/two_vs_rest.csv" --encoder_name="esm1_t34_670M_UR50S" --flatten_emb="mean" --seq_end_idx=56 --reset_param=True
# python run_pregen_emb.py --dataset_path="data/proeng/gb1/two_vs_rest.csv" --encoder_name="esm1_t34_670M_UR50S" --flatten_emb="mean" --seq_end_idx=56 --resample_param=True
# python run_pregen_emb.py --dataset_path="data/proeng/gb1/two_vs_rest.csv" --encoder_name="esm1_t34_670M_UR50S" --flatten_emb="mean" --seq_end_idx=56

# python run_pregen_emb.py --dataset_path="data/proeng/gb1/two_vs_rest.csv" --encoder_name="esm1b_t33_650M_UR50S" --flatten_emb="mean" --seq_end_idx=56 --reset_param=True
# python run_pregen_emb.py --dataset_path="data/proeng/gb1/two_vs_rest.csv" --encoder_name="esm1b_t33_650M_UR50S" --flatten_emb="mean" --seq_end_idx=56 --resample_param=True
# python run_pregen_emb.py --dataset_path="data/proeng/gb1/two_vs_rest.csv" --encoder_name="esm1b_t33_650M_UR50S" --flatten_emb="mean" --seq_end_idx=56


### AAV ###

# python run_pregen_emb.py --dataset_path="data/proeng/aav/two_vs_many.csv" --encoder_name="esm1_t6_43M_UR50S" --flatten_emb="mean" --reset_param=True
# python run_pregen_emb.py --dataset_path="data/proeng/aav/two_vs_many.csv" --encoder_name="esm1_t6_43M_UR50S" --flatten_emb="mean" --resample_param=True
# python run_pregen_emb.py --dataset_path="data/proeng/aav/two_vs_many.csv" --encoder_name="esm1_t6_43M_UR50S" --flatten_emb="mean"

# python run_pregen_emb.py --dataset_path="data/proeng/aav/two_vs_many.csv" --encoder_name="esm1_t12_85M_UR50S" --flatten_emb="mean" --reset_param=True
# python run_pregen_emb.py --dataset_path="data/proeng/aav/two_vs_many.csv" --encoder_name="esm1_t12_85M_UR50S" --flatten_emb="mean" --resample_param=True
# python run_pregen_emb.py --dataset_path="data/proeng/aav/two_vs_many.csv" --encoder_name="esm1_t12_85M_UR50S" --flatten_emb="mean"

# python run_pregen_emb.py --dataset_path="data/proeng/aav/two_vs_many.csv" --encoder_name="esm1_t34_670M_UR50S" --flatten_emb="mean" --reset_param=True
# python run_pregen_emb.py --dataset_path="data/proeng/aav/two_vs_many.csv" --encoder_name="esm1_t34_670M_UR50S" --flatten_emb="mean" --resample_param=True
# python run_pregen_emb.py --dataset_path="data/proeng/aav/two_vs_many.csv" --encoder_name="esm1_t34_670M_UR50S" --flatten_emb="mean"

# python run_pregen_emb.py --dataset_path="data/proeng/aav/two_vs_many.csv" --encoder_name="esm1b_t33_650M_UR50S" --flatten_emb="mean" --reset_param=True
# python run_pregen_emb.py --dataset_path="data/proeng/aav/two_vs_many.csv" --encoder_name="esm1b_t33_650M_UR50S" --flatten_emb="mean" --resample_param=True
# python run_pregen_emb.py --dataset_path="data/proeng/aav/two_vs_many.csv" --encoder_name="esm1b_t33_650M_UR50S" --flatten_emb="mean"


# python run_pregen_emb.py --dataset_path="data/proeng/aav/one_vs_many.csv" --encoder_name="esm1_t6_43M_UR50S" --flatten_emb="mean" --reset_param=True
# python run_pregen_emb.py --dataset_path="data/proeng/aav/one_vs_many.csv" --encoder_name="esm1_t6_43M_UR50S" --flatten_emb="mean" --resample_param=True
# python run_pregen_emb.py --dataset_path="data/proeng/aav/one_vs_many.csv" --encoder_name="esm1_t6_43M_UR50S" --flatten_emb="mean"

# python run_pregen_emb.py --dataset_path="data/proeng/aav/one_vs_many.csv" --encoder_name="esm1_t12_85M_UR50S" --flatten_emb="mean" --reset_param=True
# python run_pregen_emb.py --dataset_path="data/proeng/aav/one_vs_many.csv" --encoder_name="esm1_t12_85M_UR50S" --flatten_emb="mean" --resample_param=True
# python run_pregen_emb.py --dataset_path="data/proeng/aav/one_vs_many.csv" --encoder_name="esm1_t12_85M_UR50S" --flatten_emb="mean"

# python run_pregen_emb.py --dataset_path="data/proeng/aav/one_vs_many.csv" --encoder_name="esm1_t34_670M_UR50S" --flatten_emb="mean" --reset_param=True
# python run_pregen_emb.py --dataset_path="data/proeng/aav/one_vs_many.csv" --encoder_name="esm1_t34_670M_UR50S" --flatten_emb="mean" --resample_param=True
# python run_pregen_emb.py --dataset_path="data/proeng/aav/one_vs_many.csv" --encoder_name="esm1_t34_670M_UR50S" --flatten_emb="mean"

# python run_pregen_emb.py --dataset_path="data/proeng/aav/one_vs_many.csv" --encoder_name="esm1b_t33_650M_UR50S" --flatten_emb="mean" --reset_param=True
# python run_pregen_emb.py --dataset_path="data/proeng/aav/one_vs_many.csv" --encoder_name="esm1b_t33_650M_UR50S" --flatten_emb="mean" --resample_param=True
# python run_pregen_emb.py --dataset_path="data/proeng/aav/one_vs_many.csv" --encoder_name="esm1b_t33_650M_UR50S" --flatten_emb="mean"


### thermo ###

# python run_pregen_emb.py --dataset_path="data/proeng/thermo/mixed_split.csv" --encoder_name="esm1_t6_43M_UR50S" --flatten_emb="mean" --reset_param=True
# python run_pregen_emb.py --dataset_path="data/proeng/thermo/mixed_split.csv" --encoder_name="esm1_t6_43M_UR50S" --flatten_emb="mean" --resample_param=True
# python run_pregen_emb.py --dataset_path="data/proeng/thermo/mixed_split.csv" --encoder_name="esm1_t6_43M_UR50S" --flatten_emb="mean"

# python run_pregen_emb.py --dataset_path="data/proeng/thermo/mixed_split.csv" --encoder_name="esm1_t12_85M_UR50S" --flatten_emb="mean" --reset_param=True
# python run_pregen_emb.py --dataset_path="data/proeng/thermo/mixed_split.csv" --encoder_name="esm1_t12_85M_UR50S" --flatten_emb="mean" --resample_param=True
# python run_pregen_emb.py --dataset_path="data/proeng/thermo/mixed_split.csv" --encoder_name="esm1_t12_85M_UR50S" --flatten_emb="mean"

# python run_pregen_emb.py --dataset_path="data/proeng/thermo/mixed_split.csv" --encoder_name="esm1_t34_670M_UR50S" --flatten_emb="mean" --reset_param=True
# python run_pregen_emb.py --dataset_path="data/proeng/thermo/mixed_split.csv" --encoder_name="esm1_t34_670M_UR50S" --flatten_emb="mean" --resample_param=True
# python run_pregen_emb.py --dataset_path="data/proeng/thermo/mixed_split.csv" --encoder_name="esm1_t34_670M_UR50S" --flatten_emb="mean"

# python run_pregen_emb.py --dataset_path="data/proeng/thermo/mixed_split.csv" --encoder_name="esm1b_t33_650M_UR50S" --flatten_emb="mean" --reset_param=True
# python run_pregen_emb.py --dataset_path="data/proeng/thermo/mixed_split.csv" --encoder_name="esm1b_t33_650M_UR50S" --flatten_emb="mean" --resample_param=True
# python run_pregen_emb.py --dataset_path="data/proeng/thermo/mixed_split.csv" --encoder_name="esm1b_t33_650M_UR50S" --flatten_emb="mean"


### Annotation ### 

# python run_pregen_emb.py --dataset_path="data/annotation/scl/balanced.csv" --encoder_name="esm1_t6_43M_UR50S" --flatten_emb="mean" --reset_param=True
# python run_pregen_emb.py --dataset_path="data/annotation/scl/balanced.csv" --encoder_name="esm1_t6_43M_UR50S" --flatten_emb="mean" --resample_param=True
# python run_pregen_emb.py --dataset_path="data/annotation/scl/balanced.csv" --encoder_name="esm1_t6_43M_UR50S" --flatten_emb="mean"

# python run_pregen_emb.py --dataset_path="data/annotation/scl/balanced.csv" --encoder_name="esm1_t12_85M_UR50S" --flatten_emb="mean" --reset_param=True
# python run_pregen_emb.py --dataset_path="data/annotation/scl/balanced.csv" --encoder_name="esm1_t12_85M_UR50S" --flatten_emb="mean" --resample_param=True
# python run_pregen_emb.py --dataset_path="data/annotation/scl/balanced.csv" --encoder_name="esm1_t12_85M_UR50S" --flatten_emb="mean"

# python run_pregen_emb.py --dataset_path="data/annotation/scl/balanced.csv" --encoder_name="esm1_t34_670M_UR50S" --flatten_emb="mean" --reset_param=True
# python run_pregen_emb.py --dataset_path="data/annotation/scl/balanced.csv" --encoder_name="esm1_t34_670M_UR50S" --flatten_emb="mean" --resample_param=True
# python run_pregen_emb.py --dataset_path="data/annotation/scl/balanced.csv" --encoder_name="esm1_t34_670M_UR50S" --flatten_emb="mean"

# python run_pregen_emb.py --dataset_path="data/annotation/scl/balanced.csv" --encoder_name="esm1b_t33_650M_UR50S" --flatten_emb="mean" --reset_param=True
# python run_pregen_emb.py --dataset_path="data/annotation/scl/balanced.csv" --encoder_name="esm1b_t33_650M_UR50S" --flatten_emb="mean" --resample_param=True
# python run_pregen_emb.py --dataset_path="data/annotation/scl/balanced.csv" --encoder_name="esm1b_t33_650M_UR50S" --flatten_emb="mean"

### ss3 ###

# python run_pregen_emb.py --dataset_path="data/structure/secondary_structure/tape_ss3_processed.csv" --encoder_name="esm1_t6_43M_UR50S" --subset_list='["train","val","cb513","ts115","casp12"]'
# python run_pregen_emb.py --dataset_path="data/structure/secondary_structure/tape_ss3_processed.csv" --encoder_name="esm1_t6_43M_UR50S" --subset_list='["train","val", "cb513", "ts115", "casp12"]' --reset_param=True
# python run_pregen_emb.py --dataset_path="data/structure/secondary_structure/tape_ss3_processed.csv" --encoder_name="esm1_t6_43M_UR50S" --subset_list='["train","val", "cb513", "ts115", "casp12"]' --resample_param=True

# python run_pregen_emb.py --dataset_path="data/structure/secondary_structure/tape_ss3_processed.csv" --encoder_name="esm1_t12_85M_UR50S" --subset_list='["train","val","cb513","ts115","casp12"]'
# python run_pregen_emb.py --dataset_path="data/structure/secondary_structure/tape_ss3_processed.csv" --encoder_name="esm1_t12_85M_UR50S" --subset_list='["train","val","cb513","ts115","casp12"]' --reset_param=True
# python run_pregen_emb.py --dataset_path="data/structure/secondary_structure/tape_ss3_processed.csv" --encoder_name="esm1_t12_85M_UR50S" --subset_list='["train","val","cb513","ts115","casp12"]' --resample_param=True

# python run_pregen_emb.py --dataset_path="data/structure/secondary_structure/tape_ss3_processed.csv" --encoder_name="esm1_t34_670M_UR50S" --subset_list='["train","val","cb513","ts115","casp12"]'
# python run_pregen_emb.py --dataset_path="data/structure/secondary_structure/tape_ss3_processed.csv" --encoder_name="esm1_t34_670M_UR50S" --subset_list='["train","val","cb513","ts115","casp12"]' --reset_param=True
# python run_pregen_emb.py --dataset_path="data/structure/secondary_structure/tape_ss3_processed.csv" --encoder_name="esm1_t34_670M_UR50S" --subset_list='["train","val","cb513","ts115","casp12"]' --resample_param=True

# python run_pregen_emb.py --dataset_path="data/structure/secondary_structure/tape_ss3_processed.csv" --encoder_name="esm1b_t33_650M_UR50S" --subset_list='["train","val","cb513","ts115","casp12"]'
# python run_pregen_emb.py --dataset_path="data/structure/secondary_structure/tape_ss3_processed.csv" --encoder_name="esm1b_t33_650M_UR50S" --subset_list='["train","val","cb513","ts115","casp12"]' --reset_param=True
# python run_pregen_emb.py --dataset_path="data/structure/secondary_structure/tape_ss3_processed.csv" --encoder_name="esm1b_t33_650M_UR50S" --subset_list='["train","val","cb513","ts115","casp12"]' --resample_param=True
```
