## Supplemental data and code for "Feature Reuse and Scaling: Understanding Transfer Learning with Protein Language Models": execute_run_pytorch_carp.sh

```
# queue for running all pytorch models from embs

# if run on cpu
# export CUDA_VISIBLE_DEVICES=""

# if run on cuda:1
# export CUDA_VISIBLE_DEVICES=1

### scl ###

# python run_protran_pytorch.py --dataset_path="data/annotation/scl/balanced.csv" --encoder_name="carp_600k" --checkpoint=1 --embed_batch_size=64 --flatten_emb="mean" --embed_folder="embeddings" --loader_batch_size=256 --epochs=100 --all_plot_folder="results/pytorch_learning_curves-carp" --all_result_folder="results/pytorch-carp" --reset_param=True  &
# python run_protran_pytorch.py --dataset_path="data/annotation/scl/balanced.csv" --encoder_name="carp_600k" --checkpoint=1 --embed_batch_size=64 --flatten_emb="mean" --embed_folder="embeddings" --loader_batch_size=256 --epochs=100 --all_plot_folder="results/pytorch_learning_curves-carp" --all_result_folder="results/pytorch-carp" --resample_param=True &
# python run_protran_pytorch.py --dataset_path="data/annotation/scl/balanced.csv" --encoder_name="carp_600k" --checkpoint=1 --embed_batch_size=64 --flatten_emb="mean" --embed_folder="embeddings" --loader_batch_size=256 --epochs=100 --all_plot_folder="results/pytorch_learning_curves-carp" --all_result_folder="results/pytorch-carp" &
# python run_protran_pytorch.py --dataset_path="data/annotation/scl/balanced.csv" --encoder_name="carp_600k" --checkpoint=0.5 --embed_batch_size=64 --flatten_emb="mean" --embed_folder="embeddings" --loader_batch_size=256 --epochs=100 --all_plot_folder="results/pytorch_learning_curves-carp" --all_result_folder="results/pytorch-carp" &
# python run_protran_pytorch.py --dataset_path="data/annotation/scl/balanced.csv" --encoder_name="carp_600k" --checkpoint=0.25 --embed_batch_size=64 --flatten_emb="mean" --embed_folder="embeddings" --loader_batch_size=256 --epochs=100 --all_plot_folder="results/pytorch_learning_curves-carp" --all_result_folder="results/pytorch-carp" &
# python run_protran_pytorch.py --dataset_path="data/annotation/scl/balanced.csv" --encoder_name="carp_600k" --checkpoint=0.125 --embed_batch_size=64 --flatten_emb="mean" --embed_folder="embeddings" --loader_batch_size=256 --epochs=100 --all_plot_folder="results/pytorch_learning_curves-carp" --all_result_folder="results/pytorch-carp" &

# python run_protran_pytorch.py --dataset_path="data/annotation/scl/balanced.csv" --encoder_name="carp_38M" --checkpoint=1 --embed_batch_size=64 --flatten_emb="mean" --embed_folder="embeddings" --loader_batch_size=256 --epochs=100 --all_plot_folder="results/pytorch_learning_curves-carp" --all_result_folder="results/pytorch-carp" --reset_param=True &
# python run_protran_pytorch.py --dataset_path="data/annotation/scl/balanced.csv" --encoder_name="carp_38M" --checkpoint=1 --embed_batch_size=64 --flatten_emb="mean" --embed_folder="embeddings" --loader_batch_size=256 --epochs=100 --all_plot_folder="results/pytorch_learning_curves-carp" --all_result_folder="results/pytorch-carp" --resample_param=True &
# python run_protran_pytorch.py --dataset_path="data/annotation/scl/balanced.csv" --encoder_name="carp_38M" --checkpoint=1 --embed_batch_size=64 --flatten_emb="mean" --embed_folder="embeddings" --loader_batch_size=256 --epochs=100 --all_plot_folder="results/pytorch_learning_curves-carp" --all_result_folder="results/pytorch-carp" &
# python run_protran_pytorch.py --dataset_path="data/annotation/scl/balanced.csv" --encoder_name="carp_38M" --checkpoint=0.5 --embed_batch_size=64 --flatten_emb="mean" --embed_folder="embeddings" --loader_batch_size=256 --epochs=100 --all_plot_folder="results/pytorch_learning_curves-carp" --all_result_folder="results/pytorch-carp" &
# python run_protran_pytorch.py --dataset_path="data/annotation/scl/balanced.csv" --encoder_name="carp_38M" --checkpoint=0.25 --embed_batch_size=64 --flatten_emb="mean" --embed_folder="embeddings" --loader_batch_size=256 --epochs=100 --all_plot_folder="results/pytorch_learning_curves-carp" --all_result_folder="results/pytorch-carp" &
# python run_protran_pytorch.py --dataset_path="data/annotation/scl/balanced.csv" --encoder_name="carp_38M" --checkpoint=0.125 --embed_batch_size=64 --flatten_emb="mean" --embed_folder="embeddings" --loader_batch_size=256 --epochs=100 --all_plot_folder="results/pytorch_learning_curves-carp" --all_result_folder="results/pytorch-carp" &

# python run_protran_pytorch.py --dataset_path="data/annotation/scl/balanced.csv" --encoder_name="carp_76M" --checkpoint=1 --embed_batch_size=64 --flatten_emb="mean" --embed_folder="embeddings" --loader_batch_size=256 --epochs=100 --all_plot_folder="results/pytorch_learning_curves-carp" --all_result_folder="results/pytorch-carp" --reset_param=True &
# python run_protran_pytorch.py --dataset_path="data/annotation/scl/balanced.csv" --encoder_name="carp_76M" --checkpoint=1 --embed_batch_size=64 --flatten_emb="mean" --embed_folder="embeddings" --loader_batch_size=256 --epochs=100 --all_plot_folder="results/pytorch_learning_curves-carp" --all_result_folder="results/pytorch-carp" --resample_param=True &
# python run_protran_pytorch.py --dataset_path="data/annotation/scl/balanced.csv" --encoder_name="carp_76M" --checkpoint=1 --embed_batch_size=64 --flatten_emb="mean" --embed_folder="embeddings" --loader_batch_size=256 --epochs=100 --all_plot_folder="results/pytorch_learning_curves-carp" --all_result_folder="results/pytorch-carp" &
# python run_protran_pytorch.py --dataset_path="data/annotation/scl/balanced.csv" --encoder_name="carp_76M" --checkpoint=0.5 --embed_batch_size=64 --flatten_emb="mean" --embed_folder="embeddings" --loader_batch_size=256 --epochs=100 --all_plot_folder="results/pytorch_learning_curves-carp" --all_result_folder="results/pytorch-carp" &
# python run_protran_pytorch.py --dataset_path="data/annotation/scl/balanced.csv" --encoder_name="carp_76M" --checkpoint=0.25 --embed_batch_size=64 --flatten_emb="mean" --embed_folder="embeddings" --loader_batch_size=256 --epochs=100 --all_plot_folder="results/pytorch_learning_curves-carp" --all_result_folder="results/pytorch-carp" &
# python run_protran_pytorch.py --dataset_path="data/annotation/scl/balanced.csv" --encoder_name="carp_76M" --checkpoint=0.125 --embed_batch_size=64 --flatten_emb="mean" --embed_folder="embeddings" --loader_batch_size=256 --epochs=100 --all_plot_folder="results/pytorch_learning_curves-carp" --all_result_folder="results/pytorch-carp" &

# python run_protran_pytorch.py --dataset_path="data/annotation/scl/balanced.csv" --encoder_name="carp_640M" --checkpoint=1 --embed_batch_size=64 --flatten_emb="mean" --embed_folder="embeddings" --loader_batch_size=256 --epochs=100 --all_plot_folder="results/pytorch_learning_curves-carp" --all_result_folder="results/pytorch-carp" --reset_param=True &
# python run_protran_pytorch.py --dataset_path="data/annotation/scl/balanced.csv" --encoder_name="carp_640M" --checkpoint=1 --embed_batch_size=64 --flatten_emb="mean" --embed_folder="embeddings" --loader_batch_size=256 --epochs=100 --all_plot_folder="results/pytorch_learning_curves-carp" --all_result_folder="results/pytorch-carp" --resample_param=True &
# python run_protran_pytorch.py --dataset_path="data/annotation/scl/balanced.csv" --encoder_name="carp_640M" --checkpoint=1 --embed_batch_size=64 --flatten_emb="mean" --embed_folder="embeddings" --loader_batch_size=256 --epochs=100 --all_plot_folder="results/pytorch_learning_curves-carp" --all_result_folder="results/pytorch-carp" &
# python run_protran_pytorch.py --dataset_path="data/annotation/scl/balanced.csv" --encoder_name="carp_640M" --checkpoint=0.5 --embed_batch_size=64 --flatten_emb="mean" --embed_folder="embeddings" --loader_batch_size=256 --epochs=100 --all_plot_folder="results/pytorch_learning_curves-carp" --all_result_folder="results/pytorch-carp" &
# python run_protran_pytorch.py --dataset_path="data/annotation/scl/balanced.csv" --encoder_name="carp_640M" --checkpoint=0.25 --embed_batch_size=64 --flatten_emb="mean" --embed_folder="embeddings" --loader_batch_size=256 --epochs=100 --all_plot_folder="results/pytorch_learning_curves-carp" --all_result_folder="results/pytorch-carp" &
# python run_protran_pytorch.py --dataset_path="data/annotation/scl/balanced.csv" --encoder_name="carp_640M" --checkpoint=0.125 --embed_batch_size=64 --flatten_emb="mean" --embed_folder="embeddings" --loader_batch_size=256 --epochs=100 --all_plot_folder="results/pytorch_learning_curves-carp" --all_result_folder="results/pytorch-carp" &

### ss3 ###

# for i in 0 1 2 3 4 5 6 7 8 9 10 11 12 13 14 15 16; do
#     python run_protran_pytorch.py --dataset_path="data/structure/secondary_structure/tape_ss3_processed.csv" --encoder_name="carp_600k" --checkpoint=1 --embed_batch_size=64 --embed_folder="embeddings" --loader_batch_size=120 --epochs=100 --all_plot_folder="results/pytorch_learning_curves-carp" --all_result_folder="results/pytorch-carp" --reset_param=True --manual_layer_min="${i}" --manual_layer_max="${i}"
#     python run_protran_pytorch.py --dataset_path="data/structure/secondary_structure/tape_ss3_processed.csv" --encoder_name="carp_600k" --checkpoint=1 --embed_batch_size=64 --embed_folder="embeddings" --loader_batch_size=120 --epochs=100 --all_plot_folder="results/pytorch_learning_curves-carp" --all_result_folder="results/pytorch-carp" --resample_param=True --manual_layer_min="${i}" --manual_layer_max="${i}"
#     python run_protran_pytorch.py --dataset_path="data/structure/secondary_structure/tape_ss3_processed.csv" --encoder_name="carp_600k" --checkpoint=1 --embed_batch_size=64 --embed_folder="embeddings" --loader_batch_size=120 --epochs=100 --all_plot_folder="results/pytorch_learning_curves-carp" --all_result_folder="results/pytorch-carp" --manual_layer_min="${i}" --manual_layer_max="${i}"
#     python run_protran_pytorch.py --dataset_path="data/structure/secondary_structure/tape_ss3_processed.csv" --encoder_name="carp_600k" --checkpoint=0.5 --embed_batch_size=64 --embed_folder="embeddings" --loader_batch_size=120 --epochs=100 --all_plot_folder="results/pytorch_learning_curves-carp" --all_result_folder="results/pytorch-carp" --manual_layer_min="${i}" --manual_layer_max="${i}"
#     python run_protran_pytorch.py --dataset_path="data/structure/secondary_structure/tape_ss3_processed.csv" --encoder_name="carp_600k" --checkpoint=0.25 --embed_batch_size=64 --embed_folder="embeddings" --loader_batch_size=120 --epochs=100 --all_plot_folder="results/pytorch_learning_curves-carp" --all_result_folder="results/pytorch-carp" --manual_layer_min="${i}" --manual_layer_max="${i}"
#     python run_protran_pytorch.py --dataset_path="data/structure/secondary_structure/tape_ss3_processed.csv" --encoder_name="carp_600k" --checkpoint=0.125 --embed_batch_size=64 --embed_folder="embeddings" --loader_batch_size=120 --epochs=100 --all_plot_folder="results/pytorch_learning_curves-carp" --all_result_folder="results/pytorch-carp" --manual_layer_min="${i}" --manual_layer_max="${i}"
# done

# for i in 0 1 2 3 4 5 6 7 8 9 10 11 12 13 14 15 16; do
#     python run_protran_pytorch.py --dataset_path="data/structure/secondary_structure/tape_ss3_processed.csv" --encoder_name="carp_38M" --checkpoint=1 --embed_batch_size=64 --embed_folder="embeddings" --loader_batch_size=120 --epochs=100 --all_plot_folder="results/pytorch_learning_curves-carp" --all_result_folder="results/pytorch-carp" --reset_param=True --manual_layer_min="${i}" --manual_layer_max="${i}"
#     python run_protran_pytorch.py --dataset_path="data/structure/secondary_structure/tape_ss3_processed.csv" --encoder_name="carp_38M" --checkpoint=1 --embed_batch_size=64 --embed_folder="embeddings" --loader_batch_size=120 --epochs=100 --all_plot_folder="results/pytorch_learning_curves-carp" --all_result_folder="results/pytorch-carp" --resample_param=True --manual_layer_min="${i}" --manual_layer_max="${i}"
#     python run_protran_pytorch.py --dataset_path="data/structure/secondary_structure/tape_ss3_processed.csv" --encoder_name="carp_38M" --checkpoint=1 --embed_batch_size=64 --embed_folder="embeddings" --loader_batch_size=120 --epochs=100 --all_plot_folder="results/pytorch_learning_curves-carp" --all_result_folder="results/pytorch-carp" --manual_layer_min="${i}" --manual_layer_max="${i}"
#     python run_protran_pytorch.py --dataset_path="data/structure/secondary_structure/tape_ss3_processed.csv" --encoder_name="carp_38M" --checkpoint=0.5 --embed_batch_size=64 --embed_folder="embeddings" --loader_batch_size=120 --epochs=100 --all_plot_folder="results/pytorch_learning_curves-carp" --all_result_folder="results/pytorch-carp" --manual_layer_min="${i}" --manual_layer_max="${i}"
#     python run_protran_pytorch.py --dataset_path="data/structure/secondary_structure/tape_ss3_processed.csv" --encoder_name="carp_38M" --checkpoint=0.25 --embed_batch_size=64 --embed_folder="embeddings" --loader_batch_size=120 --epochs=100 --all_plot_folder="results/pytorch_learning_curves-carp" --all_result_folder="results/pytorch-carp" --manual_layer_min="${i}" --manual_layer_max="${i}"
#     python run_protran_pytorch.py --dataset_path="data/structure/secondary_structure/tape_ss3_processed.csv" --encoder_name="carp_38M" --checkpoint=0.125 --embed_batch_size=64 --embed_folder="embeddings" --loader_batch_size=120 --epochs=100 --all_plot_folder="results/pytorch_learning_curves-carp" --all_result_folder="results/pytorch-carp" --manual_layer_min="${i}" --manual_layer_max="${i}"
# done

# for i in 0 1 2 3 4 5 6 7 8 9 10 11 12 13 14 15 16 17 18 19 20 21 22 23 24 25 26 27 28 29 30 31 32; do
#     python run_protran_pytorch.py --dataset_path="data/structure/secondary_structure/tape_ss3_processed.csv" --encoder_name="carp_76M" --checkpoint=1 --embed_batch_size=64 --embed_folder="embeddings" --loader_batch_size=120 --epochs=100 --all_plot_folder="results/pytorch_learning_curves-carp" --all_result_folder="results/pytorch-carp" --reset_param=True --manual_layer_min="${i}" --manual_layer_max="${i}"
#     python run_protran_pytorch.py --dataset_path="data/structure/secondary_structure/tape_ss3_processed.csv" --encoder_name="carp_76M" --checkpoint=1 --embed_batch_size=64 --embed_folder="embeddings" --loader_batch_size=120 --epochs=100 --all_plot_folder="results/pytorch_learning_curves-carp" --all_result_folder="results/pytorch-carp" --resample_param=True --manual_layer_min="${i}" --manual_layer_max="${i}"
#     python run_protran_pytorch.py --dataset_path="data/structure/secondary_structure/tape_ss3_processed.csv" --encoder_name="carp_76M" --checkpoint=1 --embed_batch_size=64 --embed_folder="embeddings" --loader_batch_size=120 --epochs=100 --all_plot_folder="results/pytorch_learning_curves-carp" --all_result_folder="results/pytorch-carp" --manual_layer_min="${i}" --manual_layer_max="${i}"
#     python run_protran_pytorch.py --dataset_path="data/structure/secondary_structure/tape_ss3_processed.csv" --encoder_name="carp_76M" --checkpoint=0.5 --embed_batch_size=64 --embed_folder="embeddings" --loader_batch_size=120 --epochs=100 --all_plot_folder="results/pytorch_learning_curves-carp" --all_result_folder="results/pytorch-carp" --manual_layer_min="${i}" --manual_layer_max="${i}"
#     python run_protran_pytorch.py --dataset_path="data/structure/secondary_structure/tape_ss3_processed.csv" --encoder_name="carp_76M" --checkpoint=0.25 --embed_batch_size=64 --embed_folder="embeddings" --loader_batch_size=120 --epochs=100 --all_plot_folder="results/pytorch_learning_curves-carp" --all_result_folder="results/pytorch-carp" --manual_layer_min="${i}" --manual_layer_max="${i}"
#     python run_protran_pytorch.py --dataset_path="data/structure/secondary_structure/tape_ss3_processed.csv" --encoder_name="carp_76M" --checkpoint=0.125 --embed_batch_size=64 --embed_folder="embeddings" --loader_batch_size=120 --epochs=100 --all_plot_folder="results/pytorch_learning_curves-carp" --all_result_folder="results/pytorch-carp" --manual_layer_min="${i}" --manual_layer_max="${i}"
# done

# for i in 0 1 2 3 4 5 6 7 8 9 10 11 12 13 14 15 16 17 18 19 20 21 22 23 24 25 26 27 28 29 30 31 32 33 34 35 36 37 38 39 40 41 42 43 44 45 46 47 48 49 50 51 52 53 54 55 56; do
#     python run_protran_pytorch.py --dataset_path="data/structure/secondary_structure/tape_ss3_processed.csv" --encoder_name="carp_640M" --checkpoint=1 --embed_batch_size=64 --embed_folder="embeddings" --loader_batch_size=120 --epochs=100 --all_plot_folder="results/pytorch_learning_curves-carp" --all_result_folder="results/pytorch-carp" --reset_param=True --manual_layer_min="${i}" --manual_layer_max="${i}"
#     python run_protran_pytorch.py --dataset_path="data/structure/secondary_structure/tape_ss3_processed.csv" --encoder_name="carp_640M" --checkpoint=1 --embed_batch_size=64 --embed_folder="embeddings" --loader_batch_size=120 --epochs=100 --all_plot_folder="results/pytorch_learning_curves-carp" --all_result_folder="results/pytorch-carp" --resample_param=True --manual_layer_min="${i}" --manual_layer_max="${i}"
#     python run_protran_pytorch.py --dataset_path="data/structure/secondary_structure/tape_ss3_processed.csv" --encoder_name="carp_640M" --checkpoint=1 --embed_batch_size=64 --embed_folder="embeddings" --loader_batch_size=120 --epochs=100 --all_plot_folder="results/pytorch_learning_curves-carp" --all_result_folder="results/pytorch-carp" --manual_layer_min="${i}" --manual_layer_max="${i}"
#     python run_protran_pytorch.py --dataset_path="data/structure/secondary_structure/tape_ss3_processed.csv" --encoder_name="carp_640M" --checkpoint=0.5 --embed_batch_size=64 --embed_folder="embeddings" --loader_batch_size=100 --epochs=100 --all_plot_folder="results/pytorch_learning_curves-carp" --all_result_folder="results/pytorch-carp" --manual_layer_min="${i}" --manual_layer_max="${i}"
#     python run_protran_pytorch.py --dataset_path="data/structure/secondary_structure/tape_ss3_processed.csv" --encoder_name="carp_640M" --checkpoint=0.25 --embed_batch_size=64 --embed_folder="embeddings" --loader_batch_size=100 --epochs=100 --all_plot_folder="results/pytorch_learning_curves-carp" --all_result_folder="results/pytorch-carp" --manual_layer_min="${i}" --manual_layer_max="${i}"
#     python run_protran_pytorch.py --dataset_path="data/structure/secondary_structure/tape_ss3_processed.csv" --encoder_name="carp_640M" --checkpoint=0.125 --embed_batch_size=64 --embed_folder="embeddings" --loader_batch_size=100 --epochs=100 --all_plot_folder="results/pytorch_learning_curves-carp" --all_result_folder="results/pytorch-carp" --manual_layer_min="${i}" --manual_layer_max="${i}"
# done
```
