## Supplemental data and code for "Feature Reuse and Scaling: Understanding Transfer Learning with Protein Language Models": execute_run_pytorch_esm.sh

```
# queue for running all pytorch models from embs

# export CUDA_VISIBLE_DEVICES=""
export CUDA_VISIBLE_DEVICES=1

### scl ###

# python run_protran_pytorch.py --dataset_path="data/annotation/scl/balanced.csv" --encoder_name="esm1_t6_43M_UR50S" --checkpoint=1 --embed_batch_size=64 --flatten_emb="mean" --embed_folder="embeddings" --loader_batch_size=256 --epochs=100 --all_plot_folder="results/pytorch_learning_curves-esm" --all_result_folder="results/pytorch-esm"
# python run_protran_pytorch.py --dataset_path="data/annotation/scl/balanced.csv" --encoder_name="esm1_t6_43M_UR50S" --checkpoint=1 --embed_batch_size=64 --flatten_emb="mean" --embed_folder="embeddings" --loader_batch_size=256 --epochs=100 --all_plot_folder="results/pytorch_learning_curves-esm" --all_result_folder="results/pytorch-esm" --reset_param=True
# python run_protran_pytorch.py --dataset_path="data/annotation/scl/balanced.csv" --encoder_name="esm1_t6_43M_UR50S" --checkpoint=1 --embed_batch_size=64 --flatten_emb="mean" --embed_folder="embeddings" --loader_batch_size=256 --epochs=100 --all_plot_folder="results/pytorch_learning_curves-esm" --all_result_folder="results/pytorch-esm" --resample_param=True

# python run_protran_pytorch.py --dataset_path="data/annotation/scl/balanced.csv" --encoder_name="esm1_t12_85M_UR50S" --checkpoint=1 --embed_batch_size=64 --flatten_emb="mean" --embed_folder="embeddings" --loader_batch_size=256 --epochs=100 --all_plot_folder="results/pytorch_learning_curves-esm" --all_result_folder="results/pytorch-esm"
# python run_protran_pytorch.py --dataset_path="data/annotation/scl/balanced.csv" --encoder_name="esm1_t12_85M_UR50S" --checkpoint=1 --embed_batch_size=64 --flatten_emb="mean" --embed_folder="embeddings" --loader_batch_size=256 --epochs=100 --all_plot_folder="results/pytorch_learning_curves-esm" --all_result_folder="results/pytorch-esm" --reset_param=True
# python run_protran_pytorch.py --dataset_path="data/annotation/scl/balanced.csv" --encoder_name="esm1_t12_85M_UR50S" --checkpoint=1 --embed_batch_size=64 --flatten_emb="mean" --embed_folder="embeddings" --loader_batch_size=256 --epochs=100 --all_plot_folder="results/pytorch_learning_curves-esm" --all_result_folder="results/pytorch-esm" --resample_param=True

# python run_protran_pytorch.py --dataset_path="data/annotation/scl/balanced.csv" --encoder_name="esm1_t34_670M_UR50S" --checkpoint=1 --embed_batch_size=64 --flatten_emb="mean" --embed_folder="embeddings" --loader_batch_size=256 --epochs=100 --all_plot_folder="results/pytorch_learning_curves-esm" --all_result_folder="results/pytorch-esm"
# python run_protran_pytorch.py --dataset_path="data/annotation/scl/balanced.csv" --encoder_name="esm1_t34_670M_UR50S" --checkpoint=1 --embed_batch_size=64 --flatten_emb="mean" --embed_folder="embeddings" --loader_batch_size=256 --epochs=100 --all_plot_folder="results/pytorch_learning_curves-esm" --all_result_folder="results/pytorch-esm" --reset_param=True &
# python run_protran_pytorch.py --dataset_path="data/annotation/scl/balanced.csv" --encoder_name="esm1_t34_670M_UR50S" --checkpoint=1 --embed_batch_size=64 --flatten_emb="mean" --embed_folder="embeddings" --loader_batch_size=256 --epochs=100 --all_plot_folder="results/pytorch_learning_curves-esm" --all_result_folder="results/pytorch-esm" --resample_param=True &

# python run_protran_pytorch.py --dataset_path="data/annotation/scl/balanced.csv" --encoder_name="esm1b_t33_650M_UR50S" --checkpoint=1 --embed_batch_size=64 --flatten_emb="mean" --embed_folder="embeddings" --loader_batch_size=256 --epochs=100 --all_plot_folder="results/pytorch_learning_curves-esm" --all_result_folder="results/pytorch-esm"
# python run_protran_pytorch.py --dataset_path="data/annotation/scl/balanced.csv" --encoder_name="esm1b_t33_650M_UR50S" --checkpoint=1 --embed_batch_size=64 --flatten_emb="mean" --embed_folder="embeddings" --loader_batch_size=256 --epochs=100 --all_plot_folder="results/pytorch_learning_curves-esm" --all_result_folder="results/pytorch-esm" --reset_param=True &
# python run_protran_pytorch.py --dataset_path="data/annotation/scl/balanced.csv" --encoder_name="esm1b_t33_650M_UR50S" --checkpoint=1 --embed_batch_size=64 --flatten_emb="mean" --embed_folder="embeddings" --loader_batch_size=256 --epochs=100 --all_plot_folder="results/pytorch_learning_curves-esm" --all_result_folder="results/pytorch-esm" --resample_param=True &

### ss3 ###

# python run_protran_pytorch.py --dataset_path="data/structure/secondary_structure/tape_ss3_processed.csv" --encoder_name="" --checkpoint=1 --embed_batch_size=64 --loader_batch_size=120 --epochs=100 --all_plot_folder="results/pytorch_learning_curves-esm" --all_result_folder="results/pytorch-esm" --if_encode_all=True

# for i in 0 1 2 3 4 5 6; do
    # python run_protran_pytorch.py --dataset_path="data/structure/secondary_structure/tape_ss3_processed.csv" --encoder_name="esm1_t6_43M_UR50S" --checkpoint=1 --embed_batch_size=64 --embed_folder="embeddings" --loader_batch_size=120 --epochs=100 --all_plot_folder="results/pytorch_learning_curves-esm" --all_result_folder="results/pytorch-esm" --manual_layer_min="${i}" --manual_layer_max="${i}"
    # python run_protran_pytorch.py --dataset_path="data/structure/secondary_structure/tape_ss3_processed.csv" --encoder_name="esm1_t6_43M_UR50S" --checkpoint=1 --embed_batch_size=64 --embed_folder="embeddings" --loader_batch_size=120 --epochs=100 --all_plot_folder="results/pytorch_learning_curves-esm" --all_result_folder="results/pytorch-esm" --reset_param=True --manual_layer_min="${i}" --manual_layer_max="${i}"
    # python run_protran_pytorch.py --dataset_path="data/structure/secondary_structure/tape_ss3_processed.csv" --encoder_name="esm1_t6_43M_UR50S" --checkpoint=1 --embed_batch_size=64 --embed_folder="embeddings" --loader_batch_size=120 --epochs=100 --all_plot_folder="results/pytorch_learning_curves-esm" --all_result_folder="results/pytorch-esm" --resample_param=True --manual_layer_min="${i}" --manual_layer_max="${i}"
# done

# for i in 0 1 2 3 4 5 6 7 8 9 10 11 12; do
#     python run_protran_pytorch.py --dataset_path="data/structure/secondary_structure/tape_ss3_processed.csv" --encoder_name="esm1_t12_85M_UR50S" --checkpoint=1 --embed_batch_size=64 --embed_folder="embeddings" --loader_batch_size=120 --epochs=100 --all_plot_folder="results/pytorch_learning_curves-esm" --all_result_folder="results/pytorch-esm" --manual_layer_min="${i}" --manual_layer_max="${i}"
#     python run_protran_pytorch.py --dataset_path="data/structure/secondary_structure/tape_ss3_processed.csv" --encoder_name="esm1_t12_85M_UR50S" --checkpoint=1 --embed_batch_size=64 --embed_folder="embeddings" --loader_batch_size=120 --epochs=100 --all_plot_folder="results/pytorch_learning_curves-esm" --all_result_folder="results/pytorch-esm" --reset_param=True --manual_layer_min="${i}" --manual_layer_max="${i}"
#     python run_protran_pytorch.py --dataset_path="data/structure/secondary_structure/tape_ss3_processed.csv" --encoder_name="esm1_t12_85M_UR50S" --checkpoint=1 --embed_batch_size=64 --embed_folder="embeddings" --loader_batch_size=120 --epochs=100 --all_plot_folder="results/pytorch_learning_curves-esm" --all_result_folder="results/pytorch-esm" --resample_param=True --manual_layer_min="${i}" --manual_layer_max="${i}"
# done

# for i in 0 1 2 3 4 5 6 7 8 9 10 11 12 13 14 15 16 17 18 19 20 21 22 23 24 25 26 27 28 29 30 31 32 33 34; do
#     python run_protran_pytorch.py --dataset_path="data/structure/secondary_structure/tape_ss3_processed.csv" --encoder_name="esm1_t34_670M_UR50S" --checkpoint=1 --embed_batch_size=64 --embed_folder="embeddings" --loader_batch_size=120 --epochs=100 --all_plot_folder="results/pytorch_learning_curves-esm" --all_result_folder="results/pytorch-esm" --manual_layer_min="${i}" --manual_layer_max="${i}"
#     python run_protran_pytorch.py --dataset_path="data/structure/secondary_structure/tape_ss3_processed.csv" --encoder_name="esm1_t34_670M_UR50S" --checkpoint=1 --embed_batch_size=64 --embed_folder="embeddings" --loader_batch_size=120 --epochs=100 --all_plot_folder="results/pytorch_learning_curves-esm" --all_result_folder="results/pytorch-esm" --reset_param=True --manual_layer_min="${i}" --manual_layer_max="${i}"
#     python run_protran_pytorch.py --dataset_path="data/structure/secondary_structure/tape_ss3_processed.csv" --encoder_name="esm1_t34_670M_UR50S" --checkpoint=1 --embed_batch_size=64 --embed_folder="embeddings" --loader_batch_size=120 --epochs=100 --all_plot_folder="results/pytorch_learning_curves-esm" --all_result_folder="results/pytorch-esm" --resample_param=True --manual_layer_min="${i}" --manual_layer_max="${i}"
# done

# for i in 0 1 2 3 4 5 6 7 8 9 10 11 12 13 14 15 16 17 18 19 20 21 22 23 24 25 26 27 28 29 30 31 32 33; do
#     python run_protran_pytorch.py --dataset_path="data/structure/secondary_structure/tape_ss3_processed.csv" --encoder_name="esm1b_t33_650M_UR50S" --checkpoint=1 --embed_batch_size=64 --embed_folder="embeddings" --loader_batch_size=120 --epochs=100 --all_plot_folder="results/pytorch_learning_curves-esm" --all_result_folder="results/pytorch-esm" --manual_layer_min="${i}" --manual_layer_max="${i}"
#     python run_protran_pytorch.py --dataset_path="data/structure/secondary_structure/tape_ss3_processed.csv" --encoder_name="esm1b_t33_650M_UR50S" --checkpoint=1 --embed_batch_size=64 --embed_folder="embeddings" --loader_batch_size=120 --epochs=100 --all_plot_folder="results/pytorch_learning_curves-esm" --all_result_folder="results/pytorch-esm" --reset_param=True --manual_layer_min="${i}" --manual_layer_max="${i}"
#     python run_protran_pytorch.py --dataset_path="data/structure/secondary_structure/tape_ss3_processed.csv" --encoder_name="esm1b_t33_650M_UR50S" --checkpoint=1 --embed_batch_size=64 --embed_folder="embeddings" --loader_batch_size=120 --epochs=100 --all_plot_folder="results/pytorch_learning_curves-esm" --all_result_folder="results/pytorch-esm" --resample_param=True --manual_layer_min="${i}" --manual_layer_max="${i}"
# done
```
