## Supplemental data and code for "Feature Reuse and Scaling: Understanding Transfer Learning with Protein Language Models": execute_run_sklearn_carp.sh

```
# queue for running all sklearn models from embs

### GB1 ###

# python run_protran_sklearn.py --dataset_path="data/proeng/gb1/sampled.csv" --encoder_name="carp_600k" --checkpoint=1 --embed_batch_size=64 --flatten_emb="mean" --embed_folder="embeddings" --seq_end_idx=56 --all_embed_layers=True --sklearn_params='{"normalize":true}' --all_result_folder="results/sklearn-carp" --reset_param=True
# python run_protran_sklearn.py --dataset_path="data/proeng/gb1/sampled.csv" --encoder_name="carp_600k" --checkpoint=1 --embed_batch_size=64 --flatten_emb="mean" --embed_folder="embeddings" --seq_end_idx=56 --all_embed_layers=True --sklearn_params='{"normalize":true}' --all_result_folder="results/sklearn-carp" --resample_param=True
# python run_protran_sklearn.py --dataset_path="data/proeng/gb1/sampled.csv" --encoder_name="carp_600k" --checkpoint=1 --embed_batch_size=64 --flatten_emb="mean" --embed_folder="embeddings" --seq_end_idx=56 --all_embed_layers=True --sklearn_params='{"normalize":true}' --all_result_folder="results/sklearn-carp"
# python run_protran_sklearn.py --dataset_path="data/proeng/gb1/sampled.csv" --encoder_name="carp_600k" --checkpoint=0.5 --embed_batch_size=64 --flatten_emb="mean" --embed_folder="embeddings" --seq_end_idx=56 --all_embed_layers=True --sklearn_params='{"normalize":true}' --all_result_folder="results/sklearn-carp"
# python run_protran_sklearn.py --dataset_path="data/proeng/gb1/sampled.csv" --encoder_name="carp_600k" --checkpoint=0.25 --embed_batch_size=64 --flatten_emb="mean" --embed_folder="embeddings" --seq_end_idx=56 --all_embed_layers=True --sklearn_params='{"normalize":true}' --all_result_folder="results/sklearn-carp"
# python run_protran_sklearn.py --dataset_path="data/proeng/gb1/sampled.csv" --encoder_name="carp_600k" --checkpoint=0.125 --embed_batch_size=64 --flatten_emb="mean" --embed_folder="embeddings" --seq_end_idx=56 --all_embed_layers=True --sklearn_params='{"normalize":true}' --all_result_folder="results/sklearn-carp"

# python run_protran_sklearn.py --dataset_path="data/proeng/gb1/sampled.csv" --encoder_name="carp_38M" --checkpoint=1 --embed_batch_size=64 --flatten_emb="mean" --embed_folder="embeddings" --seq_end_idx=56 --all_embed_layers=True --sklearn_params='{"normalize":true}' --all_result_folder="results/sklearn-carp" --reset_param=True
# python run_protran_sklearn.py --dataset_path="data/proeng/gb1/sampled.csv" --encoder_name="carp_38M" --checkpoint=1 --embed_batch_size=64 --flatten_emb="mean" --embed_folder="embeddings" --seq_end_idx=56 --all_embed_layers=True --sklearn_params='{"normalize":true}' --all_result_folder="results/sklearn-carp" --resample_param=True
# python run_protran_sklearn.py --dataset_path="data/proeng/gb1/sampled.csv" --encoder_name="carp_38M" --checkpoint=1 --embed_batch_size=64 --flatten_emb="mean" --embed_folder="embeddings" --seq_end_idx=56 --all_embed_layers=True --sklearn_params='{"normalize":true}' --all_result_folder="results/sklearn-carp"
# python run_protran_sklearn.py --dataset_path="data/proeng/gb1/sampled.csv" --encoder_name="carp_38M" --checkpoint=0.5 --embed_batch_size=64 --flatten_emb="mean" --embed_folder="embeddings" --seq_end_idx=56 --all_embed_layers=True --sklearn_params='{"normalize":true}' --all_result_folder="results/sklearn-carp"
# python run_protran_sklearn.py --dataset_path="data/proeng/gb1/sampled.csv" --encoder_name="carp_38M" --checkpoint=0.25 --embed_batch_size=64 --flatten_emb="mean" --embed_folder="embeddings" --seq_end_idx=56 --all_embed_layers=True --sklearn_params='{"normalize":true}' --all_result_folder="results/sklearn-carp"
# python run_protran_sklearn.py --dataset_path="data/proeng/gb1/sampled.csv" --encoder_name="carp_38M" --checkpoint=0.125 --embed_batch_size=64 --flatten_emb="mean" --embed_folder="embeddings" --seq_end_idx=56 --all_embed_layers=True --sklearn_params='{"normalize":true}' --all_result_folder="results/sklearn-carp"

# python run_protran_sklearn.py --dataset_path="data/proeng/gb1/sampled.csv" --encoder_name="carp_76M" --checkpoint=1 --embed_batch_size=64 --flatten_emb="mean" --embed_folder="embeddings" --seq_end_idx=56 --all_embed_layers=True --sklearn_params='{"normalize":true}' --all_result_folder="results/sklearn-carp" --reset_param=True
# python run_protran_sklearn.py --dataset_path="data/proeng/gb1/sampled.csv" --encoder_name="carp_76M" --checkpoint=1 --embed_batch_size=64 --flatten_emb="mean" --embed_folder="embeddings" --seq_end_idx=56 --all_embed_layers=True --sklearn_params='{"normalize":true}' --all_result_folder="results/sklearn-carp" --resample_param=True
# python run_protran_sklearn.py --dataset_path="data/proeng/gb1/sampled.csv" --encoder_name="carp_76M" --checkpoint=1 --embed_batch_size=64 --flatten_emb="mean" --embed_folder="embeddings" --seq_end_idx=56 --all_embed_layers=True --sklearn_params='{"normalize":true}' --all_result_folder="results/sklearn-carp"
# python run_protran_sklearn.py --dataset_path="data/proeng/gb1/sampled.csv" --encoder_name="carp_76M" --checkpoint=0.5 --embed_batch_size=64 --flatten_emb="mean" --embed_folder="embeddings" --seq_end_idx=56 --all_embed_layers=True --sklearn_params='{"normalize":true}' --all_result_folder="results/sklearn-carp"
# python run_protran_sklearn.py --dataset_path="data/proeng/gb1/sampled.csv" --encoder_name="carp_76M" --checkpoint=0.25 --embed_batch_size=64 --flatten_emb="mean" --embed_folder="embeddings" --seq_end_idx=56 --all_embed_layers=True --sklearn_params='{"normalize":true}' --all_result_folder="results/sklearn-carp"
# python run_protran_sklearn.py --dataset_path="data/proeng/gb1/sampled.csv" --encoder_name="carp_76M" --checkpoint=0.125 --embed_batch_size=64 --flatten_emb="mean" --embed_folder="embeddings" --seq_end_idx=56 --all_embed_layers=True --sklearn_params='{"normalize":true}' --all_result_folder="results/sklearn-carp"

# python run_protran_sklearn.py --dataset_path="data/proeng/gb1/sampled.csv" --encoder_name="carp_640M" --checkpoint=1 --embed_batch_size=64 --flatten_emb="mean" --embed_folder="embeddings" --seq_end_idx=56 --all_embed_layers=True --sklearn_params='{"normalize":true}' --all_result_folder="results/sklearn-carp" --reset_param=True
# python run_protran_sklearn.py --dataset_path="data/proeng/gb1/sampled.csv" --encoder_name="carp_640M" --checkpoint=1 --embed_batch_size=64 --flatten_emb="mean" --embed_folder="embeddings" --seq_end_idx=56 --all_embed_layers=True --sklearn_params='{"normalize":true}' --all_result_folder="results/sklearn-carp" --resample_param=True
# python run_protran_sklearn.py --dataset_path="data/proeng/gb1/sampled.csv" --encoder_name="carp_640M" --checkpoint=1 --embed_batch_size=64 --flatten_emb="mean" --embed_folder="embeddings" --seq_end_idx=56 --all_embed_layers=True --sklearn_params='{"normalize":true}' --all_result_folder="results/sklearn-carp"
# python run_protran_sklearn.py --dataset_path="data/proeng/gb1/sampled.csv" --encoder_name="carp_640M" --checkpoint=0.5 --embed_batch_size=64 --flatten_emb="mean" --embed_folder="embeddings" --seq_end_idx=56 --all_embed_layers=True --sklearn_params='{"normalize":true}' --all_result_folder="results/sklearn-carp"
# python run_protran_sklearn.py --dataset_path="data/proeng/gb1/sampled.csv" --encoder_name="carp_640M" --checkpoint=0.25 --embed_batch_size=64 --flatten_emb="mean" --embed_folder="embeddings" --seq_end_idx=56 --all_embed_layers=True --sklearn_params='{"normalize":true}' --all_result_folder="results/sklearn-carp"
# python run_protran_sklearn.py --dataset_path="data/proeng/gb1/sampled.csv" --encoder_name="carp_640M" --checkpoint=0.125 --embed_batch_size=64 --flatten_emb="mean" --embed_folder="embeddings" --seq_end_idx=56 --all_embed_layers=True --sklearn_params='{"normalize":true}' --all_result_folder="results/sklearn-carp"


# python run_protran_sklearn.py --dataset_path="data/proeng/gb1/low_vs_high.csv" --encoder_name="carp_600k" --checkpoint=1 --embed_batch_size=64 --flatten_emb="mean" --embed_folder="embeddings" --seq_end_idx=56 --all_embed_layers=True --sklearn_params='{"normalize":true}' --all_result_folder="results/sklearn-carp" --reset_param=True
# python run_protran_sklearn.py --dataset_path="data/proeng/gb1/low_vs_high.csv" --encoder_name="carp_600k" --checkpoint=1 --embed_batch_size=64 --flatten_emb="mean" --embed_folder="embeddings" --seq_end_idx=56 --all_embed_layers=True --sklearn_params='{"normalize":true}' --all_result_folder="results/sklearn-carp" --resample_param=True
# python run_protran_sklearn.py --dataset_path="data/proeng/gb1/low_vs_high.csv" --encoder_name="carp_600k" --checkpoint=1 --embed_batch_size=64 --flatten_emb="mean" --embed_folder="embeddings" --seq_end_idx=56 --all_embed_layers=True --sklearn_params='{"normalize":true}' --all_result_folder="results/sklearn-carp"
# python run_protran_sklearn.py --dataset_path="data/proeng/gb1/low_vs_high.csv" --encoder_name="carp_600k" --checkpoint=0.5 --embed_batch_size=64 --flatten_emb="mean" --embed_folder="embeddings" --seq_end_idx=56 --all_embed_layers=True --sklearn_params='{"normalize":true}' --all_result_folder="results/sklearn-carp"
# python run_protran_sklearn.py --dataset_path="data/proeng/gb1/low_vs_high.csv" --encoder_name="carp_600k" --checkpoint=0.25 --embed_batch_size=64 --flatten_emb="mean" --embed_folder="embeddings" --seq_end_idx=56 --all_embed_layers=True --sklearn_params='{"normalize":true}' --all_result_folder="results/sklearn-carp"
# python run_protran_sklearn.py --dataset_path="data/proeng/gb1/low_vs_high.csv" --encoder_name="carp_600k" --checkpoint=0.125 --embed_batch_size=64 --flatten_emb="mean" --embed_folder="embeddings" --seq_end_idx=56 --all_embed_layers=True --sklearn_params='{"normalize":true}' --all_result_folder="results/sklearn-carp"

# python run_protran_sklearn.py --dataset_path="data/proeng/gb1/low_vs_high.csv" --encoder_name="carp_38M" --checkpoint=1 --embed_batch_size=64 --flatten_emb="mean" --embed_folder="embeddings" --seq_end_idx=56 --all_embed_layers=True --sklearn_params='{"normalize":true}' --all_result_folder="results/sklearn-carp" --reset_param=True
# python run_protran_sklearn.py --dataset_path="data/proeng/gb1/low_vs_high.csv" --encoder_name="carp_38M" --checkpoint=1 --embed_batch_size=64 --flatten_emb="mean" --embed_folder="embeddings" --seq_end_idx=56 --all_embed_layers=True --sklearn_params='{"normalize":true}' --all_result_folder="results/sklearn-carp" --resample_param=True
# python run_protran_sklearn.py --dataset_path="data/proeng/gb1/low_vs_high.csv" --encoder_name="carp_38M" --checkpoint=1 --embed_batch_size=64 --flatten_emb="mean" --embed_folder="embeddings" --seq_end_idx=56 --all_embed_layers=True --sklearn_params='{"normalize":true}' --all_result_folder="results/sklearn-carp"
# python run_protran_sklearn.py --dataset_path="data/proeng/gb1/low_vs_high.csv" --encoder_name="carp_38M" --checkpoint=0.5 --embed_batch_size=64 --flatten_emb="mean" --embed_folder="embeddings" --seq_end_idx=56 --all_embed_layers=True --sklearn_params='{"normalize":true}' --all_result_folder="results/sklearn-carp"
# python run_protran_sklearn.py --dataset_path="data/proeng/gb1/low_vs_high.csv" --encoder_name="carp_38M" --checkpoint=0.25 --embed_batch_size=64 --flatten_emb="mean" --embed_folder="embeddings" --seq_end_idx=56 --all_embed_layers=True --sklearn_params='{"normalize":true}' --all_result_folder="results/sklearn-carp"
# python run_protran_sklearn.py --dataset_path="data/proeng/gb1/low_vs_high.csv" --encoder_name="carp_38M" --checkpoint=0.125 --embed_batch_size=64 --flatten_emb="mean" --embed_folder="embeddings" --seq_end_idx=56 --all_embed_layers=True --sklearn_params='{"normalize":true}' --all_result_folder="results/sklearn-carp"

# python run_protran_sklearn.py --dataset_path="data/proeng/gb1/low_vs_high.csv" --encoder_name="carp_76M" --checkpoint=1 --embed_batch_size=64 --flatten_emb="mean" --embed_folder="embeddings" --seq_end_idx=56 --all_embed_layers=True --sklearn_params='{"normalize":true}' --all_result_folder="results/sklearn-carp" --reset_param=True
# python run_protran_sklearn.py --dataset_path="data/proeng/gb1/low_vs_high.csv" --encoder_name="carp_76M" --checkpoint=1 --embed_batch_size=64 --flatten_emb="mean" --embed_folder="embeddings" --seq_end_idx=56 --all_embed_layers=True --sklearn_params='{"normalize":true}' --all_result_folder="results/sklearn-carp" --resample_param=True
# python run_protran_sklearn.py --dataset_path="data/proeng/gb1/low_vs_high.csv" --encoder_name="carp_76M" --checkpoint=1 --embed_batch_size=64 --flatten_emb="mean" --embed_folder="embeddings" --seq_end_idx=56 --all_embed_layers=True --sklearn_params='{"normalize":true}' --all_result_folder="results/sklearn-carp"
# python run_protran_sklearn.py --dataset_path="data/proeng/gb1/low_vs_high.csv" --encoder_name="carp_76M" --checkpoint=0.5 --embed_batch_size=64 --flatten_emb="mean" --embed_folder="embeddings" --seq_end_idx=56 --all_embed_layers=True --sklearn_params='{"normalize":true}' --all_result_folder="results/sklearn-carp"
# python run_protran_sklearn.py --dataset_path="data/proeng/gb1/low_vs_high.csv" --encoder_name="carp_76M" --checkpoint=0.25 --embed_batch_size=64 --flatten_emb="mean" --embed_folder="embeddings" --seq_end_idx=56 --all_embed_layers=True --sklearn_params='{"normalize":true}' --all_result_folder="results/sklearn-carp"
# python run_protran_sklearn.py --dataset_path="data/proeng/gb1/low_vs_high.csv" --encoder_name="carp_76M" --checkpoint=0.125 --embed_batch_size=64 --flatten_emb="mean" --embed_folder="embeddings" --seq_end_idx=56 --all_embed_layers=True --sklearn_params='{"normalize":true}' --all_result_folder="results/sklearn-carp"

# python run_protran_sklearn.py --dataset_path="data/proeng/gb1/low_vs_high.csv" --encoder_name="carp_640M" --checkpoint=1 --embed_batch_size=64 --flatten_emb="mean" --embed_folder="embeddings" --seq_end_idx=56 --all_embed_layers=True --sklearn_params='{"normalize":true}' --all_result_folder="results/sklearn-carp" --reset_param=True
# python run_protran_sklearn.py --dataset_path="data/proeng/gb1/low_vs_high.csv" --encoder_name="carp_640M" --checkpoint=1 --embed_batch_size=64 --flatten_emb="mean" --embed_folder="embeddings" --seq_end_idx=56 --all_embed_layers=True --sklearn_params='{"normalize":true}' --all_result_folder="results/sklearn-carp" --resample_param=True
# python run_protran_sklearn.py --dataset_path="data/proeng/gb1/low_vs_high.csv" --encoder_name="carp_640M" --checkpoint=1 --embed_batch_size=64 --flatten_emb="mean" --embed_folder="embeddings" --seq_end_idx=56 --all_embed_layers=True --sklearn_params='{"normalize":true}' --all_result_folder="results/sklearn-carp"
# python run_protran_sklearn.py --dataset_path="data/proeng/gb1/low_vs_high.csv" --encoder_name="carp_640M" --checkpoint=0.5 --embed_batch_size=64 --flatten_emb="mean" --embed_folder="embeddings" --seq_end_idx=56 --all_embed_layers=True --sklearn_params='{"normalize":true}' --all_result_folder="results/sklearn-carp"
# python run_protran_sklearn.py --dataset_path="data/proeng/gb1/low_vs_high.csv" --encoder_name="carp_640M" --checkpoint=0.25 --embed_batch_size=64 --flatten_emb="mean" --embed_folder="embeddings" --seq_end_idx=56 --all_embed_layers=True --sklearn_params='{"normalize":true}' --all_result_folder="results/sklearn-carp"
# python run_protran_sklearn.py --dataset_path="data/proeng/gb1/low_vs_high.csv" --encoder_name="carp_640M" --checkpoint=0.125 --embed_batch_size=64 --flatten_emb="mean" --embed_folder="embeddings" --seq_end_idx=56 --all_embed_layers=True --sklearn_params='{"normalize":true}' --all_result_folder="results/sklearn-carp"


# python run_protran_sklearn.py --dataset_path="data/proeng/gb1/two_vs_rest.csv" --encoder_name="carp_600k" --checkpoint=1 --embed_batch_size=64 --flatten_emb="mean" --embed_folder="embeddings" --seq_end_idx=56 --all_embed_layers=True --sklearn_params='{"normalize":true}' --all_result_folder="results/sklearn-carp" --reset_param=True
# python run_protran_sklearn.py --dataset_path="data/proeng/gb1/two_vs_rest.csv" --encoder_name="carp_600k" --checkpoint=1 --embed_batch_size=64 --flatten_emb="mean" --embed_folder="embeddings" --seq_end_idx=56 --all_embed_layers=True --sklearn_params='{"normalize":true}' --all_result_folder="results/sklearn-carp" --resample_param=True
# python run_protran_sklearn.py --dataset_path="data/proeng/gb1/two_vs_rest.csv" --encoder_name="carp_600k" --checkpoint=1 --embed_batch_size=64 --flatten_emb="mean" --embed_folder="embeddings" --seq_end_idx=56 --all_embed_layers=True --sklearn_params='{"normalize":true}' --all_result_folder="results/sklearn-carp"
# python run_protran_sklearn.py --dataset_path="data/proeng/gb1/two_vs_rest.csv" --encoder_name="carp_600k" --checkpoint=0.5 --embed_batch_size=64 --flatten_emb="mean" --embed_folder="embeddings" --seq_end_idx=56 --all_embed_layers=True --sklearn_params='{"normalize":true}' --all_result_folder="results/sklearn-carp"
# python run_protran_sklearn.py --dataset_path="data/proeng/gb1/two_vs_rest.csv" --encoder_name="carp_600k" --checkpoint=0.25 --embed_batch_size=64 --flatten_emb="mean" --embed_folder="embeddings" --seq_end_idx=56 --all_embed_layers=True --sklearn_params='{"normalize":true}' --all_result_folder="results/sklearn-carp"
# python run_protran_sklearn.py --dataset_path="data/proeng/gb1/two_vs_rest.csv" --encoder_name="carp_600k" --checkpoint=0.125 --embed_batch_size=64 --flatten_emb="mean" --embed_folder="embeddings" --seq_end_idx=56 --all_embed_layers=True --sklearn_params='{"normalize":true}' --all_result_folder="results/sklearn-carp"

# python run_protran_sklearn.py --dataset_path="data/proeng/gb1/two_vs_rest.csv" --encoder_name="carp_38M" --checkpoint=1 --embed_batch_size=64 --flatten_emb="mean" --embed_folder="embeddings" --seq_end_idx=56 --all_embed_layers=True --sklearn_params='{"normalize":true}' --all_result_folder="results/sklearn-carp" --reset_param=True
# python run_protran_sklearn.py --dataset_path="data/proeng/gb1/two_vs_rest.csv" --encoder_name="carp_38M" --checkpoint=1 --embed_batch_size=64 --flatten_emb="mean" --embed_folder="embeddings" --seq_end_idx=56 --all_embed_layers=True --sklearn_params='{"normalize":true}' --all_result_folder="results/sklearn-carp" --resample_param=True
# python run_protran_sklearn.py --dataset_path="data/proeng/gb1/two_vs_rest.csv" --encoder_name="carp_38M" --checkpoint=1 --embed_batch_size=64 --flatten_emb="mean" --embed_folder="embeddings" --seq_end_idx=56 --all_embed_layers=True --sklearn_params='{"normalize":true}' --all_result_folder="results/sklearn-carp"
# python run_protran_sklearn.py --dataset_path="data/proeng/gb1/two_vs_rest.csv" --encoder_name="carp_38M" --checkpoint=0.5 --embed_batch_size=64 --flatten_emb="mean" --embed_folder="embeddings" --seq_end_idx=56 --all_embed_layers=True --sklearn_params='{"normalize":true}' --all_result_folder="results/sklearn-carp"
# python run_protran_sklearn.py --dataset_path="data/proeng/gb1/two_vs_rest.csv" --encoder_name="carp_38M" --checkpoint=0.25 --embed_batch_size=64 --flatten_emb="mean" --embed_folder="embeddings" --seq_end_idx=56 --all_embed_layers=True --sklearn_params='{"normalize":true}' --all_result_folder="results/sklearn-carp"
# python run_protran_sklearn.py --dataset_path="data/proeng/gb1/two_vs_rest.csv" --encoder_name="carp_38M" --checkpoint=0.125 --embed_batch_size=64 --flatten_emb="mean" --embed_folder="embeddings" --seq_end_idx=56 --all_embed_layers=True --sklearn_params='{"normalize":true}' --all_result_folder="results/sklearn-carp"

# python run_protran_sklearn.py --dataset_path="data/proeng/gb1/two_vs_rest.csv" --encoder_name="carp_76M" --checkpoint=1 --embed_batch_size=64 --flatten_emb="mean" --embed_folder="embeddings" --seq_end_idx=56 --all_embed_layers=True --sklearn_params='{"normalize":true}' --all_result_folder="results/sklearn-carp" --reset_param=True
# python run_protran_sklearn.py --dataset_path="data/proeng/gb1/two_vs_rest.csv" --encoder_name="carp_76M" --checkpoint=1 --embed_batch_size=64 --flatten_emb="mean" --embed_folder="embeddings" --seq_end_idx=56 --all_embed_layers=True --sklearn_params='{"normalize":true}' --all_result_folder="results/sklearn-carp" --resample_param=True
# python run_protran_sklearn.py --dataset_path="data/proeng/gb1/two_vs_rest.csv" --encoder_name="carp_76M" --checkpoint=1 --embed_batch_size=64 --flatten_emb="mean" --embed_folder="embeddings" --seq_end_idx=56 --all_embed_layers=True --sklearn_params='{"normalize":true}' --all_result_folder="results/sklearn-carp"
# python run_protran_sklearn.py --dataset_path="data/proeng/gb1/two_vs_rest.csv" --encoder_name="carp_76M" --checkpoint=0.5 --embed_batch_size=64 --flatten_emb="mean" --embed_folder="embeddings" --seq_end_idx=56 --all_embed_layers=True --sklearn_params='{"normalize":true}' --all_result_folder="results/sklearn-carp"
# python run_protran_sklearn.py --dataset_path="data/proeng/gb1/two_vs_rest.csv" --encoder_name="carp_76M" --checkpoint=0.25 --embed_batch_size=64 --flatten_emb="mean" --embed_folder="embeddings" --seq_end_idx=56 --all_embed_layers=True --sklearn_params='{"normalize":true}' --all_result_folder="results/sklearn-carp"
# python run_protran_sklearn.py --dataset_path="data/proeng/gb1/two_vs_rest.csv" --encoder_name="carp_76M" --checkpoint=0.125 --embed_batch_size=64 --flatten_emb="mean" --embed_folder="embeddings" --seq_end_idx=56 --all_embed_layers=True --sklearn_params='{"normalize":true}' --all_result_folder="results/sklearn-carp"

# python run_protran_sklearn.py --dataset_path="data/proeng/gb1/two_vs_rest.csv" --encoder_name="carp_640M" --checkpoint=1 --embed_batch_size=64 --flatten_emb="mean" --embed_folder="embeddings" --seq_end_idx=56 --all_embed_layers=True --sklearn_params='{"normalize":true}' --all_result_folder="results/sklearn-carp" --reset_param=True
# python run_protran_sklearn.py --dataset_path="data/proeng/gb1/two_vs_rest.csv" --encoder_name="carp_640M" --checkpoint=1 --embed_batch_size=64 --flatten_emb="mean" --embed_folder="embeddings" --seq_end_idx=56 --all_embed_layers=True --sklearn_params='{"normalize":true}' --all_result_folder="results/sklearn-carp" --resample_param=True
# python run_protran_sklearn.py --dataset_path="data/proeng/gb1/two_vs_rest.csv" --encoder_name="carp_640M" --checkpoint=1 --embed_batch_size=64 --flatten_emb="mean" --embed_folder="embeddings" --seq_end_idx=56 --all_embed_layers=True --sklearn_params='{"normalize":true}' --all_result_folder="results/sklearn-carp"
# python run_protran_sklearn.py --dataset_path="data/proeng/gb1/two_vs_rest.csv" --encoder_name="carp_640M" --checkpoint=0.5 --embed_batch_size=64 --flatten_emb="mean" --embed_folder="embeddings" --seq_end_idx=56 --all_embed_layers=True --sklearn_params='{"normalize":true}' --all_result_folder="results/sklearn-carp"
# python run_protran_sklearn.py --dataset_path="data/proeng/gb1/two_vs_rest.csv" --encoder_name="carp_640M" --checkpoint=0.25 --embed_batch_size=64 --flatten_emb="mean" --embed_folder="embeddings" --seq_end_idx=56 --all_embed_layers=True --sklearn_params='{"normalize":true}' --all_result_folder="results/sklearn-carp"
# python run_protran_sklearn.py --dataset_path="data/proeng/gb1/two_vs_rest.csv" --encoder_name="carp_640M" --checkpoint=0.125 --embed_batch_size=64 --flatten_emb="mean" --embed_folder="embeddings" --seq_end_idx=56 --all_embed_layers=True --sklearn_params='{"normalize":true}' --all_result_folder="results/sklearn-carp"


# ### AAV ###

# python run_protran_sklearn.py --dataset_path="data/proeng/aav/two_vs_many.csv" --encoder_name="carp_600k" --checkpoint=1 --embed_batch_size=64 --flatten_emb="mean" --embed_folder="embeddings" --all_embed_layers=True --sklearn_params='{"normalize":true}' --all_result_folder="results/sklearn-carp" --reset_param=True
# python run_protran_sklearn.py --dataset_path="data/proeng/aav/two_vs_many.csv" --encoder_name="carp_600k" --checkpoint=1 --embed_batch_size=64 --flatten_emb="mean" --embed_folder="embeddings" --all_embed_layers=True --sklearn_params='{"normalize":true}' --all_result_folder="results/sklearn-carp" --resample_param=True
# python run_protran_sklearn.py --dataset_path="data/proeng/aav/two_vs_many.csv" --encoder_name="carp_600k" --checkpoint=1 --embed_batch_size=64 --flatten_emb="mean" --embed_folder="embeddings" --all_embed_layers=True --sklearn_params='{"normalize":true}' --all_result_folder="results/sklearn-carp"
# python run_protran_sklearn.py --dataset_path="data/proeng/aav/two_vs_many.csv" --encoder_name="carp_600k" --checkpoint=0.5 --embed_batch_size=64 --flatten_emb="mean" --embed_folder="embeddings" --all_embed_layers=True --sklearn_params='{"normalize":true}' --all_result_folder="results/sklearn-carp"
# python run_protran_sklearn.py --dataset_path="data/proeng/aav/two_vs_many.csv" --encoder_name="carp_600k" --checkpoint=0.25 --embed_batch_size=64 --flatten_emb="mean" --embed_folder="embeddings" --all_embed_layers=True --sklearn_params='{"normalize":true}' --all_result_folder="results/sklearn-carp"
# python run_protran_sklearn.py --dataset_path="data/proeng/aav/two_vs_many.csv" --encoder_name="carp_600k" --checkpoint=0.125 --embed_batch_size=64 --flatten_emb="mean" --embed_folder="embeddings" --all_embed_layers=True --sklearn_params='{"normalize":true}' --all_result_folder="results/sklearn-carp"

# python run_protran_sklearn.py --dataset_path="data/proeng/aav/two_vs_many.csv" --encoder_name="carp_38M" --checkpoint=1 --embed_batch_size=64 --flatten_emb="mean" --embed_folder="embeddings" --all_embed_layers=True --sklearn_params='{"normalize":true}' --all_result_folder="results/sklearn-carp" --reset_param=True
# python run_protran_sklearn.py --dataset_path="data/proeng/aav/two_vs_many.csv" --encoder_name="carp_38M" --checkpoint=1 --embed_batch_size=64 --flatten_emb="mean" --embed_folder="embeddings" --all_embed_layers=True --sklearn_params='{"normalize":true}' --all_result_folder="results/sklearn-carp" --resample_param=True
# python run_protran_sklearn.py --dataset_path="data/proeng/aav/two_vs_many.csv" --encoder_name="carp_38M" --checkpoint=1 --embed_batch_size=64 --flatten_emb="mean" --embed_folder="embeddings" --all_embed_layers=True --sklearn_params='{"normalize":true}' --all_result_folder="results/sklearn-carp"
# python run_protran_sklearn.py --dataset_path="data/proeng/aav/two_vs_many.csv" --encoder_name="carp_38M" --checkpoint=0.5 --embed_batch_size=64 --flatten_emb="mean" --embed_folder="embeddings" --all_embed_layers=True --sklearn_params='{"normalize":true}' --all_result_folder="results/sklearn-carp"
# python run_protran_sklearn.py --dataset_path="data/proeng/aav/two_vs_many.csv" --encoder_name="carp_38M" --checkpoint=0.25 --embed_batch_size=64 --flatten_emb="mean" --embed_folder="embeddings" --all_embed_layers=True --sklearn_params='{"normalize":true}' --all_result_folder="results/sklearn-carp"
# python run_protran_sklearn.py --dataset_path="data/proeng/aav/two_vs_many.csv" --encoder_name="carp_38M" --checkpoint=0.125 --embed_batch_size=64 --flatten_emb="mean" --embed_folder="embeddings" --all_embed_layers=True --sklearn_params='{"normalize":true}' --all_result_folder="results/sklearn-carp"

# python run_protran_sklearn.py --dataset_path="data/proeng/aav/two_vs_many.csv" --encoder_name="carp_76M" --checkpoint=1 --embed_batch_size=64 --flatten_emb="mean" --embed_folder="embeddings" --all_embed_layers=True --sklearn_params='{"normalize":true}' --all_result_folder="results/sklearn-carp" --reset_param=True
# python run_protran_sklearn.py --dataset_path="data/proeng/aav/two_vs_many.csv" --encoder_name="carp_76M" --checkpoint=1 --embed_batch_size=64 --flatten_emb="mean" --embed_folder="embeddings" --all_embed_layers=True --sklearn_params='{"normalize":true}' --all_result_folder="results/sklearn-carp" --resample_param=True
# python run_protran_sklearn.py --dataset_path="data/proeng/aav/two_vs_many.csv" --encoder_name="carp_76M" --checkpoint=1 --embed_batch_size=64 --flatten_emb="mean" --embed_folder="embeddings" --all_embed_layers=True --sklearn_params='{"normalize":true}' --all_result_folder="results/sklearn-carp"
# python run_protran_sklearn.py --dataset_path="data/proeng/aav/two_vs_many.csv" --encoder_name="carp_76M" --checkpoint=0.5 --embed_batch_size=64 --flatten_emb="mean" --embed_folder="embeddings" --all_embed_layers=True --sklearn_params='{"normalize":true}' --all_result_folder="results/sklearn-carp"
# python run_protran_sklearn.py --dataset_path="data/proeng/aav/two_vs_many.csv" --encoder_name="carp_76M" --checkpoint=0.25 --embed_batch_size=64 --flatten_emb="mean" --embed_folder="embeddings" --all_embed_layers=True --sklearn_params='{"normalize":true}' --all_result_folder="results/sklearn-carp"
# python run_protran_sklearn.py --dataset_path="data/proeng/aav/two_vs_many.csv" --encoder_name="carp_76M" --checkpoint=0.125 --embed_batch_size=64 --flatten_emb="mean" --embed_folder="embeddings" --all_embed_layers=True --sklearn_params='{"normalize":true}' --all_result_folder="results/sklearn-carp"

# python run_protran_sklearn.py --dataset_path="data/proeng/aav/two_vs_many.csv" --encoder_name="carp_640M" --checkpoint=1 --embed_batch_size=64 --flatten_emb="mean" --embed_folder="embeddings" --all_embed_layers=True --sklearn_params='{"normalize":true}' --all_result_folder="results/sklearn-carp" --reset_param=True
# python run_protran_sklearn.py --dataset_path="data/proeng/aav/two_vs_many.csv" --encoder_name="carp_640M" --checkpoint=1 --embed_batch_size=64 --flatten_emb="mean" --embed_folder="embeddings" --all_embed_layers=True --sklearn_params='{"normalize":true}' --all_result_folder="results/sklearn-carp" --resample_param=True
# python run_protran_sklearn.py --dataset_path="data/proeng/aav/two_vs_many.csv" --encoder_name="carp_640M" --checkpoint=1 --embed_batch_size=64 --flatten_emb="mean" --embed_folder="embeddings" --all_embed_layers=True --sklearn_params='{"normalize":true}' --all_result_folder="results/sklearn-carp"
# python run_protran_sklearn.py --dataset_path="data/proeng/aav/two_vs_many.csv" --encoder_name="carp_640M" --checkpoint=0.5 --embed_batch_size=64 --flatten_emb="mean" --embed_folder="embeddings" --all_embed_layers=True --sklearn_params='{"normalize":true}' --all_result_folder="results/sklearn-carp"
# python run_protran_sklearn.py --dataset_path="data/proeng/aav/two_vs_many.csv" --encoder_name="carp_640M" --checkpoint=0.25 --embed_batch_size=64 --flatten_emb="mean" --embed_folder="embeddings" --all_embed_layers=True --sklearn_params='{"normalize":true}' --all_result_folder="results/sklearn-carp"
# python run_protran_sklearn.py --dataset_path="data/proeng/aav/two_vs_many.csv" --encoder_name="carp_640M" --checkpoint=0.125 --embed_batch_size=64 --flatten_emb="mean" --embed_folder="embeddings" --all_embed_layers=True --sklearn_params='{"normalize":true}' --all_result_folder="results/sklearn-carp"


# python run_protran_sklearn.py --dataset_path="data/proeng/aav/one_vs_many.csv" --encoder_name="carp_600k" --checkpoint=1 --embed_batch_size=64 --flatten_emb="mean" --embed_folder="embeddings" --all_embed_layers=True --sklearn_params='{"normalize":true}' --all_result_folder="results/sklearn-carp" --reset_param=True
# python run_protran_sklearn.py --dataset_path="data/proeng/aav/one_vs_many.csv" --encoder_name="carp_600k" --checkpoint=1 --embed_batch_size=64 --flatten_emb="mean" --embed_folder="embeddings" --all_embed_layers=True --sklearn_params='{"normalize":true}' --all_result_folder="results/sklearn-carp" --resample_param=True
# python run_protran_sklearn.py --dataset_path="data/proeng/aav/one_vs_many.csv" --encoder_name="carp_600k" --checkpoint=1 --embed_batch_size=64 --flatten_emb="mean" --embed_folder="embeddings" --all_embed_layers=True --sklearn_params='{"normalize":true}' --all_result_folder="results/sklearn-carp"
# python run_protran_sklearn.py --dataset_path="data/proeng/aav/one_vs_many.csv" --encoder_name="carp_600k" --checkpoint=0.5 --embed_batch_size=64 --flatten_emb="mean" --embed_folder="embeddings" --all_embed_layers=True --sklearn_params='{"normalize":true}' --all_result_folder="results/sklearn-carp"
# python run_protran_sklearn.py --dataset_path="data/proeng/aav/one_vs_many.csv" --encoder_name="carp_600k" --checkpoint=0.25 --embed_batch_size=64 --flatten_emb="mean" --embed_folder="embeddings" --all_embed_layers=True --sklearn_params='{"normalize":true}' --all_result_folder="results/sklearn-carp"
# python run_protran_sklearn.py --dataset_path="data/proeng/aav/one_vs_many.csv" --encoder_name="carp_600k" --checkpoint=0.125 --embed_batch_size=64 --flatten_emb="mean" --embed_folder="embeddings" --all_embed_layers=True --sklearn_params='{"normalize":true}' --all_result_folder="results/sklearn-carp"

# python run_protran_sklearn.py --dataset_path="data/proeng/aav/one_vs_many.csv" --encoder_name="carp_38M" --checkpoint=1 --embed_batch_size=64 --flatten_emb="mean" --embed_folder="embeddings" --all_embed_layers=True --sklearn_params='{"normalize":true}' --all_result_folder="results/sklearn-carp" --reset_param=True
# python run_protran_sklearn.py --dataset_path="data/proeng/aav/one_vs_many.csv" --encoder_name="carp_38M" --checkpoint=1 --embed_batch_size=64 --flatten_emb="mean" --embed_folder="embeddings" --all_embed_layers=True --sklearn_params='{"normalize":true}' --all_result_folder="results/sklearn-carp" --resample_param=True
# python run_protran_sklearn.py --dataset_path="data/proeng/aav/one_vs_many.csv" --encoder_name="carp_38M" --checkpoint=1 --embed_batch_size=64 --flatten_emb="mean" --embed_folder="embeddings" --all_embed_layers=True --sklearn_params='{"normalize":true}' --all_result_folder="results/sklearn-carp"
# python run_protran_sklearn.py --dataset_path="data/proeng/aav/one_vs_many.csv" --encoder_name="carp_38M" --checkpoint=0.5 --embed_batch_size=64 --flatten_emb="mean" --embed_folder="embeddings" --all_embed_layers=True --sklearn_params='{"normalize":true}' --all_result_folder="results/sklearn-carp"
# python run_protran_sklearn.py --dataset_path="data/proeng/aav/one_vs_many.csv" --encoder_name="carp_38M" --checkpoint=0.25 --embed_batch_size=64 --flatten_emb="mean" --embed_folder="embeddings" --all_embed_layers=True --sklearn_params='{"normalize":true}' --all_result_folder="results/sklearn-carp"
# python run_protran_sklearn.py --dataset_path="data/proeng/aav/one_vs_many.csv" --encoder_name="carp_38M" --checkpoint=0.125 --embed_batch_size=64 --flatten_emb="mean" --embed_folder="embeddings" --all_embed_layers=True --sklearn_params='{"normalize":true}' --all_result_folder="results/sklearn-carp"

# python run_protran_sklearn.py --dataset_path="data/proeng/aav/one_vs_many.csv" --encoder_name="carp_76M" --checkpoint=1 --embed_batch_size=64 --flatten_emb="mean" --embed_folder="embeddings" --all_embed_layers=True --sklearn_params='{"normalize":true}' --all_result_folder="results/sklearn-carp" --reset_param=True
# python run_protran_sklearn.py --dataset_path="data/proeng/aav/one_vs_many.csv" --encoder_name="carp_76M" --checkpoint=1 --embed_batch_size=64 --flatten_emb="mean" --embed_folder="embeddings" --all_embed_layers=True --sklearn_params='{"normalize":true}' --all_result_folder="results/sklearn-carp" --resample_param=True
# python run_protran_sklearn.py --dataset_path="data/proeng/aav/one_vs_many.csv" --encoder_name="carp_76M" --checkpoint=1 --embed_batch_size=64 --flatten_emb="mean" --embed_folder="embeddings" --all_embed_layers=True --sklearn_params='{"normalize":true}' --all_result_folder="results/sklearn-carp"
# python run_protran_sklearn.py --dataset_path="data/proeng/aav/one_vs_many.csv" --encoder_name="carp_76M" --checkpoint=0.5 --embed_batch_size=64 --flatten_emb="mean" --embed_folder="embeddings" --all_embed_layers=True --sklearn_params='{"normalize":true}' --all_result_folder="results/sklearn-carp"
# python run_protran_sklearn.py --dataset_path="data/proeng/aav/one_vs_many.csv" --encoder_name="carp_76M" --checkpoint=0.25 --embed_batch_size=64 --flatten_emb="mean" --embed_folder="embeddings" --all_embed_layers=True --sklearn_params='{"normalize":true}' --all_result_folder="results/sklearn-carp"
# python run_protran_sklearn.py --dataset_path="data/proeng/aav/one_vs_many.csv" --encoder_name="carp_76M" --checkpoint=0.125 --embed_batch_size=64 --flatten_emb="mean" --embed_folder="embeddings" --all_embed_layers=True --sklearn_params='{"normalize":true}' --all_result_folder="results/sklearn-carp"

# python run_protran_sklearn.py --dataset_path="data/proeng/aav/one_vs_many.csv" --encoder_name="carp_640M" --checkpoint=1 --embed_batch_size=64 --flatten_emb="mean" --embed_folder="embeddings" --all_embed_layers=True --sklearn_params='{"normalize":true}' --all_result_folder="results/sklearn-carp" --reset_param=True
# python run_protran_sklearn.py --dataset_path="data/proeng/aav/one_vs_many.csv" --encoder_name="carp_640M" --checkpoint=1 --embed_batch_size=64 --flatten_emb="mean" --embed_folder="embeddings" --all_embed_layers=True --sklearn_params='{"normalize":true}' --all_result_folder="results/sklearn-carp" --resample_param=True
# python run_protran_sklearn.py --dataset_path="data/proeng/aav/one_vs_many.csv" --encoder_name="carp_640M" --checkpoint=1 --embed_batch_size=64 --flatten_emb="mean" --embed_folder="embeddings" --all_embed_layers=True --sklearn_params='{"normalize":true}' --all_result_folder="results/sklearn-carp"
# python run_protran_sklearn.py --dataset_path="data/proeng/aav/one_vs_many.csv" --encoder_name="carp_640M" --checkpoint=0.5 --embed_batch_size=64 --flatten_emb="mean" --embed_folder="embeddings" --all_embed_layers=True --sklearn_params='{"normalize":true}' --all_result_folder="results/sklearn-carp"
# python run_protran_sklearn.py --dataset_path="data/proeng/aav/one_vs_many.csv" --encoder_name="carp_640M" --checkpoint=0.25 --embed_batch_size=64 --flatten_emb="mean" --embed_folder="embeddings" --all_embed_layers=True --sklearn_params='{"normalize":true}' --all_result_folder="results/sklearn-carp"
# python run_protran_sklearn.py --dataset_path="data/proeng/aav/one_vs_many.csv" --encoder_name="carp_640M" --checkpoint=0.125 --embed_batch_size=64 --flatten_emb="mean" --embed_folder="embeddings" --all_embed_layers=True --sklearn_params='{"normalize":true}' --all_result_folder="results/sklearn-carp"


### thermo ### 

# python run_protran_sklearn.py --dataset_path="data/proeng/thermo/mixed_split.csv" --encoder_name="carp_600k" --checkpoint=1 --embed_batch_size=64 --flatten_emb="mean" --embed_folder="embeddings" --all_embed_layers=True --sklearn_params='{"normalize":true}' --all_result_folder="results/sklearn-carp" --reset_param=True
# python run_protran_sklearn.py --dataset_path="data/proeng/thermo/mixed_split.csv" --encoder_name="carp_600k" --checkpoint=1 --embed_batch_size=64 --flatten_emb="mean" --embed_folder="embeddings" --all_embed_layers=True --sklearn_params='{"normalize":true}' --all_result_folder="results/sklearn-carp" --resample_param=True
# python run_protran_sklearn.py --dataset_path="data/proeng/thermo/mixed_split.csv" --encoder_name="carp_600k" --checkpoint=1 --embed_batch_size=64 --flatten_emb="mean" --embed_folder="embeddings" --all_embed_layers=True --sklearn_params='{"normalize":true}' --all_result_folder="results/sklearn-carp"
# python run_protran_sklearn.py --dataset_path="data/proeng/thermo/mixed_split.csv" --encoder_name="carp_600k" --checkpoint=0.5 --embed_batch_size=64 --flatten_emb="mean" --embed_folder="embeddings" --all_embed_layers=True --sklearn_params='{"normalize":true}' --all_result_folder="results/sklearn-carp"
# python run_protran_sklearn.py --dataset_path="data/proeng/thermo/mixed_split.csv" --encoder_name="carp_600k" --checkpoint=0.25 --embed_batch_size=64 --flatten_emb="mean" --embed_folder="embeddings" --all_embed_layers=True --sklearn_params='{"normalize":true}' --all_result_folder="results/sklearn-carp"
# python run_protran_sklearn.py --dataset_path="data/proeng/thermo/mixed_split.csv" --encoder_name="carp_600k" --checkpoint=0.125 --embed_batch_size=64 --flatten_emb="mean" --embed_folder="embeddings" --all_embed_layers=True --sklearn_params='{"normalize":true}' --all_result_folder="results/sklearn-carp"

# python run_protran_sklearn.py --dataset_path="data/proeng/thermo/mixed_split.csv" --encoder_name="carp_38M" --checkpoint=1 --embed_batch_size=64 --flatten_emb="mean" --embed_folder="embeddings" --all_embed_layers=True --sklearn_params='{"normalize":true}' --all_result_folder="results/sklearn-carp" --reset_param=True
# python run_protran_sklearn.py --dataset_path="data/proeng/thermo/mixed_split.csv" --encoder_name="carp_38M" --checkpoint=1 --embed_batch_size=64 --flatten_emb="mean" --embed_folder="embeddings" --all_embed_layers=True --sklearn_params='{"normalize":true}' --all_result_folder="results/sklearn-carp" --resample_param=True
# python run_protran_sklearn.py --dataset_path="data/proeng/thermo/mixed_split.csv" --encoder_name="carp_38M" --checkpoint=1 --embed_batch_size=64 --flatten_emb="mean" --embed_folder="embeddings" --all_embed_layers=True --sklearn_params='{"normalize":true}' --all_result_folder="results/sklearn-carp"
# python run_protran_sklearn.py --dataset_path="data/proeng/thermo/mixed_split.csv" --encoder_name="carp_38M" --checkpoint=0.5 --embed_batch_size=64 --flatten_emb="mean" --embed_folder="embeddings" --all_embed_layers=True --sklearn_params='{"normalize":true}' --all_result_folder="results/sklearn-carp"
# python run_protran_sklearn.py --dataset_path="data/proeng/thermo/mixed_split.csv" --encoder_name="carp_38M" --checkpoint=0.25 --embed_batch_size=64 --flatten_emb="mean" --embed_folder="embeddings" --all_embed_layers=True --sklearn_params='{"normalize":true}' --all_result_folder="results/sklearn-carp"
# python run_protran_sklearn.py --dataset_path="data/proeng/thermo/mixed_split.csv" --encoder_name="carp_38M" --checkpoint=0.125 --embed_batch_size=64 --flatten_emb="mean" --embed_folder="embeddings" --all_embed_layers=True --sklearn_params='{"normalize":true}' --all_result_folder="results/sklearn-carp"

# python run_protran_sklearn.py --dataset_path="data/proeng/thermo/mixed_split.csv" --encoder_name="carp_76M" --checkpoint=1 --embed_batch_size=64 --flatten_emb="mean" --embed_folder="embeddings" --all_embed_layers=True --sklearn_params='{"normalize":true}' --all_result_folder="results/sklearn-carp" --reset_param=True
# python run_protran_sklearn.py --dataset_path="data/proeng/thermo/mixed_split.csv" --encoder_name="carp_76M" --checkpoint=1 --embed_batch_size=64 --flatten_emb="mean" --embed_folder="embeddings" --all_embed_layers=True --sklearn_params='{"normalize":true}' --all_result_folder="results/sklearn-carp" --resample_param=True
# python run_protran_sklearn.py --dataset_path="data/proeng/thermo/mixed_split.csv" --encoder_name="carp_76M" --checkpoint=1 --embed_batch_size=64 --flatten_emb="mean" --embed_folder="embeddings" --all_embed_layers=True --sklearn_params='{"normalize":true}' --all_result_folder="results/sklearn-carp"
# python run_protran_sklearn.py --dataset_path="data/proeng/thermo/mixed_split.csv" --encoder_name="carp_76M" --checkpoint=0.5 --embed_batch_size=64 --flatten_emb="mean" --embed_folder="embeddings" --all_embed_layers=True --sklearn_params='{"normalize":true}' --all_result_folder="results/sklearn-carp"
# python run_protran_sklearn.py --dataset_path="data/proeng/thermo/mixed_split.csv" --encoder_name="carp_76M" --checkpoint=0.25 --embed_batch_size=64 --flatten_emb="mean" --embed_folder="embeddings" --all_embed_layers=True --sklearn_params='{"normalize":true}' --all_result_folder="results/sklearn-carp"
# python run_protran_sklearn.py --dataset_path="data/proeng/thermo/mixed_split.csv" --encoder_name="carp_76M" --checkpoint=0.125 --embed_batch_size=64 --flatten_emb="mean" --embed_folder="embeddings" --all_embed_layers=True --sklearn_params='{"normalize":true}' --all_result_folder="results/sklearn-carp"

# python run_protran_sklearn.py --dataset_path="data/proeng/thermo/mixed_split.csv" --encoder_name="carp_640M" --checkpoint=1 --embed_batch_size=64 --flatten_emb="mean" --embed_folder="embeddings" --all_embed_layers=True --sklearn_params='{"normalize":true}' --all_result_folder="results/sklearn-carp" --reset_param=True
# python run_protran_sklearn.py --dataset_path="data/proeng/thermo/mixed_split.csv" --encoder_name="carp_640M" --checkpoint=1 --embed_batch_size=64 --flatten_emb="mean" --embed_folder="embeddings" --all_embed_layers=True --sklearn_params='{"normalize":true}' --all_result_folder="results/sklearn-carp" --resample_param=True
# python run_protran_sklearn.py --dataset_path="data/proeng/thermo/mixed_split.csv" --encoder_name="carp_640M" --checkpoint=1 --embed_batch_size=64 --flatten_emb="mean" --embed_folder="embeddings" --all_embed_layers=True --sklearn_params='{"normalize":true}' --all_result_folder="results/sklearn-carp"
# python run_protran_sklearn.py --dataset_path="data/proeng/thermo/mixed_split.csv" --encoder_name="carp_640M" --checkpoint=0.5 --embed_batch_size=64 --flatten_emb="mean" --embed_folder="embeddings" --all_embed_layers=True --sklearn_params='{"normalize":true}' --all_result_folder="results/sklearn-carp"
# python run_protran_sklearn.py --dataset_path="data/proeng/thermo/mixed_split.csv" --encoder_name="carp_640M" --checkpoint=0.25 --embed_batch_size=64 --flatten_emb="mean" --embed_folder="embeddings" --all_embed_layers=True --sklearn_params='{"normalize":true}' --all_result_folder="results/sklearn-carp"
# python run_protran_sklearn.py --dataset_path="data/proeng/thermo/mixed_split.csv" --encoder_name="carp_640M" --checkpoint=0.125 --embed_batch_size=64 --flatten_emb="mean" --embed_folder="embeddings" --all_embed_layers=True --sklearn_params='{"normalize":true}' --all_result_folder="results/sklearn-carp"
```
