## Supplemental data and code for "Feature Reuse and Scaling: Understanding Transfer Learning with Protein Language Models": run_dataset_analysis.py

```
"""A script for running dataset analysis"""

from __future__ import annotations

from scr.preprocess.data_process import TaskProcess
from scr.vis.dataset_vis import DatasetSeqLenHist, DatasetStripHistogram

result_subfolder = "results/dataset_vis/len_hist"

summary_df = TaskProcess(forceregen=True, showplot=False).sum_file_df

for _, row in summary_df.iterrows():
    print(f"Plotting sequence length histogram for {row.csv_path}...")
    DatasetSeqLenHist(df_path=row.csv_path, result_subfolder=result_subfolder)
    

# DatasetStripHistogram(
#     dataset_folder="data/proeng/gb1",
#     split_order=["sampled", "two_vs_rest", "low_vs_high"],
#     plot_width=800,
#     showplot=False,
# )
# DatasetStripHistogram(
#     dataset_folder="data/proeng/aav",
#     split_order=["one_vs_many", "two_vs_many"],
#     showplot=False,
# )
# DatasetStripHistogram(
#     dataset_folder="data/proeng/thermo", 
#     plot_width=400,
#     showplot=False)
```
