## Supplemental data and code for "Feature Reuse and Scaling: Understanding Transfer Learning with Protein Language Models": run_pregen_emb.py

```
"""Script for pre generating all embeddings"""
from __future__ import annotations

import json
import argparse

from scr.encoding.gen_encoding import GenerateEmbeddings
from scr.params.emb import TRANSFORMER_INFO, CARP_INFO
from scr.utils import get_default_output_path

parser = argparse.ArgumentParser(description="Embedding Generation")


parser.add_argument(
    "--dataset_path",
    type=str,
    metavar="P",
    help="full path to the dataset, in pkl or panda readable format, \
    ie: data/proeng/gb1/two_vs_rest.pkl or data/annotation/scl/balanced.csv",
)

parser.add_argument(
    "--encoder_name",
    type=str,
    metavar="EN",
    help="the name of the encoder, ie: esm1b_t33_650M_UR50S",
)

parser.add_argument(
    "--checkpoint",
    type=float,
    metavar="CP",
    default=1,
    help="the fraction of the pretrain model, ie: 0.5",
)

parser.add_argument(
    "--checkpoint_folder",
    type=str,
    metavar="CPF",
    default="pretrain_checkpoints/carp",
    help="the folder for the pretrain model, ie: pretrain_checkpoints/carp",
)

parser.add_argument(
    "--reset_param",
    type=bool,
    metavar="RIP",
    default=False,
    help="if update the full model to xavier_uniform_ (default: False)",
)

parser.add_argument(
    "--resample_param",
    type=bool,
    metavar="STP",
    default=False,
    help="if update the full model to xavier_normal_ (default: False)",
)

parser.add_argument(
    "--embed_batch_size",
    type=int,
    metavar="EBS",
    default=8,
    help="the embedding batch size, set to 0 to encode all in a single batch (default: 128)",
)

parser.add_argument(
    "--flatten_emb",
    metavar="FE",
    default=False,
    help="if (False) and how ('mean', 'max']) to flatten the embedding (default: False)",
)

parser.add_argument(
    "--seq_start_idx",
    metavar="SSI",
    default=False,
    help="the index for the start of the sequence (default: False)",
)

parser.add_argument(
    "--seq_end_idx",
    metavar="SEI",
    default=False,
    help="the index for the end of the sequence (default: False)",
)

parser.add_argument(
    "--subset_list",
    metavar="SL",
    type=json.loads,
    default=['train', 'val', 'test'],
    help="the index for the end of the sequence (default: False)",
)

parser.add_argument(
    "--embed_folder",
    type=str,
    default="embeddings",
    metavar="O",
    help="the parent folder for embeddings (default: 'embeddings')",
)

# TODO add encoder_params

args = parser.parse_args()

GenerateEmbeddings(
    dataset_path=args.dataset_path,
    encoder_name=args.encoder_name,
    checkpoint=args.checkpoint,
    checkpoint_folder=args.checkpoint_folder,
    reset_param=args.reset_param,
    resample_param=args.resample_param,
    embed_batch_size=args.embed_batch_size,
    flatten_emb=args.flatten_emb,
    seq_start_idx=args.seq_start_idx,
    seq_end_idx=args.seq_end_idx,
    subset_list=args.subset_list,
    embed_folder=get_default_output_path(args.embed_folder),
    # **encoder_params,
)
```
