## Supplemental data and code for "Feature Reuse and Scaling: Understanding Transfer Learning with Protein Language Models": run_protran_pytorch.py

```
import argparse

from scr.params.sys import DEVICE, RAND_SEED
from scr.model.run_pytorch import Run_Pytorch

parser = argparse.ArgumentParser(description="Protein transfer with pytorch models")

parser.add_argument(
    "--dataset_path",
    type=str,
    metavar="P",
    help="full path to the dataset, in pkl or panda readable format, \
    ie: data/proeng/gb1/two_vs_rest.pkl or data/annotation/scl/balanced.csv",
)

parser.add_argument(
    "--encoder_name",
    type=str,
    metavar="EN",
    help="the name of the encoder, ie: esm1b_t33_650M_UR50S",
)

parser.add_argument(
    "--checkpoint",
    type=float,
    metavar="CP",
    default=1,
    help="the fraction of the pretrain model, ie: 0.5",
)

parser.add_argument(
    "--checkpoint_folder",
    type=str,
    metavar="CPF",
    default="pretrain_checkpoints/carp",
    help="the folder for the pretrain model, ie: pretrain_checkpoints/carp",
)


parser.add_argument(
    "--reset_param",
    type=bool,
    metavar="RIP",
    default=False,
    help="if update the full model to xavier_uniform_ (default: False)",
)

parser.add_argument(
    "--resample_param",
    type=bool,
    metavar="STP",
    default=False,
    help="if update the full model to xavier_normal_ (default: False)",
)

parser.add_argument(
    "--embed_batch_size",
    type=int,
    metavar="EBS",
    default=128,
    help="the embedding batch size, set to 0 to encode all in a single batch (default: 128)",
)

parser.add_argument(
    "--flatten_emb",
    metavar="FE",
    default=False,
    help="if (False) and how ('mean', 'max']) to flatten the embedding (default: 'mean')",
)

parser.add_argument(
    "--embed_folder",
    metavar="EP",
    default=None,
    help="path to presaved embedding (default: None)",
)

parser.add_argument(
    "--seq_start_idx",
    metavar="SSI",
    default=False,
    help="the index for the start of the sequence (default: False)",
)

parser.add_argument(
    "--seq_end_idx",
    metavar="SEI",
    default=False,
    help="the index for the end of the sequence (default: False)",
)

parser.add_argument(
    "--manual_layer_min",
    metavar="LMIN",
    default=False,
    help="the number of layer for manual start range (default: False)",
)

parser.add_argument(
    "--manual_layer_max",
    metavar="LMAX",
    default=False,
    help="the number of layer for manual end range (default: False)",
)
parser.add_argument(
    "--loader_batch_size",
    type=int,
    metavar="LBS",
    default=64,
    help="the batch size for train, val, and test dataloader (default: False)",
)

parser.add_argument(
    "--worker_seed",
    type=int,
    metavar="WS",
    default=RAND_SEED,
    help="the seed for dataloader (default: RAND_SEED)",
)

parser.add_argument(
    "--if_encode_all",
    type=bool,
    metavar="EA",
    default=False,
    help="if encode full dataset all layers on the fly (default: False)",
)

parser.add_argument(
    "--if_multiprocess",
    type=bool,
    metavar="MP",
    default=False,
    help="if running all layers in parallel (default: False)",
)

parser.add_argument(
    "--learning_rate",
    type=float,
    metavar="LR",
    default=1e-4,
    help="learning rate (default: 1e-4)",
)

parser.add_argument(
    "--lr_decay",
    type=float,
    metavar="LRD",
    default=0.1,
    help="factor by which to decay learning rate on plateau (default: 0.1)",
)

parser.add_argument(
    "--epochs",
    type=int,
    default=20,
    metavar="N",
    help="number of epochs to train (default: 20)",
)

parser.add_argument(
    "--early_stop",
    type=bool,
    default=True,
    metavar="ES",
    help="if initate early stopping (default: True)",
)

parser.add_argument(
    "--tolerance",
    type=int,
    default=10,
    metavar="T",
    help="tolerance for early stopping (default: 10)",
)

parser.add_argument(
    "--min_epoch",
    type=int,
    default=5,
    metavar="ME",
    help="minimal number of epochs for early stopping (default: 5)",
)

parser.add_argument(
    "--device",
    default=DEVICE,
    metavar="D",
    help="torch device (default: DEVICE)",
)

parser.add_argument(
    "--all_plot_folder",
    type=str,
    default="results/learning_curves",
    metavar="LC",
    help="the parent folder for all learning curves (default: 'results/learning_curves')",
)  

parser.add_argument(
    "--all_result_folder",
    type=str,
    default="results/pytorch",
    metavar="O",
    help="the parent folder for all results (default: 'results/pytorch')",
)

# TODO add encoder_params

args = parser.parse_args()
    
Run_Pytorch(
    dataset_path=args.dataset_path,
    encoder_name=args.encoder_name,    
    checkpoint=args.checkpoint,
    checkpoint_folder=args.checkpoint_folder,
    reset_param=args.reset_param,
    resample_param=args.resample_param,
    embed_batch_size=args.embed_batch_size,
    flatten_emb=args.flatten_emb,
    embed_folder=args.embed_folder,
    seq_start_idx=args.seq_start_idx,
    seq_end_idx=args.seq_end_idx,
    manual_layer_min=args.manual_layer_min,
    manual_layer_max=args.manual_layer_max,
    loader_batch_size=args.loader_batch_size,
    worker_seed=args.worker_seed,
    if_encode_all=args.if_encode_all,
    if_multiprocess=args.if_multiprocess,
    learning_rate=args.learning_rate,
    lr_decay=args.lr_decay,
    epochs=args.epochs,
    early_stop=args.early_stop,
    tolerance=args.tolerance,
    min_epoch=args.min_epoch,
    device=args.device,
    all_plot_folder=args.all_plot_folder,
    all_result_folder=args.all_result_folder,
    # **encoder_params,
    )
```
