## Supplemental data and code for "Feature Reuse and Scaling: Understanding Transfer Learning with Protein Language Models": run_protran_sklearn.py

```
from __future__ import annotations

import json
import argparse

import numpy as np

from scr.encoding.encoding_classes import AbstractEncoder, ESMEncoder, CARPEncoder
from scr.model.run_sklearn import RunSK
from scr.params.sys import RAND_SEED, SKLEARN_ALPHAS
from scr.params.emb import TRANSFORMER_INFO, CARP_INFO
from scr.utils import get_default_output_path

def alpha_types(alphas: np.ndarray | float):
    """
    Set the type for the ridge regression alphas

    Args:
    - alphas: np.ndarray | float
    """
    if not isinstance(alphas, np.ndarray):
        return np.array([alphas])

parser = argparse.ArgumentParser(description="Protein transfer")

parser.add_argument(
    "--dataset_path",
    type=str,
    metavar="P",
    help="full path to the dataset, in pkl or panda readable format, \
    ie: data/proeng/gb1/two_vs_rest.pkl or data/annotation/scl/balanced.csv",
)

parser.add_argument(
    "--encoder_name",
    type=str,
    metavar="EN",
    help="the name of the encoder, ie: esm1b_t33_650M_UR50S",
)

parser.add_argument(
    "--checkpoint",
    type=float,
    metavar="CP",
    default=1,
    help="the fraction of the pretrain model, ie: 0.5",
)

parser.add_argument(
    "--checkpoint_folder",
    type=str,
    metavar="CPF",
    default="pretrain_checkpoints/carp",
    help="the folder for the pretrain model, ie: pretrain_checkpoints/carp",
)

parser.add_argument(
    "--reset_param",
    type=bool,
    metavar="RIP",
    default=False,
    help="if update the full model to xavier_uniform_ (default: False)",
)

parser.add_argument(
    "--resample_param",
    type=bool,
    metavar="STP",
    default=False,
    help="if update the full model to xavier_normal_ (default: False)",
)

parser.add_argument(
    "--embed_batch_size",
    type=int,
    metavar="EBS",
    default=128,
    help="the embedding batch size, set to 0 to encode all in a single batch (default: 128)",
)

parser.add_argument(
    "--flatten_emb",
    metavar="FE",
    default="mean",
    help="if (False) and how ('mean', 'max']) to flatten the embedding (default: 'mean')",
)

parser.add_argument(
    "--embed_folder",
    metavar="EP",
    default=None,
    help="path to presaved embedding (default: None)",
)

parser.add_argument(
    "--all_embed_layers",
    type=bool,
    metavar="AEL",
    default=False,
    help="if include all embed layers (default: False)",
)

parser.add_argument(
    "--seq_start_idx",
    metavar="SSI",
    default=False,
    help="the index for the start of the sequence (default: False)",
)

parser.add_argument(
    "--seq_end_idx",
    metavar="SEI",
    default=False,
    help="the index for the end of the sequence (default: False)",
)

parser.add_argument(
    "--if_encode_all",
    type=bool,
    metavar="EA",
    default=False,
    help="if encode all embed layers all at once (default: False)",
)

parser.add_argument(
    "--alphas",
    type=alpha_types,
    metavar="A",
    default=SKLEARN_ALPHAS,
    help="arrays of alphas to be tested (default: SKLEARN_ALPHAS)",
)

parser.add_argument(
    "--sklearn_state",
    type=int,
    metavar="RS",
    default=RAND_SEED,
    help="the seed for ridge regression (default: RAND_SEED)",
)

parser.add_argument(
    "--sklearn_params",
    type=json.loads,
    metavar="RP",
    default=None,
    help="additional argument for ridge regression (default: None)",
)

parser.add_argument(
    "--all_result_folder",
    type=str,
    default="results/sklearn",
    metavar="O",
    help="the parent folder for all results (default: 'results/sklearn')",
)

# TODO add encoder_params

args = parser.parse_args()

RunSK(
    dataset_path=args.dataset_path,
    encoder_name=args.encoder_name,
    checkpoint=args.checkpoint,
    checkpoint_folder=args.checkpoint_folder,
    reset_param=args.reset_param,
    resample_param=args.resample_param,
    embed_batch_size=args.embed_batch_size,
    flatten_emb=args.flatten_emb,
    embed_folder=args.embed_folder,
    all_embed_layers=args.all_embed_layers,
    seq_start_idx=args.seq_start_idx,
    seq_end_idx=args.seq_end_idx,
    if_encode_all=args.if_encode_all,
    alphas=args.alphas,
    sklearn_state=args.sklearn_state,
    sklearn_params=args.sklearn_params,
    all_result_folder=get_default_output_path(args.all_result_folder),
     #**encoder_params,
)
```
