## Supplemental data and code for "Feature Reuse and Scaling: Understanding Transfer Learning with Protein Language Models": run_results_analysis.py

```
"""Script for running results analysis and plotting"""

import matplotlib as mpl


from scr.analysis.perlayer import LayerLoss
from scr.analysis.result_reorg import ResultReorg
from scr.vis.res_vis import PlotResultScatter
from scr.vis.res_vis_task import PlotResultByTask
from scr.params.emb import ARCH_TYPE, ARCH_CUT_DICT

notitle = False

if notitle == True:

    # Set title color to white and font size to zero
    mpl.rcParams['axes.titlecolor'] = 'white'
    mpl.rcParams['axes.titleweight'] = 0
    mpl.rcParams['axes.titlepad'] = 0
    mpl.rcParams['axes.titlesize'] = 0


"""
print("Running results analysis and plotting for sklearn CARP...")

LayerLoss(
    input_path="results/sklearn-carp",
    output_path="results/sklearn-carp_layer"
)

print("Running results analysis and plotting for sklearn ESM...")

LayerLoss(
    input_path="results/sklearn-esm",
    output_path="results/sklearn-esm_layer",
    add_checkpoint=False
)

print("Running results analysis and plotting for pytorch CARP...")

LayerLoss(
    input_path="results/pytorch-carp",
    output_path="results/pytorch-carp_layer"
)

print("Running results analysis and plotting for pytorch ESM...")

LayerLoss(
    input_path="results/pytorch-esm",
    output_path="results/pytorch-esm_layer",
    add_checkpoint=False
)
"""

# ResultReorg()
"""
plot_class = PlotResultScatter()

for metric in ["test_loss", "test_performance_1", "test_performance_2"]:
    
    plot_class.plot_emb_onehot(
            metric = metric,
        );

    for arch in ARCH_TYPE:

        for cut in ARCH_CUT_DICT[arch]:
            for s in [True, False]:
                plot_class.plot_layer_delta(
                    layer_cut=cut,
                    metric=metric,
                    arch=arch,
                    ifsimple=s,
                );
    
    # get pretrain degree for carp only
    plot_class.plot_pretrain_degree(
        metric=metric,
        arch="carp")
    
    # plot for arch size
    plot_class.plot_arch_size(
        metric=metric
    )
"""

task_class = PlotResultByTask()

for metric in ["test_loss", "test_performance_1", "test_performance_2"]:
    
    task_class._plot_by_cluster(
            metric = metric,
        );
```
