## Supplemental data and code for "Feature Reuse and Scaling: Understanding Transfer Learning with Protein Language Models": perlayer.py

```
"""Analyzing per layer output"""

from __future__ import annotations

from collections import defaultdict

import os
from glob import glob
import numpy as np

import matplotlib.pyplot as plt
from matplotlib.ticker import FormatStrFormatter

from scr.encoding.encoding_classes import get_emb_info
from scr.params.emb import TRANSFORMER_INFO, CARP_INFO
from scr.params.vis import CHECKPOINT_COLOR
from scr.analysis.utils import METRIC_DICT
from scr.utils import pickle_load, get_filename, checkNgen_folder


class LayerLoss:
    """A class for handling layer analysis"""

    def __init__(
        self,
        add_checkpoint: bool = True,
        checkpoint_list: list = [0.5, 0.25, 0.125],
        input_path: str = "results/sklearn",
        output_path: str = "results/sklearn_layer",
        metric_dict: dict[list[str]] = METRIC_DICT,
    ):
        """
        Args:
        - add_checkpoint: bool = True, if add checkpoint for carp
        - checkpoint_list: list = [0.5, 0.25, 0.125],
        - input_path: str = "results/sklearn",
        - output_path: str = "results/sklearn_layer"
        - metric_dict: list[str] = ["train_mse", "test_ndcg", "test_rho"]
        """
        self._add_checkpoint = add_checkpoint
        self._checkpoint_list = checkpoint_list
        # get rid of the last "/" if any
        self._input_path = os.path.normpath(input_path)
        # get the list of subfolders for each dataset
        self._dataset_folders = glob(f"{self._input_path}/*/*/*/*/*")
        # glob("results/train_val_test/*/*/*/*/*")

        # get rid of the last "/" if any
        self._output_path = os.path.normpath(output_path)
        self._metric_dict = metric_dict

        # init a dictionary for recording outputs
        self._onehot_baseline_dict = defaultdict(dict)
        self._layer_analysis_dict = defaultdict(dict)
        self._rand_layer_analysis_dict = defaultdict(dict)
        self._stat_layer_analysis_dict = defaultdict(dict)

        # init a dict for metric params
        self._metric_numb = defaultdict(dict)

        # init
        self._checkpoint_analysis_dict = defaultdict(dict)
        if self._add_checkpoint:
            for checkpoint in self._checkpoint_list:
                self._checkpoint_analysis_dict[checkpoint] = defaultdict(dict)

        for dataset_folder in self._dataset_folders:
            # dataset_folder = "results/train_val_test/proeng/gb1/two_vs_rest/esm1b_t33_650M_UR50S/max"
            # get the details for the dataset such as proeng/gb1/two_vs_rest
            task_subfolder = dataset_folder.split(self._input_path + "/")[-1]
            # task_subfolder = "proeng/gb1/two_vs_rest/esm1b_t33_650M_UR50S/max"
            task, dataset, split, encoder_name, flatten_emb = task_subfolder.split("/")
            # get collage_name
            collage_name = f"{task}_{dataset}_{split}_{flatten_emb}"

            # get number of metircs
            self._metric_numb[collage_name] = len(self._metric_dict[task])

            # parse results for plotting the collage and onehot
            self._layer_analysis_dict[collage_name][
                encoder_name
            ] = self.parse_result_dicts(
                dataset_folder, task, dataset, split, encoder_name, flatten_emb
            )

            # init
            # check if check points exists
            if self._add_checkpoint:
                for checkpoint in self._checkpoint_list:
                    checkpoint_path = f"{self._input_path}-{str(checkpoint)}"

                    if os.path.exists(checkpoint_path):
                        self._checkpoint_analysis_dict[checkpoint][
                            f"{task}_{dataset}_{split}_{flatten_emb}"
                        ][encoder_name] = self.parse_result_dicts(
                            dataset_folder.replace(self._input_path, checkpoint_path),
                            task,
                            dataset,
                            split,
                            encoder_name,
                            flatten_emb,
                        )

            # check if reset param experimental results exist
            reset_param_path = f"{self._input_path}-rand"

            if os.path.exists(reset_param_path):
                self._rand_layer_analysis_dict[
                    f"{task}_{dataset}_{split}_{flatten_emb}"
                ][encoder_name] = self.parse_result_dicts(
                    dataset_folder.replace(self._input_path, reset_param_path),
                    task,
                    dataset,
                    split,
                    encoder_name,
                    flatten_emb,
                )
                add_rand = True
            else:
                add_rand = False

            # check if resample param experimental results exist
            resample_param_path = f"{self._input_path}-stat"

            if os.path.exists(resample_param_path):
                self._stat_layer_analysis_dict[
                    f"{task}_{dataset}_{split}_{flatten_emb}"
                ][encoder_name] = self.parse_result_dicts(
                    dataset_folder.replace(self._input_path, resample_param_path),
                    task,
                    dataset,
                    split,
                    encoder_name,
                    flatten_emb,
                )
                add_stat = True
            else:
                add_stat = False

            # check if onehot experimental results exist
            onehot_path = f"{self._input_path}-onehot"

            if os.path.exists(onehot_path):
                if task == "structure":
                    onehot_flatten_emb_name = "noflatten"
                else:
                    onehot_flatten_emb_name = "flatten"
                self._onehot_baseline_dict[
                    f"{task}_{dataset}_{split}"
                ]["onehot"] = self.parse_result_dicts(
                    dataset_folder.replace(self._input_path, onehot_path)
                    .replace(encoder_name, "onehot")
                    .replace(flatten_emb, onehot_flatten_emb_name),
                    task,
                    dataset,
                    split,
                    "onehot",
                    onehot_flatten_emb_name,
                )
                add_onehot = True
            else:
                add_onehot = False

        # combine different model into one big plot with different encoders
        collage_folder = os.path.join(self._output_path, "collage")
        checkNgen_folder(collage_folder)

        for collage_name, encoder_dict in self._layer_analysis_dict.items():

            print(f"Plotting collage_name {collage_name}...")

            onehot_name = "_".join(collage_name.split("_")[:-1])

            if set(list(TRANSFORMER_INFO.keys())) == set(encoder_dict.keys()):
                # set the key rankings to default
                encoder_names = list(TRANSFORMER_INFO.keys())
                encoder_label = "esm"
            elif set(list(CARP_INFO.keys())) == set(encoder_dict.keys()):
                # set the key rankings to default
                encoder_names = list(CARP_INFO.keys())
                encoder_label = "carp"
            else:
                encoder_names = list(set(encoder_dict.keys()))
                encoder_label = "pretrained"

            fig, axs = plt.subplots(
                self._metric_numb[collage_name],
                len(encoder_names),
                sharey="row",
                sharex="col",
                figsize=(20, 2 * self._metric_numb[collage_name]),
                squeeze=False # not get rid off the extra dim if 1D
            )

            for m, metric in enumerate(self._metric_dict[collage_name.split("_")[0]]):

                for n, encoder_name in enumerate(encoder_names):
                    axs[m, n].plot(
                        encoder_dict[encoder_name][metric],
                        label=encoder_label,
                        color="#f79646ff",  # orange
                    )

                    # add checkpoints
                    if self._add_checkpoint:
                        for checkpoint in self._checkpoint_list:

                            checkpoint_vals = self._checkpoint_analysis_dict[
                                checkpoint
                            ][collage_name][encoder_name][metric]

                            if not np.all(checkpoint_vals == 0):
                                axs[m, n].plot(
                                    checkpoint_vals,
                                    label=f"{encoder_label}-{checkpoint}",
                                    color=CHECKPOINT_COLOR[
                                        checkpoint
                                    ],  # darker oranges
                                    linestyle="dashed",
                                )

                    # overlay random init
                    if add_rand:
                        axs[m, n].plot(
                            self._rand_layer_analysis_dict[collage_name][encoder_name][
                                metric
                            ],
                            label="random init",
                            color="#4bacc6",  # blue
                            linestyle="dashed"
                            # color="#D3D3D3",  # light grey
                        )

                    # overlay stat init
                    if add_stat:
                        axs[m, n].plot(
                            self._stat_layer_analysis_dict[collage_name][encoder_name][
                                metric
                            ],
                            label="stat transfer",
                            color="#9bbb59",  # green
                            linestyle="dashed"
                            # color="#A9A9A9",  # dark grey
                            # linestyle="dotted",
                        )

                    # overlay onehot baseline
                    if add_onehot:
                        axs[m, n].axhline(
                            self._onehot_baseline_dict[onehot_name]["onehot"][metric],
                            label="onehot",
                            color="#000000",  # black or #D3D3D3 light grey
                            linestyle="dotted",
                        )

            # add xlabels
            for ax in axs[self._metric_numb[collage_name] - 1]:
                ax.set_xlabel("layers", fontsize=16)
                ax.tick_params(axis="x", labelsize=16)

            # add column names
            for ax, col in zip(axs[0], encoder_names):
                ax.set_title(col, fontsize=16)

            # add row names
            for ax, row in zip(
                axs[:, 0], self._metric_dict[collage_name.split("_")[0]]
            ):
                ax.set_ylabel(
                    row.replace("_", " ").replace("cross-entropy", "ce"), fontsize=16
                )
                ax.tick_params(
                    axis="y",
                    which="major",
                    reset=True,
                    labelsize=16,
                    left=True,
                    right=False,  # no right side tick on the plot
                    labelleft=True,
                    labelright=False,
                )
                ax.relim()  # make sure all the data fits
                ax.autoscale()

            # set the plot yticks
            plt.gca().yaxis.set_major_formatter(FormatStrFormatter("%.2f"))
            plt.gca().yaxis.tick_left()
            plt.gca().autoscale()

            # add legend
            handles, labels = axs[0, 0].get_legend_handles_labels()

            if len(labels) == 7:

                # Add two empty dummy legend items
                # using the first label info

                axs[0, 0].axhline(
                    self._onehot_baseline_dict[onehot_name]["onehot"][
                        self._metric_dict[collage_name.split("_")[0]][0]
                    ],
                    label=" ",
                    color="w",
                    alpha=0,
                )

                axs[0, 0].axhline(
                    self._onehot_baseline_dict[onehot_name]["onehot"][
                        self._metric_dict[collage_name.split("_")[0]][0]
                    ],
                    label=" ",
                    color="w",
                    alpha=0,
                )

                adjusted_handles, adjusted_labels = axs[
                    0, 0
                ].get_legend_handles_labels()
                adjusted_y = 1.045
                ncol = 3
                legend_params = {
                    "labelspacing": 0.1,  # vertical space between the legend entries, default 0.5
                    "handletextpad": 0.2,  # space between the legend the text, default 0.8
                    "handlelength": 0.95,  # length of the legend handles, default 2.0
                    "columnspacing": 1,  # spacing between columns, default 2.0
                }

            else:
                adjusted_handles, adjusted_labels = handles, labels
                adjusted_y = 1.025
                ncol = 2
                legend_params = {}

            fig.legend(
                adjusted_handles,
                adjusted_labels,
                loc="upper left",
                bbox_to_anchor=[0.05, adjusted_y],
                fontsize=16,
                frameon=False,
                ncol=ncol,
                **legend_params,
            )

            # add whole plot level title
            fig.suptitle(
                collage_name.replace("_", " ").replace("cross-entropy", "ce"),
                y=1.0025,
                fontsize=24,
                fontweight="bold",
            )
            fig.align_labels()
            fig.tight_layout()

            for plot_ext in [".svg", ".png"]:
                plt.savefig(
                    os.path.join(collage_folder, collage_name + plot_ext),
                    bbox_inches="tight",
                )

            plt.close()

    def parse_result_dicts(
        self,
        folder_path: str,
        task: str,
        dataset: str,
        split: str,
        encoder_name: str,
        flatten_emb: bool | str,
    ):
        """
        Parse the output result dictionaries for plotting

        Args:
        - folder_path: str, the folder path for the datasets

        Returns:
        - dict, encode name as key with a dict as its value
            where metric name as keys and the array of losses as values
        - str, details for collage plot
        """

        # get the list of output pickle files
        pkl_list = glob(f"{folder_path}/*.pkl")

        _, _, max_layer_numb = get_emb_info(encoder_name)

        # init the ouput dict
        output_numb_dict = {
            metric: np.zeros([max_layer_numb]) for metric in self._metric_dict[task]
        }

        # loop through the list of the pickle files
        for pkl_file in pkl_list:
            # get the layer number
            layer_numb = int(get_filename(pkl_file).split("-")[-1].split("_")[-1])
            # load the result dictionary
            try:
                result_dict = pickle_load(pkl_file)
            except Exception as e:
                print(f"{pkl_file} with err: ", e)

            # populate the processed dictionary
            for metric in self._metric_dict[task]:
                subset, kind = metric.split("_")
                if kind == "rho":
                    output_numb_dict[metric][layer_numb] = result_dict[subset][kind][0]
                else:
                    output_numb_dict[metric][layer_numb] = result_dict[subset][kind]

        # get some details for plotting and saving
        output_subfolder = checkNgen_folder(
            folder_path.replace(self._input_path, self._output_path)
        )

        for metric in output_numb_dict.keys():

            plot_name = f"{encoder_name}_{flatten_emb}_{metric}"
            plot_prefix = f"{task}_{dataset}_{split}"

            plt.figure()
            plt.plot(output_numb_dict[metric])
            plt.title(f"{plot_prefix} \n {plot_name}")
            plt.xlabel("layers")
            plt.ylabel("loss")

            for plot_ext in [".svg", ".png"]:
                plt.savefig(
                    os.path.join(output_subfolder, plot_name + plot_ext),
                    bbox_inches="tight",
                )
            plt.close()

        return output_numb_dict

    @property
    def layer_analysis_dict(self) -> dict:
        """Return a dict with dataset name as the key"""
        return self._layer_analysis_dict

    @property
    def rand_layer_analysis_dict(self) -> dict:
        """Return a dict with dataset name as the key for rand"""
        return self._rand_layer_analysis_dict

    @property
    def stat_layer_analysis_dict(self) -> dict:
        """Return a dict with dataset name as the key for stat"""
        return self._stat_layer_analysis_dict

    @property
    def onehot_baseline_dict(self) -> dict:
        """Return a dict with dataset name as the key for onehot"""
        return self._onehot_baseline_dict

    @property
    def checkpoint_analysis_dict(self) -> dict:
        """Return a dict with dataset name as the key for checkpoints"""
        return self._checkpoint_analysis_dict
```
