## Supplemental data and code for "Feature Reuse and Scaling: Understanding Transfer Learning with Protein Language Models": result_reorg.py

```
"""A script reorg results"""

from __future__ import annotations

import os
import pandas as pd

from scr.utils import checkNgen_folder
from scr.analysis.utils import (
    metric_simplifier,
    STRUCT_TESTS,
    DEFAULT_AB_LIST,
    PRETRAIN_ARCH_LIST,
    DS_MODEL_LIST,
)
from scr.analysis.perlayer import LayerLoss


class ResultReorg:
    """A class for reorg layer wise results"""

    def __init__(
        self,
        layer_folder: str = "results",
        summary_folder: str = "results/summary",
        summary_name: str = "all_results",
    ) -> None:

        """
        Args:
        - layer_folder: str = "results", input layer wise result folder
        - summary_folder: str = "results/summary", output summary folder
        - summary_name: str = "all_results", name of output summary csv
        """

        self._layer_folder = os.path.normpath(layer_folder)
        self._summary_folder = checkNgen_folder(os.path.normpath(summary_folder))
        self._summary_name = summary_name

        self._full_results_df = self._summary_layer()

        # save master df
        print(f"Saving {self.summary_csv_path}...")
        self._full_results_df.to_csv(self.summary_csv_path, index=False)

    def _summary_layer(
        self,
    ):
        """
        A function for summary layer wise results

        ptp = pre-train percent
        """

        # init dataframe
        master_results = pd.DataFrame(
            columns=["arch", "task", "model", "ablation", "ptp", "metric", "value"]
        )

        # make value np array compatible
        master_results["value"] = master_results["value"].astype("object")

        for arch in PRETRAIN_ARCH_LIST:

            if arch == "esm":
                add_checkpoint = False
            else:
                add_checkpoint = True

            for ds_model in DS_MODEL_LIST:

                print(f"Analyzing pretrian {arch} with downstream {ds_model}...")

                layerloss = LayerLoss(
                    input_path=os.path.join(self._layer_folder, f"{ds_model}-{arch}"),
                    output_path=os.path.join(
                        self._layer_folder, f"{ds_model}-{arch}_layer"
                    ),
                    add_checkpoint=add_checkpoint,
                )

                # init ablation list
                ablation_list = DEFAULT_AB_LIST

                ablation_dict_list = [
                    layerloss.layer_analysis_dict,
                    layerloss.rand_layer_analysis_dict,
                    layerloss.stat_layer_analysis_dict,
                    layerloss.onehot_baseline_dict,
                ]

                # add checkpoints
                if arch == "carp":
                    cp_list = list(layerloss.checkpoint_analysis_dict.keys())
                    ablation_list += [f"{arch}-{str(cp)}" for cp in cp_list]
                    ablation_dict_list += [
                        layerloss.checkpoint_analysis_dict[cp] for cp in cp_list
                    ]

                for (ablation, ablation_dict) in zip(ablation_list, ablation_dict_list):
                    
                    rename_ablation = ablation

                    if ablation == "emb":
                        ptp = 1
                    # for carp checkpoints
                    elif arch == "carp" and "carp" in ablation:
                        rename_ablation = "emb"
                        ptp = float(ablation.split("-")[-1])
                    else:
                        ptp = 0

                    for task in ablation_dict.keys():
                        for model in ablation_dict[task].keys():
                            for metric in ablation_dict[task][model].keys():

                                # update metric and task if ss3
                                rename_metric = metric_simplifier(metric)
                                test_name = metric.split("_")[0]

                                if test_name in STRUCT_TESTS:
                                    rename_metric = rename_metric.replace(test_name, "test")

                                    # before update task: structure_ss3_tape_processed_noflatten
                                    if "tape_processed" in task:
                                        rename_task = task.replace(
                                            "tape_processed", test_name
                                        )
                                    else:
                                        split_list = task.split("_")
                                        rename_task = "_".join(
                                            split_list[:-1]
                                            + [test_name]
                                            + split_list[-1:]
                                        )
                                else:
                                    rename_task = task

                                master_results = pd.concat(
                                    [
                                        master_results,
                                        pd.DataFrame(
                                            {
                                                "arch": arch,
                                                "task": rename_task,
                                                "model": model,
                                                "ablation": rename_ablation,
                                                "ptp": ptp,
                                                "metric": rename_metric,
                                                "value": [
                                                    list(
                                                        ablation_dict[task][model][
                                                            metric
                                                        ]
                                                    )
                                                ],
                                            }
                                        ),
                                    ],
                                    ignore_index=True,
                                )

        return master_results

    @property
    def summary_df(self) -> pd.DataFrame:
        """Return appended summary results"""
        return self._full_results_df

    @property
    def summary_csv_path(self) -> str:
        """Return summary csv path"""

        summary_csv_path = os.path.join(
            self._summary_folder, self._summary_name + ".csv"
        )

        if os.path.exists(summary_csv_path):
            print(f"Delete existing {summary_csv_path}...")
            os.remove(summary_csv_path)

        return summary_csv_path
```
