## Supplemental data and code for "Feature Reuse and Scaling: Understanding Transfer Learning with Protein Language Models": utils.py

```
"""Utils for the results analysis"""

from __future__ import annotations

from scr.params.sys import DEFAULT_SPLIT
 
METRIC_DICT = {
            "proeng": ["train_mse", "val_mse", "test_mse", "test_ndcg", "test_rho"],
            "annotation": [
                "train_cross-entropy",
                "val_cross-entropy",
                "test_cross-entropy",
                "test_acc",
                "test_rocauc",
            ],
            "structure": [
                "train_cross-entropy",
                "val_cross-entropy",
                "casp12_cross-entropy",
                "casp12_acc",
                "casp12_rocauc",
                "cb513_cross-entropy",
                "cb513_acc",
                "cb513_rocauc",
                "ts115_cross-entropy",
                "ts115_acc",
                "ts115_rocauc",
            ],
        }

METRIC_TYPE = ["loss", "performance_1", "performance_2"] 

SIMPLE_METRIC_LIST = [f"{s}_{t}" for s in DEFAULT_SPLIT for t in METRIC_TYPE]

STRUCT_TESTS = ["casp12", "cb513", "ts115"]

# default ablation list
DEFAULT_AB_LIST = ["emb", "rand", "stat", "onehot"]

# default rand stat init dict
INIT_DICT = {
    "rand": "random init",
    "stat": "stat transfer"
}

INIT_SIMPLE_LIST = list(INIT_DICT.keys())

# pretrain arch list
PRETRAIN_ARCH_LIST = ["carp", "esm"]

# downstream model list
DS_MODEL_LIST = ["sklearn", "pytorch"]

# def a func to convert to metric to simper lables
def metric_simplifier(metric_label: str) -> str:

    """
    A function for shortening the metric lables for summary

    ie. 
    'train_mse'             --> 'train_loss'
    'train_ndcg'            --> 'train_performance_1'
    'train_rho'             --> 'train_performance_2'

    'train_cross-entropy'   --> 'train_loss'
    'train_rocauc'          --> 'train_performance_1'
    'train_acc'             --> 'train_performance_2'
    """

    split, metric = metric_label.split("_")

    if metric in ["mse", "cross-entropy"]:
        return "_".join([split, "loss"])
    elif metric in ["ndcg", "rocauc"]:
        return "_".join([split, "performance_1"])
    elif metric in ["rho", "acc"]:
        return "_".join([split, "performance_2"])
```
