## Supplemental data and code for "Feature Reuse and Scaling: Understanding Transfer Learning with Protein Language Models": encoding_classes.py

```
"""Add encoding classes with class methods"""

from __future__ import annotations

from abc import ABC, abstractmethod
from typing import Collection
from collections import Iterable, Sequence, OrderedDict

import os
import math
import random
import numpy as np
from tqdm import tqdm

import torch
from torch.nn import Parameter
from torch.nn.init import (
    xavier_uniform_,
    xavier_normal_,
    kaiming_uniform_,
    uniform_,
    normal_,
    constant_,
    ones_,
    zeros_,
    _calculate_fan_in_and_fan_out,
)
from sequence_models.pretrained import load_model_and_alphabet

from scr.params.aa import AA_NUMB, AA_TO_IND
from scr.params.emb import TRANSFORMER_INFO, CARP_INFO, CARP_CHECKPOINTS
from scr.params.sys import DEVICE, RAND_SEED


# seed everything
random.seed(RAND_SEED)
np.random.seed(RAND_SEED)
torch.manual_seed(RAND_SEED)
torch.cuda.manual_seed(RAND_SEED)
torch.cuda.manual_seed_all(RAND_SEED)
torch.backends.cudnn.deterministic = True


def cal_bound(model: torch.nn.Module, layer_name: str):
    """Return bound for reinit given model and layer name"""
    assert "bias" in layer_name, f"no bias in {layer_name}"
    fan_in, _ = _calculate_fan_in_and_fan_out(
        model.state_dict()[layer_name.replace("bias", "weight")]
    )
    return 1 / math.sqrt(fan_in) if fan_in > 0 else 0


class AbstractEncoder(ABC):
    """
    An abstract encoder class to fill in for different kinds of encoders

    All encoders will have an "encode" function
    """

    def __init__(
        self,
        encoder_name: str = "",
        reset_param: bool = False,
        resample_param: bool = False,
    ):

        """
        Args:
        - encoder_name: str, the name of the encoder, default empty for onehot
        - reset_param: bool = False, if update the full model to xavier_uniform_
        - resample_param: bool = False, if update the full model to xavier_normal_
        """

        self._encoder_name = encoder_name

        assert reset_param * resample_param != 1, "Choose reset OR resample param"

        self._reset_param = reset_param
        self._resample_param = resample_param

    def reset_resample_param(self, model: torch.nn.Module):
        """
        Initiate parameters in the PyTorch model. Following:
        https://pytorch.org/docs/stable/_modules/torch/nn/modules/transformer.html#Transformer

        Args:
        - model: torch.nn.Module, the input model

        Returns:
        - torch.nn.Module, the model with all params set with xavier_uniform
        """

        s = 0
        for p in model.parameters():
            s += np.sum(p.cpu().data.numpy())
        print(f"all param sum inside reset_resampel_param for input model: {s}")

        if self._reset_param:
            print(f"Reinit params for {self._encoder_name} ...")

            """
            ESM1b:

            layers.n.self_attn.k_proj.weight: dim 2         [nn.init.xavier_uniform_(self.k_proj.weight, gain=1 / math.sqrt(2))]
            layers.n.self_attn.v_proj.weight: dim 2         [nn.init.xavier_uniform_(self.v_proj.weight, gain=1 / math.sqrt(2))]
            layers.n.self_attn.q_proj.weight: dim 2         [nn.init.xavier_uniform_(self.q_proj.weight, gain=1 / math.sqrt(2))]
            layers.n.self_attn.out_proj.weight: dim 2       [nn.init.xavier_uniform_(self.out_proj.weight)]
            
            layers.n.self_attn.k_proj.bias: dim 1           [nn.init.uniform_(self.bias, -bound, bound)]
            layers.n.self_attn.v_proj.bias: dim 1           [nn.init.uniform_(self.bias, -bound, bound)]
            layers.n.self_attn.q_proj.bias: dim 1           [nn.init.uniform_(self.bias, -bound, bound)]
            
            layers.n.self_attn.out_proj.bias: dim 1         [nn.init.constant_(self.out_proj.bias, 0.0)]
            
            layers.n.self_attn_layer_norm.weight: dim 1     [nn.Parameter(torch.ones(hidden_size))]
            layers.n.final_layer_norm.weight: dim 1         [nn.Parameter(torch.ones(hidden_size))]
            emb_layer_norm_before.weight: dim 1             [nn.Parameter(torch.ones(hidden_size))]
            emb_layer_norm_after.weight: dim 1              [nn.Parameter(torch.ones(hidden_size))]
            
            layers.n.self_attn_layer_norm.bias: dim 1       [nn.Parameter(torch.zeros(hidden_size))]
            layers.n.final_layer_norm.bias: dim 1           [nn.Parameter(torch.zeros(hidden_size))]
            emb_layer_norm_before.bias: dim 1               [nn.Parameter(torch.zeros(hidden_size))]
            emb_layer_norm_after.bias: dim 1                [nn.Parameter(torch.zeros(hidden_size))]
            
            layers.n.fc1.weight: dim 2                      [nn.init.kaiming_uniform_(self.weight, a=math.sqrt(5))]
            layers.n.fc2.weight: dim 2                      [nn.init.kaiming_uniform_(self.weight, a=math.sqrt(5))]
            contact_head.regression.weight: dim 2           [nn.init.kaiming_uniform_(self.weight, a=math.sqrt(5))]
            
            layers.n.fc1.bias: dim 1                        [nn.init.uniform_(self.bias, -bound, bound)]
            layers.n.fc2.bias: dim 1                        [nn.init.uniform_(self.bias, -bound, bound)]
            contact_head.regression.bias: dim 1             [nn.init.uniform_(self.bias, -bound, bound)]
            
            embed_positions.weight: dim 2                   [nn.init.normal_(self.weight)]
            """
            if self._encoder_name in TRANSFORMER_INFO.keys():
                print(f"Updating esm {self._encoder_name} weights...")
                for layer_name, p in model.state_dict().items():
                    # what esm1b and esm1 have in common
                    if "_proj" in layer_name:
                        if "weight" in layer_name:
                            if "out" in layer_name:
                                xavier_uniform_(p)
                            else:
                                xavier_uniform_(p, gain=1 / math.sqrt(2))
                        elif "bias" in layer_name:
                            if "out" in layer_name:
                                constant_(p, 0.0)
                            else:
                                bound = cal_bound(model=model, layer_name=layer_name)
                                uniform_(p, -bound, bound)

                    # esm1b enced up using LayerNorm so the same
                    if "layer_norm" in layer_name:
                        if "weight" in layer_name:
                            Parameter(torch.ones_like(p))
                        elif "bias" in layer_name:
                            Parameter(torch.zeros_like(p))

                    if ("layers" and "fc" in layer_name) or (
                        "contact_head" in layer_name
                    ):
                        if "weight" in layer_name:
                            kaiming_uniform_(p, a=math.sqrt(5))
                        elif "bias" in layer_name:
                            bound = cal_bound(model=model, layer_name=layer_name)
                            uniform_(p, -bound, bound)

                    if "esm1b_" in self._encoder_name:

                        if "embed_positions" in layer_name:
                            normal_(p)

                        if layer_name == "lm_head.weight":
                            xavier_uniform_(p)

                        if layer_name == "lm_head.bias" or "lm_head.layer_norm.bias":
                            Parameter(torch.zeros_like(p))

                        if "dense" in layer_name:
                            if "weight" in layer_name:
                                kaiming_uniform_(p, a=math.sqrt(5))
                            elif "bias" in layer_name:
                                bound = cal_bound(model=model, layer_name=layer_name)
                                uniform_(p, -bound, bound)

                    elif "esm1_" and "bias_" in self._encoder_name:
                        xavier_normal_(p)

            elif self._encoder_name in CARP_INFO.keys():
                print(f"Updating carp {self._encoder_name} weights...")
                for layer_name, p in model.state_dict().items():
                    if "layers" in layer_name:
                        if "conv" in layer_name:
                            if "weight" in layer_name:
                                kaiming_uniform_(p, a=math.sqrt(5))
                            elif "bias" in layer_name:
                                fan_in, _ = _calculate_fan_in_and_fan_out(
                                    model.state_dict()[
                                        layer_name.replace("bias", "weight")
                                    ]
                                )
                                if fan_in != 0:
                                    bound = 1 / math.sqrt(fan_in)
                                    uniform_(p, -bound, bound)

                        else:
                            if "weight" in layer_name:
                                ones_(p)
                            elif "bias" in layer_name:
                                zeros_(p)
        
        elif self._resample_param:
            print(f"Resample params for {self._encoder_name} ...")

            resample_state = model.state_dict()

            if self._encoder_name in TRANSFORMER_INFO.keys():
                print(f"Updating esm {self._encoder_name} weights...")
                for layer_name, p in model.state_dict().items():
                    if (
                        ("embed_tokens" not in layer_name)
                        and ("embed_out" not in layer_name)
                        and ("_float_tensor" not in layer_name)
                    ):
                        # shuffle all dim
                        resample_state[layer_name] = p.view(-1)[
                            torch.randperm(p.view(-1).shape[0])
                        ].view(p.shape)

                        # if len(p.shape) == 1:
                        #     resample_state[layer_name] = p[torch.randperm(p.shape[0])]

                        #     """
                        #     esm1:
                        #     layers.n.self_attn.k_proj.bias: torch.Size([embed_dim])
                        #     layers.n.self_attn.v_proj.bias: torch.Size([embed_dim])
                        #     layers.n.self_attn.q_proj.bias: torch.Size([embed_dim])
                        #     layers.n.self_attn.out_proj.bias: torch.Size([embed_dim])
                        #     layers.n.self_attn_layer_norm.weight: torch.Size([embed_dim])
                        #     layers.n.self_attn_layer_norm.bias: torch.Size([embed_dim])
                        #     layers.n.fc1.bias: torch.Size([fc_dim])
                        #     layers.n.fc2.bias: torch.Size([embed_dim])
                        #     layers.n.final_layer_norm.weight: torch.Size([embed_dim])
                        #     layers.n.final_layer_norm.bias: torch.Size([embed_dim])

                        #     esm1b:
                        #     layers.n.self_attn.k_proj.bias: torch.Size([1280])
                        #     layers.n.self_attn.v_proj.bias: torch.Size([1280])
                        #     layers.n.self_attn.q_proj.bias: torch.Size([1280])
                        #     layers.n.self_attn.out_proj.bias: torch.Size([1280])
                        #     layers.n.self_attn_layer_norm.weight: torch.Size([1280])
                        #     layers.n.self_attn_layer_norm.bias: torch.Size([1280])
                        #     layers.n.fc1.bias: torch.Size([5120])
                        #     layers.n.fc2.bias: torch.Size([1280])
                        #     layers.n.final_layer_norm.weight: torch.Size([1280])
                        #     layers.n.final_layer_norm.bias: torch.Size([1280])
                        #     contact_head.regression.weight: torch.Size([1, 660])
                        #     contact_head.regression.bias: torch.Size([1])
                        #     emb_layer_norm_before.weight: torch.Size([1280])
                        #     emb_layer_norm_before.bias: torch.Size([1280])
                        #     emb_layer_norm_after.weight: torch.Size([1280])
                        #     emb_layer_norm_after.bias: torch.Size([1280])
                        #     lm_head.bias: torch.Size([33])
                        #     lm_head.dense.bias: torch.Size([1280])
                        #     lm_head.layer_norm.weight: torch.Size([1280])
                        #     lm_head.layer_norm.bias: torch.Size([1280])
                        #     """

                        # elif 1 in p.shape:

                        #     """
                        #     esm1
                        #     layers.n.self_attn.bias_k: torch.Size([1, 1, embed_dim])
                        #     layers.n.self_attn.bias_v: torch.Size([1, 1, embed_dim])
                        #     contact_head.regression.weight: torch.Size([1, reg_dim])

                        #     esm1b:
                        #     contact_head.regression.weight: torch.Size([1, 660])
                        #     contact_head.regression.bias: torch.Size([1])
                        #     """
                        #     if "bias_" in layer_name:
                        #         resample_state[layer_name] = p[
                        #             :, :, torch.randperm(self._embed_dim)
                        #         ]
                        #     elif "regression.weight" in layer_name:
                        #         resample_state[layer_name] = p[
                        #             :, torch.randperm(p.shape[-1])
                        #         ]

                        # elif (
                        #     "k_proj.weight"
                        #     or "q_proj.weight"
                        #     or "fc1.weight"
                        #     or "embed_positions.weight"
                        #     or "lm_head.weight" in layer_name
                        # ):
                        #     resample_state[layer_name] = p[torch.randperm(p.shape[0]), :]

                        #     """
                        #     esm1:
                        #     layers.n.self_attn.k_proj.weight: torch.Size([embed_dim, embed_dim])
                        #     layers.n.self_attn.q_proj.weight: torch.Size([embed_dim, embed_dim])
                        #     layers.n.fc1.weight: torch.Size([fc_dim, embed_dim])

                        #     esm1b:
                        #     layers.n.self_attn.k_proj.weight: torch.Size([1280, 1280])
                        #     layers.n.self_attn.q_proj.weight: torch.Size([1280, 1280])
                        #     layers.n.fc1.weight: torch.Size([5120, 1280])
                        #     embed_positions.weight: torch.Size([1026, 1280])
                        #     lm_head.weight: torch.Size([33, 1280])
                        #     """

                        # elif (
                        #     "v_proj.weight"
                        #     or "out_proj.weight"
                        #     or "fc2.weight"
                        #     or "lm_head.dense.weight" in layer_name
                        # ):

                        #     """
                        #     layers.n.self_attn.v_proj.weight: torch.Size([embed_dim, embed_dim])
                        #     layers.n.self_attn.out_proj.weight: torch.Size([embed_dim, embed_dim])
                        #     layers.n.fc2.weight: torch.Size([embed_dim, fc_dim])

                        #     esm1b:
                        #     layers.n.self_attn.v_proj.weight: torch.Size([1280, 1280])
                        #     layers.n.self_attn.out_proj.weight: torch.Size([1280, 1280])
                        #     layers.n.fc2.weight: torch.Size([1280, 5120])
                        #     lm_head.dense.weight: torch.Size([1280, 1280])
                        #     """

                        #     resample_state[layer_name] = p[:, torch.randperm(p.shape[1])]

            elif self._encoder_name in CARP_INFO.keys():
                print(f"Updating carp {self._encoder_name} weights...")
                for layer_name, p in model.state_dict().items():
                    # completely shuffle all weight matrix entries
                    if "layers" in layer_name:
                        resample_state[layer_name] = p.view(-1)[
                            torch.randperm(p.view(-1).shape[0])
                        ].view(p.shape)

            model.load_state_dict(resample_state)

        else:
            print("Not changing the model")

        s = 0
        for p in model.parameters():
            s += np.sum(p.cpu().data.numpy())
        print(f"all param sum after reset_resampel_param before return model: {s}")

        return model

    def encode(
        self,
        mut_seqs: Sequence[str] | str,
        batch_size: int = 0,
        flatten_emb: bool | str = False,
        mut_names: Sequence[str] | str | None = None,
    ) -> Iterable[np.ndarray]:
        """
        A function takes a list of sequences to yield a batch of encoded elements

        Args:
        - mut_seqs: list of str or str, mutant sequences of the same length
        - batch_size: int, set to 0 to encode all in a single batch
        - flatten_emb: bool or str, if and how (one of ["max", "mean"]) to flatten the embedding
        - mut_names: list of str or str or None, mutant names

        Returns:
        - generator: dict with layer number as keys and
            encoded flattened sequence with or without labels as value
        """

        if isinstance(mut_seqs, str):
            mut_seqs = [mut_seqs]

        # If the batch size is 0, then encode all at once in a single batch
        if batch_size == 0:
            yield self._encode_batch(
                mut_seqs=mut_seqs, flatten_emb=flatten_emb, mut_names=mut_names
            )

        # Otherwise, yield chunks of encoded sequence
        else:

            for i in tqdm(range(0, len(mut_seqs), batch_size)):

                # figure out what mut_names to feed in
                if mut_names is None:
                    mut_name_batch = mut_names
                else:
                    mut_name_batch = mut_names[i : i + batch_size]

                yield self._encode_batch(
                    mut_seqs=mut_seqs[i : i + batch_size],
                    flatten_emb=flatten_emb,
                    mut_names=mut_name_batch,
                )

    def flatten_encode(
        self,
        encoded_mut_seqs: np.ndarray,
        flatten_emb: bool | str,
        mut_seqs: Sequence[str] | str,
    ) -> np.ndarray:
        """
        Flatten the embedding or just return the encoded mutants.

        Args:
        - encoded_mut_seqs: np.ndarray, shape [batch_size, seq_len, embed_dim]
        - flatten_emb: bool or str, if and how (one of ["max", "mean"]) to flatten the embedding
            - True -> shape [batch_size, seq_len * embed_dim]
            - "max" or "mean" -> shape [batch_size, embed_dim]
            - False or everything else -> [batch_size, seq_len, embed_dim]

        Returns:
        - np.ndarray, shape depends on flatten_emb parameter
        """

        assert (
            encoded_mut_seqs.shape[-1] == self._embed_dim
        ), f"encode last dim {encoded_mut_seqs.shape[-1]} != embed dim {self._embed_dim}"

        if flatten_emb in [True, "flatten", "flattened", ""]:
            # shape [batch_size, seq_len * embed_dim]
            return encoded_mut_seqs.reshape(encoded_mut_seqs.shape[0], -1)

        elif isinstance(flatten_emb, str):
            # init out put seq_reps should be in dim [batch_size, embed_dim]
            seq_reps = np.empty((encoded_mut_seqs.shape[0], self._embed_dim))
            for i, encoded_mut_seq in enumerate(encoded_mut_seqs):
                
                # if the emb has label from esm
                if len(mut_seqs[i]) == 2:
                    seq_len = len(mut_seqs[i][1])
                # if the emb is carp
                elif len(mut_seqs[i]) == 1:
                    seq_len = len(mut_seqs[i][0])
                else:
                    seq_len = len(mut_seqs[i])

                assert seq_len not in [1, 2], "Check emb pooling len!"

                if flatten_emb == "mean":
                    seq_reps[i] = encoded_mut_seq[: seq_len].mean(0)
                elif flatten_emb == "max":
                    seq_reps[i] = encoded_mut_seq[: seq_len].max(0)

            return seq_reps

        else:
            # print("No embedding flattening")
            # [batch_size, seq_len, embed_dim]
            return encoded_mut_seqs

    @abstractmethod
    def _encode_batch(
        mut_seqs: Sequence[str] | str,
        flatten_emb: bool | str,
        mut_names: Sequence[str] | str | None = None,
    ) -> np.ndarray:
        """
        Encode a single batch of mut_seqs
        """
        pass

    @property
    def embed_dim(self) -> int:
        """The dim of the embedding"""
        return self._embed_dim

    @property
    def max_emb_layer(self) -> int:
        """The max layer nubmer of the embedding"""
        return self._max_emb_layer

    @property
    def encoder_name(self) -> str:
        """The name of the encoding method"""
        return self._encoder_name


class OnehotEncoder(AbstractEncoder):
    """
    Build a onehot encoder
    """

    def __init__(
        self,
        max_seq_len: int,
        encoder_name: str = "",
        reset_param: bool = False,
        resample_param: bool = False,
    ):
        """
        Args
        - encoder_name: str, the name of the encoder, one of the keys of CARP_INFO
        - max_seq_len: int, the longest sequence length
        - reset_param: bool = False, if update the full model to xavier_uniform_
        - resample_param: bool = False, if update the full model to xavier_normal_
        """
        super().__init__(encoder_name, reset_param, resample_param)

        self.max_seq_len = max_seq_len

        if encoder_name not in (TRANSFORMER_INFO.keys() and CARP_INFO.keys()):
            self._encoder_name = "onehot"
            self._embed_dim, self._max_emb_layer = AA_NUMB, 0
            self._include_input_layer = True

        # load model from torch.hub
        print(
            f"Generating {self._encoder_name} upto {self._max_emb_layer} layer embedding ..."
        )

        if reset_param or resample_param:
            self._reset_param = False
            self._resample_param = False
            print(
                f"Onehot encoding reset or resample param not allowed. /n \
                    Setting both to {self._reset_param} ..."
            )

    def _encode_batch(
        self,
        mut_seqs: Sequence[str] | str,
        flatten_emb: bool | str,
        mut_names: Sequence[str] | str | None = None,
    ) -> np.ndarray:

        encoded_mut_seqs = []

        for mut_seq in mut_seqs:
            # padding: (top, bottom), (left, right)
            encoded_mut_seqs.append(
                np.pad(
                    np.array(np.eye(AA_NUMB)[[AA_TO_IND[aa] for aa in mut_seq]]),
                    pad_width=((0, self.max_seq_len - len(mut_seq)), (0, 0)),
                )
            )

        return {
            0: self.flatten_encode(
                encoded_mut_seqs=np.array(encoded_mut_seqs),
                flatten_emb=flatten_emb,
                mut_seqs=mut_seqs,
            )
        }


class ESMEncoder(AbstractEncoder):
    """
    Build an ESM encoder
    """

    def __init__(
        self,
        encoder_name: str,
        reset_param: bool = False,
        resample_param: bool = False,
        iftrimCLS: bool = True,
        iftrimEOS: bool = True,
    ):
        """
        Args
        - encoder_name: str, the name of the encoder, one of the keys of TRANSFORMER_INFO
        - reset_param: bool = False, if update the full model to xavier_uniform_
        - resample_param: bool = False, if update the full model to xavier_normal_
        - iftrimCLS: bool, whether to trim the first classifification token
        - iftrimEOS: bool, whether to trim the end of sequence token, if exists
        """

        super().__init__(encoder_name, reset_param, resample_param)

        self._iftrimCLS = iftrimCLS
        self._iftrimEOS = iftrimEOS

        # get transformer dim and layer info
        self._embed_dim, self._max_emb_layer, _ = TRANSFORMER_INFO[self._encoder_name]

        # esm has the input representation
        self._include_input_layer = True

        # load model from torch.hub
        print(
            f"Generating {self._encoder_name} upto {self._max_emb_layer} layer embedding ..."
        )

        self.model, self.alphabet = torch.hub.load(
            "facebookresearch/esm:main", model=self._encoder_name
        )
        self.batch_converter = self.alphabet.get_batch_converter()

        s = 0
        for p in self.model.parameters():
            s += np.sum(p.cpu().data.numpy())
        print(f"all param sum for loading init esm from hub in ESMEncoder: {s}")

        # if reset or resample weights
        self.model = self.reset_resample_param(model=self.model)

        # set model to eval mode
        self.model.eval()
        self.model.to(DEVICE)

        s = 0
        for p in self.model.parameters():
            s += np.sum(p.cpu().data.numpy())
        print(f"all param sum for after reset_resample_param in ESMEncoder: {s}")

        expected_num_layers = int(self._encoder_name.split("_")[-3][1:])
        assert (
            expected_num_layers == self._max_emb_layer
        ), "Wrong ESM model name or layer"

    def _encode_batch(
        self,
        mut_seqs: Sequence[str] | str,
        flatten_emb: bool | str,
        mut_names: Sequence[str] | str | None = None,
    ) -> np.ndarray:
        """
        Encodes a batch of mutant sequences.

        Args:
        - mut_seqs: list of str or str, mutant sequences of the same length
        - flatten_emb: bool or str, if and how (one of ["max", "mean"]) to flatten the embedding
        - mut_names: list of str or str or None, mutant names

        Returns:
        - np.ndarray or a tuple(np.ndarray, list[str]) where the list is batch_labels
        """

        if isinstance(mut_names, str):
            mut_names = [mut_names]

        # pair the mut_names and mut_seqs
        if mut_names is not None:
            assert len(mut_names) == len(
                mut_seqs
            ), "mutant_name and mut_seqs different length"
            mut_seqs = [(n, m) for (n, m) in zip(mut_names, mut_seqs)]
        else:
            mut_seqs = [("", m) for m in mut_seqs]

        # convert raw mutant sequences to tokens
        batch_labels, _, batch_tokens = self.batch_converter(mut_seqs)
        batch_tokens = batch_tokens.to(DEVICE)

        # Turn off gradients and pass the batch through
        with torch.no_grad():
            # shape [batch_size, seq_len + pad, embed_dim]

            dict_encoded_mut_seqs = self.model(
                batch_tokens, repr_layers=list(range(self._max_emb_layer + 1))
            )["representations"]

        for layer, encoded_mut_seqs in dict_encoded_mut_seqs.items():

            encoded_mut_seqs = encoded_mut_seqs.cpu().numpy()
            # https://github.com/facebookresearch/esm/blob/main/esm/data.py
            # from_architecture

            # trim off initial classification token [CLS]
            # both "ESM-1" and "ESM-1b" have prepend_bos = True
            if self._iftrimCLS and self._encoder_name.split("_")[0] in [
                "esm1",
                "esm1b",
            ]:
                encoded_mut_seqs = encoded_mut_seqs[:, 1:, :]

            # trim off end-of-sequence token [EOS]
            # only "ESM-1b" has append_eos = True
            if self._iftrimEOS and self._encoder_name.split("_")[0] == "esm1b":
                encoded_mut_seqs = encoded_mut_seqs[:, :-1, :]

            if mut_names is not None:
                dict_encoded_mut_seqs[layer] = (
                    self.flatten_encode(
                        encoded_mut_seqs=encoded_mut_seqs,
                        flatten_emb=flatten_emb,
                        mut_seqs=mut_seqs,
                    ),
                    batch_labels,
                )
            else:
                dict_encoded_mut_seqs[layer] = self.flatten_encode(
                    encoded_mut_seqs=encoded_mut_seqs,
                    flatten_emb=flatten_emb,
                    mut_seqs=mut_seqs,
                )

        return dict_encoded_mut_seqs


class CARPEncoder(AbstractEncoder):
    """
    Build a CARP encoder
    """

    def __init__(
        self,
        encoder_name: str,
        checkpoint: float = 1,
        checkpoint_folder: str = "pretrain_checkpoints/carp",
        reset_param: bool = False,
        resample_param: bool = False,
    ):
        """
        Args
        - encoder_name: str, the name of the encoder, one of the keys of CARP_INFO
        - checkpoint: float = 1, the 0.5, 0.25, 0.125 checkpoint of the CARP encoder or full
        - checkpoint_folder: str = "pretrain_checkpoints/carp", folder for carp encoders
        - reset_param: bool = False, if update the full model to xavier_uniform_
        - resample_param: bool = False, if update the full model to xavier_normal_
        """

        super().__init__(encoder_name, reset_param, resample_param)

        self.model, self.collater = load_model_and_alphabet(self._encoder_name)

        s = 0
        for p in self.model.parameters():
            s += np.sum(p.cpu().data.numpy())
        print(f"all param sum for loading init carp from hub in CARPEncoder: {s}")

        # load checkpoint unless default to full
        if checkpoint != 1:

            # get the checkpoint number from the CARP_CHECKPOINTS dict
            # ie {"carp_600k": {"1/2": 239263, ...}, ...}
            # to get 'pretrain_checkpoints/carp/carp_600k/checkpoint239263.tar'

            checkpoint_path = (
                f"{os.path.normpath(checkpoint_folder)}/{encoder_name}/"
                f"checkpoint{str(CARP_CHECKPOINTS[encoder_name][checkpoint])}.tar"
            )
            
            print(
                f"Loading {encoder_name} {checkpoint} checkpoint from {checkpoint_path}..."
            )

            # get the dict with dict_keys(['model_state_dict', ...])
            checkpoint_dict = torch.load(checkpoint_path, map_location=DEVICE)

            self.model.load_state_dict(
                OrderedDict(
                    [
                        (k.replace("module", "model"), v) if "module" in k else (k, v)
                        for k, v in checkpoint_dict["model_state_dict"].items()
                    ]
                )
            )
        else:
            print("Running on fully trained model...")
        
        s = 0
        for p in self.model.parameters():
            s += np.sum(p.cpu().data.numpy())

        print(f"all param sum for after carp checkpoint loading before reset_resample_param in CARPEncoder: {s}")

        # if reset or resample weights
        self.model = self.reset_resample_param(model=self.model)

        s = 0
        for p in self.model.parameters():
            s += np.sum(p.cpu().data.numpy())
        print(f"all param sum for after carp checkpoint loading after reset_resample_param in CARPEncoder: {s}")

        # set model to eval mode
        self.model.eval()
        self.model.to(DEVICE)

        self._embed_dim, self._max_emb_layer = CARP_INFO[self._encoder_name]

        # load model from torch.hub
        print(
            f"Generating {self._encoder_name} upto {self._max_emb_layer} layer embedding ..."
        )

    def _encode_batch(
        self,
        mut_seqs: Sequence[str] | str,
        flatten_emb: bool | str,
        mut_names: Sequence[str] | str | None = None,
    ) -> np.ndarray:
        """
        Encodes a batch of mutant sequences.

        Args:
        - mut_seqs: list of str or str, mutant sequences of the same length
        - flatten_emb: bool or str, if and how (one of ["max", "mean"]) to flatten the embedding
        - mut_names: list of str or str or None, mutant names

        Returns:
        - np.ndarray or a tuple(np.ndarray, list[str]) where the list is batch_labels
        """

        mut_seqs = [[m] for m in mut_seqs]

        x = self.collater(mut_seqs)[0].to(DEVICE)
        rep = self.model(x, repr_layers=list(range(self._max_emb_layer + 1)))

        # init output dict
        dict_encoded_mut_seqs = {}

        dict_encoded_mut_seqs[0] = self.flatten_encode(
            encoded_mut_seqs=rep[0].detach().cpu().numpy(),
            flatten_emb=flatten_emb,
            mut_seqs=mut_seqs,
        )

        for layer_numb, encoded_mut_seqs in rep["representations"].items():
            dict_encoded_mut_seqs[layer_numb] = self.flatten_encode(
                encoded_mut_seqs=encoded_mut_seqs.detach().cpu().numpy(),
                flatten_emb=flatten_emb,
                mut_seqs=mut_seqs,
            )
        return dict_encoded_mut_seqs

def get_emb_info(encoder_name: str) -> Collection(str, AbstractEncoder, int):

    """
    A function return processed encoder_name and total_emb_layer

    Args:
    - encoder_name: str, input encoder_name

    Returns:
    - encoder_name: str, change anything not a transformer or carp encoder to onehot
    - encoder_class: AbstractEncoder, encoder class
    - total_emb_layer: int, number of embedding layers
    """

    if encoder_name in TRANSFORMER_INFO.keys():
        total_emb_layer = TRANSFORMER_INFO[encoder_name][1] + 1
        encoder_class = ESMEncoder
    elif encoder_name in CARP_INFO.keys():
        total_emb_layer = CARP_INFO[encoder_name][1] + 1
        encoder_class = CARPEncoder
    else:
        # for onehot
        encoder_name = "onehot"
        encoder_class = OnehotEncoder
        total_emb_layer = 1

    return encoder_name, encoder_class, total_emb_layer
```
