## Supplemental data and code for "Feature Reuse and Scaling: Understanding Transfer Learning with Protein Language Models": gen_encoding.py

```
from __future__ import annotations

import os
import tables

import numpy as np

from scr.utils import get_folder_file_names, checkNgen_folder
from scr.params.emb import MAX_SEQ_LEN
from scr.encoding.encoding_classes import get_emb_info, OnehotEncoder, CARPEncoder
from scr.preprocess.data_process import ProtranDataset


class GenerateEmbeddings:
    """A class for generating and saving embeddings"""

    def __init__(
        self,
        dataset_path: str,
        encoder_name: str,
        checkpoint: float = 1,
        checkpoint_folder: str = "pretrain_checkpoints/carp",
        reset_param: bool = False,
        resample_param: bool = False,
        embed_batch_size: int = 128,
        flatten_emb: bool | str = False,
        seq_start_idx: bool | int = False,
        seq_end_idx: bool | int = False,
        subset_list: list[str] = ["train", "val", "test"],
        embed_folder: str = "embeddings",
        **encoder_params,
    ) -> None:

        """
        Args:
        - dataset_path: str, full path to the dataset, in pkl or panda readable format
            columns include: sequence, target, set, validation,
            mut_name (optional), mut_numb (optional)
        - encoder_name: str, the name of the encoder
        - checkpoint: float = 1, the 0.5, 0.25, 0.125 checkpoint of the CARP encoder or full
        - checkpoint_folder: str = "pretrain_checkpoints/carp", folder for carp encoders
        - reset_param: bool = False, if update the full model to xavier_uniform_
        - resample_param: bool = False, if update the full model to xavier_normal_
        - embed_batch_size: int, set to 0 to encode all in a single batch
        - flatten_emb: bool or str, if and how (one of ["max", "mean"]) to flatten the embedding
        - seq_start_idx: bool | int = False, the index for the start of the sequence
        - seq_end_idx: bool | int = False, the index for the end of the sequence
        - subset_list: list of str, train, val, test, or ss3 tasks including 'cb513', 'ts115', 'casp12'
        - embed_folder: str = "embeddings", the parent folder for embeddings
        - encoder_params: kwarg, additional parameters for encoding
        """

        self.dataset_path = dataset_path
        self.encoder_name = encoder_name
        self.reset_param = reset_param
        self.resample_param = resample_param
        self.flatten_emb = flatten_emb

        self.embed_folder = embed_folder

        # append emb info
        if checkpoint != 1:
            self.embed_folder = f"{self.embed_folder}-{str(checkpoint)}"

        # append init info
        if self.reset_param and "-rand" not in self.embed_folder:
            self.embed_folder = f"{self.embed_folder}-rand"

        if self.resample_param and "-stat" not in self.embed_folder:
            self.embed_folder = f"{self.embed_folder}-stat"

        self.encoder_name, encoder_class, total_emb_layer = get_emb_info(
            self.encoder_name
        )

        # assert encoder_class != OnehotEncoder, "Generate onehot on the fly instead"
        # add in the max_seq_len for Onehot
        if encoder_class == OnehotEncoder and self.flatten_emb != False:
            encoder_params["max_seq_len"] = MAX_SEQ_LEN
            embed_rescale = MAX_SEQ_LEN
        else:
            embed_rescale = 1

        if encoder_class == CARPEncoder:
            encoder_params["checkpoint"] = checkpoint
            encoder_params["checkpoint_folder"] = checkpoint_folder

        # get the encoder
        self._encoder = encoder_class(
            encoder_name=encoder_name,
            reset_param=reset_param,
            resample_param=resample_param,
            **encoder_params,
        )

        if self.flatten_emb == False:
            flatten_emb_name = "noflatten"
        else:
            flatten_emb_name = self.flatten_emb

        # get the folder name
        dataset_folder, _ = get_folder_file_names(
            parent_folder=self.embed_folder,
            dataset_path=self.dataset_path,
            encoder_name=self.encoder_name,
            embed_layer=0,
            flatten_emb=flatten_emb_name,
        )

        # Close all the open files
        tables.file._open_files.close_all()

        for subset in subset_list:

            print(f"Generating embedding for {subset}...")

            # get the dataset to be encoded
            ds = ProtranDataset(
                dataset_path=dataset_path,
                subset=subset,
                encoder_name=encoder_name,
                reset_param=reset_param,
                resample_param=resample_param,
                embed_batch_size=embed_batch_size,
                flatten_emb=flatten_emb,
                embed_folder=None,
                embed_layer="all",
                seq_start_idx=seq_start_idx,
                seq_end_idx=seq_end_idx,
                if_encode_all=False,
                **encoder_params,
            )

            # get the max seq len from the dataset to pad the embeddings
            self._max_seq_len = ds.max_seq_len

            # get the dim of the array to be saved
            # without flattening
            if self.flatten_emb == False:
                earray_dim = (0, self._max_seq_len, self._encoder.embed_dim)
            else:
                earray_dim = (0, self._encoder.embed_dim * embed_rescale)

            init_array_list = [None] * total_emb_layer

            file_path = os.path.join(
                checkNgen_folder(os.path.join(dataset_folder, subset)), "embedding.h5"
            )

            # check all the embedding file h5 files
            # to remove old ones before generating new ones
            if os.path.isfile(file_path):
                print("Overwritting {0}".format(file_path))
                os.remove(file_path)

            # init file open
            f = tables.open_file(file_path, mode="a")
            for emb_layer in range(total_emb_layer):
                init_array_list[emb_layer] = f.create_earray(
                    f.root, "layer" + str(emb_layer), tables.Float32Atom(), earray_dim
                )
            # f.close()

            # use the encoder generator for batch emb
            # assume no labels included
            for encoded_batch_dict in self._encoder.encode(
                mut_seqs=ds.sequence,
                batch_size=embed_batch_size,
                flatten_emb=flatten_emb,
            ):
                
                for emb_layer, emb in encoded_batch_dict.items():
                    # f = tables.open_file(file_path, mode="a")
                    # if no flattening, pad all embedding to the max_seq_len
                    if flatten_emb == False:
                        padded_emb = np.pad(
                                emb,
                                pad_width=(
                                    (0, 0),
                                    (0, self._max_seq_len - emb.shape[1]),
                                    (0, 0),
                                ),
                            )
                    else:
                        padded_emb = emb
                    getattr(f.root, "layer" + str(emb_layer)).append(padded_emb)
                    # f.close()

            f.close()
```
