## Supplemental data and code for "Feature Reuse and Scaling: Understanding Transfer Learning with Protein Language Models": README.md

# Encoding
1. [Reinit (`self._reset_param = True`) ESM1b](#reset_esm1b)
2. [Resample (`self._resample_param = True`) ESM1b](#resamp_esm1b)
3. [Reinit (`self._reset_param = True`) ESM1](#reset_esm1)
4. [Resample (`self._resample_param = True`) ESM1](#resamp_esm1)
5. [Reinit (`self._reset_param = True`) CARP](#reset_carp)
6. [Resample (`self._resample_param = True`) CARP](#resamp_carp) 

<a name="reset_esm1b"></a>
## Reinit (`self._reset_param = True`) ESM1b
### ESM1b architecture
* n = 0, ..., 32

```
ProteinBertModel(
  (embed_tokens): Embedding(33, 1280, padding_idx=1)
  (layers): ModuleList(
    (n): TransformerLayer(
      (self_attn): MultiheadAttention(
        (k_proj): Linear(in_features=1280, out_features=1280, bias=True)
        (v_proj): Linear(in_features=1280, out_features=1280, bias=True)
        (q_proj): Linear(in_features=1280, out_features=1280, bias=True)
        (out_proj): Linear(in_features=1280, out_features=1280, bias=True)
      )
      (self_attn_layer_norm): LayerNorm((1280,), eps=1e-05, elementwise_affine=True)
      (fc1): Linear(in_features=1280, out_features=5120, bias=True)
      (fc2): Linear(in_features=5120, out_features=1280, bias=True)
      (final_layer_norm): LayerNorm((1280,), eps=1e-05, elementwise_affine=True)
    )
  )
  (contact_head): ContactPredictionHead(
    (regression): Linear(in_features=660, out_features=1, bias=True)
    (activation): Sigmoid()
  )
  (embed_positions): LearnedPositionalEmbedding(1026, 1280, padding_idx=1)
  (emb_layer_norm_before): LayerNorm((1280,), eps=1e-05, elementwise_affine=True)
  (emb_layer_norm_after): LayerNorm((1280,), eps=1e-05, elementwise_affine=True)
  (lm_head): RobertaLMHead(
    (dense): Linear(in_features=1280, out_features=1280, bias=True)
    (layer_norm): LayerNorm((1280,), eps=1e-05, elementwise_affine=True)
  )
)
```
### ESM1b architecture with dimensions
* Get the layers and dim with
```
for k, p in model.state_dict().items():
    print(f"{k}: dim {p.dim()}")
```
* Returns
```
embed_tokens.weight: dim 2
layers.n.self_attn.k_proj.weight: dim 2
layers.n.self_attn.k_proj.bias: dim 1
layers.n.self_attn.v_proj.weight: dim 2
layers.n.self_attn.v_proj.bias: dim 1
layers.n.self_attn.q_proj.weight: dim 2
layers.n.self_attn.q_proj.bias: dim 1
layers.n.self_attn.out_proj.weight: dim 2
layers.n.self_attn.out_proj.bias: dim 1
layers.n.self_attn_layer_norm.weight: dim 1
layers.n.self_attn_layer_norm.bias: dim 1
layers.n.fc1.weight: dim 2
layers.n.fc1.bias: dim 1
layers.n.fc2.weight: dim 2
layers.n.fc2.bias: dim 1
layers.n.final_layer_norm.weight: dim 1
layers.n.final_layer_norm.bias: dim 1
contact_head.regression.weight: dim 2
contact_head.regression.bias: dim 1
embed_positions.weight: dim 2
emb_layer_norm_before.weight: dim 1
emb_layer_norm_before.bias: dim 1
emb_layer_norm_after.weight: dim 1
emb_layer_norm_after.bias: dim 1
lm_head.weight: dim 2
lm_head.bias: dim 1
lm_head.dense.weight: dim 2
lm_head.dense.bias: dim 1
lm_head.layer_norm.weight: dim 1
lm_head.layer_norm.bias: dim 1
```

* To reinit, follow [`ProteinBertModel`](https://github.com/facebookresearch/esm/blob/839c5b82c6cd9e18baa7a88dcbed3bd4b6d48e47/esm/model/esm1.py#L22
) class in [`esm.model.esm1`](https://github.com/facebookresearch/esm/blob/main/esm/model/esm1.py)

```
 def __init__(self, args, alphabet):
    super().__init__()
    self.args = args
    self.alphabet_size = len(alphabet)
    self.padding_idx = alphabet.padding_idx
    self.mask_idx = alphabet.mask_idx
    self.cls_idx = alphabet.cls_idx
    self.eos_idx = alphabet.eos_idx
    self.prepend_bos = alphabet.prepend_bos
    self.append_eos = alphabet.append_eos
    self.emb_layer_norm_before = getattr(self.args, "emb_layer_norm_before", False)
    if self.args.arch == "roberta_large":
        self.model_version = "ESM-1b"
        self._init_submodules_esm1b()
    else:
        self.model_version = "ESM-1"
        self._init_submodules_esm1()

def _init_submodules_common(self):
    self.embed_tokens = nn.Embedding(
        self.alphabet_size, self.args.embed_dim, padding_idx=self.padding_idx
    )
    self.layers = nn.ModuleList(
        [
            TransformerLayer(
                self.args.embed_dim,
                self.args.ffn_embed_dim,
                self.args.attention_heads,
                add_bias_kv=(self.model_version != "ESM-1b"),
                use_esm1b_layer_norm=(self.model_version == "ESM-1b"),
            )
            for _ in range(self.args.layers)
        ]
    )

    self.contact_head = ContactPredictionHead(
        self.args.layers * self.args.attention_heads,
        self.prepend_bos,
        self.append_eos,
        eos_idx=self.eos_idx,
    )

def _init_submodules_esm1b(self):
    self._init_submodules_common()
    self.embed_scale = 1
    self.embed_positions = LearnedPositionalEmbedding(
        self.args.max_positions, self.args.embed_dim, self.padding_idx
    )
    self.emb_layer_norm_before = (
        ESM1bLayerNorm(self.args.embed_dim) if self.emb_layer_norm_before else None
    )
    self.emb_layer_norm_after = ESM1bLayerNorm(self.args.embed_dim)
    self.lm_head = RobertaLMHead(
        embed_dim=self.args.embed_dim,
        output_dim=self.alphabet_size,
        weight=self.embed_tokens.weight,
    )
```

#### Reinit `layer`
* Where for [`TransformerLayer`](https://github.com/facebookresearch/esm/blob/839c5b82c6cd9e18baa7a88dcbed3bd4b6d48e47/esm/modules.py#L84) class in [`esm.modules`](https://github.com/facebookresearch/esm/blob/main/esm/modules.py),
    * `add_bias_kv=(self.model_version != "ESM-1b")` would be `False`
    * `use_esm1b_layer_norm=(self.model_version == "ESM-1b")` would be `True`
    * `BertLayerNorm` would be `ESM1bLayerNorm`

```
def _init_submodules(self, add_bias_kv, use_esm1b_layer_norm):
    BertLayerNorm = ESM1bLayerNorm if use_esm1b_layer_norm else ESM1LayerNorm

    self.self_attn = MultiheadAttention(
        self.embed_dim,
        self.attention_heads,
        add_bias_kv=add_bias_kv,
        add_zero_attn=False,
        use_rotary_embeddings=self.use_rotary_embeddings,
    )
    self.self_attn_layer_norm = BertLayerNorm(self.embed_dim)

    self.fc1 = nn.Linear(self.embed_dim, self.ffn_embed_dim)
    self.fc2 = nn.Linear(self.ffn_embed_dim, self.embed_dim)

    self.final_layer_norm = BertLayerNorm(self.embed_dim)
```

* For [`MultiheadAttention`](https://github.com/facebookresearch/esm/blob/839c5b82c6cd9e18baa7a88dcbed3bd4b6d48e47/esm/multihead_attention.py#L68) class in [`esm.multihead_attention`](https://github.com/facebookresearch/esm/blob/main/esm/multihead_attention.py)

```
def __init__(
    self,
    embed_dim,
    num_heads,
    kdim=None,
    vdim=None,
    dropout=0.0,
    bias=True,
    add_bias_kv: bool = False,
    add_zero_attn: bool = False,
    self_attention: bool = False,
    encoder_decoder_attention: bool = False,
    use_rotary_embeddings: bool = False,
):
    super().__init__()
    self.embed_dim = embed_dim
    self.kdim = kdim if kdim is not None else embed_dim
    self.vdim = vdim if vdim is not None else embed_dim
    self.qkv_same_dim = self.kdim == embed_dim and self.vdim == embed_dim

    self.num_heads = num_heads
    self.dropout = dropout
    self.head_dim = embed_dim // num_heads
    assert (
        self.head_dim * num_heads == self.embed_dim
    ), "embed_dim must be divisible by num_heads"
    self.scaling = self.head_dim**-0.5

    self.self_attention = self_attention
    self.encoder_decoder_attention = encoder_decoder_attention

    assert not self.self_attention or self.qkv_same_dim, (
        "Self-attention requires query, key and " "value to be of the same size"
    )

    self.k_proj = nn.Linear(self.kdim, embed_dim, bias=bias)
    self.v_proj = nn.Linear(self.vdim, embed_dim, bias=bias)
    self.q_proj = nn.Linear(embed_dim, embed_dim, bias=bias)

    self.out_proj = nn.Linear(embed_dim, embed_dim, bias=bias)

    if add_bias_kv:
        self.bias_k = Parameter(torch.Tensor(1, 1, embed_dim))
        self.bias_v = Parameter(torch.Tensor(1, 1, embed_dim))
    else:
        self.bias_k = self.bias_v = None

    self.add_zero_attn = add_zero_attn

    self.reset_parameters()

    ...

def reset_parameters(self):
    if self.qkv_same_dim:
        # Empirically observed the convergence to be much better with
        # the scaled initialization
        nn.init.xavier_uniform_(self.k_proj.weight, gain=1 / math.sqrt(2))
        nn.init.xavier_uniform_(self.v_proj.weight, gain=1 / math.sqrt(2))
        nn.init.xavier_uniform_(self.q_proj.weight, gain=1 / math.sqrt(2))
    else:
        nn.init.xavier_uniform_(self.k_proj.weight)
        nn.init.xavier_uniform_(self.v_proj.weight)
        nn.init.xavier_uniform_(self.q_proj.weight)

    nn.init.xavier_uniform_(self.out_proj.weight)
    if self.out_proj.bias is not None:
        nn.init.constant_(self.out_proj.bias, 0.0)
    if self.bias_k is not None:
        nn.init.xavier_normal_(self.bias_k)
    if self.bias_v is not None:
        nn.init.xavier_normal_(self.bias_v)
```
* For the `bias` in `nn.Linear`, follow the `reset_parameters` in [`torch.nn.modules.linear`](https://pytorch.org/docs/stable/_modules/torch/nn/modules/linear.html#Linear)
```
def reset_parameters(self) -> None:
    # Setting a=sqrt(5) in kaiming_uniform is the same as initializing with
    # uniform(-1/sqrt(in_features), 1/sqrt(in_features)). For details, see
    # https://github.com/pytorch/pytorch/issues/57109
    init.kaiming_uniform_(self.weight, a=math.sqrt(5))
    if self.bias is not None:
        fan_in, _ = init._calculate_fan_in_and_fan_out(self.weight)
        bound = 1 / math.sqrt(fan_in) if fan_in > 0 else 0
        init.uniform_(self.bias, -bound, bound)
```
* For [`ESM1bLayerNorm`](https://github.com/facebookresearch/esm/blob/839c5b82c6cd9e18baa7a88dcbed3bd4b6d48e47/esm/modules.py#L44) class in [`esm.modules`](https://github.com/facebookresearch/esm/blob/main/esm/modules.py)
```
try:
    from apex.normalization import FusedLayerNorm as _FusedLayerNorm

    class ESM1bLayerNorm(_FusedLayerNorm):
        @torch.jit.unused
        def forward(self, x):
            if not x.is_cuda:
                return super().forward(x)
            else:
                with torch.cuda.device(x.device):
                    return super().forward(x)

except ImportError:
    from torch.nn import LayerNorm as ESM1bLayerNorm
```
* For [`FusedLayerNorm`](https://nvidia.github.io/apex/_modules/apex/normalization/fused_layer_norm.html#FusedLayerNorm)
```
    def __init__(self, normalized_shape, eps=1e-5, elementwise_affine=True):
        super(FusedLayerNorm, self).__init__()

        global fused_layer_norm_cuda
        fused_layer_norm_cuda = importlib.import_module("fused_layer_norm_cuda")

        if isinstance(normalized_shape, numbers.Integral):
            normalized_shape = (normalized_shape,)
        self.normalized_shape = torch.Size(normalized_shape)
        self.eps = eps
        self.elementwise_affine = elementwise_affine
        if self.elementwise_affine:
            self.weight = Parameter(torch.Tensor(*normalized_shape))
            self.bias = Parameter(torch.Tensor(*normalized_shape))
        else:
            self.register_parameter('weight', None)
            self.register_parameter('bias', None)
        self.reset_parameters()

    def reset_parameters(self):
        if self.elementwise_affine:
            init.ones_(self.weight)
            init.zeros_(self.bias)
```
* ESM1b `TransformerLayer` has
    * `MultiheadAttention`
        * `kdim = None`
        * `vdim = None`
        * `bias = True`
        * `add_bias_kv = False`
        * `add_zero_attn = False`
        * `self_attention = False`
    * `BertLayerNorm`
        * `use_esm1b_layer_norm = True`
        * `affine = True`
* Thus
    * `MultiheadAttention`
        * `self.kdim = kdim if kdim is not None else embed_dim` would be `embed_dim`
        * `self.vdim = vdim if vdim is not None else embed_dim` would be `embed_dim`
        * `self.qkv_same_dim = self.kdim == embed_dim and self.vdim == embed_dim` would be `True`
        * `self.bias_k = self.bias_v = None`
    * `BertLayerNorm`
        * `BertLayerNorm = ESM1bLayerNorm` where `LayerNorm` was used
* More concretely
    * `MultiheadAttention`
        * `nn.init.xavier_uniform_(self.k_proj.weight, gain=1 / math.sqrt(2))`
        * `nn.init.xavier_uniform_(self.v_proj.weight, gain=1 / math.sqrt(2))`
        * `nn.init.xavier_uniform_(self.q_proj.weight, gain=1 / math.sqrt(2))`
        * `nn.init.xavier_uniform_(self.out_proj.weight)`
        * `nn.init.constant_(self.out_proj.bias, 0.0)`
    * `BertLayerNorm`
        * `self.weight = nn.Parameter(torch.ones(hidden_size))`
        * `self.bias = nn.Parameter(torch.zeros(hidden_size))`
#### Reinit `contact_head`
* For [`ContactPredictionHead`](https://github.com/facebookresearch/esm/blob/839c5b82c6cd9e18baa7a88dcbed3bd4b6d48e47/esm/modules.py#L317) class in [`esm.modules`](https://github.com/facebookresearch/esm/blob/main/esm/modules.py)
```
def __init__(
    self,
    in_features: int,
    prepend_bos: bool,
    append_eos: bool,
    bias=True,
    eos_idx: Optional[int] = None,
):
    super().__init__()
    self.in_features = in_features
    self.prepend_bos = prepend_bos
    self.append_eos = append_eos
    if append_eos and eos_idx is None:
        raise ValueError("Using an alphabet with eos token, but no eos token was passed in.")
    self.eos_idx = eos_idx
    self.regression = nn.Linear(in_features, 1, bias)
    self.activation = nn.Sigmoid()
```
* For ESM1b `ContactPredictionHead` has
    * `bias = True`

### Reinit `embed_positions`
* For [`LearnedPositionalEmbedding`](https://github.com/facebookresearch/esm/blob/839c5b82c6cd9e18baa7a88dcbed3bd4b6d48e47/esm/modules.py#L224) class in [`esm.modules`](https://github.com/facebookresearch/esm/blob/main/esm/modules.py)
```
class LearnedPositionalEmbedding(nn.Embedding):
    """
    This module learns positional embeddings up to a fixed maximum size.
    Padding ids are ignored by either offsetting based on padding_idx
    or by setting padding_idx to None and ensuring that the appropriate
    position ids are passed to the forward function.
    """

    def __init__(self, num_embeddings: int, embedding_dim: int, padding_idx: int):
        if padding_idx is not None:
            num_embeddings_ = num_embeddings + padding_idx + 1
        else:
            num_embeddings_ = num_embeddings
        super().__init__(num_embeddings_, embedding_dim, padding_idx)
        self.max_positions = num_embeddings

    def forward(self, input: torch.Tensor):
        """Input is expected to be of size [bsz x seqlen]."""
        if input.size(1) > self.max_positions:
            raise ValueError(
                f"Sequence length {input.size(1)} above maximum "
                f" sequence length of {self.max_positions}"
            )
        mask = input.ne(self.padding_idx).int()
        positions = (torch.cumsum(mask, dim=1).type_as(mask) * mask).long() + self.padding_idx
        return F.embedding(
            positions,
            self.weight,
            self.padding_idx,
            self.max_norm,
            self.norm_type,
            self.scale_grad_by_freq,
            self.sparse,
        )
```
* For the `F.embedding`, follow the `reset_parameters` in the [`Embedding`](https://pytorch.org/docs/stable/_modules/torch/nn/modules/sparse.html#Embedding) class in `torch.nn.modules.sparse`
```
def reset_parameters(self) -> None:
    init.normal_(self.weight)
    self._fill_padding_idx_with_zero()
```
### Reinit `emb_layer_norm_before` and `emb_layer_norma_after`
```
self.emb_layer_norm_before = (
    ESM1bLayerNorm(self.args.embed_dim) if self.emb_layer_norm_before else None
)
self.emb_layer_norm_after = ESM1bLayerNorm(self.args.embed_dim)
```
### Reinit `lm_head`
* For [`RobertaLMHead`](https://github.com/facebookresearch/esm/blob/839c5b82c6cd9e18baa7a88dcbed3bd4b6d48e47/esm/modules.py#L298) class in [`esm.modules`](https://github.com/facebookresearch/esm/blob/main/esm/modules.py)
```
def __init__(self, embed_dim, output_dim, weight):
    super().__init__()
    self.dense = nn.Linear(embed_dim, embed_dim)
    self.layer_norm = ESM1bLayerNorm(embed_dim)
    self.weight = weight
    self.bias = nn.Parameter(torch.zeros(output_dim))
```
### Summary for reinit ESM1b
* To put together with proper initialization indicated in brackets, where the `bound` can be calcualted as
```
fan_in, _ = init._calculate_fan_in_and_fan_out(self.weight)
bound = 1 / math.sqrt(fan_in) if fan_in > 0 else 0
init.uniform_(self.bias, -bound, bound)
```
```
embed_tokens.weight: dim 2
layers.n.self_attn.k_proj.weight: dim 2         [nn.init.xavier_uniform_(self.k_proj.weight, gain=1 / math.sqrt(2))]
layers.n.self_attn.k_proj.bias: dim 1           [nn.init.uniform_(self.bias, -bound, bound)]
layers.n.self_attn.v_proj.weight: dim 2         [nn.init.xavier_uniform_(self.v_proj.weight, gain=1 / math.sqrt(2))]
layers.n.self_attn.v_proj.bias: dim 1           [nn.init.uniform_(self.bias, -bound, bound)]
layers.n.self_attn.q_proj.weight: dim 2         [nn.init.xavier_uniform_(self.q_proj.weight, gain=1 / math.sqrt(2))]
layers.n.self_attn.q_proj.bias: dim 1           [nn.init.uniform_(self.bias, -bound, bound)]
layers.n.self_attn.out_proj.weight: dim 2       [nn.init.xavier_uniform_(self.out_proj.weight)]
layers.n.self_attn.out_proj.bias: dim 1         [nn.init.constant_(self.out_proj.bias, 0.0)]
layers.n.self_attn_layer_norm.weight: dim 1     [nn.Parameter(torch.ones(hidden_size))]
layers.n.self_attn_layer_norm.bias: dim 1       [nn.Parameter(torch.zeros(hidden_size))]
layers.n.fc1.weight: dim 2                      [nn.init.kaiming_uniform_(self.weight, a=math.sqrt(5))]
layers.n.fc1.bias: dim 1                        [nn.init.uniform_(self.bias, -bound, bound)]
layers.n.fc2.weight: dim 2                      [nn.init.kaiming_uniform_(self.weight, a=math.sqrt(5))]
layers.n.fc2.bias: dim 1                        [nn.init.uniform_(self.bias, -bound, bound)]
layers.n.final_layer_norm.weight: dim 1         [nn.Parameter(torch.ones(hidden_size))]
layers.n.final_layer_norm.bias: dim 1           [nn.Parameter(torch.zeros(hidden_size))]
contact_head.regression.weight: dim 2           [nn.init.kaiming_uniform_(self.weight, a=math.sqrt(5))]
contact_head.regression.bias: dim 1             [nn.init.uniform_(self.bias, -bound, bound)]
embed_positions.weight: dim 2                   [nn.init.normal_(self.weight)]
emb_layer_norm_before.weight: dim 1             [nn.Parameter(torch.ones(hidden_size))]
emb_layer_norm_before.bias: dim 1               [nn.Parameter(torch.zeros(hidden_size))]
emb_layer_norm_after.weight: dim 1              [nn.Parameter(torch.ones(hidden_size))]
emb_layer_norm_after.bias: dim 1                [nn.Parameter(torch.zeros(hidden_size))]
-----------------------------------------------------------------------------------------------------------------------
[After the embeddings, no reinit needed]
lm_head.weight: dim 2                           [nn.init.xavier_uniform_()]
lm_head.bias: dim 1                             [nn.Parameter(torch.zeros(output_dim))]
lm_head.dense.weight: dim 2                     [nn.init.kaiming_uniform_(self.weight, a=math.sqrt(5))]
lm_head.dense.bias: dim 1                       [nn.init.uniform_(self.bias, -bound, bound)]
lm_head.layer_norm.weight: dim 1                [nn.Parameter(torch.ones(hidden_size))]
lm_head.layer_norm.bias: dim 1                  [nn.Parameter(torch.zeros(hidden_size))]
```

<a name="resamp_esm1b"></a>
## Resample (`self._resample_param = True`) ESM1b
* Shuffle the weights for the layers would have been reinit above
* n = 0, ..., 32

```
layers.n.self_attn.k_proj.weight: torch.Size([1280, 1280])
layers.n.self_attn.k_proj.bias: torch.Size([1280])
layers.n.self_attn.v_proj.weight: torch.Size([1280, 1280])
layers.n.self_attn.v_proj.bias: torch.Size([1280])
layers.n.self_attn.q_proj.weight: torch.Size([1280, 1280])
layers.n.self_attn.q_proj.bias: torch.Size([1280])
layers.n.self_attn.out_proj.weight: torch.Size([1280, 1280])
layers.n.self_attn.out_proj.bias: torch.Size([1280])
layers.n.self_attn_layer_norm.weight: torch.Size([1280])
layers.n.self_attn_layer_norm.bias: torch.Size([1280])
layers.n.fc1.weight: torch.Size([5120, 1280])
layers.n.fc1.bias: torch.Size([5120])
layers.n.fc2.weight: torch.Size([1280, 5120])
layers.n.fc2.bias: torch.Size([1280])
layers.n.final_layer_norm.weight: torch.Size([1280])
layers.n.final_layer_norm.bias: torch.Size([1280])
contact_head.regression.weight: torch.Size([1, 660])
contact_head.regression.bias: torch.Size([1])
embed_positions.weight: torch.Size([1026, 1280])
emb_layer_norm_before.weight: torch.Size([1280])
emb_layer_norm_before.bias: torch.Size([1280])
emb_layer_norm_after.weight: torch.Size([1280])
emb_layer_norm_after.bias: torch.Size([1280])
lm_head.weight: torch.Size([33, 1280])
lm_head.bias: torch.Size([33])
lm_head.dense.weight: torch.Size([1280, 1280])
lm_head.dense.bias: torch.Size([1280])
lm_head.layer_norm.weight: torch.Size([1280])
lm_head.layer_norm.bias: torch.Size([1280])
```
* Shuffle all weights along non-zero or embedding dimension

<a name="reinit_esm1"></a>
## Reinit (`self._reset_param = True`) ESM1
### ESM1 architecture
* `esm1_t6_43M_UR50S`
    * n = 0, ..., 5
    * embed_dim = 768
    * fc_dim = 3072
    * reg_dim = 72
* `esm1_t12_85M_UR50S`
    * n = 0, ..., 11
    * embed_dim = 768
    * fc_dim = 3072
    * reg_dim = 144
* `esm1_t34_670M_UR50S`
    * n = 0, ..., 33
    * embed_dim = 1280
    * fc_dim = 3072
    * reg_dim = 680
```
ProteinBertModel(
  (embed_tokens): Embedding(35, embed_dim, padding_idx=1)
  (layers): ModuleList(
    (n): TransformerLayer(
      (self_attn): MultiheadAttention(
        (k_proj): Linear(in_features=embed_dim, out_features=embed_dim, bias=True)
        (v_proj): Linear(in_features=embed_dim, out_features=embed_dim, bias=True)
        (q_proj): Linear(in_features=embed_dim, out_features=embed_dim, bias=True)
        (out_proj): Linear(in_features=embed_dim, out_features=embed_dim, bias=True)
      )
      (self_attn_layer_norm): ESM1LayerNorm()
      (fc1): Linear(in_features=embed_dim, out_features=fc_dim, bias=True)
      (fc2): Linear(in_features=fc_dim, out_features=embed_dim, bias=True)
      (final_layer_norm): ESM1LayerNorm()
    )
  )
  (contact_head): ContactPredictionHead(
    (regression): Linear(in_features=reg_dim, out_features=1, bias=True)
    (activation): Sigmoid()
  )
  (embed_positions): SinusoidalPositionalEmbedding()
)
```
### ESM1 architecture with dimensions
* Get the layers and dim with
```
for k, p in model.state_dict().items():
    print(f"{k}: dim {p.dim()}")
```
* Returns
```
embed_out: dim 2
embed_out_bias: dim 1
embed_tokens.weight: dim 2
layers.n.self_attn.bias_k: dim 3
layers.n.self_attn.bias_v: dim 3
layers.n.self_attn.k_proj.weight: dim 2
layers.n.self_attn.k_proj.bias: dim 1
layers.n.self_attn.v_proj.weight: dim 2
layers.n.self_attn.v_proj.bias: dim 1
layers.n.self_attn.q_proj.weight: dim 2
layers.n.self_attn.q_proj.bias: dim 1
layers.n.self_attn.out_proj.weight: dim 2
layers.n.self_attn.out_proj.bias: dim 1
layers.n.self_attn_layer_norm.weight: dim 1
layers.n.self_attn_layer_norm.bias: dim 1
layers.n.fc1.weight: dim 2
layers.n.fc1.bias: dim 1
layers.n.fc2.weight: dim 2
layers.n.fc2.bias: dim 1
layers.n.final_layer_norm.weight: dim 1
layers.n.final_layer_norm.bias: dim 1
contact_head.regression.weight: dim 2
contact_head.regression.bias: dim 1
embed_positions._float_tensor: dim 1
```

* To reinit, follow [`ProteinBertModel`](https://github.com/facebookresearch/esm/blob/839c5b82c6cd9e18baa7a88dcbed3bd4b6d48e47/esm/model/esm1.py#L22
) class in [`esm.model.esm1`](https://github.com/facebookresearch/esm/blob/main/esm/model/esm1.py)

```
 def __init__(self, args, alphabet):
    super().__init__()
    self.args = args
    self.alphabet_size = len(alphabet)
    self.padding_idx = alphabet.padding_idx
    self.mask_idx = alphabet.mask_idx
    self.cls_idx = alphabet.cls_idx
    self.eos_idx = alphabet.eos_idx
    self.prepend_bos = alphabet.prepend_bos
    self.append_eos = alphabet.append_eos
    self.emb_layer_norm_before = getattr(self.args, "emb_layer_norm_before", False)
    if self.args.arch == "roberta_large":
        self.model_version = "ESM-1b"
        self._init_submodules_esm1b()
    else:
        self.model_version = "ESM-1"
        self._init_submodules_esm1()

def _init_submodules_common(self):
    self.embed_tokens = nn.Embedding(
        self.alphabet_size, self.args.embed_dim, padding_idx=self.padding_idx
    )
    self.layers = nn.ModuleList(
        [
            TransformerLayer(
                self.args.embed_dim,
                self.args.ffn_embed_dim,
                self.args.attention_heads,
                add_bias_kv=(self.model_version != "ESM-1b"),
                use_esm1b_layer_norm=(self.model_version == "ESM-1b"),
            )
            for _ in range(self.args.layers)
        ]
    )

    self.contact_head = ContactPredictionHead(
        self.args.layers * self.args.attention_heads,
        self.prepend_bos,
        self.append_eos,
        eos_idx=self.eos_idx,
    )

def _init_submodules_esm1(self):
        self._init_submodules_common()
        self.embed_scale = math.sqrt(self.args.embed_dim)
        self.embed_positions = SinusoidalPositionalEmbedding(self.args.embed_dim, self.padding_idx)
        self.embed_out = nn.Parameter(torch.zeros((self.alphabet_size, self.args.embed_dim)))
        self.embed_out_bias = None
        if self.args.final_bias:
            self.embed_out_bias = nn.Parameter(torch.zeros(self.alphabet_size))
```

#### Reinit `layer`
* Where for [`TransformerLayer`](https://github.com/facebookresearch/esm/blob/839c5b82c6cd9e18baa7a88dcbed3bd4b6d48e47/esm/modules.py#L84) class in [`esm.modules`](https://github.com/facebookresearch/esm/blob/main/esm/modules.py),
    * `add_bias_kv=(self.model_version != "ESM-1b")` would be `True`
    * `use_esm1b_layer_norm=(self.model_version == "ESM-1b")` would be `False`
    * `BertLayerNorm` would be `ESM1LayerNorm`

```
def _init_submodules(self, add_bias_kv, use_esm1b_layer_norm):
    BertLayerNorm = ESM1bLayerNorm if use_esm1b_layer_norm else ESM1LayerNorm

    self.self_attn = MultiheadAttention(
        self.embed_dim,
        self.attention_heads,
        add_bias_kv=add_bias_kv,
        add_zero_attn=False,
        use_rotary_embeddings=self.use_rotary_embeddings,
    )
    self.self_attn_layer_norm = BertLayerNorm(self.embed_dim)

    self.fc1 = nn.Linear(self.embed_dim, self.ffn_embed_dim)
    self.fc2 = nn.Linear(self.ffn_embed_dim, self.embed_dim)

    self.final_layer_norm = BertLayerNorm(self.embed_dim)
```

* For [`MultiheadAttention`](https://github.com/facebookresearch/esm/blob/839c5b82c6cd9e18baa7a88dcbed3bd4b6d48e47/esm/multihead_attention.py#L68) class in [`esm.multihead_attention`](https://github.com/facebookresearch/esm/blob/main/esm/multihead_attention.py)

```
def __init__(
    self,
    embed_dim,
    num_heads,
    kdim=None,
    vdim=None,
    dropout=0.0,
    bias=True,
    add_bias_kv: bool = False,
    add_zero_attn: bool = False,
    self_attention: bool = False,
    encoder_decoder_attention: bool = False,
    use_rotary_embeddings: bool = False,
):
    super().__init__()
    self.embed_dim = embed_dim
    self.kdim = kdim if kdim is not None else embed_dim
    self.vdim = vdim if vdim is not None else embed_dim
    self.qkv_same_dim = self.kdim == embed_dim and self.vdim == embed_dim

    self.num_heads = num_heads
    self.dropout = dropout
    self.head_dim = embed_dim // num_heads
    assert (
        self.head_dim * num_heads == self.embed_dim
    ), "embed_dim must be divisible by num_heads"
    self.scaling = self.head_dim**-0.5

    self.self_attention = self_attention
    self.encoder_decoder_attention = encoder_decoder_attention

    assert not self.self_attention or self.qkv_same_dim, (
        "Self-attention requires query, key and " "value to be of the same size"
    )

    self.k_proj = nn.Linear(self.kdim, embed_dim, bias=bias)
    self.v_proj = nn.Linear(self.vdim, embed_dim, bias=bias)
    self.q_proj = nn.Linear(embed_dim, embed_dim, bias=bias)

    self.out_proj = nn.Linear(embed_dim, embed_dim, bias=bias)

    if add_bias_kv:
        self.bias_k = Parameter(torch.Tensor(1, 1, embed_dim))
        self.bias_v = Parameter(torch.Tensor(1, 1, embed_dim))
    else:
        self.bias_k = self.bias_v = None

    self.add_zero_attn = add_zero_attn

    self.reset_parameters()

    ...

def reset_parameters(self):
    if self.qkv_same_dim:
        # Empirically observed the convergence to be much better with
        # the scaled initialization
        nn.init.xavier_uniform_(self.k_proj.weight, gain=1 / math.sqrt(2))
        nn.init.xavier_uniform_(self.v_proj.weight, gain=1 / math.sqrt(2))
        nn.init.xavier_uniform_(self.q_proj.weight, gain=1 / math.sqrt(2))
    else:
        nn.init.xavier_uniform_(self.k_proj.weight)
        nn.init.xavier_uniform_(self.v_proj.weight)
        nn.init.xavier_uniform_(self.q_proj.weight)

    nn.init.xavier_uniform_(self.out_proj.weight)
    if self.out_proj.bias is not None:
        nn.init.constant_(self.out_proj.bias, 0.0)
    if self.bias_k is not None:
        nn.init.xavier_normal_(self.bias_k)
    if self.bias_v is not None:
        nn.init.xavier_normal_(self.bias_v)
```
* For the `bias` in `nn.Linear`, follow the `reset_parameters` in [`torch.nn.modules.linear`](https://pytorch.org/docs/stable/_modules/torch/nn/modules/linear.html#Linear)
```
def reset_parameters(self) -> None:
    # Setting a=sqrt(5) in kaiming_uniform is the same as initializing with
    # uniform(-1/sqrt(in_features), 1/sqrt(in_features)). For details, see
    # https://github.com/pytorch/pytorch/issues/57109
    init.kaiming_uniform_(self.weight, a=math.sqrt(5))
    if self.bias is not None:
        fan_in, _ = init._calculate_fan_in_and_fan_out(self.weight)
        bound = 1 / math.sqrt(fan_in) if fan_in > 0 else 0
        init.uniform_(self.bias, -bound, bound)
```
* For [`ESM1LayerNorm`](https://github.com/facebookresearch/esm/blob/839c5b82c6cd9e18baa7a88dcbed3bd4b6d48e47/esm/modules.py#L44) class in [`esm.modules`](https://github.com/facebookresearch/esm/blob/main/esm/modules.py)
```
    def __init__(self, hidden_size, eps=1e-12, affine=True):
        """Construct a layernorm layer in the TF style (eps inside the sqrt)."""
        super().__init__()
        self.hidden_size = (hidden_size,) if isinstance(hidden_size, int) else tuple(hidden_size)
        self.eps = eps
        self.affine = bool(affine)
        if self.affine:
            self.weight = nn.Parameter(torch.ones(hidden_size))
            self.bias = nn.Parameter(torch.zeros(hidden_size))
        else:
            self.weight, self.bias = None, None
```
* ESM1 `TransformerLayer` has
    * `MultiheadAttention`
        * `kdim = None`
        * `vdim = None`
        * `bias = True`
        * `add_bias_kv = True`
        * `add_zero_attn = False`
        * `self_attention = False`
    * `BertLayerNorm`
        * `affine = True`
* Thus
    * `MultiheadAttention`
        * `self.kdim = kdim if kdim is not None else embed_dim` would be `embed_dim`
        * `self.vdim = vdim if vdim is not None else embed_dim` would be `embed_dim`
        * `self.qkv_same_dim = self.kdim == embed_dim and self.vdim == embed_dim` would be `True`
        * `self.bias_k = Parameter(torch.Tensor(1, 1, embed_dim))` then `nn.init.xavier_normal_(self.bias_k)`
        * `self.bias_v = Parameter(torch.Tensor(1, 1, embed_dim))` then `nn.init.xavier_normal_(self.bias_k)`
    * `BertLayerNorm`
        * `BertLayerNorm = ESM1LayerNorm`
* More concretely
    * `MultiheadAttention`
        * `nn.init.xavier_uniform_(self.k_proj.weight, gain=1 / math.sqrt(2))`
        * `nn.init.xavier_uniform_(self.v_proj.weight, gain=1 / math.sqrt(2))`
        * `nn.init.xavier_uniform_(self.q_proj.weight, gain=1 / math.sqrt(2))`
        * `nn.init.xavier_uniform_(self.out_proj.weight)`
        * `nn.init.constant_(self.out_proj.bias, 0.0)`
        * `nn.init.xavier_normal_(self.bias_k)`
        * `nn.init.xavier_normal_(self.bias_v)`
    * `BertLayerNorm`
        * `self.weight = nn.Parameter(torch.ones(hidden_size))`
        * `self.bias = nn.Parameter(torch.zeros(hidden_size))`
#### Reinit `contact_head`
* For [`ContactPredictionHead`](https://github.com/facebookresearch/esm/blob/839c5b82c6cd9e18baa7a88dcbed3bd4b6d48e47/esm/modules.py#L317) class in [`esm.modules`](https://github.com/facebookresearch/esm/blob/main/esm/modules.py)
```
def __init__(
    self,
    in_features: int,
    prepend_bos: bool,
    append_eos: bool,
    bias=True,
    eos_idx: Optional[int] = None,
):
    super().__init__()
    self.in_features = in_features
    self.prepend_bos = prepend_bos
    self.append_eos = append_eos
    if append_eos and eos_idx is None:
        raise ValueError("Using an alphabet with eos token, but no eos token was passed in.")
    self.eos_idx = eos_idx
    self.regression = nn.Linear(in_features, 1, bias)
    self.activation = nn.Sigmoid()
```
* For ESM1 `ContactPredictionHead` has
    * `bias = True`

### Reinit `embed_positions`
* For [`SinusoidalPositionalEmbedding`](https://github.com/facebookresearch/esm/blob/839c5b82c6cd9e18baa7a88dcbed3bd4b6d48e47/esm/modules.py#L260) class in [`esm.modules`](https://github.com/facebookresearch/esm/blob/main/esm/modules.py)
```
class SinusoidalPositionalEmbedding(nn.Module):
    def __init__(self, embed_dim, padding_idx, learned=False):
        super().__init__()
        self.embed_dim = embed_dim
        self.padding_idx = padding_idx
        self.register_buffer("_float_tensor", torch.FloatTensor(1))
        self.weights = None

        def forward(self, x):
        bsz, seq_len = x.shape
        max_pos = self.padding_idx + 1 + seq_len
        if self.weights is None or max_pos > self.weights.size(0):
            self.weights = self.get_embedding(max_pos)
        self.weights = self.weights.type_as(self._float_tensor)

        positions = self.make_positions(x)
        return self.weights.index_select(0, positions.view(-1)).view(bsz, seq_len, -1).detach()

    def make_positions(self, x):
        mask = x.ne(self.padding_idx)
        range_buf = torch.arange(x.size(1), device=x.device).expand_as(x) + self.padding_idx + 1
        positions = range_buf.expand_as(x)
        return positions * mask.long() + self.padding_idx * (1 - mask.long())

    def get_embedding(self, num_embeddings):
        half_dim = self.embed_dim // 2
        emb = math.log(10000) / (half_dim - 1)
        emb = torch.exp(torch.arange(half_dim, dtype=torch.float) * -emb)
        emb = torch.arange(num_embeddings, dtype=torch.float).unsqueeze(1) * emb.unsqueeze(0)
        emb = torch.cat([torch.sin(emb), torch.cos(emb)], dim=1).view(num_embeddings, -1)
        if self.embed_dim % 2 == 1:
            # zero pad
            emb = torch.cat([emb, torch.zeros(num_embeddings, 1)], dim=1)
        if self.padding_idx is not None:
            emb[self.padding_idx, :] = 0
        return emb
```
* Leave untouched

### Summary for reinit ESM1
* To put together with proper initialization indicated in brackets, where the `bound` can be calcualted as
```
fan_in, _ = init._calculate_fan_in_and_fan_out(self.weight)
bound = 1 / math.sqrt(fan_in) if fan_in > 0 else 0
init.uniform_(self.bias, -bound, bound)
```
```
embed_out: dim 2
embed_out_bias: dim 1
embed_tokens.weight: dim 2
layers.n.self_attn.bias_k: dim 3                [nn.init.xavier_normal_(self.bias_k)]
layers.n.self_attn.bias_v: dim 3                [nn.init.xavier_normal_(self.bias_v)]
layers.n.self_attn.k_proj.weight: dim 2         [nn.init.xavier_uniform_(self.k_proj.weight, gain=1 / math.sqrt(2))]
layers.n.self_attn.k_proj.bias: dim 1           [nn.init.uniform_(self.bias, -bound, bound)]
layers.n.self_attn.v_proj.weight: dim 2         [nn.init.xavier_uniform_(self.v_proj.weight, gain=1 / math.sqrt(2))]
layers.n.self_attn.v_proj.bias: dim 1           [nn.init.uniform_(self.bias, -bound, bound)]
layers.n.self_attn.q_proj.weight: dim 2         [nn.init.xavier_uniform_(self.q_proj.weight, gain=1 / math.sqrt(2))]
layers.n.self_attn.q_proj.bias: dim 1           [nn.init.uniform_(self.bias, -bound, bound)]
layers.n.self_attn.out_proj.weight: dim 2       [nn.init.xavier_uniform_(self.out_proj.weight)]
layers.n.self_attn.out_proj.bias: dim 1         [nn.init.constant_(self.out_proj.bias, 0.0)]
layers.n.self_attn_layer_norm.weight: dim 1     [nn.Parameter(torch.ones(hidden_size))]
layers.n.self_attn_layer_norm.bias: dim 1       [nn.Parameter(torch.zeros(hidden_size))]
layers.n.fc1.weight: dim 2                      [nn.init.kaiming_uniform_(self.weight, a=math.sqrt(5))]
layers.n.fc1.bias: dim 1                        [nn.init.uniform_(self.bias, -bound, bound)]
layers.n.fc2.weight: dim 2                      [nn.init.kaiming_uniform_(self.weight, a=math.sqrt(5))]
layers.n.fc2.bias: dim 1                        [nn.init.uniform_(self.bias, -bound, bound)]
layers.n.final_layer_norm.weight: dim 1         [nn.Parameter(torch.ones(hidden_size))]
layers.n.final_layer_norm.bias: dim 1           [nn.Parameter(torch.zeros(hidden_size))]
contact_head.regression.weight: dim 2           [nn.init.kaiming_uniform_(self.weight, a=math.sqrt(5))]
contact_head.regression.bias: dim 1             [nn.init.uniform_(self.bias, -bound, bound)]
embed_positions._float_tensor: dim 1
```

<a name="resamp_esm1"></a>
## Resample (`self._resample_param = True`) ESM1
* Shuffle the weights for the layers would have been reinit above
* `esm1_t6_43M_UR50S`
    * n = 0, ..., 5
    * embed_dim = 768
    * fc_dim = 3072
    * reg_dim = 72
* `esm1_t12_85M_UR50S`
    * n = 0, ..., 11
    * embed_dim = 768
    * fc_dim = 3072
    * reg_dim = 144
* `esm1_t34_670M_UR50S`
    * n = 0, ..., 33
    * embed_dim = 1280
    * fc_dim = 3072
    * reg_dim = 680

```
layers.n.self_attn.bias_k: torch.Size([1, 1, embed_dim])
layers.n.self_attn.bias_v: torch.Size([1, 1, embed_dim])
layers.n.self_attn.k_proj.weight: torch.Size([embed_dim, embed_dim])
layers.n.self_attn.k_proj.bias: torch.Size([embed_dim])
layers.n.self_attn.v_proj.weight: torch.Size([embed_dim, embed_dim])
layers.n.self_attn.v_proj.bias: torch.Size([embed_dim])
layers.n.self_attn.q_proj.weight: torch.Size([embed_dim, embed_dim])
layers.n.self_attn.q_proj.bias: torch.Size([embed_dim])
layers.n.self_attn.out_proj.weight: torch.Size([embed_dim, embed_dim])
layers.n.self_attn.out_proj.bias: torch.Size([embed_dim])
layers.n.self_attn_layer_norm.weight: torch.Size([embed_dim])
layers.n.self_attn_layer_norm.bias: torch.Size([embed_dim])
layers.n.fc1.weight: torch.Size([fc_dim, embed_dim])
layers.n.fc1.bias: torch.Size([fc_dim])
layers.n.fc2.weight: torch.Size([embed_dim, fc_dim])
layers.n.fc2.bias: torch.Size([embed_dim])
layers.n.final_layer_norm.weight: torch.Size([embed_dim])
layers.n.final_layer_norm.bias: torch.Size([embed_dim])
contact_head.regression.weight: torch.Size([1, reg_dim])
contact_head.regression.bias: torch.Size([1])
```

* Shuffle all weights along non-zero or embedding dimension

<a name="reset_carp"></a>
## Reinit (`self._reset_param = True`) CARP

* For [`load_carp`](https://github.com/microsoft/protein-sequence-models/blob/885a24f9802aaf63bb6b554d43f912a269781514/sequence_models/pretrained.py#L16) function in [`sequence_models.pretrained`](https://github.com/microsoft/protein-sequence-models/blob/main/sequence_models/pretrained.py)

```
def load_carp(model_data):
    d_embedding = model_data['d_embed']
    d_model = model_data['d_model']
    n_layers = model_data['n_layers']
    kernel_size = model_data['kernel_size']
    activation = model_data['activation']
    slim = model_data['slim']
    r = model_data['r']
    bgc = 'bigcarp' in model_data['model']
    if not bgc:
        n_tokens = len(PROTEIN_ALPHABET)
        mask_idx = PROTEIN_ALPHABET.index(MASK)
        pad_idx = PROTEIN_ALPHABET.index(PAD)
        n_frozen = None
    else:
        n_tokens = model_data['tokens']['size']
        mask_idx = model_data['tokens']['specials'][MASK]
        pad_idx = model_data['tokens']['specials'][PAD]
        if 'frozen' in model_data['model']:
            n_frozen = 19450
        else:
            n_frozen = None
    model = ByteNetLM(n_tokens, d_embedding, d_model, n_layers, kernel_size, r, dropout=0.0,
                      activation=activation, causal=False, padding_idx=mask_idx,
                      final_ln=True, slim=slim, n_frozen_embs=n_frozen)
    sd = model_data['model_state_dict']
    model.load_state_dict(sd)
    model = CARP(model.eval(), pad_idx=pad_idx)
    return model
```

* For [`CARP`](https://github.com/microsoft/protein-sequence-models/blob/885a24f9802aaf63bb6b554d43f912a269781514/sequence_models/pretrained.py#L87) class in [`sequence_models.pretrained`](https://github.com/microsoft/protein-sequence-models/blob/main/sequence_models/pretrained.py)

```
class CARP(nn.Module):
    """Wrapper that takes care of input masking."""

    def __init__(self, model: ByteNetLM, pad_idx=PROTEIN_ALPHABET.index(PAD)):
        super().__init__()
        self.model = model
        self.pad_idx = pad_idx

    def forward(self, x, repr_layers=[-1], logits=False):
        padding_mask = (x != self.pad_idx)
        padding_mask = padding_mask.unsqueeze(-1)
        if len(repr_layers) == 1 and repr_layers[0] == -1:
            repr_layers = [len(self.model.embedder.layers)]
        result = {}
        if len(repr_layers) > 0:
            result['representations'] = {}
        x = self.model.embedder._embed(x)
        if 0 in repr_layers:
            result[0] = x
        i = 1
        for layer in self.model.embedder.layers:
            x = layer(x, input_mask=padding_mask)
            if i in repr_layers:
                result['representations'][i] = x
            i += 1
        if logits:
            result['logits'] = self.model.decoder(self.model.last_norm(x))
        return result
```

* For [`ByteNetLM`](https://github.com/microsoft/protein-sequence-models/blob/885a24f9802aaf63bb6b554d43f912a269781514/sequence_models/convolutional.py#L329) class in [`sequence_models.convolutional`](https://github.com/microsoft/protein-sequence-models/blob/main/sequence_models/convolutional.py)

```
class ByteNetLM(nn.Module):

    def __init__(self, n_tokens, d_embedding, d_model, n_layers, kernel_size, r, rank=None, n_frozen_embs=None,
                 padding_idx=None, causal=False, dropout=0.0, final_ln=False, slim=True, activation='relu',
                 tie_weights=False, down_embed=True):
        super().__init__()
        self.embedder = ByteNet(n_tokens, d_embedding, d_model, n_layers, kernel_size, r,
                                padding_idx=padding_idx, causal=causal, dropout=dropout, down_embed=down_embed,
                                slim=slim, activation=activation, rank=rank, n_frozen_embs=n_frozen_embs)
        if tie_weights:
            self.decoder = nn.Linear(d_model, n_tokens, bias=False)
            self.decoder.weight = self.embedder.embedder.weight
        else:
            self.decoder = PositionFeedForward(d_model, n_tokens)
        if final_ln:
            self.last_norm = nn.LayerNorm(d_model)
        else:
            self.last_norm = nn.Identity()

    def forward(self, x, input_mask=None):
        e = self.embedder(x, input_mask=input_mask)
        e = self.last_norm(e)
        return self.decoder(e)
```

* For [`ByteNet`](https://github.com/microsoft/protein-sequence-models/blob/885a24f9802aaf63bb6b554d43f912a269781514/sequence_models/convolutional.py#L252) class in [`sequence_models.convolutional`](https://github.com/microsoft/protein-sequence-models/blob/main/sequence_models/convolutional.py)

```
class ByteNet(nn.Module):

    """Stacked residual blocks from ByteNet paper defined by n_layers
         
         Shape:
            Input: (N, L,)
            input_mask: (N, L, 1), optional
            Output: (N, L, d)
    """

    def __init__(self, n_tokens, d_embedding, d_model, n_layers, kernel_size, r, rank=None, n_frozen_embs=None,
                 padding_idx=None, causal=False, dropout=0.0, slim=True, activation='relu', down_embed=True):
        """
        :param n_tokens: number of tokens in token dictionary
        :param d_embedding: dimension of embedding
        :param d_model: dimension to use within ByteNet model, //2 every layer
        :param n_layers: number of layers of ByteNet block
        :param kernel_size: the kernel width
        :param r: used to calculate dilation factor
        :padding_idx: location of padding token in ordered alphabet
        :param causal: if True, chooses MaskedCausalConv1d() over MaskedConv1d()
        :param rank: rank of compressed weight matrices
        :param n_frozen_embs: number of frozen embeddings
        :param slim: if True, use half as many dimensions in the NLP as in the CNN
        :param activation: 'relu' or 'gelu'
        :param down_embed: if True, have lower dimension for initial embedding than in CNN layers
        """
        super().__init__()
        if n_tokens is not None:
            if n_frozen_embs is None:
                self.embedder = nn.Embedding(n_tokens, d_embedding, padding_idx=padding_idx)
            else:
                self.embedder = DoubleEmbedding(n_tokens - n_frozen_embs, n_frozen_embs,
                                                d_embedding, padding_idx=padding_idx)
        else:
            self.embedder = nn.Identity()
        if down_embed:
            self.up_embedder = PositionFeedForward(d_embedding, d_model)
        else:
            self.up_embedder = nn.Identity()
            assert n_tokens == d_embedding
        log2 = int(np.log2(r)) + 1
        dilations = [2 ** (n % log2) for n in range(n_layers)]
        d_h = d_model
        if slim:
            d_h = d_h // 2
        layers = [
            ByteNetBlock(d_model, d_h, d_model, kernel_size, dilation=d, causal=causal, rank=rank,
                         activation=activation)
            for d in dilations
        ]
        self.layers = nn.ModuleList(modules=layers)
        self.dropout = dropout

    def forward(self, x, input_mask=None):
        """
        :param x: (batch, length)
        :param input_mask: (batch, length, 1)
        :return: (batch, length,)
        """
        e = self._embed(x)
        return self._convolve(e, input_mask=input_mask)

    def _embed(self, x):
        e = self.embedder(x)
        e = self.up_embedder(e)
        return e

    def _convolve(self, e, input_mask=None):
        for layer in self.layers:
            e = layer(e, input_mask=input_mask)
            if self.dropout > 0.0:
                e = F.dropout(e, self.dropout)
        return e
```

* For [`ByteNetBlock`](https://github.com/microsoft/protein-sequence-models/blob/885a24f9802aaf63bb6b554d43f912a269781514/sequence_models/convolutional.py#L206) class in [`sequence_models.convolutional`](https://github.com/microsoft/protein-sequence-models/blob/main/sequence_models/convolutional.py)

```
class ByteNetBlock(nn.Module):
    """Residual block from ByteNet paper (https://arxiv.org/abs/1610.10099).
         
         Shape:
            Input: (N, L, d_in)
            input_mask: (N, L, 1), optional
            Output: (N, L, d_out)
    """

    def __init__(self, d_in, d_h, d_out, kernel_size, dilation=1, groups=1, causal=False, activation='relu', rank=None):
        super().__init__()
        if causal:
            self.conv = MaskedCausalConv1d(d_h, d_h, kernel_size=kernel_size, dilation=dilation, groups=groups)
        else:
            self.conv = MaskedConv1d(d_h, d_h, kernel_size=kernel_size, dilation=dilation, groups=groups)
        if activation == 'relu':
            act = nn.ReLU
        elif activation == 'gelu':
            act = nn.GELU
        layers1 = [
            nn.LayerNorm(d_in),
            act(),
            PositionFeedForward(d_in, d_h, rank=rank),
            nn.LayerNorm(d_h),
            act()
        ]
        layers2 = [
            nn.LayerNorm(d_h),
            act(),
            PositionFeedForward(d_h, d_out, rank=rank),
        ]
        self.sequence1 = nn.Sequential(*layers1)
        self.sequence2 = nn.Sequential(*layers2)

    def forward(self, x, input_mask=None):
        """
        :param x: (batch, length, in_channels)
        :param input_mask: (batch, length, 1)
        :return: (batch, length, out_channels)
        """
        return x + self.sequence2(
            self.conv(self.sequence1(x), input_mask=input_mask)
        )
```

* For [`PositionFeedForward`](https://github.com/microsoft/protein-sequence-models/blob/885a24f9802aaf63bb6b554d43f912a269781514/sequence_models/layers.py#L63) class in [`sequence_models.layers`](https://github.com/microsoft/protein-sequence-models/blob/main/sequence_models/layers.py)

```
class PositionFeedForward(nn.Module):

    def __init__(self, d_in, d_out, rank=None):
        super().__init__()
        if rank is None:
            self.conv = nn.Conv1d(d_in, d_out, 1)
            self.factorized = False
        else:
            layer = nn.Linear(d_in, d_out)
            w = layer.weight.data
            self.bias = layer.bias
            u, s, v = torch.svd(w)
            s = torch.diag(s[:rank].sqrt())
            u = u[:, :rank]
            v = v.t()[:rank]
            self.u = nn.Parameter(u @ s)
            self.v = nn.Parameter(s @ v)
            self.factorized = True

    def forward(self, x):
        if self.factorized:
            w = self.u @ self.v
            return x @ w.t() + self.bias
        else:
            return self.conv(x.transpose(1, 2)).transpose(1, 2)
```

* For [`LayerNorm`](https://pytorch.org/docs/stable/_modules/torch/nn/modules/normalization.html#LayerNorm)

```
def reset_parameters(self) -> None:
    if self.elementwise_affine:
        init.ones_(self.weight)
        init.zeros_(self.bias)
```

* For [`MaskedConv1d`](https://github.com/microsoft/protein-sequence-models/blob/885a24f9802aaf63bb6b554d43f912a269781514/sequence_models/convolutional.py#L10) and [`MaskedCausalConv1d`](https://github.com/microsoft/protein-sequence-models/blob/885a24f9802aaf63bb6b554d43f912a269781514/sequence_models/convolutional.py#L76) both are based on [`nn.Conv1d`](https://pytorch.org/docs/stable/_modules/torch/nn/modules/conv.html#Conv1d)

```
class MaskedCausalConv1d(nn.Module):
    """Masked Causal 1D convolution based on https://github.com/Popgun-Labs/PopGen/. 
         
         Shape:
            Input: (N, L, in_channels)
            input_mask: (N, L, 1), optional
            Output: (N, L, out_channels)
    """

    def __init__(self, in_channels, out_channels, kernel_size=1, dilation=1, groups=1, init=None):
        """
        Causal 1d convolutions with caching mechanism for O(L) generation,
        as described in the ByteNet paper (Kalchbrenner et al, 2016) and "Fast Wavenet" (Paine, 2016)
        Usage:
            At train time, API is same as regular convolution. `conv = CausalConv1d(...)`
            At inference time, set `conv.sequential = True` to enable activation caching, and feed
            sequence through step by step. Recurrent state is managed internally.
        References:
            - Neural Machine Translation in Linear Time: https://arxiv.org/abs/1610.10099
            - Fast Wavenet: https://arxiv.org/abs/1611.09482
        :param in_channels: input channels
        :param out_channels: output channels
        :param kernel_size: the kernel width
        :param dilation: dilation factor
        :param groups: perform depth-wise convolutions
        :param init: optional initialisation function for nn.Conv1d module (e.g xavier)
        """
        super().__init__()

        self.in_channels = in_channels
        self.out_channels = out_channels
        self.kernel_size = kernel_size
        self.dilation = dilation
        self.groups = groups

        # if `true` enables fast generation
        self.sequential = False

        # compute required amount of padding to preserve the length
        self.zeros = (kernel_size - 1) * dilation
        self.conv = nn.Conv1d(in_channels, out_channels, kernel_size, dilation=dilation, groups=groups)

        # use supplied initialization function
        if init:
            init(self.conv)

    def forward(self, x, input_mask=None):
        """
        :param x: (batch, length, in_channels)
        :param input_mask: (batch, length, 1)
        :return: (batch, length, out_channels)
        """
        if input_mask is not None:
            x = x * input_mask
        # training mode
        x = torch.transpose(x, 1, 2)
        if not self.sequential:
            # no padding for kw=1
            if self.kernel_size == 1:
                return self.conv(x).transpose(1, 2)

            # left-pad + conv.
            out = self._pad(x)
            return self._unpad(self.conv(out)).transpose(1, 2)

        # sampling mode
        else:
            # note: x refers to a single timestep (batch, features, 1)
            if not hasattr(self, 'recurrent_state'):
                batch_size = x.size(0)
                self._init_recurrent_state(batch_size)

            return self._generate(x).transpose(1, 2)

    def _pad(self, x):
        return F.pad(x, [self.zeros, 0])

    def _unpad(self, x):
        return x

    def clear_cache(self):
        """
        Delete the recurrent state. Note: this should be called between runs, to prevent
        leftover state bleeding into future samples. Note that we delete state (instead of zeroing) to support
        changes in the inference time batch size.
        """
        if hasattr(self, 'recurrent_state'):
            del self.recurrent_state

    def _init_recurrent_state(self, batch):
        """
        Initialize the recurrent state for fast generation.
        :param batch: the batch size to generate
        """

        # extract weights and biases from nn.Conv1d module
        state = self.conv.state_dict()
        self.weight = state['weight']
        self.bias = state['bias']

        # initialize the recurrent states to zeros
        self.recurrent_state = torch.zeros(batch, self.in_channels, self.zeros, device=self.bias.device)

    def _generate(self, x_i):
        """
        Generate a single output activations, from the input activation
        and the cached recurrent state activations from previous steps.
        :param x_i: features of a single timestep (batch, in_channels, 1)
        :return: the next output value in the series (batch, out_channels, 1)
        """

        # if the kernel_size is greater than 1, use recurrent state.
        if self.kernel_size > 1:
            # extract the recurrent state and concat with input column
            recurrent_activations = self.recurrent_state[:, :, :self.zeros]
            f = torch.cat([recurrent_activations, x_i], 2)

            # update the cache for this layer
            self.recurrent_state = torch.cat(
                [self.recurrent_state[:, :, 1:], x_i], 2)
        else:
            f = x_i

        # perform convolution
        activations = F.conv1d(f, self.weight, self.bias,
                               dilation=self.dilation, groups=self.groups)

        return activations
```

```
class MaskedConv1d(nn.Conv1d):
    """ A masked 1-dimensional convolution layer.
    Takes the same arguments as torch.nn.Conv1D, except that the padding is set automatically.
         Shape:
            Input: (N, L, in_channels)
            input_mask: (N, L, 1), optional
            Output: (N, L, out_channels)
    """

    def __init__(self, in_channels: int, out_channels: int,
                 kernel_size: int, stride: int=1, dilation: int=1, groups: int=1,
                 bias: bool=True):
        """
        :param in_channels: input channels
        :param out_channels: output channels
        :param kernel_size: the kernel width
        :param stride: filter shift
        :param dilation: dilation factor
        :param groups: perform depth-wise convolutions
        :param bias: adds learnable bias to output
        """
        padding = dilation * (kernel_size - 1) // 2
        super().__init__(in_channels, out_channels, kernel_size, stride=stride, dilation=dilation,
                                           groups=groups, bias=bias, padding=padding)

    def forward(self, x, input_mask=None):
        if input_mask is not None:
            x = x * input_mask
        return super().forward(x.transpose(1, 2)).transpose(1, 2)
```

```
def reset_parameters(self) -> None:
    # Setting a=sqrt(5) in kaiming_uniform is the same as initializing with
    # uniform(-1/sqrt(k), 1/sqrt(k)), where k = weight.size(1) * prod(*kernel_size)
    # For more details see: https://github.com/pytorch/pytorch/issues/15314#issuecomment-477448573
    init.kaiming_uniform_(self.weight, a=math.sqrt(5))
    if self.bias is not None:
        fan_in, _ = init._calculate_fan_in_and_fan_out(self.weight)
        if fan_in != 0:
            bound = 1 / math.sqrt(fan_in)
            init.uniform_(self.bias, -bound, bound)
```
* Specifically,
* `carp_600k`
    * n = 0, ..., 15
    * d_model = 128
    * hid_dim = 64
* `carp_38M`
    * n = 0, ..., 15
    * d_model = 1024
    * hid_dim = 512
* `carp_76M`
    * n = 0, ..., 31
    * d_model = 1024
    * hid_dim = 512
* `carp_640M`
    * n = 0, ..., 31
    * d_model = 1280
    * hid_dim = 1280

```
model.embedder.embedder.weight: torch.Size([30, 8])
model.embedder.up_embedder.conv.weight: torch.Size([d_model, 8, 1])
model.embedder.up_embedder.conv.bias: torch.Size([d_model])
model.embedder.layers.n.conv.weight: torch.Size([hid_dim, hid_dim, 5])
model.embedder.layers.n.conv.bias: torch.Size([hid_dim])
model.embedder.layers.n.sequence1.0.weight: torch.Size([d_model])
model.embedder.layers.n.sequence1.0.bias: torch.Size([d_model])
model.embedder.layers.n.sequence1.2.conv.weight: torch.Size([hid_dim, d_model, 1])
model.embedder.layers.n.sequence1.2.conv.bias: torch.Size([hid_dim])
model.embedder.layers.n.sequence1.3.weight: torch.Size([hid_dim])
model.embedder.layers.n.sequence1.3.bias: torch.Size([hid_dim])
model.embedder.layers.n.sequence2.0.weight: torch.Size([hid_dim])
model.embedder.layers.n.sequence2.0.bias: torch.Size([hid_dim])
model.embedder.layers.n.sequence2.2.conv.weight: torch.Size([d_model, hid_dim, 1])
model.embedder.layers.n.sequence2.2.conv.bias: torch.Size([d_model])
model.decoder.conv.weight: torch.Size([30, d_model, 1])
model.decoder.conv.bias: torch.Size([30])
model.last_norm.weight: torch.Size([d_model])
model.last_norm.bias: torch.Size([d_model])
```

```
model.embedder.embedder.weight
model.embedder.up_embedder.conv.weight
model.embedder.up_embedder.conv.bias
model.embedder.layers.n.conv.weight                 [init.kaiming_uniform_(self.weight, a=math.sqrt(5))]
model.embedder.layers.n.conv.bias                   [init.uniform_(self.bias, -bound, bound)]
model.embedder.layers.n.sequence1.0.weight          [init.ones_(self.weight)]
model.embedder.layers.n.sequence1.0.bias            [init.zeros_(self.bias)]
model.embedder.layers.n.sequence1.2.conv.weight     [init.kaiming_uniform_(self.weight, a=math.sqrt(5))]
model.embedder.layers.n.sequence1.2.conv.bias       [init.uniform_(self.bias, -bound, bound)]
model.embedder.layers.n.sequence1.3.weight          [init.ones_(self.weight)]
model.embedder.layers.n.sequence1.3.bias            [init.zeros_(self.bias)]
model.embedder.layers.n.sequence2.0.weight          [init.ones_(self.weight)]
model.embedder.layers.n.sequence2.0.bias            [init.zeros_(self.bias)]
model.embedder.layers.n.sequence2.2.conv.weight     [init.kaiming_uniform_(self.weight, a=math.sqrt(5))]
model.embedder.layers.n.sequence2.2.conv.bias       [init.uniform_(self.bias, -bound, bound)]
model.decoder.conv.weight
model.decoder.conv.bias
model.last_norm.weight
model.last_norm.bias
```


<a name="resamp_carp"></a>
## Resample (`self._resample_param = True`) CARP
* Shuffle along all dim

```
model.embedder.layers.n.conv.weight: torch.Size([hid_dim, hid_dim, 5])
model.embedder.layers.n.conv.bias: torch.Size([hid_dim])
model.embedder.layers.n.sequence1.0.weight: torch.Size([d_model])
model.embedder.layers.n.sequence1.0.bias: torch.Size([d_model])
model.embedder.layers.n.sequence1.2.conv.weight: torch.Size([hid_dim, d_model, 1])
model.embedder.layers.n.sequence1.2.conv.bias: torch.Size([hid_dim])
model.embedder.layers.n.sequence1.3.weight: torch.Size([hid_dim])
model.embedder.layers.n.sequence1.3.bias: torch.Size([hid_dim])
model.embedder.layers.n.sequence2.0.weight: torch.Size([hid_dim])
model.embedder.layers.n.sequence2.0.bias: torch.Size([hid_dim])
model.embedder.layers.n.sequence2.2.conv.weight: torch.Size([d_model, hid_dim, 1])
model.embedder.layers.n.sequence2.2.conv.bias: torch.Size([d_model])
```
