## Supplemental data and code for "Feature Reuse and Scaling: Understanding Transfer Learning with Protein Language Models": pytorch_model.py

```
"""PyTorch models"""

from __future__ import annotations

import torch
from torch import nn


class LinearRegression(nn.Module):
    """Linear regression"""

    def __init__(self, input_dim: int, output_dim: int):
        super(LinearRegression, self).__init__()
        self.linear = nn.Linear(input_dim, output_dim)

    def forward(self, x):
        return self.linear(x)

    @property
    def model_name(self) -> str:
        """Return the name of the model"""
        return "LinearRegression"


class LinearClassifier(nn.Module):
    """Linear classifier"""

    def __init__(self, input_dim: int, numb_class: int, classifier_type: str):
        """
        Args:
        - input_dim: int,
        - numb_class: int, the number of classes
        - classifier_type: str, structure or annotation
        """
        super(LinearClassifier, self).__init__()
        self._classifier_type = classifier_type
        self.linear = nn.Linear(input_dim, numb_class)

    def forward(self, x: torch.tensor) -> torch.tensor:
        return self.linear(x)

    @property
    def model_name(self) -> str:
        """Return the name of the model"""
        if self._classifier_type.lower() == "structure":
            return "LinearClassifier-Structure"
        elif self._classifier_type.lower() == "annotation":
            return "LinearClassifier-Annotation"


class MultiLabelMultiClass(nn.Module):
    """Multi label multi class"""

    def __init__(self, input_dim: int, numb_class: int) -> None:
        super(MultiLabelMultiClass, self).__init__()
        self.linear = nn.Linear(input_dim, numb_class)

    def forward(self, x: torch.tensor) -> torch.tensor:
        return self.linear(x)

    @property
    def model_name(self) -> str:
        """Return the name of the model"""
        return "MultiLabelMultiClass"
```
