## Supplemental data and code for "Feature Reuse and Scaling: Understanding Transfer Learning with Protein Language Models": receptive_field.py

```
"""
A script for receptive field calc, following
https://distill.pub/2019/computing-receptive-fields/
"""

from __future__ import annotations

from collections import defaultdict

import os
import pandas as pd

from sequence_models.pretrained import load_model_and_alphabet

import matplotlib.pyplot as plt

from scr.params.emb import CARP_INFO
from scr.utils import checkNgen_folder


class ReceptiveField:

    """
    Calculate receptive field
    """

    def __init__(self, encoder_name: str):

        self._encoder_name = encoder_name

        self._model, _ = load_model_and_alphabet(self._encoder_name)

        # total number of layers
        self._layer_numb = CARP_INFO[self._encoder_name][1]

        self._conv_stat_dict = defaultdict(dict)

        for layer_name, _ in self._model.model.embedder.state_dict().items():
            # take out bias and weight in the name
            conv_layer_name = layer_name.replace(".weight", "").replace(".bias", "")

            if "conv" in conv_layer_name:

                conv_layer = self._model.model.embedder

                for sub_obj in conv_layer_name.split("."):
                    conv_layer = getattr(conv_layer, sub_obj)

                kernel_size = getattr(conv_layer, "kernel_size")
                stride = getattr(conv_layer, "stride")
                dilation = getattr(conv_layer, "dilation")

                for conv_param in [kernel_size, stride, dilation]:
                    assert len(conv_param) == 1, f"{conv_param=} not 1D"

                common_dict = {
                    "kernel_size": kernel_size[0],
                    "stride": stride[0],
                    "dilation": dilation[0],
                    "adjusted_kernel_size": self._adjust_kl(
                        kl=kernel_size[0], dl=dilation[0]
                    ),
                }

                if "up_embedder" in conv_layer_name:
                    self._conv_stat_dict["up_embedder"][conv_layer_name] = common_dict
                elif "layers" in conv_layer_name:
                    sequence_numb = ""
                    if "sequence" in conv_layer_name:
                        sequence_numb = conv_layer_name.split(".")[2]

                    self._conv_stat_dict["layers"][conv_layer_name] = {
                        **common_dict,
                        "layer_numb": conv_layer_name.split(".")[1],
                        "sequence_numb": sequence_numb,
                    }

    def _adjust_kl(self, kl: int, dl: int) -> int:

        """
        adjust the kernel_size for layer l with dilation of layer l

        dl * (kl - 1) + 1

        Args:
        - kl: int, kernel size of layer l
        - dl: int, dilation factor of layer l
        """

        if kl > 0:
            return dl * (kl - 1) + 1
        else:
            return kl

    def _get_rl(self, rl_prev: int, sl: int, kl: int, dl: int) -> int:

        """
        Calculate rl following equation (1) in
        https://distill.pub/2019/computing-receptive-fields/

        Given rl_prev = sl * rl + kl - sl
        rl = (rl_prev + sl - kl)/sl

        Args:
        - rl_prev: int, receptive field of l-1
        - sl: int, stride of layer l
        - kl: int, kernel size of layer l
        """

        rl = (rl_prev + sl - self._adjust_kl(kl=kl, dl=dl)) / sl

        assert rl.is_integer(), f"rl = {rl} should be int"

        return int(rl)

    def _get_r0(self, L: int) -> int:

        """
        Calculate rl for the first layer following equation (2) in
        https://distill.pub/2019/computing-receptive-fields/

        Note that all CARP stride = 1
        Each layer, a NyteNetBlock, is composed of:
            - conv, MaskedConv1d, kernel_size = 5
            - sequence1, with a PositionFeedForward conv layer, kernel_size = 1
            - sequence2, with a PositionFeedForward conv layer, kernel_size = 1

        Thus, the product of all sl will be 1
        kl - 1 will all be 4

        Here, we chop off the arch and ask for r0

        Args:
        - L: int, layer number for the chop off arch
        """

        r0 = 1

        for l in range(L):
            r0 += (
                self.conv_layers_df.loc[f"layers.{str(l)}.conv", "adjusted_kernel_size"]
                * self.conv_unique_stride
            )

        return r0

    @property
    def conv_stat_dict(self) -> dict:
        """Get the dict with all layer kernel and stride info"""
        return self._conv_stat_dict

    @property
    def conv_layers_df(self) -> pd.DataFrame:
        """Get the dict with all layer kernel and stride in df"""
        return pd.DataFrame.from_dict(self.conv_stat_dict["layers"]).T

    @property
    def conv_unique_stride(self) -> int:

        """Get the unique stride in df, should be all 1s for CARP"""

        conv_unique_strides = self.conv_layers_df.stride.unique()

        if self._encoder_name in CARP_INFO.keys():
            assert len(conv_unique_strides) == 1, "CARP stride should only be 1"
            assert conv_unique_strides[0] == 1, "CARP stride should only be 1"

        return conv_unique_strides[0]

    @property
    def conv_unique_kernelsize(self) -> int:

        """Get the unique kernel in df, should be 1 and 5 for CARP"""

        conv_unique_kernelsize = self.conv_layers_df.kernel_size.unique()

        if self._encoder_name in CARP_INFO.keys():
            assert (
                len(conv_unique_kernelsize) == 2
            ), "CARP stride should only have 2 kernel sizes"

        return conv_unique_kernelsize

    @property
    def conv_unique_nonone_kernelsize(self) -> int:

        """Get the unique kernel in df that is not 1"""
        conv_unique_nonone_kernelsize = self.conv_unique_kernelsize[
            self.conv_unique_kernelsize != 1
        ]

        if self._encoder_name in CARP_INFO.keys():
            assert (
                len(conv_unique_nonone_kernelsize) == 1
            ), "CARP should only have one not 1 kernel sizes"

        return conv_unique_nonone_kernelsize[0]

    @property
    def rf_dict(self) -> dict:
        """
        A dict with rf for each ByteNetBlock layer

        Note that all CARP stride = 1
        Each layer, a NyteNetBlock, is composed of:
            - conv, MaskedConv1d, kernel_size = 5
            - sequence1, with a PositionFeedForward conv layer, kernel_size = 1
            - sequence2, with a PositionFeedForward conv layer, kernel_size = 1
            - the sequence1 and sequence2 does not impact the rf of each layer
        """

        return {layer: self._get_r0(L=layer) for layer in range(self._layer_numb + 1)}

    @property
    def rf_df(self) -> pd.DataFrame:
        """Convert layer rf dict to dataframe with column names"""

        df = pd.Series(self.rf_dict).to_frame(name="rf_size")
        df.index.name = "layers"

        return df.reset_index()


def run_carp_rf(rf_folder: str = "results/rf"):

    """Save carp rf calc output as csv and plot"""

    rf_df_folder = checkNgen_folder(os.path.join(rf_folder, "dfs"))
    rf_plot_folder = checkNgen_folder(os.path.join(rf_folder, "plots"))
    encoder_names = list(CARP_INFO.keys())

    # init fig
    fig, axs = plt.subplots(
        1,
        len(encoder_names),
        sharey=True,
        figsize=(10, 2.5),
        squeeze=False,  # not get rid off the extra dim if 1D
    )

    for i, carp in enumerate(encoder_names):

        print(f"Calculating and plotting rf for {carp}...")

        rf_df = ReceptiveField(carp).rf_df

        # save df
        rf_df.to_csv(os.path.join(rf_df_folder, carp + ".csv"), index=False)

        # plot individual
        plt.figure()
        plt.plot("layers", "rf_size", data=rf_df)
        plt.xlabel("layers")
        plt.ylabel("rf_size")
        plt.title(carp)

        plt.savefig(
            os.path.join(rf_plot_folder, carp + ".png"),
            bbox_inches="tight",
        )

        plt.close()

        # add to collage
        axs[0, i].plot("layers", "rf_size", data=rf_df)

    # add xlabels
    for ax in axs.flatten():
        ax.set_xlabel("layers", fontsize=12)
        ax.tick_params(axis="x", labelsize=12)

    # add column names
    for ax, col in zip(axs[0], encoder_names):
        ax.set_title(col, fontsize=12)

    axs[0, 0].set_ylabel("rf_size", fontsize=12)

    # add whole plot level title
    fig.suptitle(
        "carp receptive field size",
        y=0.925,
        fontsize=12,
        fontweight="bold",
    )
    fig.align_labels()
    fig.tight_layout()

    plt.savefig(
        os.path.join(rf_plot_folder, "carp_all" + ".png"),
        bbox_inches="tight",
    )

    plt.close()
```
