## Supplemental data and code for "Feature Reuse and Scaling: Understanding Transfer Learning with Protein Language Models": run_pytorch.py

```
"""Script for running pytorch models"""

from __future__ import annotations

import os
import random
from tqdm import tqdm
from concurrent import futures

import numpy as np

import torch
import torch.nn as nn

from sklearn.metrics import ndcg_score, accuracy_score, roc_auc_score

from scipy.stats import spearmanr

from scr.params.aa import AA_NUMB
from scr.params.sys import RAND_SEED, DEVICE
from scr.params.emb import TRANSFORMER_INFO, CARP_INFO, MAX_SEQ_LEN

from scr.preprocess.data_process import split_protrain_loader, DatasetInfo
from scr.encoding.encoding_classes import (
    OnehotEncoder,
    ESMEncoder,
    CARPEncoder,
    get_emb_info,
)
from scr.model.pytorch_model import (
    LinearRegression,
    LinearClassifier,
    MultiLabelMultiClass,
)

from scr.model.train_test import train, test
from scr.vis.learning_vis import plot_lc
from scr.utils import get_folder_file_names, pickle_save, get_default_output_path

# seed everything
random.seed(RAND_SEED)
np.random.seed(RAND_SEED)
torch.manual_seed(RAND_SEED)
torch.cuda.manual_seed(RAND_SEED)
torch.cuda.manual_seed_all(RAND_SEED)
torch.backends.cudnn.deterministic = True


class Run_Pytorch:
    def __init__(
        self,
        dataset_path: str,
        encoder_name: str,
        checkpoint: float = 1,
        checkpoint_folder: str = "pretrain_checkpoints/carp",
        reset_param: bool = False,
        resample_param: bool = False,
        embed_batch_size: int = 128,
        flatten_emb: bool | str = False,
        embed_folder: str | None = None,
        seq_start_idx: bool | int = False,
        seq_end_idx: bool | int = False,
        manual_layer_min: bool | int = False,
        manual_layer_max: bool | int = False,
        loader_batch_size: int = 64,
        worker_seed: int = RAND_SEED,
        if_encode_all: bool = True,
        if_multiprocess: bool = False,
        if_rerun_layer: bool = False,
        learning_rate: float = 1e-4,
        lr_decay: float = 0.1,
        epochs: int = 100,
        early_stop: bool = True,
        tolerance: int = 10,
        min_epoch: int = 5,
        device: torch.device | str = DEVICE,
        all_plot_folder: str = "results/learning_curves",
        all_result_folder: str = "results/train_val_test",
        **encoder_params,
    ) -> None:

        """
        A function for running pytorch model

        Args:
        - dataset_path: str, full path to the dataset, in pkl or panda readable format
            columns include: sequence, target, set, validation, mut_name (optional), mut_numb (optional)
        - encoder_name: str, the name of the encoder
        - checkpoint: float = 1, the 0.5, 0.25, 0.125 checkpoint of the CARP encoder or full
        - checkpoint_folder: str = "pretrain_checkpoints/carp", folder for carp encoders
        - embed_batch_size: int, set to 0 to encode all in a single batch
        - flatten_emb: bool or str, if and how (one of ["max", "mean"]) to flatten the embedding
        - embed_folder: str = None, path to presaved embedding
        - seq_start_idx: bool | int = False, the index for the start of the sequence
        - seq_end_idx: bool | int = False, the index for the end of the sequence
        - manual_layer_min: bool | int = False
        - manual_layer_max: bool | int = False
        - loader_batch_size: int, the batch size for train, val, and test dataloader
        - worker_seed: int, the seed for dataloader
        - if_rerun_layer: bool, if rerun the layer if the results exist
        - learning_rate: float
        - lr_decay: float, factor by which to decay LR on plateau
        - epochs: int, number of epochs to train for
        - device: torch.device or str
        - all_plot_folder: str, the parent folder path for saving all the learning curves
        - all_result_folder: str = "results/train_val_test", the parent folder for all results
        - encoder_params: kwarg, additional parameters for encoding

        Returns:
        - result_dict: dict, with the keys and dict values
            "losses": {"train_losses": np.ndarray, "val_losses": np.ndarray}
            "train": {"mse": float,
                    "pred": np.ndarray,
                    "true": np.ndarray,
                    "ndcg": float,
                    "rho": SpearmanrResults(correlation=float, pvalue=float)}
            "val":   {"mse": float,
                    "pred": np.ndarray,
                    "true": np.ndarray,
                    "ndcg": float,
                    "rho": SpearmanrResults(correlation=float, pvalue=float)}
            "test":  {"mse": float,
                    "pred": np.ndarray,
                    "true": np.ndarray,
                    "ndcg": float,
                    "rho": SpearmanrResults(correlation=float, pvalue=float)}

        """

        self._dataset_path = dataset_path
        self._encoder_name = encoder_name

        if self._encoder_name not in (
            list(TRANSFORMER_INFO.keys()) + list(CARP_INFO.keys())
        ):
            self._encoder_name = "onehot"

        self._checkpoint = checkpoint
        self._checkpoint_folder = checkpoint_folder

        self._reset_param = reset_param
        self._resample_param = resample_param
        self._embed_batch_size = embed_batch_size
        self._flatten_emb = flatten_emb
        self._embed_folder = embed_folder

        if self._flatten_emb == False:
            self._flatten_emb_name = "noflatten"
        else:
            self._flatten_emb_name = self._flatten_emb

        self._learning_rate = learning_rate
        self._lr_decay = lr_decay
        self._epochs = epochs
        self._early_stop = early_stop
        self._tolerance = tolerance
        self._min_epoch = min_epoch
        self._device = device
        self._all_plot_folder = all_plot_folder
        self._all_result_folder = all_result_folder

        # append checkpoint fraction
        if self._checkpoint != 1:
            self._all_plot_folder += f"-{str(self._checkpoint)}"
            self._all_result_folder += f"-{str(self._checkpoint)}"
            self._embed_folder += f"-{str(self._checkpoint)}"

        if self._reset_param and "-rand" not in self._all_result_folder:
            self._all_result_folder = f"{self._all_result_folder}-rand"
            self._all_plot_folder = f"{self._all_plot_folder}-rand"

        if self._resample_param and "-stat" not in self._all_result_folder:
            self._all_result_folder = f"{self._all_result_folder}-stat"
            self._all_plot_folder = f"{self._all_plot_folder}-stat"

        self._encoder_params = encoder_params

        self._ds_info = DatasetInfo(self._dataset_path)
        self._model_type = self._ds_info.model_type
        self._numb_class = self._ds_info.numb_class
        self._subset_list = self._ds_info.subset_list

        print(f"This dataset includes subsets: {self._subset_list}...")

        encoder_name, encoder_class, total_emb_layer = get_emb_info(encoder_name)

        if encoder_class == ESMEncoder:
            self._encoder_info_dict = TRANSFORMER_INFO
        elif encoder_class == CARPEncoder:
            self._encoder_info_dict = CARP_INFO
        elif encoder_class == OnehotEncoder:

            if "-onehot" not in self._all_result_folder:
                self._all_result_folder = f"{self._all_result_folder}-onehot"
                self._all_plot_folder = f"{self._all_plot_folder}-onehot"

            # TODO aultoto
            if self._flatten_emb == False:
                self._encoder_info_dict = {"onehot": (AA_NUMB,)}
            else:
                self._encoder_info_dict = {"onehot": (MAX_SEQ_LEN * AA_NUMB,)}

        self._seq_start_idx = seq_start_idx
        self._seq_end_idx = seq_end_idx
        self._loader_batch_size = loader_batch_size
        self._worker_seed = worker_seed
        self._if_encode_all = if_encode_all
        self._encoder_params = encoder_params

        if if_multiprocess:
            print("Running different emb layer in parallel...")
            # add the thredpool max_workers=None
            with futures.ProcessPoolExecutor(max_workers=os.cpu_count() - 1) as pool:
                # for each layer train the model and save the model
                for embed_layer in tqdm(range(total_emb_layer)):
                    pool.submit(self.run_pytorch_layer, embed_layer)
        else:
            if isinstance(manual_layer_min, str) and isinstance(manual_layer_max, str):
                print(
                    f"Running pytorch model for layers from {manual_layer_min} to {manual_layer_max}..."
                )
                layer_range = range(int(manual_layer_min), int(manual_layer_max) + 1)
            else:
                layer_range = range(total_emb_layer)

            for embed_layer in layer_range:
                print(f"Running pytorch model for layer {embed_layer}...")
                if if_rerun_layer or (
                    (not if_rerun_layer)
                    and (not os.path.exists(self.get_pytorch_layer_info(embed_layer)))
                ):
                    self.run_pytorch_layer(embed_layer)
                else:
                    print(
                        f"Results for pytorch model for layer {embed_layer} already exists..."
                    )

    def get_pytorch_layer_info(self, embed_layer):
        """
        Get info on pytorch layers
        """
        dataset_subfolder, file_name = get_folder_file_names(
            parent_folder=get_default_output_path(self._all_result_folder),
            dataset_path=self._dataset_path,
            encoder_name=self._encoder_name,
            embed_layer=embed_layer,
            flatten_emb=self._flatten_emb_name,
        )
        return os.path.join(dataset_subfolder, file_name + ".pkl")

    def run_pytorch_layer(self, embed_layer):

        # init model based on datasets
        if self._model_type == "LinearRegression":
            model = LinearRegression(
                input_dim=self._encoder_info_dict[self._encoder_name][0], output_dim=1
            )
            criterion = nn.MSELoss()

        else:

            if self._model_type == "LinearClassifier-Structure":
                classifier_type = "structure"
                criterion = nn.CrossEntropyLoss(ignore_index=-1)

            elif self._model_type == "LinearClassifier-Annotation":
                classifier_type = "annotation"
                criterion = nn.CrossEntropyLoss()

            model = LinearClassifier(
                input_dim=self._encoder_info_dict[self._encoder_name][0],
                numb_class=self._numb_class,
                classifier_type=classifier_type,
            )

        model_name = model.model_name
        model = model.to(self._device, non_blocking=True)
        criterion = criterion.to(self._device, non_blocking=True)

        self._loader_dict = split_protrain_loader(
            dataset_path=self._dataset_path,
            encoder_name=self._encoder_name,
            reset_param=self._reset_param,
            resample_param=self._resample_param,
            embed_batch_size=self._embed_batch_size,
            flatten_emb=self._flatten_emb,
            embed_folder=self._embed_folder,
            embed_layer=embed_layer,
            seq_start_idx=self._seq_start_idx,
            seq_end_idx=self._seq_end_idx,
            subset_list=self._subset_list,
            loader_batch_size=self._loader_batch_size,
            worker_seed=self._worker_seed,
            if_encode_all=self._if_encode_all,
            **self._encoder_params,
        )

        train_losses, val_losses = train(
            model=model,
            criterion=criterion,
            train_loader=self._loader_dict["train"],
            val_loader=self._loader_dict["val"],
            encoder_name=self._encoder_name,
            embed_layer=embed_layer,
            reset_param=self._reset_param,
            resample_param=self._resample_param,
            embed_batch_size=self._embed_batch_size,
            flatten_emb=self._flatten_emb,
            device=self._device,
            learning_rate=self._learning_rate,
            lr_decay=self._lr_decay,
            epochs=self._epochs,
            early_stop=self._early_stop,
            tolerance=self._tolerance,
            min_epoch=self._min_epoch,
            **self._encoder_params,
        )

        # record the losses
        result_dict = {
            "losses": {"train_losses": train_losses, "val_losses": val_losses}
        }

        plot_lc(
            train_losses=train_losses,
            val_losses=val_losses,
            dataset_path=self._dataset_path,
            encoder_name=self._encoder_name,
            embed_layer=embed_layer,
            flatten_emb=self._flatten_emb_name,
            all_plot_folder=get_default_output_path(self._all_plot_folder),
        )

        # now test the model with the test data
        for subset, loader_key in zip(self._subset_list, self._loader_dict.keys()):
            print(f"testing {subset} with loader_key {loader_key}...")

            loss, pred, cls, true = test(
                model=model,
                loader=self._loader_dict[loader_key],
                embed_layer=embed_layer,
                device=self._device,
                criterion=criterion,
            )

            if model_name == "LinearRegression":
                result_dict[subset] = {
                    "mse": loss,
                    "pred": pred,
                    "true": true,
                    "ndcg": ndcg_score(true[None, :], pred[None, :]),
                    "rho": spearmanr(true, pred),
                }

            elif model_name == "LinearClassifier-Annotation":
                result_dict[subset] = {
                    "cross-entropy": loss,
                    "pred": pred,
                    "true": true,
                    "acc": accuracy_score(true, cls),
                    "rocauc": roc_auc_score(
                        true,
                        nn.Softmax(dim=1)(torch.from_numpy(pred)).numpy(),
                        multi_class="ovr",
                    ),
                }

            elif model_name == "LinearClassifier-Structure":
                print(
                    "pred.shape, true.shape, nn.Softmax(dim=1)(torch.from_numpy(pred)).numpy().shape"
                )
                print(
                    pred.shape,
                    true.shape,
                    nn.Softmax(dim=1)(torch.from_numpy(pred)).numpy().shape,
                )
                result_dict[subset] = {
                    "cross-entropy": loss,
                    "pred": pred,
                    "true": true,
                    "acc": accuracy_score(true, cls),
                    "rocauc": roc_auc_score(
                        true,
                        nn.Softmax(dim=1)(torch.from_numpy(pred)).numpy(),
                        multi_class="ovr",
                    )
                    # "rocauc": eval_rocauc(true, pred),
                }

            else:
                print(f"Unrecognize {model_name} as a model_name")

        # TODO del
        dataset_subfolder, file_name = get_folder_file_names(
            parent_folder=get_default_output_path(self._all_result_folder),
            dataset_path=self._dataset_path,
            encoder_name=self._encoder_name,
            embed_layer=embed_layer,
            flatten_emb=self._flatten_emb_name,
        )

        print(f"Saving results for {file_name} to: {dataset_subfolder}...")
        pickle_save(
            what2save=result_dict,
            where2save=os.path.join(dataset_subfolder, file_name + ".pkl"),
        )
```
