## Supplemental data and code for "Feature Reuse and Scaling: Understanding Transfer Learning with Protein Language Models": run_sklearn.py

```
"""Script for run sklearn (currently ridge) models"""

from __future__ import annotations

import os
import random
import numpy as np

from sklearn.linear_model import Ridge, LogisticRegression
from sklearn.metrics import mean_squared_error, log_loss, accuracy_score, roc_auc_score
from sklearn.preprocessing import StandardScaler, LabelEncoder
from scipy.stats import spearmanr

from scr.utils import get_folder_file_names, pickle_save, ndcg_scale
from scr.params.emb import TRANSFORMER_INFO, CARP_INFO
from scr.params.sys import RAND_SEED, SKLEARN_ALPHAS
from scr.encoding.encoding_classes import get_emb_info

from scr.preprocess.data_process import ProtranDataset, DatasetInfo

import warnings
warnings.filterwarnings("ignore")

# seed
random.seed(RAND_SEED)
np.random.seed(RAND_SEED)


class RunRidge:
    """A class for running ridge regression"""

    def __init__(
        self,
        dataset_path: str,
        encoder_name: str,
        checkpoint: float = 1,
        checkpoint_folder: str = "pretrain_checkpoints/carp",
        reset_param: bool = False,
        resample_param: bool = False,
        embed_batch_size: int = 128,
        flatten_emb: bool | str = False,
        embed_folder: str | None = None,
        all_embed_layers: bool = True,
        seq_start_idx: bool | int = False,
        seq_end_idx: bool | int = False,
        if_encode_all: bool = True,
        alphas: np.ndarray | int = SKLEARN_ALPHAS,
        ridge_state: int = RAND_SEED,
        ridge_params: dict | None = None,
        all_result_folder: str = "results/sklearn",
        **encoder_params,
    ) -> None:

        """
        Args:
        - dataset_path: str, full path to the dataset, in pkl or panda readable format
            columns include: sequence, target, set, validation,
            mut_name (optional), mut_numb (optional)
        - encoder_name: str, the name of the encoder
        - checkpoint: float = 1, the 0.5, 0.25, 0.125 checkpoint of the CARP encoder or full
        - checkpoint_folder: str = "pretrain_checkpoints/carp", folder for carp encoders
        - reset_param: bool = False, if update the full model to xavier_uniform_
        - resample_param: bool = False, if update the full model to xavier_normal_
        - embed_batch_size: int, set to 0 to encode all in a single batch
        - flatten_emb: bool or str, if and how (one of ["max", "mean"]) to flatten the embedding
        - embed_folder: str = None, path to presaved embedding
        - all_embed_layers: bool = True, if include all embed layers
        - seq_start_idx: bool | int = False, the index for the start of the sequence
        - seq_end_idx: bool | int = False, the index for the end of the sequence
        - if_encode_all: bool = True, if encode all embed layers all at once
        - alphas: np.ndarray, arrays of alphas to be tested
        - ridge_state: int = RAND_SEED, seed the ridge regression
        - ridge_params: dict | None = None, other ridge regression args
        - all_result_folder: str = "results/train_val_test", the parent folder for all results
        - encoder_params: kwarg, additional parameters for encoding
        """

        self.dataset_path = dataset_path
        self.encoder_name = encoder_name
        self.reset_param = reset_param
        self.resample_param = resample_param
        self.embed_batch_size = embed_batch_size
        self.flatten_emb = flatten_emb
        self.embed_folder = embed_folder
        self.all_embed_layers = all_embed_layers
        self.seq_start_idx = seq_start_idx
        self.seq_end_idx = seq_end_idx
        self.if_encode_all = if_encode_all
        self.encoder_params = encoder_params

        self.checkpoint = checkpoint
        self.checkpoint_folder = checkpoint_folder

        if not isinstance(alphas, np.ndarray):
            alphas = np.array([alphas])
        self.alphas = alphas

        self.ridge_state = ridge_state
        self.ridge_params = ridge_params
        self.all_result_folder = all_result_folder

        # append init
        if self.reset_param and "-rand" not in self.all_result_folder:
            self.all_result_folder += "-rand"
        if self.reset_param and "-rand" not in self.embed_folder:
            self.embed_folder += "-rand"

        if self.resample_param and "-stat" not in self.all_result_folder:
            self.all_result_folder += "-stat"
        if self.resample_param and "-stat" not in self.embed_folder:
            self.embed_folder += "-stat"

        # append checkpoint fraction
        if self.checkpoint != 1:
            self.all_result_folder += f"-{str(self.checkpoint)}"
            self.embed_folder += f"-{str(self.checkpoint)}"

        all_ridge_results = {}

        self.encoder_name, _, total_emb_layer = get_emb_info(self.encoder_name)

        if self.all_embed_layers:
            print("loading all embed layers...")
            # loader has ALL embedding layers
            self.train_ds, self.val_ds, self.test_ds = (
                ProtranDataset(
                    dataset_path=self.dataset_path,
                    subset=subset,
                    encoder_name=self.encoder_name,
                    reset_param=self.reset_param,
                    resample_param=self.resample_param,
                    embed_batch_size=self.embed_batch_size,
                    flatten_emb=self.flatten_emb,
                    embed_folder=self.embed_folder,
                    embed_layer=None,
                    seq_start_idx=self.seq_start_idx,
                    seq_end_idx=self.seq_end_idx,
                    if_encode_all=self.if_encode_all,
                    **self.encoder_params,
                )
                for subset in ["train", "val", "test"]
            )

        for layer in range(total_emb_layer):
            all_ridge_results[layer] = self.run_ridge_layer(embed_layer=layer,)

        self._all_ridge_results = all_ridge_results

    def sk_test(
        self, model: sklearn.linear_model, ds: ProtranDataset, embed_layer: int
    ):
        """
        A function for testing sklearn models for a specific layer of embeddings

        Args:
        - model: sklearn.linear_model, trained model
        - ds: ProtranDataset, train, val, or test dataset
        - embed_layer: int, specific layer of the embedding

        Returns:
        - np.concatenate(pred): np.ndarray, 1D predicted fitness values
        - np.concatenate(true): np.ndarry, 1D true fitness values
        """

        return (
            model.predict(getattr(ds, "layer" + str(embed_layer))).squeeze(),
            # getattr(train_ds, "layer" + str(embed_layer))
            ds.y.squeeze(),
        )

    def pick_model(
        self, embed_layer: int, train_ds: ProtranDataset, val_ds: ProtranDataset
    ):
        """
        A function for picking the best model for given alaphs, meaning
        lower train_mse and higher test_ndcg
        NOTE: alphas tuning is NOT currently optimal

        Args:
        - embed_layer: int, specific layer of the embedding

        Returns:
        - sklearn.linear_model, the model with the best alpha
        """

        # init values for comparison
        best_mse = np.Inf
        best_ndcg = -1
        best_rho = -1
        best_model = None

        # loop through all alphas
        for alpha in self.alphas:

            # init model for each alpha
            if self.ridge_params is None:
                self.ridge_params = {}
            model = Ridge(
                alpha=alpha, random_state=self.ridge_state, **self.ridge_params
            )

            # fit the model for a given layer of embedding
            fitness_scaler = StandardScaler()

            if self.all_embed_layers:
                train_ds = self.train_ds
                val_ds = self.val_ds

            model.fit(
                getattr(train_ds, "layer" + str(embed_layer)),
                fitness_scaler.fit_transform(train_ds.y),
            )

            # eval the model with train and test
            train_pred, train_true = self.sk_test(
                model, train_ds, embed_layer=embed_layer
            )
            val_pred, val_true = self.sk_test(model, val_ds, embed_layer=embed_layer)

            # calc the metrics
            train_mse = mean_squared_error(train_true, train_pred)
            val_ndcg = ndcg_scale(val_true, val_pred)
            val_rho = spearmanr(val_true, val_pred)[0]

            # update the model if it has lower train_mse and higher val_ndcg
            if train_mse < best_mse and val_ndcg > best_ndcg:
                best_model = model
                best_mse = train_mse
                best_ndcg = val_ndcg
                best_rho = val_rho

        print(f"best model is {best_model}")
        return best_model

    def run_ridge_layer(
        self, embed_layer: int,
    ):

        """
        A function for running ridge regression for a given layer of embedding

        Args:
        - embed_layer: int, specific layer of the embedding

        Returns:
        - dict, with the keys and dict values
            "train": {"mse": float,
                    "pred": np.ndarray,
                    "true": np.ndarray,
                    "ndcg": float,
                    "rho": SpearmanrResults(correlation=float, pvalue=float)}
            "val":   {"mse": float,
                    "pred": np.ndarray,
                    "true": np.ndarray,
                    "ndcg": float,
                    "rho": SpearmanrResults(correlation=float, pvalue=float)}
            "test":  {"mse": float,
                    "pred": np.ndarray,
                    "true": np.ndarray,
                    "ndcg": float,
                    "rho": SpearmanrResults(correlation=float, pvalue=float)}
        """

        # set up the datasets
        if self.all_embed_layers:
            ds_list = [self.train_ds, self.val_ds, self.test_ds]
        else:
            print(f"Getting embed for {embed_layer}...")
            ds_list = [
                ProtranDataset(
                    dataset_path=self.dataset_path,
                    subset=subset,
                    encoder_name=self.encoder_name,
                    checkpoint=self.checkpoint,
                    checkpoint_folder=self.checkpoint_folder,
                    reset_param=self.reset_param,
                    resample_param=self.resample_param,
                    embed_batch_size=self.embed_batch_size,
                    flatten_emb=self.flatten_emb,
                    embed_folder=self.embed_folder,
                    embed_layer=embed_layer,
                    seq_start_idx=self.seq_start_idx,
                    seq_end_idx=self.seq_end_idx,
                    if_encode_all=self.if_encode_all,
                    **self.encoder_params,
                )
                for subset in ["train", "val", "test"]
            ]

        # train and get the best alpha
        best_model = self.pick_model(
            embed_layer=embed_layer, train_ds=ds_list[0], val_ds=ds_list[1]
        )

        # init dict for resulted outputs
        result_dict = {}

        # now test the model with the test data
        for subset, ds in zip(["train", "val", "test"], ds_list):
            pred, true = self.sk_test(best_model, ds, embed_layer=embed_layer)

            result_dict[subset] = {
                "mse": mean_squared_error(true, pred),
                "pred": pred,
                "true": true,
                "ndcg": ndcg_scale(true, pred),
                "rho": spearmanr(true, pred),
            }

        dataset_subfolder, file_name = get_folder_file_names(
            parent_folder=self.all_result_folder,
            dataset_path=self.dataset_path,
            encoder_name=self.encoder_name,
            embed_layer=embed_layer,
            flatten_emb=self.flatten_emb,
        )

        print(f"Saving results for {file_name} to: {dataset_subfolder}...")
        pickle_save(
            what2save=result_dict,
            where2save=os.path.join(dataset_subfolder, file_name + ".pkl"),
        )

        return result_dict

    @property
    def all_ridge_results(self):
        """
        Returns:
        - dict, with the keys and dict values
            "layer#": {
                        "train": {"mse": float,
                                "pred": np.ndarray,
                                "true": np.ndarray,
                                "ndcg": float,
                                "rho": SpearmanrResults(correlation=float, pvalue=float)}
                        "val":   {"mse": float,
                                "pred": np.ndarray,
                                "true": np.ndarray,
                                "ndcg": float,
                                "rho": SpearmanrResults(correlation=float, pvalue=float)}
                        "test":  {"mse": float,
                                "pred": np.ndarray,
                                "true": np.ndarray,
                                "ndcg": float,
                                "rho": SpearmanrResults(correlation=float, pvalue=float)}
                        }
        """
        return self._all_ridge_results


class RunSK:
    """
    A class for running sklearn models 
    """

    def __init__(
        self,
        dataset_path: str,
        encoder_name: str,
        checkpoint: float = 1,
        checkpoint_folder: str = "pretrain_checkpoints/carp",
        reset_param: bool = False,
        resample_param: bool = False,
        embed_batch_size: int = 128,
        flatten_emb: bool | str = False,
        embed_folder: str | None = None,
        all_embed_layers: bool = True,
        seq_start_idx: bool | int = False,
        seq_end_idx: bool | int = False,
        if_encode_all: bool = True,
        alphas: np.ndarray | int = SKLEARN_ALPHAS,
        sklearn_state: int = RAND_SEED,
        sklearn_params: dict | None = None,
        all_result_folder: str = "results/sklearn",
        **encoder_params,
    ) -> None:

        """
        Args:
        - dataset_path: str, full path to the dataset, in pkl or panda readable format
            columns include: sequence, target, set, validation,
            mut_name (optional), mut_numb (optional)
        - encoder_name: str, the name of the encoder
        - checkpoint: float = 1, the 0.5, 0.25, 0.125 checkpoint of the CARP encoder or full
        - checkpoint_folder: str = "pretrain_checkpoints/carp", folder for carp encoders
        - reset_param: bool = False, if update the full model to xavier_uniform_
        - resample_param: bool = False, if update the full model to xavier_normal_
        - embed_batch_size: int, set to 0 to encode all in a single batch
        - flatten_emb: bool or str, if and how (one of ["max", "mean"]) to flatten the embedding
        - embed_folder: str = None, path to presaved embedding
        - seq_start_idx: bool | int = False, the index for the start of the sequence
        - seq_end_idx: bool | int = False, the index for the end of the sequence
        - alphas: np.ndarray, arrays of alphas to be tested
        - sklearn_state: int = RAND_SEED, seed the ridge or logistic regression
        - sklearn_params: dict | None = None, other ridge or logistic regression args
        - all_result_folder: str = "results/train_val_test", the parent folder for all results
        - encoder_params: kwarg, additional parameters for encoding
        """

        self.dataset_path = dataset_path
        self.encoder_name = encoder_name
        self.reset_param = reset_param
        self.resample_param = resample_param
        self.embed_batch_size = embed_batch_size
        self.flatten_emb = flatten_emb
        self.embed_folder = embed_folder
        self.all_embed_layers = all_embed_layers
        self.seq_start_idx = seq_start_idx
        self.seq_end_idx = seq_end_idx
        self.if_encode_all = if_encode_all
        self.encoder_params = encoder_params

        self.checkpoint = checkpoint
        self.checkpoint_folder = checkpoint_folder

        if not isinstance(alphas, np.ndarray):
            alphas = np.array([alphas])
        self.alphas = alphas

        self.sklearn_state = sklearn_state
        self.sklearn_params = sklearn_params
        self.all_result_folder = all_result_folder

        # append init
        if self.reset_param and "-rand" not in self.all_result_folder:
            self.all_result_folder += "-rand"
        if self.reset_param and "-rand" not in self.embed_folder:
            self.embed_folder += "-rand"

        if self.resample_param and "-stat" not in self.all_result_folder:
            self.all_result_folder += "-stat"
        if self.resample_param and "-stat" not in self.embed_folder:
            self.embed_folder += "-stat"

        # append checkpoint fraction
        if self.checkpoint != 1:
            self.all_result_folder += f"-{str(self.checkpoint)}"
            self.embed_folder += f"-{str(self.checkpoint)}"

        # update encoder name and total embedding layers
        if self.encoder_name in TRANSFORMER_INFO.keys():
            total_emb_layer = TRANSFORMER_INFO[encoder_name][1] + 1
        elif self.encoder_name in CARP_INFO.keys():
            total_emb_layer = CARP_INFO[encoder_name][1] + 1
        else:
            # for onehot
            self.encoder_name = "onehot"
            total_emb_layer = 1

        if self.encoder_name == "onehot" and "onehot" not in self.all_result_folder:
            self.all_result_folder = f"{self.all_result_folder}-onehot"
        # if self.all_embed_layers:
        #    print("loading all embed layers...")
        # loader has ALL embedding layers
        print("RunSK prepping all ds...")

        print(f"self.if_encode_all: {self.if_encode_all}")
        print(f"self.embed_folder: {self.embed_folder}")

        self.train_ds, self.val_ds, self.test_ds = (
            ProtranDataset(
                dataset_path=self.dataset_path,
                subset=subset,
                encoder_name=self.encoder_name,
                reset_param=self.reset_param,
                resample_param=self.resample_param,
                embed_batch_size=self.embed_batch_size,
                flatten_emb=self.flatten_emb,
                embed_folder=self.embed_folder,
                embed_layer=None,
                seq_start_idx=self.seq_start_idx,
                seq_end_idx=self.seq_end_idx,
                if_encode_all=self.if_encode_all,
                **self.encoder_params,
            )
            for subset in ["train", "val", "test"]
        )

        self._ds_info = DatasetInfo(self.dataset_path)
        self._model_type = self._ds_info.model_type
        self._numb_class = self._ds_info.numb_class

        print(f"Running {self._model_type}...")

        # pick ridge regression if y numerical
        if self._model_type == "LinearRegression":
            self.sklearn_model = Ridge

        # pick logistic regression if y is categorical
        elif self._model_type == "LinearClassifier":
            
            """le = LabelEncoder()
            self.train_ds.y, self.val_ds.y, self.test_ds.y = [
                le.fit_transform(y.flatten())
                for y in [self.train_ds.y, self.val_ds.y, self.test_ds.y]
            ]"""
            self.sklearn_model = LogisticRegression
            # convert alpha to C
            self.alphas = 1 / self.alphas
            # add other params
            if self.sklearn_params is None:
                self.sklearn_params = {}
            elif "max_iter" not in self.sklearn_params.keys():
                self.sklearn_params["max_iter"] = 10000
            
            self.sklearn_params["multi_class"] = "multinomial"

        all_sklearn_results = {}

        for layer in range(total_emb_layer):
            all_sklearn_results[layer] = self.run_sklearn_layer(embed_layer=layer,)

        self._all_sklearn_results = all_sklearn_results

    def sk_test(
        self, model: sklearn.linear_model, ds: ProtranDataset, embed_layer: int
    ):
        """
        A function for testing sklearn models for a specific layer of embeddings

        Args:
        - model: sklearn.linear_model, trained model
        - ds: ProtranDataset, train, val, or test dataset
        - embed_layer: int, specific layer of the embedding

        Returns:
        - np.concatenate(pred): np.ndarray, 1D predicted fitness values
        - np.concatenate(true): np.ndarry, 1D true fitness values
        - 
        """

        if self.sklearn_model == Ridge:
            pred_prob = None
        else:
            pred_prob = model.predict_proba(
                getattr(ds, "layer" + str(embed_layer))
            ).squeeze()

        return (
            model.predict(getattr(ds, "layer" + str(embed_layer))).squeeze(),
            ds.y.squeeze(),
            pred_prob,
        )

    def pick_model(
        self, embed_layer: int, train_ds: ProtranDataset, val_ds: ProtranDataset
    ):
        """
        A function for picking the best model for given alaphs, meaning
        lower train_mse and higher test_ndcg
        NOTE: alphas tuning is NOT currently optimal

        Args:
        - embed_layer: int, specific layer of the embedding

        Returns:
        - sklearn.linear_model, the model with the best alpha
        """

        # init values for comparison
        if self.sklearn_model == Ridge:
            best_mse = np.Inf
            best_ndcg = -1
            best_rho = -1
        else:
            best_loss = np.Inf
            best_acc = 0
            best_auc = 0

        best_model = None

        # loop through all alphas
        for alpha in self.alphas:

            # init model for each alpha
            if self.sklearn_params is None:
                self.sklearn_params = {}

            if self._model_type == "LinearRegression":
                self.sklearn_params["alpha"] = alpha
            elif self._model_type == "LinearClassifier":
                self.sklearn_params["C"] = alpha

            model = self.sklearn_model(
                random_state=self.sklearn_state, **self.sklearn_params
            )
            
            # fit the model for a given layer of embedding
            fitness_scaler = StandardScaler()

            if self.all_embed_layers:
                train_ds = self.train_ds
                val_ds = self.val_ds

            if self._model_type == "LinearRegression":
                y = fitness_scaler.fit_transform(train_ds.y)
            else:
                y = train_ds.y.ravel()

            model.fit(getattr(train_ds, "layer" + str(embed_layer)), y)

            # eval the model with train and test
            train_pred, train_true, train_prob = self.sk_test(
                model, train_ds, embed_layer=embed_layer
            )
            val_pred, val_true, val_prob = self.sk_test(
                model, val_ds, embed_layer=embed_layer
            )

            if self.sklearn_model == Ridge:
                # calc the metrics
                train_mse = mean_squared_error(train_true, train_pred)
                val_ndcg = ndcg_scale(val_true, val_pred)
                val_rho = spearmanr(val_true, val_pred)[0]

                # update the model if it has lower train_mse and higher val_ndcg
                if train_mse < best_mse and val_ndcg > best_ndcg:
                    best_model = model
                    best_mse = train_mse
                    best_ndcg = val_ndcg
                    best_rho = val_rho

            else:
                # calc the metrics
                train_loss = log_loss(train_true, train_prob)
                val_acc = accuracy_score(val_true, val_pred)
                val_auc = roc_auc_score(val_true, val_prob, multi_class="ovr")

                # update the model if it has lower log_loss and higher val_auc
                if train_loss < best_loss and val_auc > best_auc:
                    best_loss = train_loss
                    best_acc = val_acc
                    best_auc = val_auc

        print(f"best model is {best_model}")
        return best_model

    def run_sklearn_layer(
        self, embed_layer: int,
    ):

        """
        A function for running ridge or logistics regression for a given layer of embedding

        Args:
        - embed_layer: int, specific layer of the embedding

        Returns:
        - dict, with the keys and dict values
            "train": {"mse": float,
                    "pred": np.ndarray,
                    "true": np.ndarray,
                    "ndcg": float,
                    "rho": SpearmanrResults(correlation=float, pvalue=float)}
            "val":   {"mse": float,
                    "pred": np.ndarray,
                    "true": np.ndarray,
                    "ndcg": float,
                    "rho": SpearmanrResults(correlation=float, pvalue=float)}
            "test":  {"mse": float,
                    "pred": np.ndarray,
                    "true": np.ndarray,
                    "ndcg": float,
                    "rho": SpearmanrResults(correlation=float, pvalue=float)}
        """

        # set up the datasets
        if self.all_embed_layers:
            print("Getting embed for all layers...")
            ds_list = [self.train_ds, self.val_ds, self.test_ds]
        else:
            print(f"Getting embed for {embed_layer}...")
            ds_list = [
                ProtranDataset(
                    dataset_path=self.dataset_path,
                    subset=subset,
                    encoder_name=self.encoder_name,
                    reset_param=self.reset_param,
                    resample_param=self.resample_param,
                    embed_batch_size=self.embed_batch_size,
                    flatten_emb=self.flatten_emb,
                    embed_folder=self.embed_folder,
                    embed_layer=embed_layer,
                    seq_start_idx=self.seq_start_idx,
                    seq_end_idx=self.seq_end_idx,
                    if_encode_all=self.if_encode_all,
                    **self.encoder_params,
                )
                for subset in ["train", "val", "test"]
            ]

        # train and get the best alpha
        best_model = self.pick_model(
            embed_layer=embed_layer, train_ds=ds_list[0], val_ds=ds_list[1]
        )

        # init dict for resulted outputs
        result_dict = {}

        # now test the model with the test data
        for subset, ds in zip(["train", "val", "test"], ds_list,):
            pred, true, prob = self.sk_test(best_model, ds, embed_layer=embed_layer)

            if self.sklearn_model == Ridge:
                result_dict[subset] = {
                    "mse": mean_squared_error(true, pred),
                    "pred": pred,
                    "true": true,
                    "ndcg": ndcg_scale(true, pred),
                    "rho": spearmanr(true, pred),
                }

            else:
                result_dict[subset] = {
                    "log": log_loss(true, prob),
                    "pred": pred,
                    "prob": prob,
                    "true": true,
                    "acc": accuracy_score(true, pred),
                    "rocauc": roc_auc_score(true, prob, multi_class="ovr"),
                }

        dataset_subfolder, file_name = get_folder_file_names(
            parent_folder=self.all_result_folder,
            dataset_path=self.dataset_path,
            encoder_name=self.encoder_name,
            embed_layer=embed_layer,
            flatten_emb=self.flatten_emb,
        )

        print(f"Saving results for {file_name} to: {dataset_subfolder}...")
        pickle_save(
            what2save=result_dict,
            where2save=os.path.join(dataset_subfolder, file_name + ".pkl"),
        )

        return result_dict

    @property
    def all_sklearn_results(self):
        """
        Returns:
        - dict, with the keys and dict values
            "layer#": {
                        "train": {"mse": float,
                                "pred": np.ndarray,
                                "true": np.ndarray,
                                "ndcg": float,
                                "rho": SpearmanrResults(correlation=float, pvalue=float)}
                        "val":   {"mse": float,
                                "pred": np.ndarray,
                                "true": np.ndarray,
                                "ndcg": float,
                                "rho": SpearmanrResults(correlation=float, pvalue=float)}
                        "test":  {"mse": float,
                                "pred": np.ndarray,
                                "true": np.ndarray,
                                "ndcg": float,
                                "rho": SpearmanrResults(correlation=float, pvalue=float)}
                        }
        """
        return self._all_sklearn_results
```
