## Supplemental data and code for "Feature Reuse and Scaling: Understanding Transfer Learning with Protein Language Models": train_test.py

```
"""A script with Pytorch model training and testing details"""

from __future__ import annotations

import random
import numpy as np
import torch
from torch import nn
from torch.utils.data import DataLoader
from tqdm import tqdm

from sklearn.metrics import mean_squared_error, log_loss, accuracy_score, roc_auc_score
from sklearn.preprocessing import LabelEncoder

from scr.params.sys import RAND_SEED, DEVICE

# seed everything
random.seed(RAND_SEED)
np.random.seed(RAND_SEED)
torch.manual_seed(RAND_SEED)
torch.cuda.manual_seed(RAND_SEED)
torch.cuda.manual_seed_all(RAND_SEED)
torch.backends.cudnn.deterministic = True


def get_x_y(
    model_name,
    device,
    batch,
    embed_layer: int,
):
    """
    A function process x and y from the loader

    Args:
    -

    Returns:
    - x
    - y
    """

    # for each batch: y, sequence, mut_name, mut_numb, [layer0, ...]
    # or: y, seuqence, mut_name, mut_numb, layer0, ...

    embs = batch[4]
    if torch.is_tensor(embs):
        x = embs
    else:
        x = embs[embed_layer]
    y = batch[0]

    # ss3 / ss8 type
    if model_name == "LinearClassifier-Structure":

        y = torch.squeeze(y)
        x = torch.cat([seq[ss != -1, :] for (seq, ss) in zip(x, y)]).to(torch.float32)
        y = y.flatten()[y.flatten() != -1].to(torch.double).to(torch.long)

    return x.to(device, non_blocking=True), y.to(device, non_blocking=True)


def run_epoch(
    model: nn.Module,
    loader: DataLoader,
    encoder_name: str,
    embed_layer: int,
    reset_param: bool = False,
    resample_param: bool = False,
    embed_batch_size: int = 0,
    flatten_emb: bool | str = False,
    device: torch.device | str = DEVICE,
    criterion: nn.Module | None = None,
    optimizer: torch.optim.Optimizer | None = None,
    **encoder_params,
) -> float:

    """
    Runs one epoch.

    Args:
    - model: nn.Module, already moved to device
    - loader: torch.utils.data.DataLoader
    - device: torch.device or str
    - criterion: optional nn.Module, loss function, already moved to device
    - optimizer: optional torch.optim.Optimizer, must also provide criterion,
        only provided for training

    Returns:
    - float, average loss over batches
    """
    if optimizer is not None:
        assert criterion is not None
        model.train()
        is_train = True
    else:
        model.eval()
        is_train = False

    model_name = model.model_name
    cum_loss = 0.0

    with torch.set_grad_enabled(is_train):
        # if not if_encode_all:
        # for each batch: y, sequence, mut_name, mut_numb, [layer0, ...]

        for batch in loader:

            x, y = get_x_y(model_name, device, batch, embed_layer)

            outputs = model(x)

            if criterion is not None:
                if model_name == "LinearRegression":
                    loss = criterion(outputs, y.float())
                elif "LinearClassifier" in model_name:
                    loss = criterion(outputs, y.squeeze())

                if optimizer is not None:
                    optimizer.zero_grad()
                    loss.backward()
                    optimizer.step()

                cum_loss += loss.item()

    return cum_loss / len(loader)


def train(
    model: nn.Module,
    criterion: nn.Module,
    train_loader: DataLoader,
    val_loader: DataLoader,
    encoder_name: str,
    embed_layer: int,
    reset_param: bool = False,
    resample_param: bool = False,
    embed_batch_size: int = 0,
    flatten_emb: bool | str = False,
    device: torch.device | str = DEVICE,
    learning_rate: float = 1e-4,
    lr_decay: float = 0.1,
    epochs: int = 100,
    early_stop: bool = True,
    tolerance: int = 10,
    min_epoch: int = 5,
    **encoder_params,
) -> tuple[np.ndarray, np.ndarray]:

    """
    Args:
    - model: nn.Module, already moved to device
    - train_loader: torch.utils.data.DataLoader,
    - val_loader: torch.utils.data.DataLoader,
    - criterion: nn.Module, loss function, already moved to device
    - device: torch.device or str
    - learning_rate: float
    - lr_decay: float, factor by which to decay LR on plateau
    - epochs: int, number of epochs to train for
    - early_stop: bool = True,

    Returns:
    - tuple of np.ndarray, (train_losses, val_losses)
        train/val_losses: np.ndarray, shape [epochs], entries are average loss
        over batches for that epoch
    """

    optimizer = torch.optim.Adam(model.parameters(), lr=learning_rate)
    scheduler = torch.optim.lr_scheduler.ReduceLROnPlateau(
        optimizer, mode="min", factor=lr_decay
    )

    train_losses = np.zeros(epochs)
    val_losses = np.zeros(epochs)

    # init for early stopping
    counter = 0
    min_val_loss = np.Inf

    for epoch in tqdm(range(epochs)):

        train_losses[epoch] = run_epoch(
            model=model,
            loader=train_loader,
            encoder_name=encoder_name,
            embed_layer=embed_layer,
            reset_param=reset_param,
            resample_param=resample_param,
            embed_batch_size=embed_batch_size,
            flatten_emb=flatten_emb,
            device=device,
            criterion=criterion,
            optimizer=optimizer,
            **encoder_params,
        )

        val_loss = run_epoch(
            model=model,
            loader=val_loader,
            encoder_name=encoder_name,
            embed_layer=embed_layer,
            reset_param=reset_param,
            resample_param=resample_param,
            embed_batch_size=embed_batch_size,
            flatten_emb=flatten_emb,
            device=device,
            criterion=criterion,
            optimizer=None,
            **encoder_params,
        )
        val_losses[epoch] = val_loss

        scheduler.step(val_loss)

        if early_stop:
            # when val loss decrease, reset min loss and counter
            if val_loss < min_val_loss:
                min_val_loss = val_loss
                counter = 0
            else:
                counter += 1

            if epoch > min_epoch and counter == tolerance:
                break

    return train_losses, val_losses


def test(
    model: nn.Module,
    loader: DataLoader,
    embed_layer: int,
    criterion: nn.Module | None,
    device: torch.device | str = DEVICE,
    print_every: int = 1000,
) -> tuple[float, np.ndarray, np.ndarray]:
    """
    Runs one epoch of testing, returning predictions and labels.

    Args:
    - model: nn.Module, already moved to device
    - device: torch.device or str
    - loader: torch.utils.data.DataLoader
    - criterion: optional nn.Module, loss function, already moved to device
    - print_every: int, how often (number of batches) to print avg loss

    Returns: tuple (avg_loss, preds, labels)
    - avg_loss: float, average loss per training example
    - preds: np.ndarray, shape [num_examples, ...], predictions over dataset
    - labels: np.ndarray, shape [num_examples, ...], dataset labels
    """
    model.eval()
    msg = "[{step:5d}] loss: {loss:.3f}"

    model_name = model.model_name
    cum_loss = 0.0

    pred_probs = []
    pred_classes = []
    labels = []

    with torch.no_grad():

        for i, batch in enumerate(tqdm(loader)):
            # for each batch: y, sequence, mut_name, mut_numb, [layer0, ...]
            x, y = get_x_y(model_name, device, batch, embed_layer)

            # forward + backward + optimize
            outputs = model(x)

            # append results
            labels.append(y.detach().cpu().squeeze().numpy())

            # append class
            if "LinearClassifier" in model_name:
                pred_classes.append(
                    outputs.detach()
                    .cpu()
                    .data.max(1, keepdim=True)[1]
                    .squeeze()
                    .numpy()
                )

            pred_probs.append(outputs.detach().cpu().squeeze().numpy())

            if criterion is not None:
                if model_name == "LinearRegression":
                    loss = criterion(outputs, y)
                elif "LinearClassifier" in model_name:
                    loss = criterion(outputs, y.squeeze())

                cum_loss += loss.item()

                if ((i + 1) % print_every == 0) or (i + 1 == len(loader)):
                    tqdm.write(msg.format(step=i + 1, loss=cum_loss / len(loader)))

    avg_loss = cum_loss / len(loader)

    if pred_classes == []:
        pred_classes_conc = pred_classes
    else:
        pred_classes_conc = np.concatenate(pred_classes)

    return (
        avg_loss,
        np.concatenate(pred_probs),
        pred_classes_conc,
        np.concatenate(labels),
    )
```
