## Supplemental data and code for "Feature Reuse and Scaling: Understanding Transfer Learning with Protein Language Models": aa.py

```
"""describe properties of amino acids"""

from __future__ import annotations

import pandas as pd

# Amino acid code conversion
AA_DICT = {
    "Ala": "A",
    "Asx": "B",
    "Cys": "C",
    "Asp": "D",
    "Glu": "E",
    "Phe": "F",
    "Gly": "G",
    "His": "H",
    "Ile": "I",
    "Lys": "K",
    "Leu": "L",
    "Met": "M",
    "Asn": "N",
    "Pro": "P",
    "Gln": "Q",
    "Arg": "R",
    "Ser": "S",
    "Thr": "T",
    "Scy": "U",
    "Val": "V",
    "Trp": "W",
    "Xxx": "X",
    "Tyr": "Y",
    "Ter": "*",
}

# the upper case three letter code for the amino acids
ALL_AAS_TLC_DICT = {k.upper(): v for k, v in AA_DICT.items() if v != "*"}

# the upper case three letter code for the amino acids
ALL_AAS_TLC = list(ALL_AAS_TLC_DICT.keys())

# propertyies of the amino acids
AA_PROP_DICT = {
    "R": "Positive",
    "H": "Positive",
    "K": "Positive",
    "D": "Negative",
    "E": "Negative",
    "S": "Polar uncharged",
    "T": "Polar uncharged",
    "N": "Polar uncharged",
    "Q": "Polar uncharged",
    "B": "Polar",
    "C": "Special",
    "X": "Unknown",
    "U": "Special",
    "G": "Special",
    "P": "Special",
    "A": "Hydrophobic",
    "V": "Hydrophobic",
    "I": "Hydrophobic",
    "L": "Hydrophobic",
    "M": "Hydrophobic",
    "F": "Hydrophobic",
    "Y": "Hydrophobic",
    "W": "Hydrophobic",
}

# All canonical amino acids
ALL_AAS = AA_PROP_DICT.keys()
AA_NUMB = len(ALL_AAS)
ALLOWED_AAS = set(ALL_AAS)

# Create a dictionary that links each amino acid to an index
AA_TO_IND = {aa: i for i, aa in enumerate(ALL_AAS)}

# Convert the AA_PROP dictionary to dataframe
AA_PROP_DF = pd.DataFrame(
    AA_PROP_DICT.values(), index=ALL_AAS, columns=["property"]
)
AA_PROP_DF.index.names = ["residue"]
```
