## Supplemental data and code for "Feature Reuse and Scaling: Understanding Transfer Learning with Protein Language Models": emb.py

```
"""Embedding constants"""

from copy import deepcopy

ARCH_TYPE = ["carp", "esm"]

ARCH_AB_DICT = {"rand": "random init", "stat": "stat transfer"}
ARCH_AB = sorted(deepcopy(list(ARCH_AB_DICT.keys())))

ARCH_BAR_LAYER = [0, 2, 4, 6]

ARCH_CUT_DICT = {"": [2, 3, 4, 6], "carp": [2, 4, 6, 12], "esm": [2, 3, 4, 6]}

# the longest seq for esm1b is 1022 with CLS and EOS
MAX_SEQ_LEN = 1022

# encoder_name: (embedding dim, num layers, token dim)
TRANSFORMER_INFO = {
    "esm1_t6_43M_UR50S": (768, 6, 2),
    "esm1_t12_85M_UR50S": (768, 12, 2),
    "esm1_t34_670M_UR50S": (1280, 34, 2),
    "esm1b_t33_650M_UR50S": (1280, 33, 2),
}

# encoder_name: (d_model, n_layers)
CARP_INFO = {
    "carp_600k": (128, 16),
    "carp_38M": (1024, 16),
    "carp_76M": (1024, 32),
    "carp_640M": (1280, 56),
}

# model parameter number in M
MODEL_SIZE = {
    "esm1_t6_43M_UR50S": 43,
    "esm1_t12_85M_UR50S": 85,
    "esm1_t34_670M_UR50S": 670,
    "esm1b_t33_650M_UR50S": 650,
    "carp_600k": 0.6,
    "carp_38M": 38,
    "carp_76M": 76,
    "carp_640M": 640,
    "onehot": 0.02,
}

EMB_SIMPLE_MAP = {
    "esm1_t6_43M_UR50S": "ESM1 43M",
    "esm1_t12_85M_UR50S": "ESM1 85M",
    "esm1_t34_670M_UR50S": "ESM1 670M",
    "esm1b_t33_650M_UR50S": "ESM1b 650M",
    "carp_600k": "CARP 0.6M",
    "carp_38M": "CARP 38M",
    "carp_76M": "CARP 76M",
    "carp_640M": "CARP 640M",
    "onehot": "Onehot"
}

EMB_SIZE_SIMPLE = {
    "esm1_t6_43M_UR50S": "Small",
    "esm1_t12_85M_UR50S": "Medium",
    "esm1_t34_670M_UR50S": "Large",
    "esm1b_t33_650M_UR50S": "Large*",
    "carp_600k": "Mini",
    "carp_38M": "Small",
    "carp_76M": "Medium",
    "carp_640M": "Large",
}

BASELINE_NAME_DICT = {"onehot": "One-hot", "rand": "Random init", "stat": "Stat transfer"}
EMB_SIZE_NAME_SIMPLE = ["Small", "Medium", "Large"]

# the embeddings to use for task oriented plots
EMB4TASK = ["onehot", 
            "esm1_t6_43M_UR50S", "esm1_t12_85M_UR50S", "esm1b_t33_650M_UR50S",
            "carp_38M", "carp_76M", "carp_640M"
            ]

# emb model parameter number in M
EMB_MODEL_SIZE = {k: v for k, v in deepcopy(MODEL_SIZE).items() if k != "onehot"}

MODEL_LAYER = {
    model_name: model_dets[1]
    for info_dict in [deepcopy(TRANSFORMER_INFO), deepcopy(CARP_INFO), {"onehot": (1, 1)}]
    for model_name, model_dets in info_dict.items()
}

EMB_MODEL_LAYER = {k: v for k, v in deepcopy(MODEL_LAYER).items() if k != "onehot"}

CARP_MODEL_LAYER = {k: v[-1] for k, v in deepcopy(CARP_INFO).items()}

CHECKPOINT_PERCENT = [0.125, 0.25, 0.5, 1]

# TODO integrate to be auto from sheet
CARP_CHECKPOINTS = {
    "carp_600k": {0.5: 239263, 0.25: 114344, 0.125: 52039},
    "carp_38M": {0.5: 517622, 0.25: 256897, 0.125: 129575},
    "carp_76M": {0.5: 327960, 0.25: 162959, 0.125: 83180},
    "carp_640M": {0.5: 311757, 0.25: 154698, 0.125: 78810},
}

CARP_CHECKPOINT_LOSSES = {
    "carp_600k": {
        1: 2.5051483969586483,
        0.5: 2.5123053303486858,
        0.25: 2.517630364015801,
        0.125: 2.5268538140067247,
    },
    "carp_38M": {
        1: 2.3030167711945997,
        0.5: 2.3189422531432275,
        0.25: 2.338586726549564,
        0.125: 2.3630260306320774,
    },
    "carp_76M": {
        1: 2.2056100474366542,
        0.5: 2.224771098829285,
        0.25: 2.248077081483563,
        0.125: 2.277597215778113,
    },
    "carp_640M": {
        1: 2.0194828284458466,
        0.5: 2.0535276480066376,
        0.25: 2.094229900229448,
        0.125: 2.1455246604919718,
    },
}

CARP_600K = {
    "experiment": "pretrain",
    "task": "mlm",
    "dataset": "uniref50",
    "d_embed": 8,
    "d_model": 128,
    "activation": "gelu",
    "slim": True,
    "epochs": 100,
    "n_layers": 16,
    "kernel_size": 5,
    "r": 128,
    "max_tokens": 600000,
    "max_batch_size": 1600,
    "bucket_size": 10000,
    "opt_level": "O2",
    "lr": 1e-3,
    "warmup_steps": 16000,
    "train_steps": 8000,
}

CARP_38M = {
    "experiment": "pretrain",
    "task": "mlm",
    "dataset": "uniref50",
    "d_embed": 8,
    "d_model": 1024,
    "activation": "gelu",
    "slim": True,
    "epochs": 100,
    "n_layers": 16,
    "kernel_size": 5,
    "r": 128,
    "max_tokens": 40000,
    "max_batch_size": 800,
    "bucket_size": 1000,
    "opt_level": "O2",
    "lr": 1e-3,
    "warmup_steps": 16000,
    "train_steps": 8000,
}

CARP_76M = {
    "experiment": "pretrain",
    "task": "mlm",
    "dataset": "uniref50",
    "d_embed": 8,
    "d_model": 1024,
    "activation": "gelu",
    "slim": True,
    "epochs": 100,
    "n_layers": 32,
    "kernel_size": 5,
    "r": 128,
    "max_tokens": 60000,
    "max_batch_size": 800,
    "bucket_size": 1000,
    "opt_level": "O2",
    "lr": 1e-3,
    "warmup_steps": 16000,
    "train_steps": 8000,
}

CARP_640M = {
    "experiment": "pretrain",
    "task": "mlm",
    "dataset": "uniref50",
    "d_embed": 8,
    "d_model": 1280,
    "activation": "gelu",
    "slim": False,
    "epochs": 100,
    "n_layers": 56,
    "kernel_size": 5,
    "r": 128,
    "max_tokens": 10000,
    "max_batch_size": 800,
    "bucket_size": 1000,
    "opt_level": "O2",
    "lr": 4e-4,
    "warmup_steps": 16000,
    "train_steps": 8000,
}
```
