## Supplemental data and code for "Feature Reuse and Scaling: Understanding Transfer Learning with Protein Language Models": sys.py

```
"""Parameters for training and testing"""

import numpy as np
import torch

# Set up cuda variables
DEVICE = "cuda" if torch.cuda.is_available() else "cpu"

NUM_WORKERS = 8

RAND_SEED = 42

ALPHA_MAG_LOW = -3
ALPHA_MAG_HIGH = 1
SKLEARN_ALPHAS = np.logspace(
    ALPHA_MAG_LOW, ALPHA_MAG_HIGH, ALPHA_MAG_HIGH - ALPHA_MAG_LOW + 1
)

DEFAULT_SPLIT = ["train", "val", "test"]
```
