## Supplemental data and code for "Feature Reuse and Scaling: Understanding Transfer Learning with Protein Language Models": vis.py

```
"""Parameters for plotting"""

from __future__ import annotations

from copy import deepcopy

import numpy as np

import seaborn as sns

from scr.params.emb import CARP_INFO

# allowed dimension reduction types
ALLOWED_DIM_RED_TYPES = ["pca", "tsne", "umap"]
PLOT_EXTS = [".png", ".svg"]

PRESENTATION_PALETTE_SATURATE_DICT = {
    "blue": "#4bacc6",
    "orange": "#f79646",
    "green": "#9bbb59",
    "purple": "#8064a2",
    "gray": "#666666",
}

# blue, orange, green, yellow, purple, gray
PRESENTATION_PALETTE_SATURATE6 = [
    "#4bacc6",
    "#f79646ff",
    "#9bbb59",
    "#f9be00",
    "#8064a2",
    "#666666",
]

TASK_COLORS = (
    sns.dark_palette(PRESENTATION_PALETTE_SATURATE6[1], 4).as_hex()[1:]
    + sns.dark_palette(PRESENTATION_PALETTE_SATURATE6[0], 3).as_hex()[1:]
    + [PRESENTATION_PALETTE_SATURATE6[3], PRESENTATION_PALETTE_SATURATE6[4]]
    + sns.dark_palette(PRESENTATION_PALETTE_SATURATE6[2], 4).as_hex()[1:]
)

# blue, orange, green, purple, gray
PRESENTATION_PALETTE_SATURATE = list(PRESENTATION_PALETTE_SATURATE_DICT.keys())

# lgihter organes: 45%, 30%, 15%
# from https://www.w3schools.com/colors/colors_picker.asp
CHECKPOINT_COLOR = {0.5: "#dc6809", 0.25: "#934506", 0.125: "#492303"}

CARP_ALPHA = {c: a for (c,a) in zip(deepcopy(CARP_INFO).keys(), np.linspace(0.25, 1, 4))}

# note that "structure_ss3_tape_processed" is not considered
# as only for train and val

# the order for plotting legend
ORDERED_TASK_LIST = [
    "proeng_gb1_sampled",
    "proeng_gb1_low_vs_high",
    "proeng_gb1_two_vs_rest",
    "proeng_aav_two_vs_many",
    "proeng_aav_one_vs_many",
    "proeng_thermo_mixed_split",
    "annotation_scl_balanced",
    "structure_ss3_casp12",
    "structure_ss3_cb513",
    "structure_ss3_ts115",
]

# assume same order as the above
ORDERED_TASK_LIST_SIMPLE = [
    "GB1 - sampled",
    "GB1 - low vs high",
    "GB1 - two vs rest",
    "AAV - two vs many",
    "AAV - one vs many",
    "Thermostability",
    "Subcellular localization",
    "SS3 - CASP12",
    "SS3 - CB513",
    "SS3 - TS115",
]

# assume same order
TASK_LEGEND_MAP = {k: v for k, v in zip(ORDERED_TASK_LIST, ORDERED_TASK_LIST_SIMPLE)}

# assume same order
TASK_SIMPLE_COLOR_MAP = {k: v for k, v in zip(ORDERED_TASK_LIST_SIMPLE, TASK_COLORS)}

# for lines
ARCH_LINE_STYLE_DICT = deepcopy({
    "carp": {"linestyle": "solid", "mec": "none"},
    "esm": {"linestyle": "dotted", "mfc": "none"},
})

# for dots
ARCH_DOT_STYLE_DICT = deepcopy({
    "carp": {"edgecolors": "none", "alpha": 0.8},
    "esm": {"facecolors": "none"},
})

ARCH_AB_DOT_STYLE_DICT = deepcopy({
    "rand": ARCH_DOT_STYLE_DICT["esm"],
    "stat": ARCH_DOT_STYLE_DICT["carp"]
})

# for scatter
ARCH_SCATTER_STYLE_DICT = {
    "carp": "o",
    "esm": "X",
}

LAYER_ALPHAS = [0.1, 0.25, 0.4, 0.65, 1]
```
