## Supplemental data and code for "Feature Reuse and Scaling: Understanding Transfer Learning with Protein Language Models": data_process.py

```
"""Pre-processing the dataset"""

from __future__ import annotations

from collections import defaultdict

import os
from glob import glob
import tables
import pandas as pd
import numpy as np

from sklearn.preprocessing import LabelEncoder

import torch
from torch.utils.data import Dataset, DataLoader

from scr.utils import (
    pickle_save,
    pickle_load,
    replace_ext,
    read_std_csv,
    get_folder_file_names,
)
from scr.params.sys import RAND_SEED
from scr.params.emb import TRANSFORMER_INFO, CARP_INFO, MAX_SEQ_LEN
from scr.vis.dataset_vis import DatasetECDF, DatasetStripHistogram
from scr.preprocess.seq_loader import SeqLoader
from scr.encoding.encoding_classes import (
    AbstractEncoder,
    ESMEncoder,
    CARPEncoder,
    OnehotEncoder,
)


def get_mut_name(mut_seq: str, parent_seq: str) -> str:
    """
    A function for returning the mutant name

    Args:
    - mut_seq: str, the full mutant sequence
    - parent_seq: str, the full parent sequence

    Returns:
    - str, parent, indel, or mutant name in the format of
        ParentAAMutLocMutAA:ParentAAMutLocMutAA:..., ie. W39W:D40G:G41C:V54Q
    """

    mut_list = []
    if parent_seq == mut_seq:
        return "parent"
    elif len(parent_seq) == len(mut_seq):
        for i, (p, m) in enumerate(zip(list(parent_seq), list(mut_seq))):
            if p != m:
                mut_list.append(f"{p}{i+1}{m}")
        return ":".join(mut_list)
    else:
        return "indel"


def get_parent_fit(df: pd.DataFrame) -> float:
    """Return the parent fitness value"""

    assert "target" in df.columns, "target is not a column in the dataframe"
    assert "sequence" in df.columns, "sequence is not a column in the dataframe"

    return df["target"][df["mut_name"] == "parent"].values.mean()


class AddMutInfo:
    """A class for appending mutation info for mainly protein engineering tasks"""

    def __init__(self, parent_seq_path: str, csv_path: str):

        """
        Args:
        - parent_seq_path: str, path for the parent sequence
        - csv_path: str, path for the fitness csv file
        """

        # Load the parent sequence from the fasta file
        self._parent_seq = SeqLoader(fasta_loc=parent_seq_path).seq

        # load the dataframe
        self._init_df = read_std_csv(csv_path)
        self._df = self._init_df.copy()

        # add a column with the mutant names
        self._df["mut_name"] = self._init_df["sequence"].apply(
            get_mut_name, parent_seq=self._parent_seq
        )
        # add a column with the number of mutations
        self._df["mut_numb"] = (
            self._df["mut_name"].str.split(":").map(len, na_action="ignore")
        )

        # check number of parents
        parent_idx = self._df[self._df["mut_name"] == "parent"].index.tolist()
        numb_parent = len(parent_idx)

        if numb_parent < 1:
            print("no parent")
        elif numb_parent > 1:
            print(f"{numb_parent} parents observed")

        # change the mut_numb of parent to 0
        self._df.loc[parent_idx, "mut_numb"] = 0

        # get the pickle file path
        self._pkl_path = replace_ext(input_path=csv_path, ext=".pkl")

        pickle_save(what2save=self._df, where2save=self._pkl_path)

    @property
    def parent_seq(self) -> str:
        """Return the parent sequence"""
        return self._parent_seq

    @property
    def pkl_path(self) -> str:
        """Return the pkl file path for the processed dataframe"""
        return self._pkl_path

    @property
    def df(self) -> pd.DataFrame:
        """Return the processed dataframe"""
        return self._df

    @property
    def parent_fit(self) -> float:
        """Return the parent fitness value"""
        return get_parent_fit(self._df)


def std_split_ssdf(
    ssdf_path: str = "data/structure/ss3/tape.csv",
    split_test: bool = True,
) -> None:
    """
    A function that standardize secondary structure dataset
    to set up as columns as sequence, target, set, validation
    where set is train or test and add validation as true
    """
    folder_path = os.path.dirname(ssdf_path)

    df = pd.read_csv(ssdf_path)

    # add validation column
    df["validation"] = df["split"].apply(lambda x: True if x == "valid" else "")
    # now replace valid to train
    df = df.replace("valid", "train")
    # rename all columns
    df.columns = ["sequence", "target", "set", "validation"]

    if split_test:
        # get all kinds of test sets
        ss_tests = set(df["set"].unique()) - set(["train"])

        for ss_test in ss_tests:
            df.loc[~df["set"].isin(set(ss_tests) - set([ss_test]))].replace(
                ss_test, "test"
            ).to_csv(os.path.join(folder_path, ss_test + ".csv"), index=False)

    else:
        df.to_csv(f"{os.path.splitext(ssdf_path)[0]}_processed.csv", index=False)


def split_df_sets(df: pd.DataFrame) -> dict[pd.DataFrame]:
    """
    Return split dataframe for training, validation, and testing

    Args:
    - df: pd.DataFrame, input dataframe

    Returns:
    - a dict of dataframes for train, val, test (or ss3 tasks)
    """

    assert "set" in df.columns, f"set is not a column in the dataframe"
    assert "validation" in df.columns, f"validation is not a column in the dataframe"

    # init split df dict output
    df_dict = {}

    df_dict["train"] = df.loc[
        (df["set"] == "train") & (df["validation"] != True)
    ].reset_index(drop=True)
    df_dict["val"] = df.loc[
        (df["set"] == "train") & (df["validation"] == True)
    ].reset_index(drop=True)

    test_tasks = set(df["set"].unique()) - set(["train"])

    for test_task in test_tasks:
        df_dict[test_task] = df.loc[(df["set"] == test_task)].reset_index(drop=True)

    return df_dict


class DatasetInfo:
    """
    A class returns the information of a dataset
    """

    def __init__(self, dataset_path: str) -> None:
        """
        Args:
        - dataset_path: str, the path for the csv
        """

        self._df = read_std_csv(dataset_path)
        self._df_dict = split_df_sets(self._df)

    def get_model_type(self) -> str:
        # pick linear regression if y numerical
        if self._df.target.dtype.kind in "iufc":
            return "LinearRegression"
        else:
            # ss3
            if "[" in self._df.target[0]:
                return "LinearClassifier-Structure"
            # annotation
            else:
                return "LinearClassifier-Annotation"

    def get_numb_class(self) -> int:
        """
        A function to get number of class
        """
        # annotation class number
        if self.model_type == "LinearClassifier-Annotation":
            return self._df.target.nunique()
        # ss3 or ss8 secondary structure states plus padding
        elif self.model_type == "LinearClassifier-Structure":
            # ss3 secondary structure states WITHOUT padding
            return len(np.unique(np.array(self._df["target"][0][1:-1].split(", "))))
        else:
            return np.nan

    @property
    def df_dict(self) -> dict:
        """Return split dataset based on train, val, test"""
        return self._df_dict

    @property
    def train_numb(self) -> int:
        """Number of train data"""
        return len(self._df_dict["train"])

    @property
    def val_numb(self) -> int:
        """Number of val data"""
        return len(self._df_dict["val"])

    @property
    def test_numb(self) -> int:
        """Number of test data"""
        return len(self._df_dict["test"])

    @property
    def model_type(self) -> str:
        """Return the pytorch model type"""
        return self.get_model_type()

    @property
    def numb_class(self) -> int:
        """Return number of classes for classification"""
        return self.get_numb_class()

    @property
    def subset_list(self) -> list[str]:
        """Return a list of subset"""
        subset_list = list(self._df["set"].unique())
        subset_list.insert(1, "val")
        return subset_list


class TaskProcess:
    """A class for handling different downstream tasks"""

    def __init__(
        self,
        data_folder: str = "data/",
        forceregen: bool = False,
        showplot: bool = False,
    ):
        """
        Args:
        - data_folder: str, a folder path with all the tasks as subfolders where
            all the subfolders have datasets as the subsubfolders, ie

            {data_folder}/
                proeng/
                    aav/
                        one_vs_many.csv
                        two_vs_many.csv
                        P03135.fasta
                    thermo/
                        mixed.csv
        - forceregen: bool = False, if force regenerate the pkl files
        """

        self._data_folder = os.path.normpath(data_folder) + "/"
        self._forceregen = forceregen
        self._showplot = showplot

        # save the sumamarize for all files in the data folder
        self.sum_file_df.to_csv(self.sum_file_df_path)


    def _sum_files(self) -> pd.DataFrame:
        """
        Summarize all files in the data folder

        Returns:
        - A dataframe with "task", "dataset", "split",
            "csv_path", "fasta_path", "pkl_path" as columns, ie.
            (proeng, gb1, low_vs_high, data/proeng/gb1/low_vs_high.csv,
            data/proeng/gb1/5LDE_1.fasta)
            note that csv_path is the list of lmdb files for the structure task
        """
        dataset_folders = glob(f"{self._data_folder}*/*")
        # need a list of tuples in the order of:
        # (task, dataset, split, csv_path, fasta_path)
        list_for_df = []
        for dataset_folder in dataset_folders:
            _, task, dataset = dataset_folder.split("/")

            csv_paths = glob(f"{dataset_folder}/*.csv")

            if task == "structure":
                csv_paths = set(csv_paths) - set(
                    glob(f"{dataset_folder}/tape*.csv")
                )

            fasta_paths = glob(f"{dataset_folder}/*.fasta")
            pkl_paths = glob(f"{dataset_folder}/*.pkl")

            assert len(csv_paths) >= 1, "Less than one csv"
            assert len(fasta_paths) <= 1, "More than one fasta"

            for csv_path in csv_paths:

                # if numerical target
                ds_info = DatasetInfo(dataset_path=csv_path)
                if ds_info.model_type == "LinearRegression":
                    # plot ecdf for each csv file
                    DatasetECDF(dataset_path=csv_path, showplot=self._showplot)

                # if parent seq fasta exists
                if len(fasta_paths) == 1:
                    fasta_path = fasta_paths[0]

                    # if no existing pkl file, generate and save
                    if len(pkl_paths) == 0 or self._forceregen:
                        print(f"Adding mutation info to {csv_path}...")
                        addmutinfo_class = AddMutInfo(
                            parent_seq_path=fasta_path, csv_path=csv_path
                        )
                        pkl_path = addmutinfo_class.pkl_path
                        parent_fit = addmutinfo_class.parent_fit

                    # pkl file exits
                    else:
                        pkl_path = replace_ext(input_path=csv_path, ext=".pkl")
                        parent_fit = get_parent_fit(pickle_load(pkl_path))
                # no parent fasta no pkl file
                else:
                    fasta_path = ""
                    pkl_path = ""
                    parent_fit = np.nan

                df_dict = split_df_sets(read_std_csv(csv_path))

                list_for_df.append(
                    tuple(
                        [
                            task,
                            dataset,
                            os.path.basename(os.path.splitext(csv_path)[0]),
                            ds_info.train_numb,
                            ds_info.val_numb,
                            ds_info.test_numb,
                            ds_info.model_type,
                            ds_info.numb_class,
                            parent_fit,
                            csv_path,
                            fasta_path,
                            pkl_path,
                        ]
                    )
                )

        return pd.DataFrame(
            list_for_df,
            columns=[
                "task",
                "dataset",
                "split",
                "train_numb",
                "val_numb",
                "test_numb",
                "model_type",
                "numb_class",
                "parent_fit",
                "csv_path",
                "fasta_path",
                "pkl_path",
            ],
        )

    @property
    def sum_file_df(self) -> pd.DataFrame:
        """A summary table for all files in the data folder"""
        return self._sum_files()
    
    @property
    def sum_file_df_path(self) -> pd.DataFrame:
        """A summary table for all files in the data folder"""
        return f"{self._data_folder}summary.csv"
    

class ProtranDataset(Dataset):

    """A dataset class for processing protein transfer data"""

    def __init__(
        self,
        dataset_path: str,
        subset: str,
        encoder_name: str,
        reset_param: bool = False,
        resample_param: bool = False,
        embed_batch_size: int = 0,
        flatten_emb: bool | str = False,
        embed_folder: str = None,
        embed_layer: int | None = None,
        seq_start_idx: bool | int = False,
        seq_end_idx: bool | int = False,
        if_encode_all: bool = True,
        **encoder_params,
    ):

        """
        Args:
        - dataset_path: str, full path to the dataset, in pkl or panda readable format, ie
            "data/proeng/gb1/low_vs_high.csv"
            columns include: sequence, target, set, validation,
            mut_name (optional), mut_numb (optional)
        - subset: str, train, val, test
        - encoder_name: str, the name of the encoder
        - reset_param: bool = False, if update the full model to xavier_uniform_
        - resample_param: bool = False, if update the full model to xavier_normal_
        - embed_batch_size: int, set to 0 to encode all in a single batch
        - flatten_emb: bool or str, if and how (one of ["max", "mean"]) to flatten the embedding
        - embed_folder: str = None, path to presaved embedding folder, ie
            "embeddings/proeng/gb1/low_vs_high"
            for which then can add the subset to be, ie
            "embeddings/proeng/gb1/low_vs_high/esm1_t6_43M_UR50S/mean/test/embedding.h5"
        - seq_start_idx: bool | int = False, the index for the start of the sequence
        - seq_end_idx: bool | int = False, the index for the end of the sequence
        - if_encode_all: bool = True, if encode full dataset all layers on the fly
        - encoder_params: kwarg, additional parameters for encoding, including checkpoint info
        """

        # with additional info mut_name, mut_numb
        if os.path.splitext(dataset_path)[-1] in [".pkl", ".PKL", ""]:
            self._df = pickle_load(dataset_path)
            self._add_mut_info = True
        # without such info
        else:
            self._df = read_std_csv(dataset_path)
            self._ds_info = DatasetInfo(dataset_path)
            self._model_type = self._ds_info.model_type
            self._numb_class = self._ds_info.numb_class

            self._add_mut_info = False

        self._df_dict = split_df_sets(self._df)

        assert subset in list(
            self._df_dict.keys()
        ), "split can only be 'train', 'val', 'test' or 'cb513', 'ts115', 'casp12'"
        self._subset = subset

        self._subdf_len = len(self._df_dict[self._subset])

        # not specified seq start will be from 0
        if seq_start_idx == False:
            self._seq_start_idx = 0
        else:
            self._seq_start_idx = int(seq_start_idx)
        # not specified seq end will be the full sequence length
        if seq_end_idx == False:
            self._seq_end_idx = -1
            self._max_seq_len = self._df.sequence.str.len().max()
        else:
            self._seq_end_idx = int(seq_end_idx)
            self._max_seq_len = self._seq_end_idx - self._seq_start_idx

        # get unencoded string of input sequence
        # will need to convert data type
        self.sequence = self._get_column_value("sequence")

        self.if_encode_all = if_encode_all
        self._embed_folder = embed_folder
        
        if self._embed_folder is not None:
            # append init info
            if reset_param and "-rand" not in self._embed_folder:
                self._embed_folder = f"{self._embed_folder}-rand"

            if resample_param and "-stat" not in self._embed_folder:
                self._embed_folder = f"{self._embed_folder}-stat"

        self._encoder_name = encoder_name
        self._flatten_emb = flatten_emb

        # convert binary self._flatten_emb to string
        if self._flatten_emb == False:
            self._flatten_emb_name = "noflatten"
        else:
            self._flatten_emb_name = self._flatten_emb

        if "checkpoint" in encoder_params.keys():
            self._checkpoint = encoder_params["checkpoint"]
        else:
            self._checkpoint = 1

        # get the encoder class
        if self._encoder_name in TRANSFORMER_INFO.keys():
            encoder_class = ESMEncoder

        elif self._encoder_name in CARP_INFO.keys():
            encoder_class = CARPEncoder

        else:
            self._encoder_name == "onehot"
            encoder_class = OnehotEncoder
            encoder_params["max_seq_len"] = self._max_seq_len

        # get the encoder
        self._encoder = encoder_class(
            encoder_name=self._encoder_name,
            reset_param=reset_param,
            resample_param=resample_param,
            **encoder_params,
        )
        self._total_emb_layer = self._encoder.max_emb_layer + 1
        self._embed_layer = embed_layer

        # init 
        self._emb_dataset_folder = ""
        self._emb_table_path = ""

        print(f"self.if_encode_all: {self.if_encode_all}")
        print(f"self._embed_folder: {self._embed_folder}")
        print(f"self._embed_layer: {self._embed_layer}")

        # encode all and load in memory
        if self.if_encode_all or (
            self._embed_folder is None and self._embed_layer is None
        ):
            print("Encoding all...")
            # encode the sequences without the mut_name
            # init an empty dict with empty list to append emb
            encoded_dict = defaultdict(list)

            # use the encoder generator for batch emb
            # assume no labels included
            for encoded_batch_dict in self._encoder.encode(
                mut_seqs=self.sequence,
                batch_size=embed_batch_size,
                flatten_emb=self._flatten_emb,
            ):

                for layer, emb in encoded_batch_dict.items():
                    encoded_dict[layer].append(emb)
            print(encoded_dict.keys())
            # assign each layer as its own variable
            for layer, emb in encoded_dict.items():
                setattr(self, "layer" + str(layer), np.vstack(emb))

        # pre gen all emb in batches
        if not (os.path.exists(self._emb_table_path)):
            print(
                f"{self._emb_table_path} not exist. Need to pre-encoding all in batches..."
            )

        # load from pre saved emb
        if self._embed_folder is not None:

            # get emb folder and path for specific dataset
            self._emb_dataset_folder, _ = get_folder_file_names(
                parent_folder=self._embed_folder,
                dataset_path=dataset_path,
                encoder_name=self._encoder_name,
                embed_layer=0,
                flatten_emb=self._flatten_emb_name,
            )

            self._emb_table_path = os.path.join(
                os.path.join(self._emb_dataset_folder, subset),
                "embedding.h5",
            )
            print(f"self._emb_table_path: {self._emb_table_path}")

            # append emb info
            if self._checkpoint != 1 and "_0." not in self._embed_folder:
                self._embed_folder += f"-{str(self._checkpoint)}"

            """
            dataset_folder, _ = get_folder_file_names(
                parent_folder=self._embed_folder,
                dataset_path=dataset_path,
                encoder_name=self._encoder_name,
                embed_layer=0,
                flatten_emb=self._flatten_emb_name,
            )

            self._emb_table_path = os.path.join(
                        os.path.join(dataset_folder, subset),
                        "embedding.h5",
                    )
            """

            # return all
            if self._embed_layer is None:

                print(f"Load all layers from {self._emb_table_path}...")

                emb_table = tables.open_file(self._emb_table_path)
                emb_table.flush()

                for layer in range(self._total_emb_layer):
                    setattr(
                        self,
                        "layer" + str(layer),
                        getattr(emb_table.root, "layer" + str(layer))[:],
                    )

                emb_table.close()

            # load full one layer embedding
            else:

                print(f"Load {self._embed_layer} from {self._emb_table_path}...")

                emb_table = tables.open_file(self._emb_table_path)
                emb_table.flush()

                setattr(
                    self,
                    "layer" + str(self._embed_layer),
                    getattr(emb_table.root, "layer" + str(self._embed_layer))[:],
                )

                emb_table.close()
        # get and format the fitness or secondary structure values
        # can be numbers or string
        # will need to convert data type
        # make 1D tensor 2D
        self.y = np.expand_dims(self._get_column_value("target"), 1)

        # add mut_name and mut_numb for relevant proeng datasets
        if self._add_mut_info:
            self.mut_name = self._get_column_value("mut_name")
            self.mut_numb = self._get_column_value("mut_numb")
        else:
            self.mut_name = [""] * self._subdf_len
            self.mut_numb = [np.nan] * self._subdf_len

    def __len__(self):
        """Return the length of the selected subset of the dataframe"""
        return self._subdf_len

    def __getitem__(self, idx: int):

        """
        Return the item in the order of
        target (y), sequence, mut_name (optional), mut_numb (optional),
        embedding per layer upto the max number of layer for the encoder

        Args:
        - idx: int
        """
        if self.if_encode_all and self._embed_folder is None:
            return (
                self.y[idx],
                self.sequence[idx],
                self.mut_name[idx],
                self.mut_numb[idx],
                *(
                    getattr(self, "layer" + str(layer))[idx]
                    for layer in range(self._total_emb_layer)
                ),
            )
        elif self._embed_folder is not None:
            # load the .h5 file with the embeddings
            """
            gb1_emb = tables.open_file("embeddings/proeng/gb1/low_vs_high/esm1_t6_43M_UR50S/mean/test/embedding.h5")
            gb1_emb.flush()
            gb1_emb.root.layer0[0:5]
            """
            # append emb info
            if self._checkpoint != 1 and "_0." not in self._embed_folder:
                self._embed_folder += f"-{str(self._checkpoint)}"

            # return all
            if self._embed_layer is None:

                emb_table = tables.open_file(self._emb_table_path)

                emb_table.flush()

                layer_embs = [
                    getattr(emb_table.root, "layer" + str(layer))[idx]
                    for layer in range(self._total_emb_layer)
                ]

                emb_table.close()

                return (
                    self.y[idx],
                    self.sequence[idx],
                    self.mut_name[idx],
                    self.mut_numb[idx],
                    layer_embs,
                )
            # only pick particular embeding layer
            else:

                return (
                    self.y[idx],
                    self.sequence[idx],
                    self.mut_name[idx],
                    self.mut_numb[idx],
                    getattr(self, "layer" + str(self._embed_layer))[idx],
                )
        else:
            return (
                self.y[idx],
                self.sequence[idx],
                self.mut_name[idx],
                self.mut_numb[idx],
            )

    def _get_column_value(self, column_name: str) -> np.ndarray:
        """
        Check and return the column values of the selected dataframe subset

        Args:
        - column_name: str, the name of the dataframe column
        """
        if column_name in self._df.columns:

            y = self._df_dict[self._subset][column_name]

            if column_name == "sequence":

                return (
                    y.astype(str)
                    .str[self._seq_start_idx : self._seq_end_idx]
                    .apply(
                        lambda x: x[: int(MAX_SEQ_LEN // 2)]
                        + x[-int(MAX_SEQ_LEN // 2) :]
                        if len(x) > MAX_SEQ_LEN
                        else x
                    )
                    .values
                )
            elif self._model_type == "LinearClassifier-Annotation":
                print("Converting classes into int...")
                le = LabelEncoder()
                return le.fit_transform(y.values.flatten())
            elif self._model_type == "LinearClassifier-Structure":
                print("Converting ss3/ss8 into np.array and pad -1...")
                np_y = y.apply(lambda x: np.array(x[1:-1].split(", ")).astype("int"))
                return np.stack(
                    [
                        np.pad(
                            i,
                            pad_width=(0, self._max_seq_len - len(i)),
                            constant_values=-1,
                        )
                        for i in np_y
                    ]
                )
            else:
                return y.values

    @property
    def df_full(self) -> pd.DataFrame:
        """Return the full loaded dataset"""
        return self._df

    @property
    def df_train(self) -> pd.DataFrame:
        """Return the dataset for training only"""
        return self._df_dict["train"]

    @property
    def df_val(self) -> pd.DataFrame:
        """Return the dataset for validation only"""
        return self._df_dict["val"]

    @property
    def df_dict(self) -> pd.DataFrame:
        """Return the dict with different dataframe split"""
        return self._df_dict

    @property
    def max_seq_len(self) -> int:
        """Longest sequence length"""
        return self._max_seq_len

    @property
    def emb_table_path(self) -> str:
        """The full path for the emb table for dataset subset"""
        return self._emb_table_path

    @property
    def emb_dataset_folder(self) -> str:
        """The emb folder path for specific dataset"""
        return self._emb_dataset_folder


def split_protrain_loader(
    dataset_path: str,
    encoder_name: str,
    reset_param: bool = False,
    resample_param: bool = False,
    embed_batch_size: int = 128,
    flatten_emb: bool | str = False,
    embed_folder: str | None = None,
    embed_layer: int | None = None,
    seq_start_idx: bool | int = False,
    seq_end_idx: bool | int = False,
    subset_list: list[str] = ["train", "val", "test"],
    loader_batch_size: int = 64,
    worker_seed: int = RAND_SEED,
    if_encode_all: bool = True,
    **encoder_params,
):

    """
    A function encode and load the data from a path

    Args:
    - dataset_path: str, full path to the dataset, in pkl or panda readable format, ie
        "data/proeng/gb1/low_vs_high.csv"
        columns include: sequence, target, set, validation,
        mut_name (optional), mut_numb (optional)
    - encoder_name: str, the name of the encoder
    - reset_param: bool = False, if update the full model to xavier_uniform_
    - resample_param: bool = False, if update the full model to xavier_normal_
    - embed_batch_size: int, set to 0 to encode all in a single batch
    - flatten_emb: bool or str, if and how (one of ["max", "mean"]) to flatten the embedding
    - embed_folder: str = None, path to presaved embedding
    - seq_start_idx: bool | int = False, the index for the start of the sequence
    - seq_end_idx: bool | int = False, the index for the end of the sequence
    - subset_list: list of str, train, val, test
    - loader_batch_size: int, the batch size for train, val, and test dataloader
    - worker_seed: int, the seed for dataloader
    - if_encode_all: bool = True, if encode full dataset all layers on the fly
    - encoder_params: kwarg, additional parameters for encoding, including checkpoint info
    """

    assert set(subset_list) <= set(
        ["train", "val", "test", "cb513", "ts115", "casp12"]
    ), "subset_list can only contain 'train', 'val', 'test', or 'cb513', 'ts115', 'casp12'"

    # specify no shuffling for validation and test
    if_shuffle_list = [True if subset == "train" else False for subset in subset_list]

    return {
        subset: DataLoader(
            dataset=ProtranDataset(
                dataset_path=dataset_path,
                subset=subset,
                encoder_name=encoder_name,
                reset_param=reset_param,
                resample_param=resample_param,
                embed_batch_size=embed_batch_size,
                flatten_emb=flatten_emb,
                embed_folder=embed_folder,
                embed_layer=embed_layer,
                seq_start_idx=seq_start_idx,
                seq_end_idx=seq_end_idx,
                if_encode_all=if_encode_all,
                **encoder_params,
            ),
            batch_size=loader_batch_size,
            shuffle=if_shuffle,
            worker_init_fn=worker_seed,
        )
        for subset, if_shuffle in zip(subset_list, if_shuffle_list)
    }
```
