## Supplemental data and code for "Feature Reuse and Scaling: Understanding Transfer Learning with Protein Language Models": seq_loader.py

```
"""Load seuqences"""

from __future__ import annotations

from abc import ABC

from Bio import SeqIO

from scr.params.aa import ALLOWED_AAS


class SeqLoader(ABC):
    """Class for loading sequences."""

    def __init__(self, fasta_loc: str = "", fasta_seq: str = ""):

        assert fasta_loc != "" or fasta_seq != "", "Need either fasta loc or sequence"

        # load the reference sequence from fasta file or direclty
        self._fasta_loc = fasta_loc

        if fasta_seq != "":
            self._seq = fasta_seq
            if fasta_loc != "":
                print("Both fasta loc and seq are passed in. Take the seq.")
        # only location
        else:
            self._seq = self.load_seq()

        seq_aa_set = set(list(self._seq))

        assert (
            set(list(self._seq)) <= ALLOWED_AAS
        ), f"Sequence contains non-canonical amino acids {ALLOWED_AAS-seq_aa_set}"

    def load_seq(self) -> str:
        """
        Load one fasta sequence

        Returns:
        - str, the full parent sequence in the fasta file
        """

        with open(self._fasta_loc) as handle:
            recs = []
            for rec in SeqIO.parse(handle, "fasta"):
                recs.append(rec)

        # Confirm that there is only one seq file
        assert len(recs) == 1, "More than 1 sequence in the seq file"
        return str(recs[0].seq)

    @property
    def seq(self) -> str:
        """Return the full sequence of the fasta file"""
        return self._seq

    @property
    def seq_len(self) -> int:
        """Return the length of the seqeucne"""
        return len(self._seq)
```
