## Supplemental data and code for "Feature Reuse and Scaling: Understanding Transfer Learning with Protein Language Models": utils.py

```
from __future__ import annotations

# from collections import Sequence

import os
import sys
import pickle

import numpy as np
import pandas as pd

from sklearn.metrics import ndcg_score


def get_default_output_path(envpath : str = "", default_output_path: str = "results/sklearn"):
    """Set default output folder path"""
    if os.getenv(envpath) is None:
        return default_output_path
    else:
        return os.path.join(os.getenv(envpath), default_output_path)


def checkNgen_folder(folder_path: str, envpath: str = "") -> str:
    """
    Check if the folder or the subfolder exists
    to create a new directory if not

    Args:
    - folder_path: str, the folder path
    """
    if os.getenv(envpath) is None:
        split_list = folder_path.split("/")
        for p, _ in enumerate(split_list):
            subfolder_path = "/".join(split_list[: p + 1])
            if not os.path.exists(subfolder_path):
                print(f"Making {subfolder_path} ...")
                os.mkdir(subfolder_path)
        return folder_path

    else:
        _, local_path = folder_path.split(os.getenv(envpath) + "/")
        split_local_list = local_path.split("/")
        for p, _ in enumerate(split_local_list):
            subfolder_path = os.path.join(
                os.getenv(envpath), "/".join(split_local_list[: p + 1])
            )
            if not os.path.exists(subfolder_path):
                print(f"Making {subfolder_path}...")
                os.mkdir(subfolder_path)

        return folder_path


def get_filename(file_path: str) -> str:
    """
    Get the filename without the extension from a full path
    
    Args:
    - file_path: str, the full path for the input file

    Returns:
    - str, the file name without extension
    """
    return os.path.splitext(os.path.basename(file_path))[0]


def replace_ext(input_path: str, ext: str) -> str:

    """
    Replace a file extention of a path to another

    Args:
    - input_path: str, the input path of the file
    - ext: str, the new file extension

    Returns:
    - str, the output path with the new file extension
    """

    if ext[0] == ".":
        return os.path.splitext(input_path)[0] + ext
    else:
        return os.path.splitext(input_path)[0] + "." + ext


def read_std_csv(dataset_path: str) -> pd.DataFrame:
    """
    A function read and check the csv with standard form
    """
    df = pd.read_csv(dataset_path)

    for col in ["sequence", "target", "set", "validation"]:
        assert col in df.columns, f"{col} is not a column in {dataset_path}"

    return df.dropna(subset=["set"])


def get_task_data_split(dataset_path: str,) -> list[str]:
    """
    Return the task, dataset, and the split given the dataset_path is
    data/task/dataset/split.csv, ie. data/proeng/gb1/two_vs_rest.csv
    Args:
    - dataset_path: str, full path to the input dataset, in pkl or panda readable format
        columns include: sequence, target, set, validation, mut_name (optional), mut_numb (optional)
    
    Returns:
    - list[str]: [task, dataset, split]
    """
    return os.path.splitext(dataset_path)[0].split("/")[1:]


def get_folder_file_names(
    parent_folder: str,
    dataset_path: str,
    encoder_name: str,
    embed_layer: int,
    flatten_emb: bool | str,
) -> list[str]:
    """
    A function for specify folder and file names for the output given input dataset

    Args:
    - parent_folder: str, the parent result folder, such as results/train_val_test
    - dataset_path: str, full path to the input dataset, in pkl or panda readable format
        columns include: sequence, target, set, validation, mut_name (optional), mut_numb (optional)
    - encoder_name: str, the name of the encoder
    - embed_layer: int, the layer number of the embedding
    - flatten_emb: bool or str, if and how (one of ["max", "mean"]) to flatten the embedding

    Returns:
    - dataset_subfolder: str, the full path for the dataset based subfolder
    - file_name: str, the name of the file with embedding details without file extension
    """
    # path for the subfolder
    dataset_subfolder = os.path.join(
        parent_folder,
        "/".join(os.path.splitext(dataset_path)[0].split("/")[1:]),
        encoder_name,
        flatten_emb,
    )

    # check and generate the folder
    checkNgen_folder(dataset_subfolder)

    file_name = f"{encoder_name}-{flatten_emb}-layer_{embed_layer}"

    return dataset_subfolder, file_name


def pickle_save(what2save, where2save: str) -> None:

    """
    Save variable to a pickle file

    Args:
    - what2save, the varible that needs to be saved
    - where2save: str, the .pkl path for saving
    """

    with open(where2save, "wb") as f:
        pickle.dump(what2save, f)


def pickle_load(path2load: str):

    """
    Load pickle file

    Args:
    - path2load: str, the .pkl path for loading
    """

    with open(path2load, "rb") as f:
        return pickle.load(f)


def blockPrint():
    """Block printing"""
    sys.stdout = open(os.devnull, "w")


def enablePrint():
    """Restore printing"""
    sys.stdout = sys.__stdout__


def ndcg_scale(true: np.ndarray, pred: np.ndarray):
    """Calculate the ndcg_score with neg correction"""
    if min(true) < 0:
        true = true - min(true)
    return ndcg_score(true[None, :], pred[None, :])
```
