## Supplemental data and code for "Feature Reuse and Scaling: Understanding Transfer Learning with Protein Language Models": dataset_vis.py

```
"""For dataset vis"""

from __future__ import annotations

import os

from glob import glob

import pandas as pd

import matplotlib.pyplot as plt

import iqplot

from scr.params.vis import PLOT_EXTS, PRESENTATION_PALETTE_SATURATE_DICT
from scr.vis.vis_utils import BokehSave
from scr.vis.iqplot_striphis import striphistogram
from scr.utils import get_task_data_split, read_std_csv, pickle_load, checkNgen_folder

class DatasetSeqLenHist:
    """
    A class for plotting dataset length histograms
    """

    def __init__(
        self,
        df_path: str,
        result_subfolder: str = "results/dataset_vis/len_hist",
    ) -> None:

        self._df_path = df_path
        self._result_subfolder = checkNgen_folder(result_subfolder)

        for iflog in [True, False]:
            self._plot_len_hist(iflog=iflog)

    def _plot_len_hist(self, iflog: bool = False):
        """
        plot histogram of seq len with if log option
        """

        plot_title = f"{self.df_dets} seq length histogram"

        if iflog:
            log_det = " log scale"
        else:
            log_det = ""

        len_hist = self.seqlen.hist(bins=self.bin_numb, log=iflog)
        len_hist.set_xlabel("Length")
        len_hist.set_ylabel("Counts")
        len_hist.set_title(plot_title)
        
        plt.grid(False)

        len_hist.get_figure().savefig(
            os.path.join(
                self._result_subfolder, f"{plot_title}{log_det}.png".replace(" ", "_")
            ),
            bbox_inches="tight",
        )

        plt.close()
    

    @property
    def df(self) -> pd.DataFrame:
        """Return the dataframe from the path"""
        return pd.read_csv(self._df_path)

    @property
    def df_dets(self) -> str:
        """Details for the dataset"""
        return os.path.splitext(self._df_path)[0].split("data/")[-1].replace("/", " ")

    @property
    def seqlen(self) -> pd.Series:
        """Get a series for seq len"""
        return self.df.sequence.str.len()

    @property
    def bin_numb(self) -> int:
        """Number of bins for the hist"""
        return int(self.seqlen.max() / 10)


class DatasetECDF(BokehSave):
    def __init__(
        self,
        dataset_path: str,
        path2folder: str = "results/dataset_vis",
        plot_exts: list = PLOT_EXTS,
        plot_height: int = 300,
        plot_width: int = 450,
        axis_font_size: str = "10pt",
        title_font_size: str = "10pt",
        x_name: str = "fitness",
        y_name: str = "ecdf",
        gridoff: bool = True,
        showplot: bool = True
    ) -> None:

        df = read_std_csv(dataset_path)

        df.loc[df["validation"] == True, "set"] = "val"

        self.bokeh_plot = iqplot.ecdf(
            df,
            q="target",
            cats="set",
            conf_int=True,
            # style="staircase",
            palette=[
                PRESENTATION_PALETTE_SATURATE_DICT["blue"],
                PRESENTATION_PALETTE_SATURATE_DICT["green"],
                PRESENTATION_PALETTE_SATURATE_DICT["orange"],
            ],
            order=["train", "val", "test"],
            legend_location="bottom_right",
            marker_kwargs={"alpha": 0.5},
            fill_kwargs={"fill_alpha": 0.1}
            # line_kwargs={"line_width": 2.5},
        )

        super(DatasetECDF, self).__init__(
            bokeh_plot=self.bokeh_plot,
            path2folder=path2folder,
            plot_name="-".join(get_task_data_split(dataset_path)),
            plot_exts=plot_exts,
            plot_height=plot_height,
            plot_width=plot_width,
            axis_font_size=axis_font_size,
            title_font_size=title_font_size,
            x_name=x_name,
            y_name=y_name,
            gridoff=gridoff,
            showplot=showplot
        )


class DatasetStripHistogram(BokehSave):
    def __init__(
        self,
        dataset_folder: str,
        split_order: list[str] | None = None,
        path2folder: str = "results/dataset_vis",
        plot_exts: list = PLOT_EXTS,
        plot_height: int = 400,
        plot_width: int = 600,
        axis_font_size: str = "10pt",
        title_font_size: str = "10pt",
        x_name: str = "",
        y_name: str = "fitness",
        gridoff: bool = True,
        showplot: bool = True
    ) -> None:
        """
        Args:
        - dataset_folder: str, ie. data/proeng/gb1
        - split_order: list[str], ie. ["low_vs_high", "two_vs_rest", "sampled"]
        """
        self._dataset_folder = os.path.normpath(dataset_folder)
        self._dataset_paths = glob(f"{self._dataset_folder}/*.pkl")

        assert "proeng" in self._dataset_folder, "only support proeng datasets"

        if len(self._dataset_paths) == 0:
            self._dataset_paths = glob(f"{self._dataset_folder }/*.csv")
            self._cat_dfs = read_std_csv(
                glob(f"{self._dataset_folder }/*.csv")[0]
            )
            self._cat_dfs.loc[self._cat_dfs["validation"] == True, "set"] = "val"
        else:
            dfs = []
            for pkl in self._dataset_paths:
                task, data, split = get_task_data_split(pkl)
                df = pickle_load(pkl)
                df["split"] = split
                df.loc[df["validation"] == True, "set"] = "val"
                dfs.append(df)

            self._cat_dfs = pd.concat(dfs, ignore_index=True, axis=0)

        set_order = ["train", "val", "test"]

        if split_order is None:
            cat_orders = set_order
            cats_list = ["set"]
        else:
            cat_orders = [(i, j) for i in split_order for j in set_order]
            cats_list = ["split", "set"]

        self.bokeh_plot = striphistogram(
            self._cat_dfs,
            q="target",
            cats=cats_list,
            spread="jitter",
            # jitter=True,
            color_column="set",
            palette=[
                PRESENTATION_PALETTE_SATURATE_DICT["orange"],
                PRESENTATION_PALETTE_SATURATE_DICT["blue"],
                PRESENTATION_PALETTE_SATURATE_DICT["green"],
            ],
            top_level="histogram",
            marker_kwargs={"alpha": 0.1},
            fill_kwargs={"fill_alpha": 0.1},
            order=cat_orders,
            # spread_kwargs={'distribution': 'normal', 'width': 0.1},
            q_axis="y",
        )

        super(DatasetStripHistogram, self).__init__(
            bokeh_plot=self.bokeh_plot,
            path2folder=path2folder,
            plot_name="-".join(get_task_data_split(self._dataset_folder)[:2]),
            plot_exts=plot_exts,
            plot_height=plot_height,
            plot_width=plot_width,
            axis_font_size=axis_font_size,
            title_font_size=title_font_size,
            x_name=x_name,
            y_name=y_name,
            gridoff=gridoff,
            showplot=showplot
        )
```
