## Supplemental data and code for "Feature Reuse and Scaling: Understanding Transfer Learning with Protein Language Models": emb_vis.py

```
"""Visulization for the embeddings"""

from __future__ import annotations

from collections import Sequence

from scr.params.aa import AA_PROP_DF, ALL_AAS

import os
import matplotlib.pyplot as plt

import numpy as np

import anndata as ad
import scanpy as sc

from scr.params.vis import ALLOWED_DIM_RED_TYPES, PLOT_EXTS

"""
# allowed dimension reduction types
ALLOWED_DIM_RED_TYPES = ["pca", "tsne", "umap"]
DEFAULT_FORMATS = [".png", ".svg"]
"""

class Emb_Vis:
    """Class for embbeding visulization"""

    def __init__(
        self,
        aa_emb: np.ndarray,
        emb_name: str,
        dim_red_types: Sequence(str) = ALLOWED_DIM_RED_TYPES,
        folder_path: str = "results/emb_vis",
        subfolder: str = "",
        plot_formats: Sequence(str) = PLOT_EXTS,
    ) -> None:
        """
        Args
        - aa_emb: np.ndarray, (number of aa, embedding dim), amino acid embeddings
        - emb_name: str, name of the embedding, ie. esm1b_t33_650M_UR50S input amino acids
        - dim_red_type: Sequence(str), types of dimension reduction
        - folder_path: str, folder path for embedding plots
        - subfolder: str, name of the subfolder, ie. esm
        - plot_formats: Sequence(str), types of plots
        """

        # annotate the embedding with amino acid properties
        aa_emb_adata = ad.AnnData(aa_emb, obs=AA_PROP_DF)

        # reduce embedding dimensions
        sc.pp.pca(aa_emb_adata, n_comps=2)
        sc.pp.neighbors(aa_emb_adata, use_rep="X")
        sc.tl.umap(aa_emb_adata)
        sc.tl.tsne(aa_emb_adata)

        self._emb_name = emb_name

        # check dimension reduction type list
        if set(dim_red_types) > set(ALLOWED_DIM_RED_TYPES):
            print(
                f"Including not supported dim_red_types, set to {ALLOWED_DIM_RED_TYPES}"
            )
            self._dim_red_types = ALLOWED_DIM_RED_TYPES
        else:
            self._dim_red_types = dim_red_types

        # create the path for saving the plots
        self._subfolder_path = os.path.join(folder_path, subfolder)
        self._plot_formats = plot_formats

        for dim_red_type in self._dim_red_types:
            self.save_emb_vis(aa_emb_adata=aa_emb_adata, dim_red_type=dim_red_type)

    def save_emb_vis(self, aa_emb_adata: ad.AnnData, dim_red_type: str) -> None:
        """
        Generate and save the embedding visulization plots

        Args:
        - aa_emb_adata: ad.AnaData, annotated embedding after dim red
        - dim_red_type: str, dimension reduction type, ie. pca
        """

        fig, ax = plt.subplots(dpi=300)
        ax = getattr(sc.pl, dim_red_type)(
            aa_emb_adata, color="property", size=250, ax=ax, show=False
        )
        obsm_name = "X_" + dim_red_type

        for i, aa in enumerate(ALL_AAS):
            ax.annotate(
                aa,
                (
                    aa_emb_adata.obsm[obsm_name][i, 0],
                    aa_emb_adata.obsm[obsm_name][i, 1],
                ),
            )
        ax.set(
            title=f"{self._emb_name} embeddings",
        )
        fig.show()

        mod_emb_name = self._emb_name.replace(" ", "_")

        # save plots for given formats
        for plot_format in self._plot_formats:
            # check if the suffix include the dot
            if plot_format[0] != ".":
                plot_format = "." + plot_format
            plt.savefig(
                os.path.join(
                    self._subfolder_path, f"{mod_emb_name}_{dim_red_type}{plot_format}"
                ),
                bbox_inches="tight"
            )
```
