## Supplemental data and code for "Feature Reuse and Scaling: Understanding Transfer Learning with Protein Language Models": emb_vis_esm.py

```
"""Visulization for the ESM embeddings"""

from __future__ import annotations

from collections import Sequence

import torch

from scr.vis.emb_vis import Emb_Vis
from scr.params.emb import TRANSFORMER_INFO

ESM_MODEL_LIST = list(TRANSFORMER_INFO.keys())

def esm_emb_vis(
    esm_model_list: list(str) = ESM_MODEL_LIST,
    emb_start_ind: int = 0,
    emb_end_ind: int = 20,
) -> list(Emb_Vis):
    """
    Plot and save the esm embedding

    Args:
    - esm_model_list: list(str), a list of esm models
    - emb_start_ind: int, the start index of the esm embedding
    - emb_end_ind: int, the ending index of the esm embedding

    Returns:
    - emb_vis_class_list: list(Emb_Vis), a lsit of Emb_Vis class
    """

    emb_vis_class_list = [None] * len(esm_model_list)

    for m, esm_model in enumerate(esm_model_list):
        model, _ = torch.hub.load("facebookresearch/esm:main", esm_model)
        aa_emb = (
            model.embed_tokens.weight.detach()
            .cpu()
            .numpy()[emb_start_ind:emb_end_ind, :]
        )

        print(f"Plotting and saving {esm_model}")
        emb_vis_class_list[m] = Emb_Vis(
            aa_emb=aa_emb,
            emb_name=f"{esm_model} input amino acids",
            subfolder="esm",
        )

    return emb_vis_class_list
```
