## Supplemental data and code for "Feature Reuse and Scaling: Understanding Transfer Learning with Protein Language Models": iqplot_striphis.py

```
"""Temp to fix iqplot overlay"""

import copy
import warnings

import numpy as np
import pandas as pd
import xarray

import colorcet

import bokeh.models
import bokeh.plotting


def _fig_dimensions(kwargs):
    if (
        "width" not in kwargs
        and "plot_width" not in kwargs
        and "frame_width" not in kwargs
    ):
        kwargs["frame_width"] = 375
    if (
        "height" not in kwargs
        and "plot_height" not in kwargs
        and "frame_height" not in kwargs
    ):
        kwargs["frame_height"] = 275

    if "toolbar_location" not in kwargs:
        kwargs["toolbar_location"] = "above"

    return kwargs


def _parse_deprecations(
    q,
    q_axis,
    val,
    horizontal,
    horiz_q_axis,
    click_policy,
    legend_click_policy,
    conf_int_kwargs,
    fill_kwargs,
):
    if q_axis not in ("x", "y"):
        raise RuntimeError("Invalid `q_axis`. Must by 'x' or 'y'.")

    if horizontal is not None:
        if (horizontal and q_axis != horiz_q_axis) or (
            not horizontal and q_axis == horiz_q_axis
        ):
            raise RuntimeError(
                "`horizontal` and `q_axis` kwargs in disagreement. "
                "Use `q_axis`; `horizontal` is deprecated."
            )

        warnings.warn(f"`horizontal` is deprecated. Use `q_axis`.", DeprecationWarning)

    if val is not None:
        if q is None:
            q = val
        elif q != val:
            raise RuntimeError(
                "`val` and `q` in disagreement. Use `q`; `val` is deprecated."
            )

        warnings.warn(f"`val` is deprecated. Use `q`. Using q={q}.", DeprecationWarning)

    if click_policy is not None:
        if legend_click_policy is None:
            legend_click_policy = click_policy
        elif click_policy != legend_click_policy:
            raise RuntimeError(
                "`click_policy` and `legend_click_policy` in disagreement. Use `legend_click_policy`; `click_policy` is deprecated."
            )

        warnings.warn(
            f"`click_policy` is deprecated. Use `legend_click_policy`. Using legend_click_policy='{legend_click_policy}'."
        )

    if conf_int_kwargs is not None:
        if fill_kwargs is None:
            fill_kwargs = copy.copy(conf_int_kwargs)
            warnings.warn(f"`conf_int_kwargs is deprecated. Use `fill_kwargs`.")
        elif conf_int_kwargs != fill_kwargs:
            raise RuntimeError(
                "`fill_kwargs` and `conf_int_kwargs` in disagreement. Use `fill_kwargs`; `conf_int_kwargs` is deprecated."
            )

        warnings.warn(f"`conf_int_kwargs is deprecated. Use `fill_kwargs`.")

    return q, legend_click_policy, fill_kwargs


def _data_cats(data, q, cats, show_legend, legend_label):
    if type(data) == xarray.core.dataarray.DataArray:
        if q is None:
            if data.name is None:
                q = "x"
            else:
                q = data.name
        data = pd.DataFrame({q: data.squeeze().values})
    elif type(data) == np.ndarray:
        if q is None:
            q = "x"
        data = pd.DataFrame({q: data.squeeze()})
        if cats is not None:
            raise RuntimeError("If `data` is a Numpy array, `cats` must be None.")

    if cats is None:
        data = data.copy()
        if legend_label is None:
            data["__dummy_cat"] = " "
            show_legend = False
        else:
            data["__dummy_cat"] = legend_label
        cats = "__dummy_cat"

    return data, q, cats, show_legend


def _fill_between(p, x1=None, y1=None, x2=None, y2=None, **kwargs):
    """
    Create a filled region between two curves.
    Parameters
    ----------
    p : bokeh.plotting.Figure instance
        Figure to be populated.
    x1 : array_like
        Array of x-values for first curve
    y1 : array_like
        Array of y-values for first curve
    x2 : array_like
        Array of x-values for second curve
    y2 : array_like
        Array of y-values for second curve
    kwargs
        Any kwargs passed to p.patch.
    Returns
    -------
    output : bokeh.plotting.Figure instance
        Plot populated with fill-between.
    """
    x = list(x1) + list(x2)[::-1]
    y = list(y1) + list(y2)[::-1]
    patch = p.patch(x=x, y=y, **kwargs)

    # Old way; only works for Numpy arrays
    # patch = p.patch(
    #     x=np.concatenate((x1, x2[::-1])), y=np.concatenate((y1, y2[::-1])), **kwargs
    # )

    return p, patch


def _get_marker(p, marker):
    if marker.lower() == "asterisk":
        return p.asterisk
    if marker.lower() == "circle":
        return p.circle
    if marker.lower() == "circle_cross":
        return p.circle_cross
    if marker.lower() == "circle_x":
        return p.circle_x
    if marker.lower() == "cross":
        return p.cross
    if marker.lower() == "dash":
        return p.dash
    if marker.lower() == "diamond":
        return p.diamond
    if marker.lower() == "diamond_cross":
        return p.diamond_cross
    if marker.lower() == "hex":
        return p.hex
    if marker.lower() == "inverted_triangle":
        return p.inverted_triangle
    if marker.lower() == "square":
        return p.square
    if marker.lower() == "square_cross":
        return p.square_cross
    if marker.lower() == "square_x":
        return p.square_x
    if marker.lower() == "triangle":
        return p.triangle
    if marker.lower() == "x":
        return p.x

    raise RuntimeError(
        f"{marker} is an invalid marker specification. Acceptable values are ['asterisk', 'circle', 'circlecross', 'circle_x', 'cross', 'dash', 'diamond', 'diamondcross', 'hex', 'invertedtriangle', 'square', 'squarecross', 'squarex', 'triangle', 'x']"
    )


def _source_and_labels_from_cats(df, cats):
    if type(cats) in [list, tuple]:
        cat_source = list(zip(*tuple([df[cat].astype(str) for cat in cats])))
        return cat_source, [", ".join(cat) for cat in cat_source]
    else:
        cat_source = list(df[cats].astype(str).values)
        return cat_source, cat_source


def _tooltip_cols(tooltips):
    if tooltips is None:
        return []
    if type(tooltips) not in [list, tuple]:
        raise RuntimeError("`tooltips` must be a list or tuple of two-tuples.")

    cols = []
    for tip in tooltips:
        if type(tip) not in [list, tuple] or len(tip) != 2:
            raise RuntimeError("Invalid tooltip.")
        if tip[1][0] == "@":
            if tip[1][1] == "{":
                cols.append(tip[1][2 : tip[1].find("}")])
            elif "{" in tip[1]:
                cols.append(tip[1][1 : tip[1].find("{")])
            else:
                cols.append(tip[1][1:])

    return cols


def _cols_to_keep(cats, q, color_column, tooltips):
    cols = _tooltip_cols(tooltips)
    cols += [q]

    if type(cats) in [list, tuple]:
        cols += list(cats)
    else:
        cols += [cats]

    if color_column is not None:
        cols += [color_column]

    return list(set(cols))


def _check_cat_input(
    df, cats, q, color_column, parcoord_column, tooltips, palette, order, kwargs
):
    if df is None:
        raise RuntimeError("`df` argument must be provided.")
    if cats is None:
        raise RuntimeError("`cats` argument must be provided.")
    if q is None:
        raise RuntimeError("`q` argument must be provided.")

    if type(palette) not in [list, tuple, str]:
        raise RuntimeError("`palette` must be a list, tuple or string.")

    if q not in df.columns:
        raise RuntimeError(f"{q} is not a column in the inputted data frame")

    cats_array = type(cats) in [list, tuple]
    if cats_array and len(cats) == 1:
        cats = cats[0]
        cats_array = False

    if cats_array:
        for cat in cats:
            if cat not in df.columns:
                raise RuntimeError(f"{cat} is not a column in the inputted data frame")
    else:
        if type(cats) == tuple:
            raise RuntimeError(
                "Cannot have tuples as data frame column names if there is only one categorical variable."
            )
        if cats not in df.columns:
            raise RuntimeError(f"{cats} is not a column in the inputted data frame")

    if color_column is not None and color_column not in df.columns:
        raise RuntimeError(f"{color_column} is not a column in the inputted data frame")

    cols = _cols_to_keep(cats, q, color_column, tooltips)

    for col in cols:
        if col not in df.columns:
            raise RuntimeError(f"{col} is not a column in the inputted data frame")

    bad_kwargs = ["x", "y", "source", "cat", "legend"]
    if kwargs is not None and any([key in kwargs for key in bad_kwargs]):
        raise RuntimeError(", ".join(bad_kwargs) + " are not allowed kwargs.")

    if q == "cat":
        raise RuntimeError("`'cat'` cannot be used as `q`.")

    if q == "__label" or (cats == "__label" or (cats_array and "__label" in cats)):
        raise RuntimeError("'__label' cannot be used for `q` or `cats`.")

    if order is not None:
        grouped = df.groupby(cats)
        if grouped.ngroups > len(order):
            raise RuntimeError(
                "`order` must have at least as many elements as the number of unique groups in `cats`."
            )
        for entry in order:
            if entry not in grouped.groups.keys():
                raise RuntimeError(
                    f"Entry {entry} in grouping of input data but not present in as a group in the inputted data."
                )

    if parcoord_column is not None:
        if parcoord_column not in df.columns:
            raise RuntimeError(
                f"{parcoord_column} is not a column in the inputted data frame"
            )
        if cats == "__dummy_cat":
            raise RuntimeError(
                "`cats` must be provided in `parcoord_column` is provided."
            )
        grouped = df.groupby(parcoord_column)

    return cats, cols


def _convert_data(data, inf_ok=False, min_len=1):
    """
    Convert inputted 1D data set into NumPy array of floats.
    All nan's are dropped.
    Parameters
    ----------
    data : int, float, or array_like
        Input data, to be converted.
    inf_ok : bool, default False
        If True, np.inf values are allowed in the arrays.
    min_len : int, default 1
        Minimum length of array.
    Returns
    -------
    output : ndarray
        `data` as a one-dimensional NumPy array, dtype float.
    """
    # If it's scalar, convert to array
    if np.isscalar(data):
        data = np.array([data], dtype=np.float64)

    # Convert data to NumPy array
    data = np.array(data, dtype=np.float64)

    # Make sure it is 1D
    if len(data.shape) != 1:
        raise RuntimeError("Input must be a 1D array or Pandas series.")

    # Remove NaNs
    data = data[~np.isnan(data)]

    # Check for infinite entries
    if not inf_ok and np.isinf(data).any():
        raise RuntimeError("All entries must be finite.")

    # Check to minimal length
    if len(data) < min_len:
        raise RuntimeError(
            "Array must have at least {0:d} non-NaN entries.".format(min_len)
        )

    return data


def _sort_df(df, cats, order):
    if order is not None and cats is not None:
        if type(cats) in [list, tuple]:
            df["__sort"] = df.apply(lambda r: order.index(tuple(r[cats])), axis=1)
        else:
            df["__sort"] = df.apply(lambda r: order.index(r[cats]), axis=1)
        df = df.sort_values(by="__sort")

    return df

def _stacked_histograms(
    df,
    grouped,
    q,
    bins,
    density,
    palette,
    q_axis,
    order,
    p,
    mirror,
    hist_height,
    style,
    conf_int,
    ptiles,
    n_bs_reps,
    rug,
    rug_height,
    tooltips,
    line_kwargs,
    fill_kwargs,
    rug_kwargs,
    kwargs,
):
    # Protect against mutability and get copies
    line_kwargs = copy.copy(line_kwargs)
    fill_kwargs = copy.copy(fill_kwargs)

    line_line_color_supplied = "line_color" in line_kwargs
    fill_fill_color_supplied = "fill_color" in fill_kwargs

    if p is None:
        p, _, _ = cat._cat_figure(df, grouped, q, order, None, q_axis, kwargs)

    for i, (name, g) in enumerate(grouped):
        if not fill_fill_color_supplied:
            fill_kwargs["fill_color"] = palette[i % len(palette)]
        if not line_line_color_supplied:
            line_kwargs["line_color"] = palette[i % len(palette)]

        numerical_bins, e, f = _compute_histogram(g[q].values, bins, density)
        e0, f0 = _hist_for_plotting(e, f)

        if conf_int:
            q_vals = g[q].values
            f_reps = np.empty((n_bs_reps, len(f)))
            for j in range(n_bs_reps):
                _, _, f_reps[j, :] = _compute_histogram(
                    np.random.choice(q_vals, len(q_vals)), numerical_bins, density
                )

            f_ptiles = np.percentile(f_reps, [2.5, 97.5], axis=0)

            _, f0_low = _hist_for_plotting(e, f_ptiles[0, :])
            _, f0_high = _hist_for_plotting(e, f_ptiles[1, :])

            # Scale to fit nicely in categorical axes
            scale = 1.0 / np.max(f0_high) * hist_height / 2

            f0_low_cat = [
                (*name, f0_val) if type(name) == tuple else (name, f0_val)
                for f0_val in scale * f0_low
            ]
            f0_high_cat = [
                (*name, f0_val) if type(name) == tuple else (name, f0_val)
                for f0_val in scale * f0_high
            ]

            if q_axis == "y":
                p, patch = utils._fill_between(
                    p, f0_low_cat, e0, f0_high_cat, e0, **fill_kwargs
                )
            else:
                p, patch = utils._fill_between(
                    p, e0, f0_low_cat, e0, f0_high_cat, **fill_kwargs
                )

            if mirror:
                f0_low_cat = [
                    (*name, f0_val) if type(name) == tuple else (name, f0_val)
                    for f0_val in -scale * f0_low
                ]
                f0_high_cat = [
                    (*name, f0_val) if type(name) == tuple else (name, f0_val)
                    for f0_val in -scale * f0_high
                ]
                if q_axis == "y":
                    p, patch = utils._fill_between(
                        p, f0_low_cat, e0, f0_high_cat, e0, **fill_kwargs
                    )
                else:
                    p, patch = utils._fill_between(
                        p, e0, f0_low_cat, e0, f0_high_cat, **fill_kwargs
                    )
        else:
            scale = 1.0 / np.max(f0) * hist_height / 2

        # y-values for histogram, appropriately scaled
        f0 *= scale
        f0_cat = [
            (*name, f0_val) if type(name) == tuple else (name, f0_val) for f0_val in f0
        ]

        if mirror:
            f0_cat += list(
                reversed(
                    [
                        (*name, -f0_val) if type(name) == tuple else (name, -f0_val)
                        for f0_val in f0
                    ]
                )
            )
            e0 = np.concatenate((e0, e0[::-1]))

        # Line of histogram
        if q_axis == "y":
            p.line(f0_cat, e0, **line_kwargs)
            if style == "step_filled":
                p.patch(f0_cat, e0, **fill_kwargs)
        else:
            p.line(e0, f0_cat, **line_kwargs)
            if style == "step_filled":
                p.patch(e0, f0_cat, **fill_kwargs)

        # Add rug, only if not mirrored. Otherwise, use striphistogram().
        if rug:
            for i, (name, g) in enumerate(grouped):
                xs = [[x, x] for x in g[q]]
                y0_cat = [
                    (*name, rug_height * hist_height / 2)
                    if type(name) == tuple
                    else (name, rug_height * hist_height / 2)
                    for _ in range(len(g))
                ]

                if mirror:
                    ys = [(y0, (y0[0], -y0[1])) for y0 in y0_cat]
                else:
                    ys = [(y0, (y0[0], 0)) for y0 in y0_cat]

                cds = bokeh.models.ColumnDataSource(g)
                cds.data["__xs"] = xs
                cds.data["__ys"] = ys

                if "color" not in rug_kwargs and "line_color" not in rug_kwargs:
                    p.multi_line(
                        source=cds,
                        xs="__xs",
                        ys="__ys",
                        line_color=palette[i % len(palette)],
                        **rug_kwargs,
                    )
                else:
                    p.multi_line(source=cds, xs="__xs", ys="__ys", **rug_kwargs)

    if rug and tooltips is not None:
        p.add_tools(bokeh.models.HoverTool(tooltips=tooltips, names=["hover_glyphs"]))

    return p


def _compute_histogram(data, bins, density):
    """Computes the bins and edges of a histogram."""
    if type(bins) == str and bins == "sqrt":
        bins = int(np.ceil(np.sqrt(len(data))))
    elif type(bins) == str and bins == "freedman-diaconis":
        h = 2 * (np.percentile(data, 75) - np.percentile(data, 25)) / np.cbrt(len(data))
        if h == 0.0:
            bins = 3
        else:
            bins = int(np.ceil((data.max() - data.min()) / h))

    f, e = np.histogram(data, bins=bins, density=density)

    return bins, e, f


def _hist_for_plotting(e, f):
    """Takes output e and f from _compute_histogram(), and generates
    x, y values for plotting the histogram."""
    e0 = np.empty(2 * len(e))
    f0 = np.empty(2 * len(e))
    e0[::2] = e
    e0[1::2] = e
    f0[0] = 0
    f0[-1] = 0
    f0[1:-1:2] = f
    f0[2:-1:2] = f

    return e0, f0

def strip(
    data=None,
    q=None,
    cats=None,
    q_axis="x",
    palette=None,
    order=None,
    p=None,
    show_legend=None,
    legend_location="right",
    legend_orientation="vertical",
    legend_click_policy="hide",
    color_column=None,
    parcoord_column=None,
    tooltips=None,
    marker="circle",
    spread=None,
    cat_grid=False,
    marker_kwargs=None,
    jitter_kwargs=None,
    swarm_kwargs=None,
    parcoord_kwargs=None,
    jitter=None,
    horizontal=None,
    val=None,
    click_policy=None,
    **kwargs,
):
    """
    Make a strip plot.
    Parameters
    ----------
    data : Pandas DataFrame, 1D Numpy array, or xarray
        DataFrame containing tidy data for plotting.  If a Numpy array,
        a single category is assumed and a strip plot generated from
        data.
    q : hashable
        Name of column to use as quantitative variable if `data` is a
        Pandas DataFrame. Otherwise, `q` is used as the quantitative
        axis label.
    cats : hashable or list of hashables
        Name of column(s) to use as categorical variable(s).
    q_axis : str, either 'x' or 'y', default 'x'
        Axis along which the quantitative value varies.
    palette : list of strings of hex colors, or single hex string
        If a list, color palette to use. If a single string representing
        a hex color, all glyphs are colored with that color. Default is
        colorcet.b_glasbey_category10 from the colorcet package.
    order : list or None
        If not None, must be a list of unique group names when the input
        data frame is grouped by `cats`. The order of the list specifies
        the ordering of the categorical variables on the categorical
        axis and legend. If None, the categories appear in the order in
        which they appeared in the inputted data frame.
    p : bokeh.plotting.Figure instance, or None (default)
        If None, create a new figure. Otherwise, populate the existing
        figure `p`.
    show_legend : bool, default False
        If True, display legend.
    legend_location : str, default 'right'
        Location of legend. If one of "right", "left", "above", or
        "below", the legend is placed outside of the plot area. If one
        of "top_left", "top_center", "top_right", "center_right",
        "bottom_right", "bottom_center", "bottom_left", "center_left",
        or "center", the legend is placed within the plot area. If a
        2-tuple, legend is placed according to the coordinates in the
        tuple.
    legend_orientation : str, default 'vertical'
        Either 'horizontal' or 'vertical'.
    legend_click_policy : str, default 'hide'
        Either 'hide', 'mute', or None; how the glyphs respond when the
        corresponding category is clicked in the legend.
    color_column : hashable, default None
        Column of `data` to use in determining color of glyphs. If None,
        then `cats` is used.
    parcoord_column : hashable, default None
        Column of `data` to use to construct a parallel coordinate plot.
        Data points with like entries in the parcoord_column are
        connected with lines.
    tooltips : list of 2-tuples
        Specification for tooltips as per Bokeh specifications. For
        example, if we want `col1` and `col2` tooltips, we can use
        `tooltips=[('label 1': '@col1'), ('label 2': '@col2')]`.
    marker : str, default 'circle'
        Name of marker to be used in the plot (ignored if `formal` is
        False). Must be one of['asterisk', 'circle', 'circle_cross',
        'circle_x', 'cross', 'dash', 'diamond', 'diamond_cross', 'hex',
        'inverted_triangle', 'square', 'square_cross', 'square_x',
        'triangle', 'x']
    spread : str or None, default None
        If 'jitter', spread points out using a jitter transform. If
        'swarm', spread points in beeswarm style. In None or 'none', do
        not spread.
    cat_grid : bool, default False
        If True, show grid lines for categorical axis.
    marker_kwargs : dict
        Keyword arguments to pass when adding markers to the plot.
        ["x", "y", "source", "cat", "legend"] are note allowed because
        they are determined by other inputs.
    jitter_kwargs : dict
        Keyword arguments to be passed to `bokeh.transform.jitter()`. If
        not specified, default is
        `{'distribution': 'normal', 'width': 0.1}`. If the user
        specifies `{'distribution': 'uniform'}`, the `'width'` entry is
        adjusted to 0.4.
    swarm_kwargs : dict
        Keyword arguments for use in generating swarm. Keys with allowed
        values are:
            - 'corral': Either 'gutter' (default) or 'wrap'. This
            specifies how points that are moved too far out are dealt
            with. Using 'gutter', points are overlayed at the maximum
            allowed distance. Using 'wrap', points are reflected inwards
            form the maximal extent and possibly overlayed with other
            points.
            - 'priority': Either 'ascending' (default) or 'descending'.
            Sort order when determining which points get moved in the
            y-direction first.
            - marker_pad_px : Gap between markers in units of pixels,
            default 0.
    jitter : bool, default False
        Deprecated, use `spread`.
    horizontal : bool or None, default None
        Deprecated. Use `q_axis`.
    val : hashable
        Deprecated, use `q`.
    click_policy : str, default 'hide'
        Deprecated. Use `legend_click_policy`.
    kwargs
        Any kwargs to be passed to `bokeh.plotting.figure()` when
        instantiating the figure.
    Returns
    -------
    output : bokeh.plotting.Figure instance
        Plot populated with a strip plot.
    """
    # Protect against mutability of dicts
    jitter_kwargs = copy.copy(jitter_kwargs)
    swarm_kwargs = copy.copy(swarm_kwargs)
    marker_kwargs = copy.copy(marker_kwargs)

    q, legend_click_policy, _ = _parse_deprecations(
        q, q_axis, val, horizontal, "x", click_policy, legend_click_policy, None, None
    )

    # Hand check jitter deprecation
    if jitter is not None:
        if jitter:
            if spread is None:
                spread = "jitter"
                warnings.warn("`jitter` is deprecated. Use spread='jitter'.")
            if spread == "jitter":
                warnings.warn("`jitter` is deprecated. Use spread='jitter'.")
            else:
                raise RuntimeError(
                    "`jitter` is deprecated. Use spread='jitter'. `jitter` and `spread` are in conflict."
                )
        else:
            if spread == "jitter":
                raise RuntimeError(
                    "`jitter` is deprecated. Use `spread`. `jitter` and `spread` are in conflict."
                )
            else:
                warnings.warn("`jitter` is deprecated. Use spread='jitter'.")

    if spread is not None and spread != 'none' and parcoord_column is not None:
        raise NotImplementedError("Parallel coordinate plots are not implemented with jitter or swarm spreading.")

    if palette is None:
        palette = colorcet.b_glasbey_category10

    if show_legend is None:
        if color_column is None:
            show_legend = False
        else:
            show_legend = not _color_column_hexcodes(data, color_column)

    data, q, cats, show_legend = _data_cats(data, q, cats, show_legend, None)

    cats, cols = _check_cat_input(
        data, cats, q, color_column, parcoord_column, tooltips, palette, order, kwargs
    )

    grouped = data.groupby(cats, sort=False)

    if p is None:
        p, factors, color_factors = _cat_figure(
            data, grouped, q, order, color_column, q_axis, kwargs
        )
    else:
        if type(p.x_range) == bokeh.models.ranges.FactorRange and q_axis == "x":
            raise RuntimeError("`q_axis` is 'x', but `p` has a categorical x-axis.")
        elif type(p.y_range) == bokeh.models.ranges.FactorRange and q_axis == "y":
            raise RuntimeError("`q_axis` is 'y', but `p` has a categorical y-axis.")

        _, factors, color_factors = _get_cat_range(
            data, grouped, order, color_column, q_axis
        )

    if tooltips is not None:
        p.add_tools(bokeh.models.HoverTool(tooltips=tooltips, names=["hover_glyphs"]))

    if jitter_kwargs is None:
        jitter_kwargs = dict(width=0.1, mean=0, distribution="normal")
    elif type(jitter_kwargs) != dict:
        raise RuntimeError("`jitter_kwargs` must be a dict.")
    elif "width" not in jitter_kwargs:
        if (
            "distribution" not in jitter_kwargs
            or jitter_kwargs["distribution"] == "uniform"
        ):
            jitter_kwargs["width"] = 0.4
        else:
            jitter_kwargs["width"] = 0.1

    if swarm_kwargs is None:
        swarm_kwargs = dict(corral="gutter", priority="ascending", marker_pad_px=0)
    elif type(swarm_kwargs) != dict:
        raise RuntimeError("`swarm_kwargs` must be a dict.")
    if "corral" not in swarm_kwargs:
        swarm_kwargs["corral"] = "gutter"
    if "priority" not in swarm_kwargs:
        swarm_kwargs["priority"] = "ascending"
    if "marker_pad_px" not in swarm_kwargs:
        swarm_kwargs["marker_pad_px"] = 0

    if marker_kwargs is None:
        marker_kwargs = {}
    elif type(marker_kwargs) != dict:
        raise RuntimeError("`marker_kwargs` must be a dict.")

    if "name" not in marker_kwargs:
        marker_kwargs["name"] = "hover_glyphs"
    if (
        "color" not in marker_kwargs
        and "fill_color" not in marker_kwargs
        and "line_color" not in marker_kwargs
    ):
        if color_column is None:
            color_column = "cat"
            if show_legend:
                warnings.warn(
                    "`color_column` is not specified. No legend will be generated."
                )
                show_legend = False
        if color_factors == "hex":
            marker_kwargs["line_color"] = color_column
            marker_kwargs["fill_color"] = color_column
            if show_legend:
                warnings.warn(
                    "`color_column` consists of hex colors. No legend will be generated."
                )
                show_legend = False
        elif not show_legend:
            marker_kwargs["fill_color"] = bokeh.transform.factor_cmap(
                color_column, palette=palette, factors=color_factors
            )
            marker_kwargs["line_color"] = bokeh.transform.factor_cmap(
                color_column, palette=palette, factors=color_factors
            )

    if marker == "tick":
        marker = "dash"
    marker_fun = _get_marker(p, marker)

    if marker == "dash":
        if spread == "swarm":
            raise RuntimeError(
                "Cannot have 'swarm' spreading with dash or tick markers."
            )
        if "angle" not in marker_kwargs and q_axis == "x":
            marker_kwargs["angle"] = np.pi / 2
        if "size" not in marker_kwargs:
            if q_axis == "x":
                marker_kwargs["size"] = p.frame_height * 0.25 / len(grouped)
            else:
                marker_kwargs["size"] = p.frame_width * 0.25 / len(grouped)
    else:
        if "size" not in marker_kwargs:
            marker_kwargs["size"] = 4
        if "line_width" not in marker_kwargs:
            marker_kwargs["line_width"] = 1

    source_dict = _cat_source_dict(data, cats, cols, color_column)

    if spread == "swarm":
        r = (marker_kwargs["size"] + marker_kwargs["line_width"]) / 2

        if q_axis == 'x' and p.x_range.start is not None and p.x_range.end is not None:
            x_range = [p.x_range.start, p.x_range.end]
        elif q_axis == 'y' and p.y_range.start is not None and p.y_range.end is not None:
            x_range = [p.y_range.start, p.y_range.end]
        else:
            x_range_width = data[q].max() - data[q].min()
            x_range = [
                data[q].min() - 0.05 * x_range_width,
                data[q].max() + 0.05 * x_range_width,
            ]

        y_swarm = (
            grouped[q].transform(_swarm, p, r, x_range, q_axis, **swarm_kwargs).values
        )
        source_dict["__y_swarm"] = [
            (*cat, y_val) if type(cat) == tuple else (cat, y_val)
            for cat, y_val in zip(source_dict["cat"], y_swarm)
        ]

    if q_axis == "x":
        x = q
        if spread == "jitter":
            jitter_kwargs["range"] = p.y_range
            y = bokeh.transform.jitter("cat", **jitter_kwargs)
        elif spread == "swarm":
            y = "__y_swarm"
        else:
            y = "cat"
        if not cat_grid:
            p.ygrid.grid_line_color = None
    else:
        y = q
        if spread == "jitter":
            jitter_kwargs["range"] = p.x_range
            x = bokeh.transform.jitter("cat", **jitter_kwargs)
        elif spread == "swarm":
            x = "__y_swarm"
        else:
            x = "cat"
        if not cat_grid:
            p.xgrid.grid_line_color = None

    if parcoord_column is not None:
        source_pc = _parcoord_source(data, q, cats, q_axis, parcoord_column, factors)

        if parcoord_kwargs is None:
            line_color = "gray"
            parcoord_kwargs = {}
        elif type(parcoord_kwargs) != dict:
            raise RuntimeError("`parcoord_kwargs` must be a dict.")

        if "color" in parcoord_kwargs and "line_color" not in parcoord_kwargs:
            line_color = parcoord_kwargs.pop("color")
        else:
            line_color = parcoord_kwargs.pop("line_color", "gray")

        p.multi_line(
            source=source_pc, xs="xs", ys="ys", line_color=line_color, **parcoord_kwargs
        )

    if color_factors == "hex" or color_column == "cat" or not show_legend:
        marker_fun(
            source=bokeh.models.ColumnDataSource(source_dict),
            x=x,
            y=y,
            **marker_kwargs,
        )
    else:
        items = []
        df = pd.DataFrame(source_dict)
        for i, (name, g) in enumerate(df.groupby(color_column)):
            marker_kwargs["color"] = palette[i % len(palette)]
            mark = marker_fun(source=g, x=x, y=y, **marker_kwargs)
            items.append((g["__label"].iloc[0], [mark]))

        if len(p.legend) == 1:
            for item in items:
                p.legend.items.append(
                    bokeh.models.LegendItem(label=item[0], renderers=item[1])
                )
        else:
            if len(p.legend) > 1:
                warnings.warn(
                    "Ambiguous which legend to add glyphs to. Creating new legend."
                )
            if legend_location in ["right", "left", "above", "below"]:
                legend = bokeh.models.Legend(
                    items=items,
                    location="center",
                    orientation=legend_orientation,
                    title=color_column,
                )
                p.add_layout(legend, legend_location)
            elif (
                legend_location
                in [
                    "top_left",
                    "top_center",
                    "top_right",
                    "center_right",
                    "bottom_right",
                    "bottom_center",
                    "bottom_left",
                    "center_left",
                    "center",
                ]
                or type(legend_location) == tuple
            ):
                legend = bokeh.models.Legend(
                    items=items,
                    location=legend_location,
                    orientation=legend_orientation,
                    title=color_column,
                )
                p.add_layout(legend, "center")
            else:
                raise RuntimeError(
                    'Invalid `legend_location`. Must be a 2-tuple specifying location or one of ["right", "left", "above", "below", "top_left", "top_center", "top_right", "center_right", "bottom_right", "bottom_center", "bottom_left", "center_left", "center"]'
                )

        p.legend.click_policy = legend_click_policy

    return p


def box(
    data=None,
    q=None,
    cats=None,
    q_axis="x",
    palette=None,
    order=None,
    p=None,
    whisker_caps=False,
    display_points=True,
    outlier_marker="circle",
    min_data=5,
    cat_grid=False,
    box_kwargs=None,
    median_kwargs=None,
    whisker_kwargs=None,
    outlier_kwargs=None,
    display_outliers=None,
    horizontal=None,
    val=None,
    **kwargs,
):
    """
    Make a box-and-whisker plot.
    Parameters
    ----------
    data : Pandas DataFrame, 1D Numpy array, or xarray
        DataFrame containing tidy data for plotting.  If a Numpy array,
        a single category is assumed and a box plot with a single box is
        generated from data.
    q : hashable
        Name of column to use as quantitative variable if `data` is a
        Pandas DataFrame. Otherwise, `q` is used as the quantitative
        axis label.
    cats : hashable or list of hashables
        Name of column(s) to use as categorical variable(s).
    q_axis : str, either 'x' or 'y', default 'x'
        Axis along which the quantitative value varies.
    palette : list of strings of hex colors, or single hex string
        If a list, color palette to use. If a single string representing
        a hex color, all glyphs are colored with that color. Default is
        colorcet.b_glasbey_category10 from the colorcet package.
    order : list or None
        If not None, must be a list of unique group names when the input
        data frame is grouped by `cats`. The order of the list specifies
        the ordering of the categorical variables on the categorical
        axis and legend. If None, the categories appear in the order in
        which they appeared in the inputted data frame.
    p : bokeh.plotting.Figure instance, or None (default)
        If None, create a new figure. Otherwise, populate the existing
        figure `p`.
    whisker_caps : bool, default False
        If True, put caps on whiskers. If False, omit caps.
    display_points : bool, default True
        If True, display outliers and any other points that arise from
        categories with fewer than `min_data` data points; otherwise
        suppress them. This should only be False when using the boxes
        as annotation on another plot.
    outlier_marker : str, default 'circle'
        Name of marker to be used in the plot (ignored if `formal` is
        False). Must be one of['asterisk', 'circle', 'circle_cross',
        'circle_x', 'cross', 'dash', 'diamond', 'diamond_cross', 'hex',
        'inverted_triangle', 'square', 'square_cross', 'square_x',
        'triangle', 'x']
    min_data : int, default 5
        Minimum number of data points in a given category in order to
        make a box and whisker. Otherwise, individual data points are
        plotted as in a strip plot.
    cat_grid : bool, default False
        If True, display grid line for categorical axis.
    box_kwargs : dict, default None
        A dictionary of kwargs to be passed into `p.hbar()` or
        `p.vbar()` when constructing the boxes for the box plot.
    median_kwargs : dict, default None
        A dictionary of kwargs to be passed into `p.hbar()` or
        `p.vbar()` when constructing the median line for the box plot.
    whisker_kwargs : dict, default None
        A dictionary of kwargs to be passed into `p.segment()`
        when constructing the whiskers for the box plot.
    outlier_kwargs : dict, default None
        A dictionary of kwargs to be passed into `p.circle()`
        when constructing the outliers for the box plot.
    horizontal : bool or None, default None
        Deprecated. Use `q_axis`.
    val : hashable
        Deprecated, use `q`.
    kwargs
        Kwargs that are passed to bokeh.plotting.figure() in contructing
        the figure.
    Returns
    -------
    output : bokeh.plotting.Figure instance
        Plot populated with box-and-whisker plot.
    Notes
    -----
    Uses the Tukey convention for box plots. The top and bottom of
    the box are respectively the 75th and 25th percentiles of the
    data. The line in the middle of the box is the median. The top
    whisker extends to the maximum of the set of data points that are
    less than 1.5 times the IQR beyond the top of the box, with an
    analogous definition for the lower whisker. Data points not
    between the ends of the whiskers are considered outliers and are
    plotted as individual points.
    """
    # Protect against mutability of dicts
    box_kwargs = copy.copy(box_kwargs)
    median_kwargs = copy.copy(median_kwargs)
    whisker_kwargs = copy.copy(whisker_kwargs)
    outlier_kwargs = copy.copy(outlier_kwargs)

    q, _, _ = _parse_deprecations(
        q, q_axis, val, horizontal, "x", None, None, None, None
    )

    if display_outliers is not None:
        warnings.warn(
            f"`display_outliers` is deprecated. Use `display_points`. Using `display_points={display_outliers}.",
            DeprecationWarning,
        )
        display_points = display_outliers

    if palette is None:
        palette = colorcet.b_glasbey_category10

    data, q, cats, _ = _data_cats(data, q, cats, False, None)

    cats, cols = _check_cat_input(
        data, cats, q, None, None, None, palette, order, box_kwargs
    )

    if outlier_kwargs is None:
        outlier_kwargs = dict()
    elif type(outlier_kwargs) != dict:
        raise RuntimeError("`outlier_kwargs` must be a dict.")

    if box_kwargs is None:
        box_kwargs = {"line_color": None}
        box_width = 0.4
    elif type(box_kwargs) != dict:
        raise RuntimeError("`box_kwargs` must be a dict.")
    else:
        box_width = box_kwargs.pop("width", 0.4)
        if "line_color" not in box_kwargs:
            box_kwargs["line_color"] = None

    if whisker_kwargs is None:
        if "fill_color" in box_kwargs:
            whisker_kwargs = {"line_color": box_kwargs["fill_color"]}
        else:
            whisker_kwargs = {"line_color": "black"}
    elif type(whisker_kwargs) != dict:
        raise RuntimeError("`whisker_kwargs` must be a dict.")

    if median_kwargs is None:
        median_kwargs = {"line_color": "white"}
    elif type(median_kwargs) != dict:
        raise RuntimeError("`median_kwargs` must be a dict.")
    elif "line_color" not in median_kwargs:
        median_kwargs["line_color"] = "white"

    if q_axis == "x":
        if "height" in box_kwargs:
            warnings.warn("'height' entry in `box_kwargs` ignored; using `box_width`.")
            del box_kwargs["height"]
    else:
        if "width" in box_kwargs:
            warnings.warn("'width' entry in `box_kwargs` ignored; using `box_width`.")
            del box_kwargs["width"]

    grouped = data.groupby(cats, sort=False)

    if p is None:
        p, factors, color_factors = _cat_figure(
            data, grouped, q, order, None, q_axis, kwargs
        )
    else:
        _, factors, color_factors = _get_cat_range(data, grouped, order, None, q_axis)

    marker_fun = _get_marker(p, outlier_marker)

    source_box, source_outliers = _box_source(data, cats, q, cols, min_data)

    if "color" in outlier_kwargs:
        if "line_color" in outlier_kwargs or "fill_color" in outlier_kwargs:
            raise RuntimeError(
                "If `color` is in `outlier_kwargs`, `line_color` and `fill_color` cannot be."
            )
    else:
        if "fill_color" in box_kwargs:
            if "fill_color" not in outlier_kwargs:
                outlier_kwargs["fill_color"] = box_kwargs["fill_color"]
            if "line_color" not in outlier_kwargs:
                outlier_kwargs["line_color"] = box_kwargs["fill_color"]
        else:
            if "fill_color" not in outlier_kwargs:
                outlier_kwargs["fill_color"] = bokeh.transform.factor_cmap(
                    "cat", palette=palette, factors=factors
                )
            if "line_color" not in outlier_kwargs:
                outlier_kwargs["line_color"] = bokeh.transform.factor_cmap(
                    "cat", palette=palette, factors=factors
                )

    if "fill_color" not in box_kwargs:
        box_kwargs["fill_color"] = bokeh.transform.factor_cmap(
            "cat", palette=palette, factors=factors
        )

    if q_axis == "x":
        p.segment(
            source=source_box,
            y0="cat",
            y1="cat",
            x0="top",
            x1="top_whisker",
            **whisker_kwargs,
        )
        p.segment(
            source=source_box,
            y0="cat",
            y1="cat",
            x0="bottom",
            x1="bottom_whisker",
            **whisker_kwargs,
        )
        if whisker_caps:
            p.hbar(
                source=source_box,
                y="cat",
                left="top_whisker",
                right="top_whisker",
                height=box_width / 4,
                **whisker_kwargs,
            )
            p.hbar(
                source=source_box,
                y="cat",
                left="bottom_whisker",
                right="bottom_whisker",
                height=box_width / 4,
                **whisker_kwargs,
            )
        p.hbar(
            source=source_box,
            y="cat",
            left="bottom",
            right="top",
            height=box_width,
            **box_kwargs,
        )
        p.hbar(
            source=source_box,
            y="cat",
            left="middle",
            right="middle",
            height=box_width,
            **median_kwargs,
        )
        if display_points:
            marker_fun(source=source_outliers, y="cat", x=q, **outlier_kwargs)
        if not cat_grid:
            p.ygrid.grid_line_color = None
    else:
        p.segment(
            source=source_box,
            x0="cat",
            x1="cat",
            y0="top",
            y1="top_whisker",
            **whisker_kwargs,
        )
        p.segment(
            source=source_box,
            x0="cat",
            x1="cat",
            y0="bottom",
            y1="bottom_whisker",
            **whisker_kwargs,
        )
        if whisker_caps:
            p.vbar(
                source=source_box,
                x="cat",
                bottom="top_whisker",
                top="top_whisker",
                width=box_width / 4,
                **whisker_kwargs,
            )
            p.vbar(
                source=source_box,
                x="cat",
                bottom="bottom_whisker",
                top="bottom_whisker",
                width=box_width / 4,
                **whisker_kwargs,
            )
        p.vbar(
            source=source_box,
            x="cat",
            bottom="bottom",
            top="top",
            width=box_width,
            **box_kwargs,
        )
        p.vbar(
            source=source_box,
            x="cat",
            bottom="middle",
            top="middle",
            width=box_width,
            **median_kwargs,
        )
        if display_points:
            marker_fun(source=source_outliers, x="cat", y=q, **outlier_kwargs)
        if not cat_grid:
            p.xgrid.grid_line_color = None

    return p


def stripbox(
    data=None,
    q=None,
    cats=None,
    q_axis="x",
    palette=None,
    order=None,
    p=None,
    show_legend=False,
    legend_location="right",
    legend_orientation="vertical",
    legend_click_policy="hide",
    top_level="strip",
    color_column=None,
    parcoord_column=None,
    tooltips=None,
    marker="circle",
    spread=None,
    cat_grid=False,
    marker_kwargs=None,
    jitter_kwargs=None,
    spread_kwargs=None,
    parcoord_kwargs=None,
    whisker_caps=True,
    display_points=True,
    min_data=5,
    box_kwargs=None,
    median_kwargs=None,
    whisker_kwargs=None,
    jitter=None,
    horizontal=None,
    val=None,
    click_policy=None,
    **kwargs,
):
    """
    Make a strip plot with a box plot as annotation.
    Parameters
    ----------
    data : Pandas DataFrame, 1D Numpy array, or xarray
        DataFrame containing tidy data for plotting.  If a Numpy array,
        a single category is assumed and a strip plot generated from
        data.
    q : hashable
        Name of column to use as quantitative variable if `data` is a
        Pandas DataFrame. Otherwise, `q` is used as the quantitative
        axis label.
    cats : hashable or list of hashables
        Name of column(s) to use as categorical variable(s).
    q_axis : str, either 'x' or 'y', default 'x'
        Axis along which the quantitative value varies.
    palette : list of strings of hex colors, or single hex string
        If a list, color palette to use. If a single string representing
        a hex color, all glyphs are colored with that color. Default is
        colorcet.b_glasbey_category10 from the colorcet package.
    order : list or None
        If not None, must be a list of unique group names when the input
        data frame is grouped by `cats`. The order of the list specifies
        the ordering of the categorical variables on the categorical
        axis and legend. If None, the categories appear in the order in
        which they appeared in the inputted data frame.
    p : bokeh.plotting.Figure instance, or None (default)
        If None, create a new figure. Otherwise, populate the existing
        figure `p`.
    top_level : str, default 'strip'
        If 'box', the box plot is overlaid. If 'strip', the strip plot
        is overlaid.
    show_legend : bool, default False
        If True, display legend.
    legend_location : str, default 'right'
        Location of legend. If one of "right", "left", "above", or
        "below", the legend is placed outside of the plot area. If one
        of "top_left", "top_center", "top_right", "center_right",
        "bottom_right", "bottom_center", "bottom_left", "center_left",
        or "center", the legend is placed within the plot area. If a
        2-tuple, legend is placed according to the coordinates in the
        tuple.
    legend_orientation : str, default 'vertical'
        Either 'horizontal' or 'vertical'.
    legend_click_policy : str, default 'hide'
        Either 'hide', 'mute', or None; how the glyphs respond when the
        corresponding category is clicked in the legend.
    color_column : hashable, default None
        Column of `data` to use in determining color of glyphs. The data
        in the color_column are assumed to be categorical. If the data
        in color_column consist entirely of hex colors, then those
        colors are directly used to color the glyphs. If None,
        then `cats` is used.
    parcoord_column : hashable, default None
        Column of `data` to use to construct a parallel coordinate plot.
        Data points with like entries in the parcoord_column are
        connected with lines in the strip plot.
    tooltips : list of 2-tuples
        Specification for tooltips as per Bokeh specifications. For
        example, if we want `col1` and `col2` tooltips, we can use
        `tooltips=[('label 1': '@col1'), ('label 2': '@col2')]`.
    marker : str, default 'circle'
        Name of marker to be used in the plot (ignored if `formal` is
        False). Must be one of['asterisk', 'circle', 'circle_cross',
        'circle_x', 'cross', 'dash', 'diamond', 'diamond_cross', 'hex',
        'inverted_triangle', 'square', 'square_cross', 'square_x',
        'triangle', 'x']
    spread : str or None, default None
        If 'jitter', spread points out using a jitter transform. If
        'swarm', spread points in beeswarm style. In None or 'none', do
        not spread.
    cat_grid : bool, default False
        If True, display grid line for categorical axis.
    marker_kwargs : dict
        Keyword arguments to pass when adding markers to the plot.
        ["x", "y", "source", "cat", "legend"] are note allowed because
        they are determined by other inputs.
    jitter_kwargs : dict
        Keyword arguments to be passed to `bokeh.transform.jitter()`. If
        not specified, default is
        `{'distribution': 'normal', 'width': 0.1}`. If the user
        specifies `{'distribution': 'uniform'}`, the `'width'` entry is
        adjusted to 0.4.
    swarm_kwargs : dict
        Keyword arguments for use in generating swarm. Keys with allowed
        values are:
            - 'corral': Either 'gutter' (default) or 'wrap'. This
            specifies how points that are moved too far out are dealt
            with. Using 'gutter', points are overlayed at the maximum
            allowed distance. Using 'wrap', points are reflected inwards
            form the maximal extent and possibly overlayed with other
            points.
            - 'priority': Either 'ascending' (default) or 'descending'.
            Sort order when determining which points get moved in the
            y-direction first.
            - marker_pad_px : Gap between markers in units of pixels,
            default 0.
    whisker_caps : bool, default True
        If True, put caps on whiskers. If False, omit caps.
    min_data : int, default 5
        Minimum number of data points in a given category in order to
        make a box and whisker. Otherwise, individual data points are
        plotted as in a strip plot.
    box_kwargs : dict, default None
        A dictionary of kwargs to be passed into `p.hbar()` or
        `p.vbar()` when constructing the boxes for the box plot.
    median_kwargs : dict, default None
        A dictionary of kwargs to be passed into `p.hbar()` or
        `p.vbar()` when constructing the median line for the box plot.
    whisker_kwargs : dict, default None
        A dictionary of kwargs to be passed into `p.segment()`
        when constructing the whiskers for the box plot.
    jitter : bool, default False
        Deprecated, use `spread`.
    horizontal : bool or None, default None
        Deprecated. Use `q_axis`.
    val : hashable
        Deprecated, use `q`.
    click_policy : str, default 'hide'
        Deprecated. Use `legend_click_policy`.
    kwargs
        Any kwargs to be passed to `bokeh.plotting.figure()` when
        instantiating the figure.
    Returns
    -------
    output : bokeh.plotting.Figure instance
        Plot populated with a strip-box plot.
    """
    # Protect against mutability of dicts
    box_kwargs = copy.copy(box_kwargs)
    median_kwargs = copy.copy(median_kwargs)
    whisker_kwargs = copy.copy(whisker_kwargs)
    jitter_kwargs = copy.copy(jitter_kwargs)
    spread_kwargs = copy.copy(spread_kwargs)
    marker_kwargs = copy.copy(marker_kwargs)
    parcoord_kwargs = copy.copy(parcoord_kwargs)

    # Set defaults
    if box_kwargs is None:
        box_kwargs = dict(line_color="gray", fill_alpha=0)
    if "color" not in box_kwargs and "line_color" not in box_kwargs:
        box_kwargs["line_color"] = "gray"
    if ("fill_alpha" not in box_kwargs) and ("fill_color" not in box_kwargs):
        box_kwargs["fill_alpha"] = 0
    elif ("fill_color" in box_kwargs) and ("fill_alpha" not in box_kwargs):
        box_kwargs["fill_alpha"] = 0.5

    if median_kwargs is None:
        median_kwargs = dict(line_color="gray")
    if "color" not in box_kwargs and "line_color" not in median_kwargs:
        median_kwargs["line_color"] = "gray"

    if whisker_kwargs is None:
        whisker_kwargs = dict(line_color="gray")
    if "color" not in box_kwargs and "line_color" not in whisker_kwargs:
        whisker_kwargs["line_color"] = "gray"

    if top_level == "box":
        p = strip(
            data=data,
            q=q,
            cats=cats,
            q_axis=q_axis,
            palette=palette,
            order=order,
            p=p,
            show_legend=show_legend,
            legend_location=legend_location,
            legend_orientation=legend_orientation,
            legend_click_policy=legend_click_policy,
            color_column=color_column,
            parcoord_column=parcoord_column,
            tooltips=tooltips,
            marker=marker,
            spread=spread,
            cat_grid=cat_grid,
            marker_kwargs=marker_kwargs,
            jitter_kwargs=jitter_kwargs,
            # spread_kwargs=spread_kwargs,
            parcoord_kwargs=parcoord_kwargs,
            jitter=jitter,
            horizontal=horizontal,
            val=val,
            click_policy=click_policy,
            **kwargs,
        )

        p = box(
            data=data,
            q=q,
            cats=cats,
            q_axis=q_axis,
            palette=palette,
            order=order,
            p=p,
            display_points=False,
            whisker_caps=whisker_caps,
            min_data=min_data,
            box_kwargs=box_kwargs,
            median_kwargs=median_kwargs,
            whisker_kwargs=whisker_kwargs,
            horizontal=horizontal,
            val=val,
        )
    elif top_level == "strip":
        p = box(
            data=data,
            q=q,
            cats=cats,
            q_axis=q_axis,
            palette=palette,
            order=order,
            p=p,
            display_points=False,
            cat_grid=cat_grid,
            whisker_caps=whisker_caps,
            min_data=min_data,
            box_kwargs=box_kwargs,
            median_kwargs=median_kwargs,
            whisker_kwargs=whisker_kwargs,
            horizontal=horizontal,
            val=val,
            **kwargs,
        )

        p = strip(
            data=data,
            q=q,
            cats=cats,
            q_axis=q_axis,
            palette=palette,
            order=order,
            p=p,
            show_legend=show_legend,
            legend_location=legend_location,
            legend_orientation=legend_orientation,
            legend_click_policy=legend_click_policy,
            color_column=color_column,
            parcoord_column=parcoord_column,
            tooltips=tooltips,
            marker=marker,
            spread=spread,
            cat_grid=cat_grid,
            marker_kwargs=marker_kwargs,
            jitter_kwargs=jitter_kwargs,
            # spread_kwargs=spread_kwargs,
            parcoord_kwargs=parcoord_kwargs,
            jitter=jitter,
            horizontal=horizontal,
            val=val,
            click_policy=click_policy,
            **kwargs,
        )
    else:
        raise RuntimeError("Invalid `top_level`. Allowed values are 'box' and 'strip'.")

    return p


def striphistogram(
    data=None,
    q=None,
    cats=None,
    q_axis="x",
    palette=None,
    order=None,
    p=None,
    show_legend=None,
    legend_location="right",
    legend_orientation="vertical",
    legend_click_policy="hide",
    top_level="strip",
    color_column=None,
    parcoord_column=None,
    tooltips=None,
    marker="circle",
    spread=None,
    cat_grid=True,
    marker_kwargs=None,
    jitter_kwargs=None,
    spread_kwargs=None,
    parcoord_kwargs=None,
    bins="freedman-diaconis",
    style=None,
    mirror=True,
    hist_height=0.75,
    conf_int=False,
    ptiles=[2.5, 97.5],
    n_bs_reps=10000,
    line_kwargs=None,
    fill_kwargs=None,
    conf_int_kwargs=None,
    kind=None,
    jitter=None,
    horizontal=None,
    val=None,
    click_policy=None,
    **kwargs,
):
    """
    Make a strip plot with a histogram as annotation.
    Parameters
    ----------
    data : Pandas DataFrame, 1D Numpy array, or xarray
        DataFrame containing tidy data for plotting.  If a Numpy array,
        a single category is assumed and a strip plot generated from
        data.
    q : hashable
        Name of column to use as quantitative variable if `data` is a
        Pandas DataFrame. Otherwise, `q` is used as the quantitative
        axis label.
    cats : hashable or list of hashables
        Name of column(s) to use as categorical variable(s).
    q_axis : str, either 'x' or 'y', default 'x'
        Axis along which the quantitative value varies.
    palette : list of strings of hex colors, or single hex string
        If a list, color palette to use. If a single string representing
        a hex color, all glyphs are colored with that color. Default is
        colorcet.b_glasbey_category10 from the colorcet package.
    order : list or None
        If not None, must be a list of unique group names when the input
        data frame is grouped by `cats`. The order of the list specifies
        the ordering of the categorical variables on the categorical
        axis and legend. If None, the categories appear in the order in
        which they appeared in the inputted data frame.
    p : bokeh.plotting.Figure instance, or None (default)
        If None, create a new figure. Otherwise, populate the existing
        figure `p`.
    top_level : str, default 'strip'
        If 'box', the box plot is overlaid. If 'strip', the strip plot
        is overlaid.
    show_legend : bool, default False
        If True, display legend.
    legend_location : str, default 'right'
        Location of legend. If one of "right", "left", "above", or
        "below", the legend is placed outside of the plot area. If one
        of "top_left", "top_center", "top_right", "center_right",
        "bottom_right", "bottom_center", "bottom_left", "center_left",
        or "center", the legend is placed within the plot area. If a
        2-tuple, legend is placed according to the coordinates in the
        tuple.
    legend_orientation : str, default 'vertical'
        Either 'horizontal' or 'vertical'.
    legend_click_policy : str, default 'hide'
        Either 'hide', 'mute', or None; how the glyphs respond when the
        corresponding category is clicked in the legend.
    color_column : hashable, default None
        Column of `data` to use in determining color of glyphs. The data
        in the color_column are assumed to be categorical. If the data
        in color_column consist entirely of hex colors, then those
        colors are directly used to color the glyphs. If None,
        then `cats` is used.
    parcoord_column : hashable, default None
        Column of `data` to use to construct a parallel coordinate plot.
        Data points with like entries in the parcoord_column are
        connected with lines in the strip plot.
    tooltips : list of 2-tuples
        Specification for tooltips as per Bokeh specifications. For
        example, if we want `col1` and `col2` tooltips, we can use
        `tooltips=[('label 1': '@col1'), ('label 2': '@col2')]`.
    marker : str, default 'circle'
        Name of marker to be used in the plot (ignored if `formal` is
        False). Must be one of['asterisk', 'circle', 'circle_cross',
        'circle_x', 'cross', 'dash', 'diamond', 'diamond_cross', 'hex',
        'inverted_triangle', 'square', 'square_cross', 'square_x',
        'triangle', 'x']
    jitter : bool, default False
        If True, apply a jitter transform to the glyphs.
    cat_grid : bool, default True
        If True, display grid line for categorical axis.
    marker_kwargs : dict
        Keyword arguments to pass when adding markers to the plot.
        ["x", "y", "source", "cat", "legend"] are note allowed because
        they are determined by other inputs.
    jitter_kwargs : dict
        Keyword arguments to be passed to `bokeh.transform.jitter()`. If
        not specified, default is
        `{'distribution': 'normal', 'width': 0.1}`. If the user
        specifies `{'distribution': 'uniform'}`, the `'width'` entry is
        adjusted to 0.4.
    bins : int, array_like, or str, default 'freedman-diaconis'
        If int or array_like, setting for `bins` kwarg to be passed to
        `np.histogram()`. If 'exact', then each unique value in the
        data gets its own bin. If 'integer', then integer data is
        assumed and each integer gets its own bin. If 'sqrt', uses the
        square root rule to determine number of bins. If
        `freedman-diaconis`, uses the Freedman-Diaconis rule for number
        of bins.
    style : None or one of ['step', 'step_filled']
        Default for overlayed histograms is 'step' and for stacked
        histograms 'step_filled'. The exception is when `cont_int` is
        True, in which case `style` must be 'step'.
    mirror : bool, default True
        If True, reflect the histogram through zero.
    hist_height : float, default 0.75
        Maximal height of histogram of its confidence interval as a
        fraction of available height along categorical axis. Only active
        when `arrangement` is 'stack'.
    conf_int : bool, default False
        If True, display confidence interval of ECDF.
    ptiles : list, default [2.5, 97.5]
        The percentiles to use for the confidence interval of the
        histogram. Ignored if `conf_int` is False.
    n_bs_reps : int, default 10,000
        Number of bootstrap replicates to do to compute confidence
        interval of histogram. Ignored if `conf_int` is False.
    line_kwargs : dict
        Keyword arguments to pass to `p.line()` in constructing the
        histograms. By default, {"line_width": 2}.
    fill_kwargs : dict
        Keyword arguments to pass to `p.patch()` when making the fill
        for the step-filled histogram or confidence intervals. Ignored
        if `style = 'step'` and `conf_int` is False. By default
        {"fill_alpha": 0.3, "line_alpha": 0}.
    horizontal : bool or None, default None
        Deprecated. Use `q_axis`.
    val : hashable
        Deprecated, use `q`.
    click_policy : str, default 'hide'
        Deprecated. Use `legend_click_policy`.
    conf_int_kwargs : dict
        Deprecated. Use `fill_kwargs`.
    kind : str, default 'step_filled'
        Deprecated. Use `style`.
    kwargs
        Any kwargs to be passed to `bokeh.plotting.figure()` when
        instantiating the figure.
    Returns
    -------
    output : bokeh.plotting.Figure instance
        Plot populated with a strip-histogram plot.
    Notes
    -----
    .. Histograms are all normalized, as would be the case using the
       `density=True` kwargs of iqplot.histogram()`. This is necessary
       because there is no quantitative axis for the height of the
       histogram in a strip plot.
    """
    # Protect against mutability of dicts
    jitter_kwargs = copy.copy(jitter_kwargs)
    spread_kwargs = copy.copy(spread_kwargs)
    marker_kwargs = copy.copy(marker_kwargs)
    parcoord_kwargs = copy.copy(parcoord_kwargs)
    line_kwargs = copy.copy(line_kwargs)
    fill_kwargs = copy.copy(fill_kwargs)

    # Set defaults
    if color_column is not None and color_column != cats:
        if line_kwargs is None:
            line_kwargs = {}
        if fill_kwargs is None:
            fill_kwargs = {}
        if "color" not in line_kwargs and "line_color" not in line_kwargs:
            line_kwargs["line_color"] = "gray"
        if "color" not in fill_kwargs and "fill_color" not in fill_kwargs:
            fill_kwargs["fill_color"] = "gray"

    if style is None:
        if conf_int:
            style = "step"
        else:
            style = "step_filled"

    if top_level == "histogram":
        p = strip(
            data=data,
            q=q,
            cats=cats,
            q_axis=q_axis,
            palette=palette,
            order=order,
            p=p,
            show_legend=show_legend,
            color_column=color_column,
            parcoord_column=parcoord_column,
            tooltips=tooltips,
            marker=marker,
            spread=spread,
            cat_grid=cat_grid,
            marker_kwargs=marker_kwargs,
            jitter_kwargs=jitter_kwargs,
            # spread_kwargs=spread_kwargs,
            parcoord_kwargs=parcoord_kwargs,
            jitter=jitter,
            horizontal=horizontal,
            val=val,
            click_policy=click_policy,
            **kwargs,
        )

        p = histogram(
            data=data,
            q=q,
            cats=cats,
            q_axis=q_axis,
            palette=palette,
            order=order,
            p=p,
            rug=False,
            show_legend=False,
            bins=bins,
            density=True,
            style=style,
            arrangement="stack",
            mirror=mirror,
            hist_height=hist_height,
            conf_int=conf_int,
            ptiles=ptiles,
            n_bs_reps=n_bs_reps,
            line_kwargs=line_kwargs,
            fill_kwargs=fill_kwargs,
            conf_int_kwargs=conf_int_kwargs,
            kind=kind,
        )
    elif top_level == "strip":
        p = histogram(
            data=data,
            q=q,
            cats=cats,
            q_axis=q_axis,
            palette=palette,
            order=order,
            p=p,
            rug=False,
            show_legend=False,
            bins=bins,
            density=True,
            style=style,
            arrangement="stack",
            mirror=mirror,
            hist_height=hist_height,
            conf_int=conf_int,
            ptiles=ptiles,
            n_bs_reps=n_bs_reps,
            line_kwargs=line_kwargs,
            fill_kwargs=fill_kwargs,
            conf_int_kwargs=conf_int_kwargs,
            kind=kind,
            **kwargs,
        )

        p = strip(
            data=data,
            q=q,
            cats=cats,
            q_axis=q_axis,
            palette=palette,
            order=order,
            p=p,
            cat_grid=cat_grid,
            show_legend=show_legend,
            color_column=color_column,
            parcoord_column=parcoord_column,
            tooltips=tooltips,
            marker=marker,
            spread=spread,
            marker_kwargs=marker_kwargs,
            jitter_kwargs=jitter_kwargs,
            # spread_kwargs=spread_kwargs,
            parcoord_kwargs=parcoord_kwargs,
            jitter=jitter,
            horizontal=horizontal,
            val=val,
        )

        if not cat_grid:
            p.xgrid.grid_line_color = None
    else:
        raise RuntimeError(
            "Invalid `top_level`. Allowed values are 'histogram' and 'strip'."
        )

    return p


def _get_cat_range(df, grouped, order, color_column, q_axis):
    if order is None:
        if isinstance(list(grouped.groups.keys())[0], tuple):
            factors = tuple(
                [tuple([str(k) for k in key]) for key in grouped.groups.keys()]
            )
        else:
            factors = tuple([str(key) for key in grouped.groups.keys()])
    else:
        if type(order[0]) in [list, tuple]:
            factors = tuple([tuple([str(k) for k in key]) for key in order])
        else:
            factors = tuple([str(entry) for entry in order])

    if q_axis == "x":
        cat_range = bokeh.models.FactorRange(*(factors[::-1]))
    elif q_axis == "y":
        cat_range = bokeh.models.FactorRange(*factors)

    if color_column is None:
        color_factors = factors
    elif _color_column_hexcodes(df, color_column):
        color_factors = "hex"
    else:
        color_factors = tuple(sorted(list(df[color_column].unique().astype(str))))

    return cat_range, factors, color_factors


def _color_column_hexcodes(df, color_column):
    """Return True of the color column consists of all hex codes"""
    try:
        return df[color_column].str.match(r"^#([A-Fa-f0-9]{6}|[A-Fa-f0-9]{3})$").all()
    except:
        return False


def _cat_figure(df, grouped, q, order, color_column, q_axis, kwargs):
    cat_range, factors, color_factors = _get_cat_range(
        df, grouped, order, color_column, q_axis
    )

    kwargs = _fig_dimensions(kwargs)

    if q_axis == "x":
        if "x_axis_label" not in kwargs:
            kwargs["x_axis_label"] = q

        if "y_axis_type" in kwargs:
            warnings.warn("`y_axis_type` specified for categorical axis. Ignoring.")
            del kwargs["y_axis_type"]

        kwargs["y_range"] = cat_range
    elif q_axis == "y":
        if "y_axis_label" not in kwargs:
            kwargs["y_axis_label"] = q

        if "x_axis_type" in kwargs:
            warnings.warn("`x_axis_type` specified for categorical axis. Ignoring.")
            del kwargs["x_axis_type"]

        kwargs["x_range"] = cat_range

    return bokeh.plotting.figure(**kwargs), factors, color_factors


def _cat_source_dict(df, cats, cols, color_column):
    cat_source, labels = _source_and_labels_from_cats(df, cats)

    if type(cols) in [list, tuple, pd.core.indexes.base.Index]:
        source_dict = {col: list(df[col].values) for col in cols}
    else:
        source_dict = {cols: list(df[cols].values)}

    source_dict["cat"] = cat_source
    if color_column in [None, "cat"]:
        source_dict["__label"] = labels
    else:
        source_dict["__label"] = list(df[color_column].astype(str).values)
        source_dict[color_column] = list(df[color_column].astype(str).values)

    return source_dict


def _parcoord_source(data, q, cats, q_axis, parcoord_column, factors):
    if type(cats) not in [list, tuple]:
        cats = [cats]
        tuple_factors = False
    else:
        tuple_factors = True

    grouped_parcoord = data.groupby(parcoord_column)
    xs = []
    ys = []
    for t, g in grouped_parcoord:
        xy = []
        for _, r in g.iterrows():
            if tuple_factors:
                xy.append([tuple([r[cat] for cat in cats]), r[q]])
            else:
                xy.append([r[cats[0]], r[q]])

        if len(xy) > 1:
            xy.sort(key=lambda a: factors.index(a[0]))
            xs_pc = []
            ys_pc = []
            for pair in xy:
                xs_pc.append(pair[0])
                ys_pc.append(pair[1])

            if q_axis == "y":
                xs.append(xs_pc)
                ys.append(ys_pc)
            else:
                xs.append(ys_pc)
                ys.append(xs_pc)

    return bokeh.models.ColumnDataSource(dict(xs=xs, ys=ys))


def _outliers(data, min_data):
    if len(data) >= min_data:
        bottom, middle, top = np.percentile(data, [25, 50, 75])
        iqr = top - bottom
        outliers = data[(data > top + 1.5 * iqr) | (data < bottom - 1.5 * iqr)]
        return outliers
    else:
        return data


def _box_and_whisker(data, min_data):
    if len(data) >= min_data:
        middle = data.median()
        bottom = data.quantile(0.25)
        top = data.quantile(0.75)
        iqr = top - bottom
        top_whisker = max(data[data <= top + 1.5 * iqr].max(), top)
        bottom_whisker = min(data[data >= bottom - 1.5 * iqr].min(), bottom)
        return pd.Series(
            {
                "middle": middle,
                "bottom": bottom,
                "top": top,
                "top_whisker": top_whisker,
                "bottom_whisker": bottom_whisker,
            }
        )
    else:
        return pd.Series(
            {
                "middle": np.nan,
                "bottom": np.nan,
                "top": np.nan,
                "top_whisker": np.nan,
                "bottom_whisker": np.nan,
            }
        )


def _box_source(df, cats, q, cols, min_data):
    """Construct a data frame for making box plot."""
    # Need to reset index for use in slicing outliers
    df_source = df.reset_index(drop=True)

    if cats is None:
        grouped = df_source
    else:
        grouped = df_source.groupby(cats, sort=False)

    # Data frame for boxes and whiskers
    df_box = grouped[q].apply(_box_and_whisker, min_data).unstack().reset_index()
    df_box = df_box.dropna()

    source_box = bokeh.models.ColumnDataSource(
        _cat_source_dict(
            df_box,
            cats,
            ["middle", "bottom", "top", "top_whisker", "bottom_whisker"],
            None,
        )
    )

    # Data frame for outliers
    s_outliers = grouped[q].apply(_outliers, min_data)

    # If no cat has enough data, just use everything as an "outlier"
    if len(s_outliers) == len(df_source):
        df_outliers = df_source.copy()
        inds = df_source.index
    else:
        df_outliers = s_outliers.reset_index()
        inds = s_outliers.index.get_level_values(-1)

    df_outliers.index = inds
    df_outliers[cols] = df_source.loc[inds, cols]

    source_outliers = bokeh.models.ColumnDataSource(
        _cat_source_dict(df_outliers, cats, cols, None)
    )

    return source_box, source_outliers


def _out_every_interval(y, intervals, epsilon=1e-6):
    """Check to see if a value `y` list outside every interval in a
    list of 2-tuples `intervals`."""
    for interval in intervals:
        if y > interval[0] + epsilon and y < interval[1] - epsilon:
            return False

    return True


def _swarm_px(
    x,
    frame_width,
    r,
    x_range,
    max_y_px=np.inf,
    corral="gutter",
    priority="ascending",
    marker_pad_px=0,
):
    """Computes y-coordinates in pixel units for a swarm plot, where x
    is the quantitative axis.
    Parameters
    ----------
    x : array_like
        Array of values of quantitative varaible.
    frame_width : int or float
        Width of plot frame in pixels.
    r : float
        Radius of marker, which is typically the (marker size + 1) / 2,
        where the +1 is due to the standard line width for a marker of
        one pixel.
    x_range : list
        List of length 2, where the first entry is the lower limit of
        the quantitative axis and the second entry is the upper limit of
        the quantitative axis.
    max_y_px : float, default np.inf
        Maximum allowed displacement. Any points with computed y-values
        beyond this will be corraled.
    corral : str, default 'gutter'
        Either 'gutter' or 'wrap'. How to corral points beyond the
        maximum displacement.
    priority : str, either
        Sort order when determining which points get moved in the
        y-direction first. Either 'ascending' or 'descending'.
    marker_pad_px : int of float
        Gap between markers in units of pixels.
    Returns
    -------
    y : array_like
        Array of y-values in units of pixels.
    n_overrun : int
        Number of data points that overrun max_y_px.
    """
    # Sort x according to priority
    if priority == "ascending":
        inds = [i[0] for i in sorted(enumerate(x), key=lambda x: x[1])]
    elif priority == "descending":
        inds = [i[0] for i in sorted(enumerate(x), key=lambda x: -x[1])]
    elif priority == "random":
        raise NotImplementedError("'random' priority is not yet implemented.")
    else:
        raise NotImplementedError("Custom `priority` not yet implemented.")

    x_pixels = (x[inds] - x_range[0]) * frame_width / (x_range[1] - x_range[0])
    y_pixels = np.inf * np.ones_like(x_pixels)

    for i in range(len(x_pixels)):
        intervals = []

        # Scan points to the right
        for j in range(i + 1, len(x_pixels)):
            dist = abs(x_pixels[i] - x_pixels[j])
            if dist > 2 * r:
                if priority in ["ascending", "descending"]:
                    break
                else:
                    continue
            if y_pixels[j] < np.inf:
                offset = np.sqrt(4 * r**2 - dist**2) + marker_pad_px
                intervals.append([y_pixels[j] - offset, y_pixels[j] + offset])

        # Scan points to the left
        for j in range(i - 1, -1, -1):
            dist = abs(x_pixels[i] - x_pixels[j])
            if dist > 2 * r:
                if priority in ["ascending", "descending"]:
                    break
                else:
                    continue
            if y_pixels[j] < np.inf:
                offset = np.sqrt(4 * r**2 - dist**2) + marker_pad_px
                intervals.append([y_pixels[j] - offset, y_pixels[j] + offset])

        # Any y-position must be outside all intervals and should be at the edge of one of the intervals
        # Need to find first candidate the satisfies this
        y_cand = 0
        if len(intervals) > 0:
            candidates = sorted(np.array(intervals).flatten(), key=abs)
            for cand in candidates:
                if _out_every_interval(cand, intervals):
                    y_cand = cand
                    break
        y_pixels[i] = y_cand

    # Clean up points landing too far out
    n_overrun = 0
    for i in range(len(x)):
        if abs(y_pixels[i]) > max_y_px:
            n_overrun += 1
            if corral == "gutter":
                y_pixels[i] = np.sign(y_pixels[i]) * max_y_px
            elif corral == "wrap":
                y_pixels[i] = np.sign(y_pixels[i]) * 2 * max_y_px - y_pixels[i]

    # Build output for y in pixels
    y_out = np.empty_like(x)
    for i in range(len(x)):
        y_out[inds[i]] = y_pixels[i]

    return y_out, n_overrun


def _swarm(
    x, p, r, x_range, q_axis, corral="gutter", priority="ascending", marker_pad_px=0
):
    if q_axis == "x":
        extra_padding = 0
        if type(p.y_range.factors[0]) == tuple:
            if len(p.y_range.factors[0]) >= 2:
                extra_padding += p.y_range.group_padding
            if len(p.y_range.factors[0]) > 2:
                extra_padding += (len(p.y_range.factors[0]) - 2) * p.y_range.subgroup_padding
        h = p.frame_height
        w = p.frame_width
        n_factors = len(p.y_range.factors) + extra_padding
    else:
        extra_padding = 0
        if type(p.x_range.factors[0]) == tuple:
            if len(p.x_range.factors[0]) >= 2:
                extra_padding += p.x_range.group_padding
            if len(p.x_range.factors[0]) > 2:
                extra_padding += (len(p.x_range.factors[0]) - 2) * p.x_range.subgroup_padding
        w = p.frame_height
        h = p.frame_width
        n_factors = len(p.x_range.factors) + extra_padding

    max_y_px = h / n_factors / 2 - 2 * r
    y_pixels, n_overrun = _swarm_px(
        np.array(x), w, r, x_range, max_y_px=max_y_px, marker_pad_px=marker_pad_px
    )

    if n_overrun > 0:
        hw = 'height' if q_axis == 'x' else 'width'
        warnings.warn(
            f"{n_overrun} data points exceed maximum {hw}. Consider using spread='jitter' or increasing the frame {hw}."
        )

    return y_pixels / h * n_factors

def histogram(
    data=None,
    q=None,
    cats=None,
    palette=None,
    order=None,
    q_axis="x",
    p=None,
    rug=None,
    rug_height=None,
    show_legend=None,
    legend_label=None,
    legend_location="right",
    legend_orientation="vertical",
    legend_click_policy="hide",
    tooltips=None,
    bins="freedman-diaconis",
    density=None,
    style=None,
    arrangement="stack",
    mirror=False,
    hist_height=0.75,
    conf_int=False,
    ptiles=[2.5, 97.5],
    n_bs_reps=10000,
    line_kwargs=None,
    fill_kwargs=None,
    rug_kwargs=None,
    conf_int_kwargs=None,
    horizontal=None,
    val=None,
    click_policy=None,
    kind=None,
    **kwargs,
):
    """
    Make a plot of histograms.
    Parameters
    ----------
    data : Pandas DataFrame, 1D Numpy array, or xarray
        DataFrame containing tidy data for plotting.  If a Numpy array,
        a single category is assumed and a box plot generated from
        data.
    q : hashable
        Name of column to use as quantitative variable if `data` is a
        Pandas DataFrame. Otherwise, `q` is used as the quantitative
        axis label.
    cats : hashable or list of hashables
        Name of column(s) to use as categorical variable(s).
    q_axis : str, either 'x' or 'y', default 'x'
        Axis along which the quantitative value varies.
    palette : list colors, or single color string
        If a list, color palette to use. If a single string representing
        a color, all glyphs are colored with that color. Default is
        colorcet.b_glasbey_category10 from the colorcet package.
    order : list or None
        If not None, must be a list of unique group names when the input
        data frame is grouped by `cats`. The order of the list specifies
        the ordering of the categorical variables in the legend. If
        None, the categories appear in the order in which they appeared
        in the inputted data frame.
    p : bokeh.plotting.Figure instance, or None (default)
        If None, create a new figure. Otherwise, populate the existing
        figure `p`.
    rug : bool, default True
        If True, also include a rug plot. If, however, `bins` is 'exact'
        or 'integer', the `rug` kwarg is ignored.
    rug_height : float, default None
        Height of the rug plot as a fraction of the highest point in the
        histograms. For 'overlay' arrangement, default is 0.05. For
        'stacked' arrangement, default is 0.2.
    legend_label : str, default None
        If `cats` is None and `show_legend` is True, then if
        `legend_label` is not None, a legend is created for the glyph
        on the plot and labeled with `legend_label`. Otherwise, no
        legend is created if `cats` is None.
    legend_location : str, default 'right'
        Location of legend. If one of "right", "left", "above", or
        "below", the legend is placed outside of the plot area. If one
        of "top_left", "top_center", "top_right", "center_right",
        "bottom_right", "bottom_center", "bottom_left", "center_left",
        or "center", the legend is placed within the plot area. If a
        2-tuple, legend is placed according to the coordinates in the
        tuple.
    legend_orientation : str, default 'vertical'
        Either 'horizontal' or 'vertical'.
    legend_click_policy : str, default 'hide'
        Either 'hide', 'mute', or None; how the glyphs respond when the
        corresponding category is clicked in the legend.
    tooltips : list of 2-tuples
        Specification for tooltips as per Bokeh specifications. For
        example, if we want `col1` and `col2` tooltips, we can use
        `tooltips=[('label 1': '@col1'), ('label 2': '@col2')]`. Ignored
        if `rug` is False.
    bins : int, array_like, or str, default 'freedman-diaconis'
        If int or array_like, setting for `bins` kwarg to be passed to
        `np.histogram()`. If 'exact', then each unique value in the
        data gets its own bin. If 'integer', then integer data is
        assumed and each integer gets its own bin. If 'sqrt', uses the
        square root rule to determine number of bins. If
        `freedman-diaconis`, uses the Freedman-Diaconis rule for number
        of bins.
    density : None or bool
        If True, normalize the histograms. Otherwise, base the
        histograms on counts. Must be True for arrangement = 'stack'
        because there is no absolute scale along the categorical axis.
        For arrangement = 'overlay', `density` is assumed to be False if
        None is given.
    style : None or one of ['step', 'step_filled']
        Default for overlayed histograms is 'step' and for stacked
        histograms 'step_filled'. The exception is when `cont_int` is
        True, in which case `style` must be 'step'.
    arrangement : 'stack' or 'overlay', default 'stack'
        Arrangement of histograms. If 'overlay', histograms are overlaid
        on the same plot. If 'stack', histograms are stacked one on top
        of the other.
    mirror : bool, default False
        If True, reflect the histogram through zero. Ignored if
        `arrangement == 'overlay'`.
    hist_height : float, default 0.75
        Maximal height of histogram of its confidence interval as a
        fraction of available height along categorical axis. Only active
        when `arrangement` is 'stack'.
    conf_int : bool, default False
        If True, display confidence interval of ECDF.
    ptiles : list, default [2.5, 97.5]
        The percentiles to use for the confidence interval. Ignored if
        `conf_int` is False.
    n_bs_reps : int, default 10,000
        Number of bootstrap replicates to do to compute confidence
        interval. Ignored if `conf_int` is False.
    line_kwargs : dict
        Keyword arguments to pass to `p.line()` in constructing the
        histograms. By default, {"line_width": 2}.
    fill_kwargs : dict
        Keyword arguments to pass to `p.patch()` when making the fill
        for the step-filled histogram or confidence intervals. Ignored
        if `style = 'step'` and `conf_int` is False. By default
        {"fill_alpha": 0.3, "line_alpha": 0}.
    rug_kwargs : dict
        Keyword arguments to pass to `p.multi_line()` when making the
        rug plot.
    horizontal : bool or None, default None
        Deprecated. Use `q_axis`.
    val : hashable
        Deprecated, use `q`.
    click_policy : str, default 'hide'
        Deprecated. Use `legend_click_policy`.
    conf_int_kwargs : dict
        Deprecated. Use `fill_kwargs`.
    kind : str, default 'step_filled'
        Deprecated. Use `style`.
    kwargs
        Any kwargs to be passed to `bokeh.plotting.figure()` when making
        the plot.
    Returns
    -------
    output : Bokeh figure
        Figure populated with histograms.
    Notes
    -----
    .. Confidence intervals for the histogram are computed using
       nonparametric bootstrap as follows. The bins are established as
       per user input via the `bins` kwarg. These bins are fixed for all
       bootstrap replicates. Then, for each bootstrap sample drawn, the
       histogram is computed for the bins. The confidence interval is
       then computed from these bootstrap samples.
    """
    # Protect against mutability of dicts
    line_kwargs = copy.copy(line_kwargs)
    fill_kwargs = copy.copy(fill_kwargs)
    rug_kwargs = copy.copy(rug_kwargs)

    # Check the deprecation of `kind` kwarg (not in utils._parse_deprecations because
    # `kind` is a valid kwarg for ECDFs)
    if kind is not None:
        if style != kind:
            raise RuntimeError(
                "`kind` and `style` in disagreement. Use `style`; `kind` is deprecated."
            )
        warnings.warn(
            f"`kind` is deprecated. Use `style`. Using style='{style}'.",
            DeprecationWarning,
        )

    if style is None:
        if conf_int:
            style = "step"
        else:
            if arrangement == "stack":
                style = "step_filled"
            else:
                style = "step"

    if density is None:
        if arrangement == "stack":
            density = True
        else:
            density = False

    if arrangement == "stack":
        if rug_height is None:
            rug_height = 0.2
        if not density:
            raise RuntimeError("Must have density = True for 'stack' arrangement.")
    elif arrangement != "overlay":
        raise RuntimeError(
            "Only allowed values for `arrangement` are 'stack' and 'overlay'."
        )
    elif rug_height is None:
        rug_height = 0.05

    if style == "step_filled" and conf_int:
        raise RuntimeError(
            "`style` must be 'step' when confidence intervals are displayed."
        )

    if conf_int and style == "step_filled":
        raise RuntimeError(
            f"`style` must be 'step' when confidence intervals are included."
        )

    q, legend_click_policy, fill_kwargs = _parse_deprecations(
        q,
        q_axis,
        val,
        horizontal,
        "y",
        click_policy,
        legend_click_policy,
        conf_int_kwargs,
        fill_kwargs,
    )

    if type(bins) == str and bins in ["integer", "exact"]:
        if rug is None:
            rug = False

        if rug:
            warnings.warn("Rug plot not generated for integer or exact bins.")
            rug = False
    elif rug is None:
        rug = True

    if palette is None:
        palette = colorcet.b_glasbey_category10
    elif type(palette) == str:
        palette = [palette]

    df, q, cats, show_legend = _data_cats(
        data, q, cats, show_legend, legend_label
    )

    if arrangement == "stack":
        if show_legend is None:
            show_legend = False

        if show_legend:
            warnings.warn(
                "Cannot show legend with arrangement='stack'. There is no legend to show."
            )

    if show_legend is None:
        if cats is None:
            show_legend = False
        else:
            show_legend = True

    if type(bins) == str and bins not in [
        "integer",
        "exact",
        "sqrt",
        "freedman-diaconis",
    ]:
        raise RuntimeError("Invalid bin specification.")

    if cats is None:
        df["__cat"] = "__dummy_cat"
        if show_legend:
            raise RuntimeError("No legend to show if `cats` is None.")
        if order is not None:
            raise RuntimeError("No `order` is allowed if `cats` is None.")
        cats = "__cat"

    cats, cols = _check_cat_input(
        df, cats, q, None, None, None, palette, order, kwargs
    )

    # Defaults for histogram
    if line_kwargs is None:
        line_kwargs = {"line_width": 2}
    if fill_kwargs is None:
        fill_kwargs = {}
    if "fill_alpha" not in fill_kwargs:
        fill_kwargs["fill_alpha"] = 0.3
    if "line_alpha" not in fill_kwargs:
        fill_kwargs["line_alpha"] = 0

    # Defaults for rug_kwargs
    if rug_kwargs is None:
        rug_kwargs = dict(alpha=0.5, line_width=0.5)
    elif type(rug_kwargs) != dict:
        raise RuntimeError("`rug_kwargs` must be a dictionary.")
    if "alpha" not in rug_kwargs and "line_alpha" not in rug_kwargs:
        rug_kwargs["alpha"] = 0.5
    if "line_width" not in rug_kwargs:
        rug_kwargs["line_width"] = 0.5
    if "name" not in rug_kwargs:
        rug_kwargs["name"] = "hover_glyphs"

    _, df["__label"] = _source_and_labels_from_cats(df, cats)
    cols += ["__label"]

    df = _sort_df(df, cats, order)

    if type(bins) == str and bins == "exact":
        a = np.unique(df[q])
        if len(a) == 1:
            bins = np.array([a[0] - 0.5, a[0] + 0.5])
        else:
            bins = np.concatenate(
                (
                    (a[0] - (a[1] - a[0]) / 2,),
                    (a[1:] + a[:-1]) / 2,
                    (a[-1] + (a[-1] - a[-2]) / 2,),
                )
            )
    elif type(bins) == str and bins == "integer":
        if np.any(df[q] != np.round(df[q])):
            raise RuntimeError("'integer' bins chosen, but data are not integer.")
        bins = np.arange(df[q].min() - 1, df[q].max() + 1) + 0.5

    if arrangement == "stack":
        if cats is not None:
            grouped = df.groupby(cats, sort=False)
            return _stacked_histograms(
                df,
                grouped,
                q,
                bins,
                density,
                palette,
                q_axis,
                order,
                p,
                mirror,
                hist_height,
                style,
                conf_int,
                ptiles,
                n_bs_reps,
                rug,
                rug_height,
                tooltips,
                line_kwargs,
                fill_kwargs,
                rug_kwargs,
                kwargs,
            )

    if p is None:
        kwargs = _fig_dimensions(kwargs)

        if "x_axis_label" not in kwargs:
            if q_axis == "y":
                if density:
                    kwargs["x_axis_label"] = "density"
                else:
                    kwargs["x_axis_label"] = "count"
            else:
                kwargs["x_axis_label"] = q

        if "y_axis_label" not in kwargs:
            if q_axis == "y":
                kwargs["y_axis_label"] = q
            else:
                if density:
                    kwargs["y_axis_label"] = "density"
                else:
                    kwargs["y_axis_label"] = "count"

        if q_axis == "y":
            if "x_range" not in kwargs:
                kwargs["x_range"] = bokeh.models.DataRange1d(start=0)
        else:
            if "y_range" not in kwargs:
                kwargs["y_range"] = bokeh.models.DataRange1d(start=0)

        p = bokeh.plotting.figure(**kwargs)

    # Explicitly loop to enable click policies on the legend (not possible with factors)
    max_height = 0
    lines = []
    labels = []
    patches = []
    for i, (name, g) in enumerate(df.groupby(cats, sort=False)):
        numerical_bins, e, f = _compute_histogram(g[q], bins, density)
        e0, f0 = _hist_for_plotting(e, f)

        if conf_int:
            fill_kwargs["fill_color"] = palette[i % len(palette)]

            q_vals = g[q].values
            f_reps = np.empty((n_bs_reps, len(f)))
            for j in range(n_bs_reps):
                _, _, f_reps[j, :] = _compute_histogram(
                    np.random.choice(q_vals, len(q_vals)), numerical_bins, density
                )

            f_ptiles = np.percentile(f_reps, [2.5, 97.5], axis=0)

            _, f0_low = _hist_for_plotting(e, f_ptiles[0, :])
            _, f0_high = _hist_for_plotting(e, f_ptiles[1, :])
            if q_axis == "y":
                p, patch = _fill_between(
                    p, f0_low, e0, f0_high, e0, **fill_kwargs
                )
            else:
                p, patch = _fill_between(
                    p, e0, f0_low, e0, f0_high, **fill_kwargs
                )
            patches.append(patch)

        max_height = max(f0.max(), max_height)

        line_kwargs["color"] = palette[i % len(palette)]

        if q_axis == "y":
            lines.append(p.line(f0, e0, **line_kwargs))
        else:
            lines.append(p.line(e0, f0, **line_kwargs))
        labels.append(g["__label"].iloc[0])

        if style == "step_filled":
            x2 = [e0.min(), e0.max()]
            y2 = [0, 0]
            fill_kwargs["color"] = palette[i % len(palette)]
            if q_axis == "y":
                p, patch = _fill_between(p, f0, e0, y2, x2, **fill_kwargs)
            else:
                p, patch = _fill_between(p, e0, f0, x2, y2, **fill_kwargs)
            patches.append(patch)

    # Put in the rug plot
    if rug:
        y = [0, max_height * rug_height]

        for i, (name, g) in enumerate(df.groupby(cats, sort=False)):
            xs = [[q_val, q_val] for q_val in g[q].values]
            ys = [y] * len(g)

            if q_axis == "y":
                xs, ys = ys, xs

            cds = bokeh.models.ColumnDataSource(g)
            cds.data["__xs"] = xs
            cds.data["__ys"] = ys

            if "color" not in rug_kwargs and "line_color" not in rug_kwargs:
                p.multi_line(
                    source=cds,
                    xs="__xs",
                    ys="__ys",
                    line_color=palette[i % len(palette)],
                    **rug_kwargs,
                )
            else:
                p.multi_line(source=cds, xs="__xs", ys="__ys", **rug_kwargs)

    if tooltips is not None:
        p.add_tools(bokeh.models.HoverTool(tooltips=tooltips, names=["hover_glyphs"]))

    return _dist_legend(
        p,
        show_legend,
        legend_location,
        legend_orientation,
        legend_click_policy,
        labels,
        [],
        lines,
        patches,
        [],
        [],
        [],
        [],
        [],
    )
```
