## Supplemental data and code for "Feature Reuse and Scaling: Understanding Transfer Learning with Protein Language Models": learning_vis.py

```
"""Script for visulizing the learning related process"""

from __future__ import annotations

import os
import numpy as np
import matplotlib.pyplot as plt

from scr.utils import get_folder_file_names, get_task_data_split


def plot_lc(
    train_losses: np.ndarray,
    val_losses: np.ndarray,
    dataset_path: str,
    encoder_name: str,
    embed_layer: int,
    flatten_emb: bool | str,
    all_plot_folder: str = "results/learning_curves",
) -> None:
    """
    Plot and save learning curves

    Args:
    - train_losses: np.ndarry, an array of training loss
    - val_losses: np.ndarry, an array of validation loss
    - dataset_path: str, full path to the dataset, in pkl or panda readable format
        columns include: sequence, target, set, validation, mut_name (optional), mut_numb (optional)
    - encoder_name: str, the name of the encoder
    - embed_layer: int, the layer number of the embedding
    - flatten_emb: bool or str, if and how (one of ["max", "mean"]) to flatten the embedding
    - all_plot_folder: str, the parent folder path for saving all the learning curves
    """

    epochs = len(train_losses)

    plot_dataset_folder, plotname = get_folder_file_names(
        parent_folder=all_plot_folder,
        dataset_path=dataset_path,
        encoder_name=encoder_name,
        embed_layer=embed_layer,
        flatten_emb=flatten_emb,
    )

    task, dataset, split = get_task_data_split(dataset_path=dataset_path)

    plt.figure()
    plt.plot(range(epochs), train_losses, label="train")
    plt.plot(range(epochs), val_losses, label="val")
    plt.title(f"{task}_{dataset}_{split} \n {plotname}")
    plt.xlabel("epochs")
    plt.ylabel("loss")
    plt.legend(loc="upper right")

    # TODO: split the encoder and flatten to be subfolders

    plt.savefig(
        os.path.join(plot_dataset_folder, plotname + ".png"), bbox_inches="tight"
    )
    plt.close()
```
