## Supplemental data and code for "Feature Reuse and Scaling: Understanding Transfer Learning with Protein Language Models": res_vis.py

```
# a script for vis results from summary data df
from __future__ import annotations

import ast

import os

import math

import numpy as np
import pandas as pd

import matplotlib as mpl
import matplotlib.pyplot as plt
from matplotlib.lines import Line2D
from matplotlib.patches import Rectangle
from matplotlib.gridspec import GridSpec

import seaborn as sns
import holoviews as hv
from holoviews import dim

hv.extension("bokeh")

from scr.vis.vis_utils import BokehSave, save_plt
from scr.params.emb import (
    MODEL_SIZE,
    EMB_MODEL_SIZE,
    MODEL_LAYER,
    EMB_MODEL_LAYER,
    CARP_MODEL_LAYER,
    CARP_INFO,
    ARCH_TYPE,
    ARCH_BAR_LAYER,
    ARCH_AB,
    ARCH_AB_DICT,
    CHECKPOINT_PERCENT,
    CARP_CHECKPOINT_LOSSES,
)
from scr.params.vis import (
    ORDERED_TASK_LIST,
    TASK_LEGEND_MAP,
    TASK_COLORS,
    TASK_SIMPLE_COLOR_MAP,
    PLOT_EXTS,
    CARP_ALPHA,
    ARCH_LINE_STYLE_DICT,
    ARCH_DOT_STYLE_DICT,
    ARCH_AB_DOT_STYLE_DICT,
    ARCH_SCATTER_STYLE_DICT,
    LAYER_ALPHAS,
)
from scr.analysis.utils import INIT_DICT, INIT_SIMPLE_LIST, SIMPLE_METRIC_LIST
from scr.utils import checkNgen_folder


class PlotResultScatter:
    """
    A class handling plotting results in scatter plots, including:
    - PlotLayerDelta
    - PlotEmbvsOnehot
    """

    def __init__(
        self,
        sum_folder: str = "results/summary",
        sum_df_name: str = "all_results",
    ) -> None:

        self._sum_folder = checkNgen_folder(os.path.normpath(sum_folder))
        self._sum_df_name = sum_df_name

    def _get_best_metric_df_dicts(self) -> list[dict]:

        """
        A method for spliting the df into dict based on the best metric

        'all' means best within emb and onehot
        'emb' means best slicing out emb from all
        'carp' or 'esm' means best given arch
        """

        best_metric_df_dict = {"all": {}, "emb": {}}

        for arch in ARCH_TYPE:
            best_metric_df_dict[arch] = {}

        for m in SIMPLE_METRIC_LIST:

            slice_df = get_bestorlast_metric_df(df=self.prepped_df.copy(), metric=m)

            best_metric_df_dict["all"][m] = slice_df
            best_metric_df_dict["emb"][m] = slice_df[
                slice_df["ablation"] == "emb"
            ].copy()

            for arch in ARCH_TYPE:
                arch_df = get_bestorlast_metric_df(
                    df=self.prepped_df.copy(), metric=m, arch=arch
                )
                best_metric_df_dict[arch][m] = arch_df[
                    arch_df["ablation"] == "emb"
                ].copy()

        return best_metric_df_dict

    def _get_last_metric_df_dict(self) -> list[dict]:

        """
        A method for spliting the df into dict gettin last layer performance
        based on the metric

        'all' means best within emb and onehot
        'emb' means best slicing out emb from all
        'carp' or 'esm' means best given arch
        """

        last_metric_dict = {}

        for arch in ARCH_TYPE + [""]:

            last_metric_dict[arch] = {}

        for m in SIMPLE_METRIC_LIST:

            for arch in ARCH_TYPE + [""]:
                last_metric_dict[arch][m] = get_bestorlast_metric_df(
                    df=self.prepped_df.copy(), metric=m, arch=arch, bestorlast="last"
                )

        return last_metric_dict

    def _get_last_layer_bar_df(
        self, metric: str, ablation: str = "emb", layers: list[int] = []
    ) -> list[pd.DataFrame, pd.DataFrame, list[str]]:

        """
        A method for slicing df with last layer info for bar plots
        """

        df = get_bestorlast_metric_df(
            df=self.prepped_df,
            metric=metric,
            arch="",
            bestorlast="last",
            ablation=ablation,
            layers=layers,
        )

        get_val_col = [v for v in df.columns if "_value" in v]

        # do not consider diff deg of pretrain just yet in this case
        df = df[df["ptp"].isin([0, 1])][
            [
                "arch",
                "task",
                "model",
                "ablation",
                "model_size",
                "model_layer",
                *get_val_col,
            ]
        ]

        # prep clean emb_df
        emb_df = df[df["ablation"] == ablation].reset_index(drop=True)
        # Convert 'model' to categorical with custom order
        emb_df["model"] = pd.Categorical(
            emb_df["model"], categories=list(EMB_MODEL_SIZE.keys()), ordered=True
        )

        # Sort the DataFrame based on the custom order in 'model'
        emb_df = emb_df.sort_values(by=["task", "model"]).reset_index(drop=True)

        # prep clean emb_df
        onehot_df = df[df["ablation"] == "onehot"].reset_index(drop=True)
        # onehot should be same for either arch
        onehot_df = onehot_df.drop("arch", axis=1)
        # Drop duplicate rows based on all columns
        onehot_df = onehot_df.drop_duplicates().reset_index(drop=True)

        return emb_df, onehot_df, sorted(get_val_col.copy())

    def _get_df_for_best_layer(self, metric: str, arch: str = "") -> pd.DataFrame:

        """
        A method for getting best layer df

        Args:
        - arch: str = "", defalt for esm and carp with out ptp, if carp keep ptp
        """

        # plot the best layer achieved with
        best_layer_append = get_bestorlast_metric_df(
            df=self.prepped_df,
            metric=metric,
            arch="",
            bestorlast="best",
            ifgroupbestbytask=False,
            ablation="emb",
            layers=[],
        )

        # drop onehot
        best_layer_append = best_layer_append[best_layer_append["ablation"] == "emb"]

        # drop ptp if for esm or both
        if arch == "carp":
            best_layer_append = best_layer_append[best_layer_append["arch"] == arch]
        else:
            best_layer_append = best_layer_append[best_layer_append["ptp"].isin([0, 1])]

        # Convert 'model' to categorical with custom order
        best_layer_append["model"] = pd.Categorical(
            best_layer_append["model"],
            categories=list(EMB_MODEL_SIZE.keys()),
            ordered=True,
        )

        # Sort the DataFrame based on the custom order in 'model'
        return best_layer_append.sort_values(by=["task", "model"]).reset_index(
            drop=True
        )

    def _append_randstat(self, emb_df: pd.DataFrame) -> pd.DataFrame:

        """
        A method for adding rand stat and delta onehot info to
        best performance based on given metric
        """

        # add rand stat
        for ab in INIT_SIMPLE_LIST:
            emb_df[ab] = np.nan
            emb_df[f"{ab} - onehot"] = np.nan

        emb_df["emb - onehot"] = np.nan

        emb_df = emb_df.reset_index(drop=True)

        # get the corresponding stat and rand value at the layer
        for i, row in emb_df.iterrows():

            # Convert the row to a dictionary
            row_dict = row.to_dict()

            # Select certain keys from the dictionary
            selected_keys = ["arch", "task", "metric", "model"]

            # pick the row to match
            row_to_match = {key: row_dict[key] for key in selected_keys}

            # get the onehot baseline
            onehot_val = get_layer_value(
                df=self.prepped_df,
                row_to_match={
                    key: row_dict[key] if i < 3 else "onehot"
                    for i, key in enumerate(selected_keys)
                },
                ablation="onehot",
                layer_numb=0,
            )

            # calc best emb perf del
            emb_df.at[i, "emb - onehot"] = emb_df.at[i, "best_value"] - onehot_val

            for ab in INIT_SIMPLE_LIST:

                randstat_val = get_layer_value(
                    df=self.prepped_df,
                    row_to_match=row_to_match,
                    ablation=ab,
                    layer_numb=row_dict["best_layer"],
                )

                emb_df.at[i, ab] = randstat_val
                emb_df.at[i, f"{ab} - onehot"] = randstat_val - onehot_val

        return emb_df

    def _append_ptp(self, emb_df: pd.DataFrame) -> pd.DataFrame:

        """A method for adding pretrain percent best performance"""

        for ptp in CHECKPOINT_PERCENT:
            emb_df[str(ptp)] = np.nan
            emb_df[f"{str(ptp)} - onehot"] = np.nan

        emb_df = emb_df.reset_index(drop=True)

        # figure out if last or best
        if "best_layer" in emb_df.columns:
            bestoflast = "best_layer"
        else:
            bestoflast = "model_layer"

        # get the corresponding stat and rand value at the layer
        for i, row in emb_df.iterrows():

            # Convert the row to a dictionary
            row_dict = row.to_dict()

            # Select certain keys from the dictionary
            selected_keys = ["arch", "task", "metric", "model"]

            # pick the row to match
            row_to_match = {key: row_dict[key] for key in selected_keys}

            # get the onehot baseline
            onehot_val = get_layer_value(
                df=self.prepped_df,
                row_to_match={
                    key: row_dict[key] if i < 3 else "onehot"
                    for i, key in enumerate(selected_keys)
                },
                ablation="onehot",
                layer_numb=0,
            )

            # add in the checkpoint
            for ptp in CHECKPOINT_PERCENT:

                statorrand_val = get_layer_value(
                    df=self.prepped_df,
                    row_to_match=row_to_match,
                    ablation="emb",
                    ptp=ptp,
                    layer_numb=row_dict[bestoflast],
                )

                emb_df.at[i, str(ptp)] = statorrand_val
                emb_df.at[i, f"{str(ptp)} - onehot"] = statorrand_val - onehot_val

        # drop the dup of carp last since all ptp info included
        if bestoflast == "model_layer":
            emb_df = (
                emb_df.drop(columns=["ptp", "value", "last_value"])
                .drop_duplicates()
                .reset_index(drop=True)
            )

        return emb_df

    def plot_emb_onehot(
        self, metric: str = "test_performance_1", layers: list[int] = [2, 4, 6]
    ) -> pd.DataFrame:

        """
        A method for getting df for emb vs onehot

        Args:
        - ifclean_metric: bool = True, simplify test performance
        """

        # now plot and save
        vs_plot = plot_emb_onehot(
            df=self.best_metric_df_dict["all"][metric],
            metric=metric,
            path2folder=checkNgen_folder(
                os.path.join(self._sum_folder, "best", "embvsonehot", metric)
            ),
        )

        # get a bar plot with percent performance achieved
        emb_df = self.best_metric_df_dict["emb"][metric]
        emb_df["model_layer_percent"] = emb_df["best_layer"] / emb_df["model_layer"]

        best_layer = plot_best_layer_bar(
            df=emb_df,
            metric=metric,
            path2folder=checkNgen_folder(
                os.path.join(self._sum_folder, "best", "bestemblayer", metric)
            ),
        )

        best_layer_all_df = self._get_df_for_best_layer(metric=metric, arch="")

        # best layer for all task and model combo
        best_layer_all = plot_emb_df_best_layer_bar(
            emb_df=best_layer_all_df,
            metric=metric,
            path2folder=checkNgen_folder(
                os.path.join(self._sum_folder, "best", "bestemblayer_all", metric)
            ),
        )

        # prep to merge for summary df
        merge_df = best_layer_all_df[["task", "model", "model_layer", "best_layer"]].copy()
        merge_df["best_layer_percent"] = (
            merge_df["best_layer"] / merge_df["model_layer"]
        )
        merge_df = merge_df.drop(columns=["model_layer"])

        best_layer_carp_df = self._get_df_for_best_layer(metric=metric, arch="carp")
        # best layer for carp with differen pretrain degree
        plot_carp_best_layer_bar(
            df=best_layer_carp_df,
            metric=metric,
            path2folder=checkNgen_folder(
                os.path.join(self._sum_folder, "best", "bestemblayer_carp", metric)
            ),
        )

        # cross pretrain loss
        for ifpercent in [0, 1]:
            plot_carp_best_layer_pretrain_loss(
                df=best_layer_carp_df,
                metric=metric,
                ifpercent=ifpercent,
                path2folder=checkNgen_folder(
                    os.path.join(
                        self._sum_folder, "best", "bestemblayer_carp_loss", metric
                    )
                ),
            )

        last_df = self.last_metric_df_dict[""][metric]

        # plot more detailed last layer emb vs onehot and save
        vs_plot_det = plot_emb_onehot_det(
            df=last_df[last_df["ptp"].isin([0, 1])],  # ignore ptp not 1 or 0
            metric=metric,
            path2folder=checkNgen_folder(
                os.path.join(self._sum_folder, "last", "embvsonehot", metric)
            ),
        )

        # plot embvsonehot in bar plots
        bar_emb_df, bar_onehot_df, _ = self._get_last_layer_bar_df(metric=metric)

        last_layer_bar = plot_emb_df_last_layer_bar(
            emb_df=bar_emb_df,
            onehot_df=bar_onehot_df,
            metric=metric,
            path2folder=checkNgen_folder(
                os.path.join(self._sum_folder, "last", "embvsonehot_bar", metric)
            ),
        )

        # get diff layer emb vs onehot
        layer_emb_df, layer_onehot_df, layer_val_col = self._get_last_layer_bar_df(
            metric=metric, ablation="emb", layers=ARCH_BAR_LAYER
        )

        # minue onehot
        layer_emb_delta_df = subtract_onehot(
            df=layer_emb_df.copy(),
            onehot_df=layer_onehot_df.copy(),
            val_col=layer_val_col,
        ).copy()

        # append to merge for summary df
        merge_df = pd.merge(
            merge_df.copy(),
            layer_emb_delta_df[["task", "model", "last_value"]].rename(
                columns={"last_value": "Emb > OH"}
            ).copy(),
            on=["task", "model"],
            how="outer",
        )

        plot_emb_layer_bar(
            df=layer_emb_delta_df,
            val_col=layer_val_col,
            metric=metric,
            ifratio=False,
            path2folder=checkNgen_folder(
                os.path.join(self._sum_folder, "last", "emblayersvsonehot_bar", metric)
            ),
        )

        # dict for rand stat df before subtract onehot
        layer_ab_df_dict = {}

        # dict for rand stat df after subtract onehot
        layer_ab_delta_df_dict = {}

        # now do rand and stat
        for ablation in ARCH_AB:
            # get rand stat bar with layers
            ab_df, _, _ = self._get_last_layer_bar_df(
                metric=metric, layers=ARCH_BAR_LAYER, ablation=ablation
            )
            layer_ab_df_dict[ablation] = ab_df

            layer_ab_delta_df_dict[ablation] = subtract_onehot(
                ab_df, layer_onehot_df, layer_val_col
            )

            for r in [False, True]:

                if r:
                    df = take_ratio(
                        ablation_df=layer_ab_delta_df_dict[ablation].copy(),
                        emb_df=layer_emb_delta_df.copy(),
                        val_col=layer_val_col,
                    )
                else:
                    df = layer_ab_delta_df_dict[ablation].copy()

                plot_emb_layer_bar(
                    df=df,
                    val_col=layer_val_col,
                    metric=metric,
                    ifratio=r,
                    path2folder=checkNgen_folder(
                        os.path.join(
                            self._sum_folder, "last", "emblayersvsonehot_bar", metric
                        )
                    ),
                )

        randstat_col = [f"{a}_value" for a in ARCH_AB]

        # do randstat vs onehot with or without emb ratio
        ablation_onehot_df = merge_rand_stat(
            rand_df=layer_ab_delta_df_dict[ARCH_AB[0]].copy(),
            stat_df=layer_ab_delta_df_dict[ARCH_AB[1]].copy(),
            onehot_df=layer_onehot_df.copy(),
            val_col=layer_val_col,
            ifdelta=True,
        ).copy()

        for (df, r) in zip(
            [
                take_ratio(
                    ablation_df=ablation_onehot_df,
                    emb_df=merge_rand_stat(
                        rand_df=layer_emb_delta_df.copy(),
                        stat_df=layer_emb_delta_df.copy(),
                        onehot_df=layer_onehot_df.copy(),
                        val_col=layer_val_col,
                        ifdelta=True,
                    ).copy(),
                    val_col=randstat_col,
                ),
                ablation_onehot_df,
            ],
            [True, False],
        ):

            # now do the rand stat combined simpler one
            plot_emb_layer_bar(
                df=df,
                val_col=randstat_col,
                metric=metric,
                iflast=True,
                ifratio=r,
                path2folder=checkNgen_folder(
                    os.path.join(
                        self._sum_folder, "last", "emblayersvsonehot_bar", metric
                    )
                ),
            )

        # append simplified df for summary df
        merge_df = pd.merge(
            merge_df.copy(),
            ablation_onehot_df[["task", "model", "rand_value", "stat_value"]].rename(
                columns={"rand_value": "RI > OH", "stat_value": "ST > OH"}
            ).copy(),
            on=["task", "model"],
            how="outer",
        )

        # just rand and stat vs emb (no onehot)
        sel_col = [
            "arch",
            "task",
            "ablation",
            "model",
            "model_size",
            "model_layer",
            "last_value",
        ]

        for ab in ARCH_AB:

            plot_emb_layer_bar(
                df=subtract_emb(
                    layer_ab_df_dict[ab], layer_emb_df, val_col=layer_val_col
                ),
                val_col=layer_val_col,
                metric=metric,
                iflast=False,
                ifratio=False,
                ifemb=True,
                path2folder=checkNgen_folder(
                    os.path.join(self._sum_folder, "last", f"{ab}vsemb_bar", metric)
                ),
            )

        ablation_emb_df = merge_rand_stat(
            rand_df=subtract_emb(
                layer_ab_df_dict["rand"], layer_emb_df, val_col=["last_value"]
            ),
            stat_df=subtract_emb(
                layer_ab_df_dict["stat"], layer_emb_df, val_col=["last_value"]
            ),
            onehot_df=layer_emb_df[sel_col],
            val_col=["last_value"],
            ifdelta=True,
        ).copy()

        # merge but only last
        plot_emb_layer_bar(
            df=ablation_emb_df,
            val_col=["rand_value", "stat_value"],
            metric=metric,
            iflast=True,
            ifratio=False,
            ifemb=True,
            path2folder=checkNgen_folder(
                os.path.join(self._sum_folder, "last", "randstatvsemb_bar", metric)
            ),
        )

        # append simplified df for summary df
        merge_df = pd.merge(
            merge_df.copy(),
            ablation_emb_df[["task", "model", "rand_value", "stat_value"]].rename(
                columns={"rand_value": "RI > Emb", "stat_value": "ST > Emb"}
            ).copy(),
            on=["task", "model"],
            how="outer",
        )

        # get rand - stat
        rand_stat = subtract_emb(
                layer_ab_df_dict["rand"], layer_ab_df_dict["stat"], val_col=["last_value"]
            ).copy()

        merge_df = pd.merge(
            merge_df.copy(),
            rand_stat[["task", "model", "last_value"]].rename(
                columns={"last_value": "RI > ST"}
            ).copy(),
            on=["task", "model"],
            how="outer",
        )

        # add rand stat info to emb_df
        emb_df = self._append_randstat(emb_df=emb_df)

        randstat_plot_dict = {}

        for delta_onehot in [0, 1]:
            randstat_plot_dict[delta_onehot] = {}

            for randstat in INIT_SIMPLE_LIST + [""]:

                randstat_plot_dict[delta_onehot][randstat] = plot_randstat(
                    df=emb_df,
                    metric=metric,
                    randstat=randstat,
                    delta_onehot=delta_onehot,
                    path2folder=checkNgen_folder(
                        os.path.join(self._sum_folder, "best", "randstat", metric)
                    ),
                )

        # process df to be simpler and save
        simpler_df = simplify_sum_delta_df(df=merge_df)
        simpler_df.to_csv(
            os.path.join(
                checkNgen_folder(
                    os.path.join(self._sum_folder, "all_results_last_layer", metric)
                ),
                "last_layer_delta_summary.csv",
            )
        )

        return (
            vs_plot,
            best_layer,
            best_layer_all,
            vs_plot_det,
            last_layer_bar,
            merge_df,
        )

    def plot_layer_delta(
        self,
        layer_cut: int,
        metric: str = "test_performance_1",
        arch: str = "esm",
        ifsimple: bool = True,
    ) -> pd.DataFrame:

        """
        A method for getting the sliced dataframe

        Add per-train degree as a col

        Args:
        - ifsimple: bool = True, if selecting the best
        """

        assert arch in ARCH_TYPE, f"{arch} not in {ARCH_TYPE}"

        df = self.prepped_df.copy()

        # get rid of onehot
        df = df[df["ablation"] == "emb"].copy()

        if ifsimple or arch is None:

            slice_df = get_bestorlast_metric_df(df=df, metric=metric)

            # Apply the function and generate two new columns
            slice_df["x-0"], slice_df["f-x"] = zip(
                *slice_df.apply(
                    lambda row: delta_layer(
                        layer_cut=layer_cut, value_array=row["value"]
                    ),
                    axis=1,
                )
            )

            # now plot and save
            delta_plot = plot_layer_delta_simple(
                df=slice_df,
                layer_cut=layer_cut,
                metric=metric,
                path2folder=checkNgen_folder(
                    os.path.join(self._sum_folder, "best", "layerdelta_simple", metric)
                ),
            )

            return slice_df, delta_plot

        else:

            slice_df = df[(df["metric"] == metric) & (df["arch"] == arch)].copy()

            # Apply the function and generate two new columns
            slice_df["x-0"], slice_df["f-x"] = zip(
                *slice_df.apply(
                    lambda row: delta_layer(
                        layer_cut=layer_cut, value_array=row["value"]
                    ),
                    axis=1,
                )
            )

            # now plot and save
            delta_plot = plot_layer_delta_det(
                df=slice_df,
                layer_cut=layer_cut,
                arch=arch,
                metric=metric,
                path2folder=checkNgen_folder(
                    os.path.join(self._sum_folder, "best", "layerdelta_det", metric)
                ),
            )

        return slice_df, delta_plot

    def plot_pretrain_degree(
        self,
        metric: str = "test_performance_1",
        arch: str = "carp",
        ifloss: bool = True,
    ):

        """
        A method for plotting the pretraining arch

        Args:
        - ifloss: bool = True, over loss or pretrain degree
        """

        for (bestorlast, df) in zip(
            ["best", "last"],
            [
                self.best_metric_df_dict[arch][metric],
                self.last_metric_df_dict[arch][metric],
            ],
        ):

            for delta_onehot in [0, 1]:
                for ifloss in [0, 1]:
                    plot_pretrain_degree(
                        emb_df=self._append_ptp(df[df["ablation"] == "emb"].copy()),
                        metric=metric,
                        arch=arch,
                        delta_onehot=delta_onehot,
                        ifloss=ifloss,
                        path2folder=checkNgen_folder(
                            os.path.join(
                                self._sum_folder,
                                bestorlast,
                                "pretraindegree",
                                metric,
                                arch,
                            )
                        ),
                    )

    def plot_arch_size(
        self,
        metric: str = "test_performance_1",
    ):

        """A method for plotting arch size"""

        for delta_onehot in [0, 1]:
            for arch in ARCH_TYPE + [""]:
                plot_arch_size(
                    arch_df_dict=self.last_metric_df_dict,
                    metric=metric,
                    arch=arch,
                    delta_onehot=delta_onehot,
                    path2folder=checkNgen_folder(
                        os.path.join(self._sum_folder, "archsize", metric, arch)
                    ),
                )

    @property
    def result_df_path(self) -> str:
        """Return full summary result csv path"""
        df_path = os.path.join(
            os.path.normpath(self._sum_folder), self._sum_df_name + ".csv"
        )

        assert os.path.exists(df_path), f"{df_path} does not exist"

        return df_path

    @property
    def result_df(self) -> pd.DataFrame:

        """Return full result df with value cleaned up"""

        result_df = pd.read_csv(self.result_df_path)

        # check column name existance
        for c in ["metric", "ablation", "arch", "value", "task", "model", "ptp"]:
            assert c in result_df.columns, f"{c} not in df from {self.result_df_path}"

        # Convert the string of lists to NumPy arrays
        result_df["value"] = result_df["value"].apply(ast.literal_eval).apply(np.array)

        # make ptp float
        result_df["ptp"] = result_df["ptp"].astype(float)

        return result_df

    @property
    def prepped_df(self) -> pd.DataFrame:

        """Return a more plotting compatible df"""

        prepped_df = self.result_df.copy()

        # add task type and model size details for plotting legends
        prepped_df["task_type"] = prepped_df["task"].str.split("_").str[0]
        prepped_df["model_size"] = prepped_df["model"].map(MODEL_SIZE)
        prepped_df["model_layer"] = prepped_df["model"].map(MODEL_LAYER)

        # get rid of pooling details
        prepped_df["task"] = prepped_df["task"].str.replace("_mean", "")
        prepped_df["task"] = prepped_df["task"].str.replace("_noflatten", "")

        # sort based on given task order for plot legend
        prepped_df["task"] = pd.Categorical(
            prepped_df["task"], categories=ORDERED_TASK_LIST, ordered=True
        ).map(TASK_LEGEND_MAP)
        prepped_df = prepped_df.sort_values(["task", "ptp"], ascending=[True, False])

        return prepped_df

    @property
    def best_metric_df_dict(self) -> dict:
        """Return a dict splicing the df with best metric based on best metric"""
        return self._get_best_metric_df_dicts()

    @property
    def last_metric_df_dict(self) -> dict:
        """Return a dict splicin the df with the last layer value given metric"""
        return self._get_last_metric_df_dict()


def simplify_sum_delta_df(df: pd.DataFrame) -> pd.DataFrame:

    """
    A function to simply the last layer df
    """

    # Applying the rules to the respective columns
    for col in ["Emb > OH", "RI > OH", "ST > OH", "RI > ST"]:
        df[col] = df[col].apply(lambda value: True if value > 0 else False)

    # flip order
    for (col, ncol) in zip(["RI > Emb", "ST > Emb"], ["Emb > RI", "Emb > ST"]):
        df[ncol] = df[col].apply(lambda value: False if value > 0 else True)
        df = df.drop(columns=col)

    return df


def simplify_test_metric(metric: str) -> str:

    """
    A function to unify metric for plotting
    """

    for t in ["test_performance_1", "test_performance_2", "test_loss"]:
        metric = metric.replace(t, "test performance")

    return metric


def get_bestorlast_metric_df(
    df: pd.DataFrame,
    metric: str = "test_performance_1",
    arch: str = "",
    bestorlast: str = "best",
    ifgroupbestbytask: bool = True,
    ablation: str = "emb",
    layers: list[int] = [],
) -> pd.DataFrame:

    """
    A function for cleaning up the df to get best layer based on chosen metric

    Args:
    - ifgroupbestbytask: bool: True, if best of each task
    - layers: list[int] = [], a list of ints for what other layers to extract
    """

    slice_df = df[(df["metric"] == metric)].copy()

    slice_df = slice_df[
        (slice_df["ablation"] == "onehot") | (slice_df["ablation"] == ablation)
    ]

    # comb carp and esm
    if arch != "":
        slice_df = slice_df[(slice_df["arch"] == arch)].copy()

    if bestorlast == "best":

        if metric == "test_loss":
            # get the max perform layer
            slice_df["best_value"] = slice_df["value"].apply(np.min)
            slice_df["best_layer"] = slice_df["value"].apply(np.argmin)

            if ifgroupbestbytask:
                # Find the index of the maximum value in 'value_column' for each group
                min_indices = (
                    slice_df.groupby(["task", "ablation"])["best_value"]
                    .idxmin()
                    .dropna()
                )

                # Use loc to select the rows corresponding to the max indices
                slice_df = slice_df.loc[min_indices]

        else:
            # get the max perform layer
            slice_df["best_value"] = slice_df["value"].apply(np.max)
            slice_df["best_layer"] = slice_df["value"].apply(np.argmax)

            if ifgroupbestbytask:

                # Find the index of the maximum value in 'value_column' for each group
                max_indices = (
                    slice_df.groupby(["task", "ablation"])["best_value"]
                    .idxmax()
                    .dropna()
                )

                # Use loc to select the rows corresponding to the max indices
                slice_df = slice_df.loc[max_indices]

    elif bestorlast == "last":

        # get last layer
        slice_df["last_value"] = slice_df["value"].apply(
            lambda x: x[-1] if len(x) > 0 else None
        )

        if len(layers) != 0:
            for l in layers:
                # get last layer
                slice_df[f"{l}_value"] = slice_df["value"].apply(
                    lambda x: x[l] if len(x) > l else None
                )

    else:
        print(f"{bestorlast} is not 'best' or 'last'")

    return slice_df.copy()


def subtract_onehot(df: pd.DataFrame, onehot_df: pd.DataFrame, val_col: list[str]):

    """A function for subtracting onehot from emb vals"""

    # Merge dataframes on common columns
    merged_df = pd.merge(
        df,
        onehot_df[["task", "last_value"]].rename(
            columns={"last_value": "onehot_value"}
        ),
        on=["task"],
        how="left",
    )

    # Perform the desired subtraction for all val col

    for c in val_col:
        merged_df[c] = merged_df[c] - merged_df["onehot_value"]

    # Drop the onehot val columns
    return merged_df.drop(columns=["onehot_value"])


def subtract_emb(df: pd.DataFrame, emb_df: pd.DataFrame, val_col: list[str]):

    """A function for subtracting onehot from emb vals"""

    # Merge dataframes on common columns
    merged_df = df.copy()[
        ["arch", "task", "model", "ablation", "model_size", "model_layer"] + val_col
    ]

    for c in val_col:
        merged_df[c] = df[c] - emb_df[c]

    # Drop the onehot val columns
    return merged_df


def take_ratio(ablation_df: pd.DataFrame, emb_df: pd.DataFrame, val_col: list[str]):
    """
    Return df ratio
    """
    ablation_df[val_col] = ablation_df[val_col] / emb_df[val_col]
    return ablation_df


def merge_rand_stat(
    rand_df: pd.DataFrame,
    stat_df: pd.DataFrame,
    onehot_df: pd.DataFrame,
    val_col: list[str],
    ifdelta: bool = True,
) -> pd.DataFrame:
    """
    A function merge rand and stat df subtract onehot

    Args:
    - ifdelta: bool = True, if the input dataframe already gone through `subtract_onehot`
    """
    if ifdelta:
        return pd.merge(
            rand_df.drop(columns=[v for v in val_col if "last" not in v])
            .rename(columns={"last_value": "rand_value"})
            .drop(columns=["ablation"]),
            stat_df.drop(columns=[v for v in val_col if "last" not in v])
            .rename(columns={"last_value": "stat_value"})
            .drop(columns=["ablation"]),
            on=["arch", "task", "model", "model_size", "model_layer"],
            how="left",
        )
    else:
        return pd.merge(
            subtract_onehot(rand_df, onehot_df, ["last_value"])
            .drop(columns=[v for v in val_col if "last" not in v])
            .rename(columns={"last_value": "rand_value"})
            .drop(columns=["ablation"]),
            subtract_onehot(stat_df, onehot_df, ["last_value"])
            .drop(columns=[v for v in val_col if "last" not in v])
            .rename(columns={"last_value": "stat_value"})
            .drop(columns=["ablation"]),
            on=["arch", "task", "model", "model_size", "model_layer"],
            how="left",
        )


def get_layer_value(
    df: pd.DataFrame,
    row_to_match: dict,
    ablation: str,
    layer_numb: int,
    ptp: float = -1,
) -> float:

    """
    A function to get value of a given layer and other specifics

    Args:
    - ptp: float = -1, include ptp only when ptp not default -1
    """

    row_to_match["ablation"] = ablation

    # overwrite default
    if ptp > -1:
        row_to_match["ptp"] = ptp

    # Create a boolean mask for each condition
    conditions = [df[col] == value for col, value in row_to_match.items()]

    # Combine the conditions with AND (use np.all)
    mask = np.all(conditions, axis=0)

    # Use the mask to select the matching row(s)
    matching_rows = df[mask]

    assert len(matching_rows) == 1, f"{matching_rows} len not 1!"

    return matching_rows["value"].to_numpy()[0][int(layer_numb)]


def delta_layer(layer_cut: int, value_array: np.array) -> np.array:
    """
    A function return the difference between a given layer performance
    to 0th and the last layer

    Args:
    - layer_cut: int, the layer whose performance will be compared
    - value_array: np.array, the array of all layer performances

    Returns:
    - np.arrary, the performance difference between
        [layer_cut - layer0, final_layer - layer_cut]
    """

    last_layer_numb = len(value_array)

    assert (
        0 < layer_cut < last_layer_numb
    ), f"{layer_cut} not in between 0 and {last_layer_numb}"

    layer_perf = value_array[layer_cut]

    return np.array([layer_perf - value_array[0], value_array[-1] - layer_perf])


def plot_best_layer_bar(
    df: pd.DataFrame, metric: str, path2folder: str = "results/summary/bestemblayer"
):

    """
    A function for plotting a bar plot for
    """

    plot_title = "Best {} achieved at percent depth of pretrain model".format(
        simplify_test_metric(metric)
    )

    print(f"Plotting {plot_title}...")

    fig, ax = plt.subplots()

    df.plot(
        kind="bar",
        x="task",
        y="model_layer_percent",
        color=[TASK_SIMPLE_COLOR_MAP.get(task, "gray") for task in df["task"]],
        ax=ax,
        legend=None,
    )

    ax.set_ylim(0, 1)

    # set labels and title
    plt.xlabel("Task")
    plt.ylabel("Percent")
    plt.title(plot_title, pad=10)

    path2folder = os.path.normpath(path2folder)

    print(f"Saving to {path2folder}...")

    save_plt(fig, plot_title=plot_title, path2folder=path2folder)

    return fig


# TODO clean up combine with plot_emb_df_last_layer_bar
def plot_emb_df_best_layer_bar(
    emb_df: pd.DataFrame,
    metric: str,
    path2folder: str = "results/summary/best/bestemblayer_all",
):

    if "model_layer_percent" not in emb_df.columns:
        emb_df["model_layer_percent"] = emb_df["best_layer"] / emb_df["model_layer"]

    plot_title = "Best embedding layer {} achieved cross pretrain models".format(
        simplify_test_metric(metric)
    )

    print(f"Plotting {plot_title}...")

    # Create a three-degree nested multiclass bar plot
    fig, ax = plt.subplots()
    fig.set_size_inches(24, 6)

    # set up over lay grid
    gs = GridSpec(2, 1, height_ratios=[0.5, 4], hspace=0)

    # Scatter plot above the main bar plot
    ax = plt.subplot(gs[1])
    ax_scatter = plt.subplot(gs[0], sharex=ax)

    # add a second y axis for percent
    ax_percent = ax.twinx()

    # Plotting the bars with different levels of customization
    for i, (t, st, sst, ms, val) in enumerate(
        zip(
            emb_df["task"],
            emb_df["arch"],
            emb_df["model"],
            emb_df["model_layer"],
            emb_df["best_layer"],
        )
    ):

        x = (
            1.2 * i + 1 + math.ceil((i + 1) / 4) * 0.5 + math.ceil((i + 1) / 8) * 0.5
        )  # x-position for each bar

        if st == "esm":
            bar_style = {
                "color": "none",
                "edgecolor": TASK_SIMPLE_COLOR_MAP[t],
                "hatch": "\\",
            }
            color = "none"
        else:
            bar_style = {
                "color": TASK_SIMPLE_COLOR_MAP[t] + "80",
                "edgecolor": TASK_SIMPLE_COLOR_MAP[t],
            }
            color = "gray"

        # Plot the bars with different shading and alpha values
        bar = ax.bar(x, val, linewidth=1.2, **bar_style)

        # add the percent
        ax_percent.scatter(x, val / ms, color="gray", marker="X", s=80, alpha=0.8)

        # Overlay scatter plot indicating model size
        ax_scatter.scatter(
            x, 0.95, s=np.log(ms + 1) * 15, edgecolor="gray", marker="o", color=color
        )

    # Manually create legend elements
    legend_elements = [
        Rectangle(
            (0, 0),
            1,
            1,
            facecolor="none",
            hatch="/",
            edgecolor="gray",
            linewidth=1.2,
            label="ESM",
        ),
        Rectangle(
            (0, 0),
            1,
            1,
            facecolor="gray",
            edgecolor="gray",
            alpha=0.8,
            linewidth=1.2,
            label="CARP",
        ),
        Line2D(
            [0],
            [0],
            marker="X",
            color="w",
            label="Percent of layer",
            markersize=12,
            alpha=0.8,
            markerfacecolor="gray",
            markeredgecolor="gray",
        ),
    ]

    # Create legend
    ax.add_artist(
        ax.legend(
            handles=legend_elements,
            bbox_to_anchor=(1.0425, 1.012),
            loc="upper left",
            title="Embedding type",
        )
    )

    # add legend for arch size
    arch_size_scatter = [None] * (len(EMB_MODEL_LAYER))

    for i, (model, tl) in enumerate(EMB_MODEL_LAYER.items()):

        if "esm" in model:
            mfc = "none"
        else:
            mfc = "gray"

        arch_size_scatter[i] = Line2D(
            [0],
            [0],
            marker="o",
            color="w",
            label=f"{model}: {tl} layer",
            markersize=np.log(tl) * 2.5,
            markerfacecolor="gray",
            markeredgecolor="gray",
            mfc=mfc,
        )

    ax.add_artist(
        ax.legend(
            handles=arch_size_scatter,
            bbox_to_anchor=(1.0425, 0.55),
            loc="upper left",
            title="Pretrained architecture total layer",
        )
    )

    # Remove x and y axis labels for scatter plot
    ax_scatter.xaxis.set_visible(False)
    ax_scatter.set_yticks([])

    # Set y-axis range for scatter plot
    ax_scatter.set_ylim(0, 2)

    # Remove box around the scatter plot
    ax_scatter.spines["top"].set_visible(False)
    ax_scatter.spines["right"].set_visible(False)
    ax_scatter.spines["bottom"].set_visible(False)
    ax_scatter.spines["left"].set_visible(False)

    ax_scatter.set_title(plot_title, pad=0)

    # Set x and y value ranges
    ax.set_xlim(0.36, 112.6)
    ax.set_ylim(0, 56)

    ax_percent.set_ylim(0, 1)

    # Set x-axis ticks and align them to the middle of the corresponding labels
    ax.set_xticks(np.linspace(0.36 + 6, 112.6 - 6, 10))
    ax.set_xticklabels(
        [
            k.replace(" - ", "\n").replace("r l", "r\nl")
            for k in TASK_SIMPLE_COLOR_MAP.keys()
        ],
        ha="center",
    )

    # Set labels and title
    ax.set_xlabel("Tasks")
    ax.set_ylabel("Layer number")
    ax_percent.set_ylabel("Layer percent")

    # Adjust layout for better visibility of legends
    plt.subplots_adjust(right=0.7, bottom=0.2)

    path2folder = checkNgen_folder(os.path.normpath(path2folder))

    print(f"Saving to {path2folder}...")

    save_plt(fig, plot_title=plot_title, path2folder=path2folder)

    return fig


# TODO clean up
def plot_carp_best_layer_bar(
    df: pd.DataFrame,
    metric: str,
    path2folder: str = "results/summary/best/bestemblayer_carp",
):
    """
    A function for plotting best carp layer number

    Args:
    - iflast: bool = False, if take only the last layer but combine rand and stat
    - ifratio: bool = False, if take the ratio against emb
    """

    plot_title = "Best CARP embedding {} achieved cross pretrain degrees".format(
        simplify_test_metric(metric)
    )
    # Create a three-degree nested multiclass bar plot
    fig, ax = plt.subplots()
    fig.set_size_inches(18, 6)

    # set up over lay grid
    gs = GridSpec(2, 1, height_ratios=[0.5, 4], hspace=0)

    # Scatter plot above the main bar plot
    ax = plt.subplot(gs[1])
    ax_scatter = plt.subplot(gs[0], sharex=ax)

    # add a second y axis for percent
    ax_percent = ax.twinx()

    for i, (category, group) in enumerate(df.groupby(["task", "model"])):

        # x-position for each bar
        x = 2 * i + math.ceil((i + 1) / 4) * 3.2

        # Plotting the bars with different levels of customization
        for t, st, sst, ml, ptp, val in zip(
            group["task"],
            group["arch"],
            group["model"],
            group["model_layer"],
            group["ptp"],
            group["best_layer"],
        ):

            # plot the scatter for diff emb layer and percent
            ax.scatter(
                x,  # make x y same size
                val,
                s=80,
                marker="o",
                color=TASK_SIMPLE_COLOR_MAP[t],
                alpha=ptp,
            )

            # add the percent
            ax_percent.scatter(x, val / ml, color="gray", marker="X", s=80, alpha=ptp)

            # Overlay scatter plot indicating model size
            ax_scatter.scatter(
                x,
                0.95,
                s=ml * 1.5,
                edgecolor="gray",
                marker=ARCH_SCATTER_STYLE_DICT[st],
                color="grey",
                alpha=0.8,
            )

    # add legned for emb or percent
    legend_elements = [
        Line2D(
            [0],
            [0],
            marker="o",
            color="w",
            label="CARP",
            markersize=12,
            alpha=0.8,
            markerfacecolor="gray",
            markeredgecolor="gray",
        ),
        Line2D(
            [0],
            [0],
            marker="X",
            color="w",
            label="Percent of layer",
            markersize=12,
            alpha=0.8,
            markerfacecolor="gray",
            markeredgecolor="gray",
        ),
    ]

    # Create legend
    ax.add_artist(
        ax.legend(
            handles=legend_elements,
            bbox_to_anchor=(1.05, 1.012),
            loc="upper left",
            title="Embedding type",
        )
    )

    # add legned for ptp
    alpha_legend = [None] * len(CHECKPOINT_PERCENT)

    for i, a in enumerate(CHECKPOINT_PERCENT):
        alpha_legend[i] = Line2D(
            [0],
            [0],
            marker="o",
            color="w",
            label=a,
            alpha=a,
            markerfacecolor="k",
            markersize=12,
        )

    ax.add_artist(
        ax.legend(
            handles=alpha_legend,
            bbox_to_anchor=(1.05, 0.8),
            loc="upper left",
            title="Pretrain degree",
        )
    )
    # add legend for arch size
    arch_size_scatter = [None] * (len(CARP_MODEL_LAYER))

    for i, (model, tl) in enumerate(CARP_MODEL_LAYER.items()):

        arch_size_scatter[i] = Line2D(
            [0],
            [0],
            marker="o",
            color="w",
            label=f"{model}: {tl} layer",
            markersize=np.log(tl) * 2.5,
            markerfacecolor="gray",
            markeredgecolor="gray",
        )

    ax.add_artist(
        ax.legend(
            handles=arch_size_scatter,
            bbox_to_anchor=(1.05, 0.325),
            loc="upper left",
            title="Pretrained architecture total layer",
        )
    )

    # Remove x and y axis labels for scatter plot
    ax_scatter.xaxis.set_visible(False)
    ax_scatter.set_yticks([])

    # Set y-axis range for scatter plot
    ax_scatter.set_ylim(0, 2)

    # Remove box around the scatter plot
    ax_scatter.spines["top"].set_visible(False)
    ax_scatter.spines["right"].set_visible(False)
    ax_scatter.spines["bottom"].set_visible(False)
    ax_scatter.spines["left"].set_visible(False)

    ax_scatter.set_title(plot_title, pad=0)

    # Set x and y value ranges
    ax.set_xlim(0.36, 112.6)
    ax.set_ylim(0, 56)
    ax_percent.set_ylim(0, 1)

    # Set x-axis ticks and align them to the middle of the corresponding labels
    ax.set_xticks(np.linspace(0.36 + 6, 112.6 - 6, 10))
    ax.set_xticklabels(
        [
            k.replace(" - ", "\n").replace("r l", "r\nl")
            for k in TASK_SIMPLE_COLOR_MAP.keys()
        ],
        ha="center",
    )

    # Set labels and title
    ax.set_xlabel("Tasks")
    ax.set_ylabel("Layer number")
    ax_percent.set_ylabel("Layer percent")

    # Adjust layout for better visibility of legends
    plt.subplots_adjust(right=0.7, bottom=0.2)

    path2folder = checkNgen_folder(os.path.normpath(path2folder))

    print(f"Saving to {path2folder}...")

    save_plt(fig, plot_title=plot_title, path2folder=path2folder)

    return fig


def plot_carp_best_layer_pretrain_loss(
    df: pd.DataFrame,
    metric: str = "test_performance_1",
    ifpercent: bool = True,
    path2folder: str = "results/summar/best/bestemblayer_carp/loss",
):

    """A method for plotting the pretraining arch"""

    if ifpercent:
        append = "percent "
    else:
        append = ""

    plot_title = (
        "Best CARP embedding {}achieved at {} layer cross pretrain degrees".format(
            append, simplify_test_metric(metric)
        )
    )

    # Plot dots with colors corresponding to the category
    fig, ax = plt.subplots()
    fig.set_size_inches(6, 6)

    for i, (category, group) in enumerate(df.groupby(["task", "model"])):

        t, m = category

        c = TASK_SIMPLE_COLOR_MAP[t]

        xs = [CARP_CHECKPOINT_LOSSES[m][x] for x in group["ptp"]]

        if ifpercent:
            ys = group["best_layer"] / group["model_layer"]
        else:
            ys = group["best_layer"]

        ax.plot(
            xs,
            ys,
            marker="o",
            markersize=12,
            linestyle="dotted",
            color=c,
            markerfacecolor=c,
            markeredgecolor=c,
            alpha=CARP_ALPHA[m],
        )

    # add task legend
    task_legend_list = [None] * len(TASK_SIMPLE_COLOR_MAP)

    for i, (t, c) in enumerate(TASK_SIMPLE_COLOR_MAP.items()):

        task_legend_list[i] = Line2D(
            [0],
            [0],
            marker="o",
            color=c,
            label=t,
            markersize=12,
            linestyle="dotted",
            markerfacecolor=c,
            markeredgecolor=c,
        )

    ax.add_artist(
        ax.legend(
            handles=task_legend_list,
            bbox_to_anchor=(1, 1.012),
            loc="upper left",
            title="Task",
        )
    )

    # add alpha legend for carp sizes
    arch_legend_list = [None] * len(CARP_ALPHA)

    for i, (c, a) in enumerate(CARP_ALPHA.items()):

        arch_legend_list[i] = Line2D(
            [0],
            [0],
            marker="o",
            color="gray",
            alpha=a,
            linestyle="dotted",
            label=c,
            markersize=12,
            markerfacecolor="gray",
            markeredgecolor="gray",
        )

    ax.add_artist(
        ax.legend(
            handles=arch_legend_list,
            bbox_to_anchor=(1, 0.5025),
            loc="upper left",
            title="CARP models",
        )
    )

    # Flip the x-axis so loss from left to right are reducing
    ax.invert_xaxis()

    if ifpercent:
        ax.set_ylim(0, 1)
        ax.set_ylabel("Layer percent")
    else:
        ax.set_ylim(0, 56)
        ax.set_ylabel("Layer number")

    # Set labels and title
    ax.set_xlabel("Pretrain loss")

    ax.set_title(plot_title, pad=10)

    path2folder = checkNgen_folder(os.path.join(os.path.normpath(path2folder), append))

    print(f"Saving to {path2folder}...")

    save_plt(fig, plot_title=plot_title, path2folder=path2folder)

    return fig


def plot_emb_df_last_layer_bar(
    emb_df: pd.DataFrame,
    onehot_df: pd.DataFrame,
    metric: str,
    path2folder: str = "results/summary/last/embvsonehot_bar",
):

    plot_title = "Last layer embedding {} against onehot".format(
        simplify_test_metric(metric)
    )

    # Create a three-degree nested multiclass bar plot
    fig, ax = plt.subplots()
    fig.set_size_inches(24, 6)

    # set up over lay grid
    gs = GridSpec(2, 1, height_ratios=[0.5, 4], hspace=0)

    # Scatter plot above the main bar plot
    ax = plt.subplot(gs[1])
    ax_scatter = plt.subplot(gs[0], sharex=ax)

    # plot horizontal lines for onehot baselines
    for i, t in enumerate(TASK_SIMPLE_COLOR_MAP.keys()):

        baseline_value = onehot_df[onehot_df["task"] == t][
            "last_value"
        ].max()  # Adjust as needed
        ax.axhline(
            y=baseline_value,
            linestyle="--",
            color=TASK_SIMPLE_COLOR_MAP[t],
            xmin=0.008 + i * 0.09875,
            xmax=i * 0.09875 + 0.103,
            linewidth=1.2,
        )

    # Plotting the bars with different levels of customization
    for i, (t, st, sst, ms, val) in enumerate(
        zip(
            emb_df["task"],
            emb_df["arch"],
            emb_df["model"],
            emb_df["model_size"],
            emb_df["last_value"],
        )
    ):

        x = (
            1.2 * i + 1 + math.ceil((i + 1) / 4) * 0.5 + math.ceil((i + 1) / 8) * 0.5
        )  # x-position for each bar

        if st == "esm":
            bar_style = {
                "color": "none",
                "edgecolor": TASK_SIMPLE_COLOR_MAP[t],
                "hatch": "\\",
            }
            color = "none"
        else:
            bar_style = {
                "color": TASK_SIMPLE_COLOR_MAP[t] + "80",
                "edgecolor": TASK_SIMPLE_COLOR_MAP[t],
            }
            color = "gray"

        # Plot the bars with different shading and alpha values
        bar = ax.bar(x, val, linewidth=1.2, **bar_style)

        # Overlay scatter plot indicating another parameter
        scatter = ax_scatter.scatter(
            x, 0.95, s=np.log(ms + 1) * 15, edgecolor="gray", marker="o", color=color
        )

    # Manually create legend elements
    legend_elements = [
        Rectangle(
            (0, 0),
            1,
            1,
            facecolor="none",
            hatch="/",
            edgecolor="gray",
            linewidth=1.2,
            label="ESM",
        ),
        Rectangle(
            (0, 0),
            1,
            1,
            facecolor="gray",
            edgecolor="gray",
            alpha=0.8,
            linewidth=1.2,
            label="CARP",
        ),
        Line2D([0], [0], linestyle="--", color="gray", label="Onehot"),
    ]

    # Create legend
    ax.add_artist(
        ax.legend(
            handles=legend_elements,
            bbox_to_anchor=(1, 1.012),
            loc="upper left",
            title="Embedding type",
        )
    )

    # add legend for arch size
    arch_size_scatter = [None] * (len(EMB_MODEL_SIZE))

    for i, (model, size) in enumerate(EMB_MODEL_SIZE.items()):

        if "esm" in model:
            mfc = "none"
        else:
            mfc = "gray"

        arch_size_scatter[i] = Line2D(
            [0],
            [0],
            marker="o",
            color="w",
            label=f"{model}: {size} M",
            markersize=np.log(size + 1) + 2.75,
            markerfacecolor="gray",
            markeredgecolor="gray",
            mfc=mfc,
        )

    ax.add_artist(
        ax.legend(
            handles=arch_size_scatter,
            bbox_to_anchor=(1, 0.55),
            loc="upper left",
            title="Pretrained architecture sizes",
        )
    )

    # Remove x and y axis labels for scatter plot
    ax_scatter.xaxis.set_visible(False)
    ax_scatter.set_yticks([])

    # Set y-axis range for scatter plot
    ax_scatter.set_ylim(0, 2)

    # Remove box around the scatter plot
    ax_scatter.spines["top"].set_visible(False)
    ax_scatter.spines["right"].set_visible(False)
    ax_scatter.spines["bottom"].set_visible(False)
    ax_scatter.spines["left"].set_visible(False)

    ax_scatter.set_title(plot_title, pad=0)

    # Set x and y value ranges
    ax.set_xlim(0.36, 112.6)

    if metric == "test_loss":
        ax.set_yscale("log")
    else:
        ax.set_ylim(0, 1)

    # Set x-axis ticks and align them to the middle of the corresponding labels
    ax.set_xticks(np.linspace(0.36 + 6, 112.6 - 6, 10))
    ax.set_xticklabels(
        [
            k.replace(" - ", "\n").replace("r l", "r\nl")
            for k in TASK_SIMPLE_COLOR_MAP.keys()
        ],
        ha="center",
    )

    # Set labels and title
    ax.set_xlabel("Tasks")
    ax.set_ylabel("Last layer embedding test performance")

    # Adjust layout for better visibility of legends
    plt.subplots_adjust(right=0.7, bottom=0.2)

    path2folder = checkNgen_folder(os.path.normpath(path2folder))

    print(f"Saving to {path2folder}...")

    save_plt(fig, plot_title=plot_title, path2folder=path2folder)

    return fig


def plot_emb_layer_bar(
    df: pd.DataFrame,
    val_col: list[str],
    metric: str,
    iflast: bool = False,
    ifratio: bool = False,
    ifonehot: bool = True,
    ifemb: bool = False,
    path2folder: str = "results/summary/last/emblayersvsonehot_bar",
):
    """
    A function for plotting different layer of emb

    Args:
    - iflast: bool = False, if take only the last layer but combine rand and stat
    - ifratio: bool = False, if take the ratio against emb
    - ifemb: bool = False, if compared against emb
    """

    if ifemb:
        baseline = "Embedding"
        embonehot = "emb"
    else:
        baseline = "Onehot"
        embonehot = "onehot"

    if iflast:

        difflast = "Last"

        alpha_style = ARCH_AB_DOT_STYLE_DICT.copy()

        title_ab = "embedding ablation"

        if ifratio:
            ylabel = "(Embedding ablation - Onehot) / (Embedding - Onehot)"
            path2folder = path2folder.replace(
                "emblayersvsonehot_bar", "randstatembvsonehot_bar"
            )
        else:
            ylabel = f"Embedding ablation - {baseline}"
            path2folder = path2folder.replace(
                "emblayersvsonehot_bar", f"randstatvs{embonehot}_bar"
            )

    else:

        difflast = "Different"

        alpha_style = {"alpha": LAYER_ALPHAS, "edgecolor": "none"}

        if "emb" in df["ablation"].values:
            title_ab = "embedding"
            ylabel = f"Embedding - Onehot"

        elif "rand" in df["ablation"].values:
            title_ab = "random init"
            if ifratio:
                path2folder = path2folder.replace(
                    "emblayersvsonehot_bar", "randembvsonehot_bar"
                )
                ylabel = "(Random init - Onehot) / (Embedding - Onehot)"
            else:
                path2folder = path2folder.replace(
                    "emblayersvsonehot_bar", f"randvs{embonehot}_bar"
                )
                ylabel = f"Random init - {baseline}"

        elif "stat" in df["ablation"].values:
            title_ab = "stat transfer"
            if ifratio:
                path2folder = path2folder.replace(
                    "emblayersvsonehot_bar", "statembvsonehot_bar"
                )
                ylabel = "(Stat transfer - Onehot) / (Embedding - Onehot)"
            else:
                path2folder = path2folder.replace(
                    "emblayersvsonehot_bar", f"statvs{embonehot}_bar"
                )
                ylabel = f"Stat transfer - {baseline}"

    plot_title = "{} layer {} {} compared against {}".format(
        difflast, title_ab, simplify_test_metric(metric), baseline.lower()
    )

    # Create a three-degree nested multiclass bar plot
    fig, ax = plt.subplots()
    fig.set_size_inches(24, 6)

    # set up over lay grid
    gs = GridSpec(2, 1, height_ratios=[0.5, 4], hspace=0)

    # Scatter plot above the main bar plot
    ax = plt.subplot(gs[1])
    ax_scatter = plt.subplot(gs[0], sharex=ax)

    # Plotting the bars with different levels of customization
    for i, (t, st, sst, ml, vals) in enumerate(
        zip(
            df["task"],
            df["arch"],
            df["model"],
            df["model_layer"],
            df[val_col].to_numpy(),
        )
    ):

        # x-position for each bar
        x = 1.2 * i + 1 + math.ceil((i + 1) / 4) * 0.5 + math.ceil((i + 1) / 8) * 0.5

        if iflast:

            for v, col in enumerate(val_col):

                val = col.split("_")[0]

                # plot the scatter for diff emb layer
                ax.scatter(
                    x,
                    vals[v],
                    s=80,
                    marker=ARCH_SCATTER_STYLE_DICT[st],
                    color=TASK_SIMPLE_COLOR_MAP[t],
                    **alpha_style[val],
                )

        else:
            # plot the scatter for diff emb layer
            ax.scatter(
                [x] * len(val_col),  # make x y same size
                vals,
                s=80,
                marker=ARCH_SCATTER_STYLE_DICT[st],
                color=TASK_SIMPLE_COLOR_MAP[t],
                **alpha_style,
            )

        # Overlay scatter plot indicating model size
        ax_scatter.scatter(
            x,
            0.95,
            s=ml * 1.5,
            edgecolor="gray",
            marker=ARCH_SCATTER_STYLE_DICT[st],
            color="grey",
            alpha=0.8,
        )

    # Manually create legend elements
    arch_legend_list = [None] * len(ARCH_TYPE)

    for i, a in enumerate(ARCH_TYPE):

        arch_legend_list[i] = Line2D(
            [0],
            [0],
            marker=ARCH_SCATTER_STYLE_DICT[a],
            color="w",
            linestyle=ARCH_LINE_STYLE_DICT[a]["linestyle"],
            label=a.upper(),
            markersize=10,
            markerfacecolor="gray",
            markeredgecolor="none",
        )

    # add onehot y=0 line

    if ifratio:
        ref_y = 1
        ref_label = "Emb = Ablation"
    else:
        ref_y = 0
        ref_label = baseline

    ax.axhline(
        y=ref_y,
        linestyle="--",
        color="gray",
        linewidth=1.2,
    )
    arch_legend_list.append(
        Line2D([0], [0], linestyle="--", color="gray", label=ref_label)
    )

    ax.add_artist(
        ax.legend(
            handles=arch_legend_list,
            bbox_to_anchor=(1, 1.012),
            loc="upper left",
            title="Pretrained architectures",
        )
    )

    # add legend for arch size
    arch_size_scatter = [None] * (len(EMB_MODEL_LAYER))

    for i, (model, tl) in enumerate(EMB_MODEL_LAYER.items()):

        if "esm" in model:
            arch_type = "esm"
        else:
            arch_type = "carp"

        arch_size_scatter[i] = Line2D(
            [0],
            [0],
            marker=ARCH_SCATTER_STYLE_DICT[arch_type],
            color="w",
            label=f"{model}: {tl} layer",
            markersize=np.log(tl) * 2.5,
            markerfacecolor="gray",
            markeredgecolor="none",
            # mfc=mfc,
        )

    ax.add_artist(
        ax.legend(
            handles=arch_size_scatter,
            bbox_to_anchor=(1, 0.55),
            loc="upper left",
            title="Pretrained architecture total layer",
        )
    )

    if iflast:
        alpha_title = "Embedding ablation"

        alpha_legend = [None] * len(val_col)

        for i, c in enumerate(sorted(val_col.copy())):

            ab = c.replace("_value", "")

            if ab == "rand":
                mfc = "none"
            else:
                mfc = "gray"

            alpha_legend[i] = Line2D(
                [0],
                [0],
                marker="o",
                color="w",
                label=ARCH_AB_DICT[ab].capitalize(),
                markersize=10,
                markerfacecolor="gray",
                markeredgecolor="gray",
                mfc=mfc,
            )

    else:
        alpha_title = "Layer"
        # add alpha legend
        alpha_legend = [None] * len(LAYER_ALPHAS)

        for i, (c, a) in enumerate(zip(sorted(val_col.copy()), sorted(LAYER_ALPHAS))):
            alpha_legend[i] = Line2D(
                [0],
                [0],
                marker="o",
                color="w",
                label=c.replace("_value", ""),
                alpha=a,
                markerfacecolor="k",
                markersize=10,
            )

    ax.add_artist(
        ax.legend(
            handles=alpha_legend,
            bbox_to_anchor=(1.135, 1.012),
            loc="upper left",
            title=alpha_title,
        )
    )

    # Remove x and y axis labels for scatter plot
    ax_scatter.xaxis.set_visible(False)
    ax_scatter.set_yticks([])

    # Set y-axis range for scatter plot
    ax_scatter.set_ylim(0, 2)

    # Remove box around the scatter plot
    ax_scatter.spines["top"].set_visible(False)
    ax_scatter.spines["right"].set_visible(False)
    ax_scatter.spines["bottom"].set_visible(False)
    ax_scatter.spines["left"].set_visible(False)

    ax_scatter.set_title(plot_title, pad=0)

    # Set x and y value ranges
    ax.set_xlim(0.36, 112.6)

    if metric == "test_loss" or ifratio:
        ax.set_yscale("symlog")
    ax.autoscale(axis="y")

    # Set x-axis ticks and align them to the middle of the corresponding labels
    ax.set_xticks(np.linspace(0.36 + 6, 112.6 - 6, 10))
    ax.set_xticklabels(
        [
            k.replace(" - ", "\n").replace("r l", "r\nl")
            for k in TASK_SIMPLE_COLOR_MAP.keys()
        ],
        ha="center",
    )

    # Set labels and title
    ax.set_xlabel("Tasks")
    ax.set_ylabel(ylabel)

    # Adjust layout for better visibility of legends
    plt.subplots_adjust(right=0.7, bottom=0.2)

    path2folder = checkNgen_folder(os.path.normpath(path2folder))

    print(f"Saving to {path2folder}...")

    save_plt(fig, plot_title=plot_title, path2folder=path2folder)

    return fig


def plot_emb_onehot(
    df: pd.DataFrame,
    metric: str,
    path2folder: str = "results/summary/best/embonehot",
):
    """A function for plotting best emb vs onehot"""

    if "best" in path2folder:
        bestorlast = "Best"
    else:
        bestorlast = "Last layer"

    plot_title = "{} {} embedding against onehot baseline".format(
        bestorlast, simplify_test_metric(metric)
    )

    print(f"Plotting {plot_title}...")

    fig, ax = plt.subplots()
    fig.set_size_inches(6, 6)

    # get the min x or y for the diagnol line
    diag_min = 1

    for (task, c) in TASK_SIMPLE_COLOR_MAP.items():

        sliced_df = df[df["task"] == task]

        x = sliced_df[sliced_df["ablation"] == "onehot"]["best_value"].values
        y = sliced_df[sliced_df["ablation"] == "emb"]["best_value"].values

        if metric != "test_loss":
            min_xy = min(min(y), min(x))
            if min_xy < diag_min:
                diag_min = min_xy

        scatter = ax.scatter(x, y, c=c, s=200, alpha=0.8, label=task, edgecolors="none")

    if metric != "test_loss":
        # diag min to smallest one decimal
        diag_min = math.floor(diag_min * 10) / 10

        # Add a diagonal line
        plt.plot(
            [diag_min, 1],
            [diag_min, 1],
            linestyle=":",
            color="grey",
        )

    ax.add_artist(ax.legend(title="Tasks", bbox_to_anchor=(1, 1.012), loc="upper left"))

    if metric == "test_loss":
        plt.xscale("log")
        plt.yscale("log")

    plt.ylabel(f"{bestorlast} embedding test performance")
    plt.xlabel("Onehot")
    plt.title(plot_title)

    path2folder = checkNgen_folder(os.path.normpath(path2folder))

    print(f"Saving to {path2folder}...")

    save_plt(fig, plot_title=plot_title, path2folder=path2folder)

    return fig


# make df no ptp dets already
def plot_emb_onehot_det(
    df: pd.DataFrame,
    metric: str,
    path2folder: str = "results/summary/last/embonehot",
):

    """
    A function for plotting emb with different size and arch vs onehot
    """

    if "best" in path2folder:
        bestorlast = "Best"
    else:
        bestorlast = "Last layer"

    plot_title = "{} {} embedding against onehot baseline".format(
        bestorlast, simplify_test_metric(metric)
    )

    print(f"Plotting {plot_title}...")

    fig, ax = plt.subplots()
    fig.set_size_inches(6, 6)

    if "best_value" in df.columns:
        val_col = "best_value"
    elif "last_value" in df.columns:
        val_col = "last_value"

    # get the min x or y for the diagnol line
    diag_min = 1

    for arch in ARCH_TYPE:

        for (task, c) in TASK_SIMPLE_COLOR_MAP.items():

            sliced_df = df[(df["task"] == task) & (df["arch"] == arch)]

            onehot_df = sliced_df[sliced_df["ablation"] == "onehot"]
            emb_df = sliced_df[sliced_df["ablation"] == "emb"]

            x = onehot_df[val_col].values
            y = emb_df[val_col].values
            # x will be onehot for esm or carp for the given task
            # and y will be all different arch
            x = np.repeat(x, len(y))

            if metric != "test_loss":
                min_xy = min(min(y), min(x))
                if min_xy < diag_min:
                    diag_min = min_xy

            s = np.log(emb_df["model_size"].values + 1) * 18

            arch_dict = ARCH_DOT_STYLE_DICT[arch]

            if arch == "carp":
                arch_dict["c"] = c
                arch_dict["label"] = task
            elif arch == "esm":
                arch_dict["edgecolor"] = c

            scatter = ax.scatter(x, y, s=s, **arch_dict)

    if metric != "test_loss":
        # diag min to smallest one decimal
        diag_min = math.floor(diag_min * 10) / 10

        # Add a diagonal line
        plt.plot(
            [diag_min, 1],
            [diag_min, 1],
            linestyle=":",
            color="grey",
        )

    ax.add_artist(ax.legend(title="Tasks", bbox_to_anchor=(1, 1.012), loc="upper left"))

    # add legend for arch size
    arch_size_scatter = [None] * len(EMB_MODEL_SIZE)

    for i, (model, size) in enumerate(EMB_MODEL_SIZE.items()):

        if "carp" in model:
            mfc = "gray"
        else:
            mfc = "none"

        arch_size_scatter[i] = Line2D(
            [0],
            [0],
            marker="o",
            color="w",
            label=f"{model}: {size} M",
            markersize=np.log(size + 1) + 2.75,
            markerfacecolor="gray",
            markeredgecolor="gray",
            mfc=mfc,
        )

    ax.add_artist(
        ax.legend(
            handles=arch_size_scatter,
            bbox_to_anchor=(1, 0.49),
            loc="upper left",
            title="Pretrained architecture sizes",
        )
    )

    if metric == "test_loss":
        plt.xscale("log")
        plt.yscale("log")

    plt.ylabel(f"{bestorlast} embedding test performance")
    plt.xlabel("Onehot")
    plt.title(plot_title)

    path2folder = checkNgen_folder(os.path.normpath(path2folder))

    print(f"Saving to {path2folder}...")

    save_plt(fig, plot_title=plot_title, path2folder=path2folder)

    return fig


def plot_randstat(
    df: pd.DataFrame,
    metric: str,
    randstat: str,
    delta_onehot: bool = True,
    path2folder: str = "results/summary/last/randstat",
):

    """
    A function for plotting emb vs rand or stat
    or rand vs stat
    """

    if "best" in path2folder:
        bestorlast = "Best"
    else:
        bestorlast = "Last layer"

    if randstat == "":
        comp_det = " vs ".join(list(INIT_DICT.values()))
        plot_title = "{} {} embedding same layer {}".format(
            bestorlast, simplify_test_metric(metric), comp_det
        )
        pathrandstat = "vs"
    else:
        plot_title = "{} {} embedding against same layer {}".format(
            bestorlast, simplify_test_metric(metric), INIT_DICT[randstat]
        )
        pathrandstat = randstat

    print(f"Plotting {plot_title}...")

    fig, ax = plt.subplots()
    fig.set_size_inches(6, 6)

    # get the min x or y for the diagnol line
    diag_min = 1

    if delta_onehot:
        diag_max = 0
        label_append = "- onehot"
        path_append = "onehot"

    else:
        diag_max = 1
        label_append = ""
        path_append = ""

    for (task, c) in TASK_SIMPLE_COLOR_MAP.items():

        task_df = df[df["task"] == task]

        if delta_onehot:
            if randstat == "":
                x = task_df["rand - onehot"].values
                y = task_df["stat - onehot"].values
            else:
                x = task_df["emb - onehot"].values
                y = task_df[f"{randstat} - onehot"].values

            if metric != "test_loss":

                max_xy = max(max(x), max(y))

                if max_xy > diag_max:
                    diag_max = max_xy

        else:
            if randstat == "":
                x = task_df["rand"].values
                y = task_df["stat"].values
            else:
                x = task_df["best_value"].values
                y = task_df[randstat].values

        if metric != "test_loss":

            min_xy = min(min(y), min(x))

            if min_xy < diag_min:
                diag_min = min_xy

            min_xy = min(min(y), min(x))
            if min_xy < diag_min:
                diag_min = min_xy

        scatter = ax.scatter(x, y, c=c, s=200, alpha=0.8, label=task, edgecolors="none")

    if metric != "test_loss":

        # diag min to smallest one decimal and max to largest
        diag_min = math.floor(diag_min * 10) / 10
        diag_max = math.ceil(diag_max * 10) / 10

        # Add a diagonal line
        plt.plot(
            [diag_min, diag_max],
            [diag_min, diag_max],
            linestyle=":",
            color="grey",
        )

    ax.add_artist(ax.legend(title="Tasks", bbox_to_anchor=(1, 1.012), loc="upper left"))

    if randstat == "":
        plt.xlabel("{} embedding random init {}".format(bestorlast, label_append))
        plt.ylabel("{} embedding stat transfer {}".format(bestorlast, label_append))
    else:
        plt.xlabel("{} embedding {}".format(bestorlast, label_append))
        plt.ylabel(f"{INIT_DICT[randstat].capitalize()} {label_append}")

    plt.title(plot_title)

    path2folder = checkNgen_folder(
        os.path.normpath(os.path.join(path2folder, pathrandstat, path_append))
    )

    print(f"Saving to {path2folder}...")

    save_plt(fig, plot_title=plot_title, path2folder=path2folder)

    return fig


def plot_pretrain_degree(
    emb_df: pd.DataFrame,
    metric: str = "test_performance_1",
    arch: str = "carp",
    delta_onehot: bool = True,
    ifloss: bool = True,
    path2folder: str = "results/summar/last/pretraindegree",
):

    """A method for plotting the pretraining arch"""

    if "best" in path2folder:
        bestorlast = "Best"
    else:
        bestorlast = "Last layer"

    if ifloss:
        loss_append = "_loss"
    else:
        loss_append = ""

    plot_title = "{} {} cross different pretrain degrees of {}".format(
        bestorlast, simplify_test_metric(metric), arch.upper()
    )

    # set up the id var keys for melt
    melt_id_cols = ["task", "model"]

    if delta_onehot:
        melt_cols = melt_id_cols + [str(p) + " - onehot" for p in CHECKPOINT_PERCENT]
        label_append = " - onehot"
        path_append = "onehot"
        y_max = None
    else:
        melt_cols = melt_id_cols + [str(p) for p in CHECKPOINT_PERCENT]
        label_append = ""
        path_append = ""
        y_max = 1

    x_name = "Pretrain degree"
    y_name = "{} test performance{}".format(bestorlast, label_append)

    emb_df_melt = pd.melt(
        emb_df[melt_cols],
        id_vars=melt_id_cols,
        var_name=x_name,
        value_name=y_name,
    )

    # Plot dots with colors corresponding to the category
    fig, ax = plt.subplots()
    fig.set_size_inches(6, 6)

    for category, group in emb_df_melt.groupby("task"):
        for subc, subg in group.groupby("model"):

            if ifloss:
                xs = [
                    CARP_CHECKPOINT_LOSSES[subc][float(x.replace(label_append, ""))]
                    for x in subg[x_name]
                ]
            else:
                xs = [float(x.replace(label_append, "")) for x in subg[x_name]]

            c = TASK_SIMPLE_COLOR_MAP[category]

            ax.plot(
                xs,
                subg[y_name],
                marker="o",
                markersize=12,
                linestyle="solid",
                color=c,
                markerfacecolor=c,
                markeredgecolor=c,
                alpha=CARP_ALPHA[subc],
            )

    # add onehot baseline if delta
    if delta_onehot:
        ax.axhline(
            y=0,
            linestyle="--",
            color="gray",
            linewidth=1.2,
        )

    # add task legend
    task_legend_list = [None] * len(TASK_SIMPLE_COLOR_MAP)

    for i, (t, c) in enumerate(TASK_SIMPLE_COLOR_MAP.items()):

        task_legend_list[i] = Line2D(
            [0],
            [0],
            marker="o",
            color=c,
            label=t,
            markersize=12,
            linestyle="solid",
            markerfacecolor=c,
            markeredgecolor=c,
        )

    ax.add_artist(
        ax.legend(
            handles=task_legend_list,
            bbox_to_anchor=(1, 1.012),
            loc="upper left",
            title="Task",
        )
    )

    # add alpha legend for carp sizes
    arch_legend_list = [None] * len(CARP_ALPHA)

    for i, (c, a) in enumerate(CARP_ALPHA.items()):

        arch_legend_list[i] = Line2D(
            [0],
            [0],
            marker="o",
            color="gray",
            alpha=a,
            linestyle="solid",
            label=c,
            markersize=12,
            markerfacecolor="gray",
            markeredgecolor="gray",
        )

    if delta_onehot:
        arch_legend_list.append(
            Line2D([0], [0], linestyle="--", color="gray", label="Onehot")
        )

    ax.add_artist(
        ax.legend(
            handles=arch_legend_list,
            bbox_to_anchor=(1, 0.5025),
            loc="upper left",
            title="CARP models",
        )
    )

    if ifloss:
        x_label = "Pretrain loss"
        # Flip the x-axis so loss from left to right are reducing
        ax.invert_xaxis()

    else:
        x_label = x_name
        ax.set_xticks(CHECKPOINT_PERCENT)
        ax.set_xticklabels([str(tick) for tick in CHECKPOINT_PERCENT])

        ax.set_xlim(0, 1.125)

    if metric != "test_loss":
        ax.set_ylim(None, y_max)
    else:
        if not delta_onehot:
            plt.yscale("symlog")
            ax.autoscale(axis="y")

    # Set labels and title
    ax.set_xlabel(x_label)
    ax.set_ylabel(y_name)
    ax.set_title(plot_title, pad=10)

    path2folder = checkNgen_folder(
        os.path.join(
            os.path.normpath(
                path2folder.replace("pretraindegree", "pretraindegree" + loss_append)
            ),
            path_append,
        )
    )

    print(f"Saving to {path2folder}...")

    save_plt(fig, plot_title=plot_title, path2folder=path2folder)

    return fig


def plot_arch_size(
    arch_df_dict: dict,
    metric: str = "test_performance_1",
    arch: str = "",
    delta_onehot: bool = True,
    path2folder: str = "results/summary/archsize",
):

    """
    A function for plotting performance vs arch size

    Args:
    - arch: str = "", for both arch combined
    """

    if arch == "":
        arch_name = "architectures"
        arch_list = ARCH_TYPE
    else:
        arch_name = arch.upper()
        arch_list = [arch]

    plot_title = "Last layer {} cross different sizes of pretrain {}".format(
        simplify_test_metric(metric), arch_name
    )

    if delta_onehot:
        label_append = " - onehot"
        path_append = "onehot"
        y_max = None
    else:
        label_append = ""
        path_append = ""
        y_max = 1

    # Plot dots with colors corresponding to the category
    fig, ax = plt.subplots()
    fig.set_size_inches(6, 6)

    for i, a in enumerate(arch_list):

        arch_df = arch_df_dict[a][metric]

        if a == "carp":
            # ignore ptp not 1 or 0
            # Select rows where 'Column2' has a value from the list
            arch_df = arch_df[
                arch_df["ptp"].isin(arch_df_dict["esm"][metric]["ptp"].unique())
            ]

        # get rid of esm1b
        elif a == "esm":
            arch_df = arch_df[~arch_df["model"].isin(["esm1b_t33_650M_UR50S"])]

        else:
            print(f"{a} not in {ARCH_TYPE}")

        for category, group in arch_df.sort_values(["model_size"]).groupby("task"):

            # to not duplicate label
            if a == "carp" or len(arch_list) == 1:
                label = category
            else:
                label = None

            if delta_onehot:

                group = group.reset_index(drop=True).copy()

                # Identify the onehot row
                onehot_row = group[group["ablation"] == "onehot"].index
                onehot_val = group.loc[
                    group["ablation"] == "onehot", "last_value"
                ].iloc[0]

                # subtract the onehot value from other rows
                group["last_value"] -= onehot_val

                # drop the onehot row
                group = group.drop(index=onehot_row)

            ax.plot(
                group["model_size"],
                group["last_value"],
                marker="o",
                markersize=12,
                alpha=0.8,
                label=label,
                color=TASK_SIMPLE_COLOR_MAP.get(category, "gray"),
                **ARCH_LINE_STYLE_DICT[a],
            )

    # add onehot baseline if delta
    if delta_onehot:
        ax.axhline(
            y=0,
            linestyle="--",
            color="gray",
            linewidth=1.2,
        )

    # add additional legend if for both
    if arch == "":
        arch_legend_list = [None] * len(ARCH_TYPE)

        for i, a in enumerate(ARCH_TYPE):

            if a == "esm":
                mfc = "none"
            else:
                mfc = "gray"

            arch_legend_list[i] = Line2D(
                [0],
                [0],
                marker="o",
                color="gray",
                linestyle=ARCH_LINE_STYLE_DICT[a]["linestyle"],
                label=a.upper(),
                markersize=12,
                mfc=mfc,
            )

        if delta_onehot:
            arch_legend_list.append(
                Line2D([0], [0], linestyle="--", color="gray", label="Onehot")
            )

        ax.add_artist(
            ax.legend(
                handles=arch_legend_list,
                bbox_to_anchor=(1, 0.5025),
                loc="upper left",
                title="Pretrained architectures",
            )
        )
    else:
        ax.legend(
            bbox_to_anchor=(1, 0.5025),
            loc="upper left",
            title="Pretrained architectures",
        )

    plt.xscale("log")

    if metric == "test_loss":
        plt.yscale("symlog")
        ax.autoscale(axis="y")

    if metric != "test_loss":
        # Set y-axis limits
        ax.set_ylim(bottom=None, top=y_max)

    # Set labels and title
    ax.set_xlabel("Log model size (M)")
    ax.set_ylabel(f"Last layer test performance{label_append}")
    ax.set_title(plot_title)

    ax.add_artist(ax.legend(title="Tasks", bbox_to_anchor=(1, 1.012), loc="upper left"))

    path2folder = checkNgen_folder(
        os.path.join(os.path.normpath(path2folder), path_append)
    )

    print(f"Saving to {path2folder}...")

    save_plt(fig, plot_title=plot_title, path2folder=path2folder)

    return fig


def plot_layer_delta_simple(
    df: pd.DataFrame,
    layer_cut: int,
    metric: str,
    path2folder: str = "results/summary/last/layerdelta_simple",
):
    """
    A function for plotting and saving layer delta
    after selecting the best performance based on given metric
    """

    if "best" in path2folder:
        bestorlast = "Best"
    else:
        bestorlast = "Last layer"

    plot_title = "{} {} at x = {}".format(
        bestorlast, simplify_test_metric(metric), layer_cut
    )

    print(f"Plotting {plot_title}...")

    fig, ax = plt.subplots()
    fig.set_size_inches(6, 6)

    for (task, c) in TASK_SIMPLE_COLOR_MAP.items():

        sliced_df = df[df["task"] == task]
        x = sliced_df["x-0"].values
        y = sliced_df["f-x"].values

        scatter = ax.scatter(x, y, c=c, label=task, s=200, alpha=0.8, edgecolors="none")

    # add colored task legend
    ax.add_artist(ax.legend(title="Tasks", bbox_to_anchor=(1, 1.012), loc="upper left"))

    plt.xlabel("x-0")
    plt.ylabel("f-x")
    plt.title(plot_title)

    path2folder = os.path.normpath(path2folder)

    print(f"Saving to {path2folder}...")

    save_plt(fig, plot_title=plot_title, path2folder=path2folder)

    return fig


def plot_layer_delta_det(
    df: pd.DataFrame,
    layer_cut: int,
    arch: str,
    metric: str,
    path2folder: str = "results/summary/layerdelta_det",
):
    """A function for plotting and saving layer delta"""

    plot_title = "{} layer {} at x = {}".format(
        arch.upper(), simplify_test_metric(metric), layer_cut
    )

    print(f"Plotting {plot_title}...")

    if arch == "esm":
        alaph_values = [0.8]
        alpha_unique = alaph_values
        alpha_label = ["1"]
        size_legend_label = list(MODEL_SIZE.keys())[:4]
    else:
        alaph_values = df["ptp"].values
        alpha_unique = list(df["ptp"].unique())
        alpha_label = [str(a) for a in alpha_unique]
        size_legend_label = list(MODEL_SIZE.keys())[-5:-1]

    fig, ax = plt.subplots()
    fig.set_size_inches(10, 7.5)

    for (task, c) in TASK_SIMPLE_COLOR_MAP.items():
        sliced_df = df[df["task"] == task]
        x = sliced_df["x-0"].values
        y = sliced_df["f-x"].values
        s = np.log(sliced_df["model_size"].values + 1) * 18
        scatter = ax.scatter(
            x, y, c=c, s=s, label=task, alpha=alaph_values, edgecolors="none"
        )

    # add colored task legend
    ax.add_artist(ax.legend(title="Tasks", bbox_to_anchor=(1, 1.012), loc="upper left"))

    # add size legend
    handles, labels = scatter.legend_elements(prop="sizes", color="k", alpha=0.8)
    legend2 = ax.legend(
        handles,
        size_legend_label,
        bbox_to_anchor=(1, 0.59),
        loc="upper left",
        title="Model sizes",
    )
    ax.add_artist(legend2)

    # add alpha legend
    alpha_legend = [None] * len(alpha_unique)

    for i, (a_value, a_lable) in enumerate(zip(alpha_unique, alpha_label)):
        alpha_legend[i] = Line2D(
            [0],
            [0],
            marker="o",
            color="w",
            label=a_lable,
            alpha=a_value,
            markerfacecolor="k",
            markersize=10,
        )

    ax.add_artist(
        ax.legend(
            handles=alpha_legend,
            bbox_to_anchor=(1, 0.39),
            loc="upper left",
            title="Pretraining degree",
        )
    )

    plt.xlabel("x-0")
    plt.ylabel("f-x")
    plt.title(plot_title)

    path2folder = os.path.normpath(path2folder)

    print(f"Saving to {path2folder}...")

    save_plt(fig, plot_title=plot_title, path2folder=path2folder)

    return fig


def plot_layer_delta_hv(
    df: pd.DataFrame,
    layer_cut: int,
    arch: str,
    metric: str,
    path2folder: str = "results/summary",
):
    """A function for plotting and saving layer delta"""

    plot_title = "{} layer {} at x = {}".format(arch.upper(), metric, layer_cut)

    print(f"Plotting {plot_title}...")

    if arch == "esm":
        alpha = 0.8
    else:
        alpha = "ptp"

    delta_scatter = hv.render(
        hv.Scatter(df, kdims=["x-0"], vdims=["f-x", "task", "model_size", "ptp"],).opts(
            color="task",
            cmap={
                l: c
                for l, c in zip(
                    list(TASK_LEGEND_MAP.values()),
                    TASK_COLORS,
                )
            },
            alpha=alpha,
            line_width=2,
            width=800,
            height=400,
            legend_position="right",
            legend_offset=(1, 0),
            size=np.log(dim("model_size") + 1) * 1.5,
            title=plot_title,
        )
    )

    # turn off legend box line
    delta_scatter.legend.border_line_alpha = 0

    print(f"Saving to {path2folder}...")

    BokehSave(
        bokeh_plot=delta_scatter,
        path2folder=path2folder,
        plot_name=plot_title,
        plot_width=800,
    )
```
