## Supplemental data and code for "Feature Reuse and Scaling: Understanding Transfer Learning with Protein Language Models": res_vis_task.py

```
"""A tempo clean up for summary plot"""

# a script for vis results from summary data df
from __future__ import annotations

import ast

import os
import itertools

import math

import numpy as np
import pandas as pd


from scipy.stats import spearmanr

import matplotlib as mpl
import matplotlib.pyplot as plt
import matplotlib.ticker as ticker
from matplotlib.lines import Line2D
from matplotlib.patches import Rectangle
from matplotlib.gridspec import GridSpec
from matplotlib.colors import ListedColormap


import seaborn as sns
import holoviews as hv
from holoviews import dim

hv.extension("bokeh")

from scr.vis.vis_utils import BokehSave, save_plt
from scr.params.emb import (
    MODEL_SIZE,
    EMB_MODEL_SIZE,
    EMB4TASK,
    MODEL_LAYER,
    EMB_MODEL_LAYER,
    CARP_MODEL_LAYER,
    CARP_INFO,
    ARCH_TYPE,
    ARCH_BAR_LAYER,
    ARCH_AB,
    ARCH_AB_DICT,
    CHECKPOINT_PERCENT,
    CARP_CHECKPOINT_LOSSES,
    EMB_SIMPLE_MAP,
    EMB_SIZE_SIMPLE,
    BASELINE_NAME_DICT,
    EMB_SIZE_NAME_SIMPLE
)
from scr.params.vis import (
    ORDERED_TASK_LIST,
    TASK_LEGEND_MAP,
    TASK_COLORS,
    TASK_SIMPLE_COLOR_MAP,
    PLOT_EXTS,
    CARP_ALPHA,
    ARCH_LINE_STYLE_DICT,
    ARCH_DOT_STYLE_DICT,
    ARCH_AB_DOT_STYLE_DICT,
    ARCH_SCATTER_STYLE_DICT,
    LAYER_ALPHAS,
)
from scr.analysis.utils import INIT_DICT, INIT_SIMPLE_LIST, SIMPLE_METRIC_LIST
from scr.utils import checkNgen_folder


task_cluster = {
    "well-aligned": ["SS3 - CB513", "SS3 - TS115", "SS3 - CASP12"],
    "benefit": ["Thermostability", "GB1 - low vs high", "AAV - two vs many", "AAV - one vs many"],
    "nobenefit": ["Subcellular localization", "GB1 - sampled", "GB1 - two vs rest"],
}

# if include small and medium
add_sm = True

selected_emb_s = {"esm": "esm1_t6_43M_UR50S", "carp": "carp_38M"}
selected_emb_m = {"esm": "esm1_t12_85M_UR50S", "carp": "carp_76M"}
selected_emb = {"esm": "esm1b_t33_650M_UR50S", "carp": "carp_640M"}

# if for main text filter again for model in EMB4TASK
concise = True
large_esm = "esm1b_t33_650M_UR50S"

mild_3 = ["#f1d384", "#9dae88", "#a3bfd6"]  # lighter yellow green blue s m l
bright_3 = ["#F9BE00", "#73A950", "#00A1DF"]  # yellow green blue s m l
dark_3 = ["#cabc8c", "#4c5835", "#376977"]  # yellow green blue
deep_3 = ["#cabc8c", "#005851", "#003B4C"] # may need new dark yellow


class PlotResultByTask:
    """
    A class handling plotting results in summary task class
    """

    def __init__(
        self,
        sum_folder: str = "results/summary",
        sum_df_name: str = "all_results",
    ) -> None:

        self._sum_folder = checkNgen_folder(os.path.normpath(sum_folder))
        self._sum_df_name = sum_df_name

    def _plot_by_cluster(self, metric: str = "test_performance_2"):
        """
        A method for plotting based on task behavior cluster, where for each task per row:
        - bar plot for onehot, L - rand, L - stat, S, M, L for both CARP and ESM
        - layer by layer for S, M, L CARP and ESM with onehot
        - CARP scale with pretrain loss
        """
        
        all_task_summary = {}

        for cluster, selected_tasks in task_cluster.items():

            numb_task = len(selected_tasks)

            # Create a three-degree nested multiclass bar plot
            fig, ax = plt.subplots(
                nrows=3, ncols=numb_task, figsize=(numb_task * 2.75 + 0.25, 9), sharey="row"
            )

            # Adjust layout with space between rows
            fig.subplots_adjust(hspace=0.3)  # Adjust the space here

            # Flip the x-axis so loss from left to right are reducing
            ax[2, 0].invert_xaxis()

            # Share x-axis within each column
            for row in range(3):
                for col in range(1, numb_task):
                    ax[row, col].sharex(ax[row, 0])

            # set up task / row wise
            for i, (l, t) in enumerate(
                {
                    "a)": "Last layer baseline and embedding performance",
                    "b)": "Layer by layer performance",
                    "c)": "Pretraining loss vs. last layer performance",
                }.items()
            ):

                if i == 0:
                    added_space = 0.02
                else:
                    added_space = -0.02

                title_y = ax[i, 0].get_position().y0 + 0.205 + added_space
                
                # add subfig label
                fig.text(
                    0.05,
                    title_y+0.005,
                    l,
                    ha="left",
                    va="bottom",
                    fontsize="large",
                )
                ax[i, 0].set_ylabel("Test performance")

            for i, task in enumerate(selected_tasks):

                df1, df2 = get_taskdf2plot(
                    df=self.prepped_df, metric=metric, task=task, large_esm=large_esm, concise=concise
                )

                c = TASK_SIMPLE_COLOR_MAP[task]

                bar_x_labels = list(BASELINE_NAME_DICT.values()) + [""] + EMB_SIZE_NAME_SIMPLE
                # there are two ones
                onehot_val = df1[(df1["task"] == task) & (df1["model"] == "onehot")][
                    "last_value"
                ].values[0]

                if task == "AAV - two vs many":
                    onehot_val = round(onehot_val) + 0.005

                # do summary dict before plotting
                all_task_summary[task] = task_summary(
                    task=task,
                    df1=df1,
                    df2=df2,
                    large_esm=large_esm,
                    error_margin=0.1,
                    tolerance=0.2,
                    window_size=6
                )

                ax[0, i].scatter(
                    bar_x_labels[0],
                    onehot_val,
                    s=150,
                    linewidth=1.2,
                    color="#666666",
                    facecolor="#666666",
                    edgecolor="#666666",
                    marker="s",
                    alpha=0.8,
                )

                for arch in ARCH_TYPE:

                    arch_df = df1[df1["arch"] == arch].copy()

                    # rand
                    ax[0, i].scatter(
                        bar_x_labels[1],
                        pair_taskwy(df=arch_df, val_col="last_value")[1],
                        s=150,
                        linewidth=1.2,
                        marker=ARCH_SCATTER_STYLE_DICT[arch],
                        facecolor="#666666",
                        edgecolor=bright_3[-1],
                        alpha=0.4,
                    )

                    # stat
                    ax[0, i].scatter(
                        bar_x_labels[2],
                        pair_taskwy(df=arch_df, val_col="last_value")[2],
                        s=150,
                        linewidth=1.2,
                        marker=ARCH_SCATTER_STYLE_DICT[arch],
                        edgecolor=bright_3[-1],
                        facecolor=dark_3[-1],
                        alpha=0.8,
                    )
                    # add some space in the middle
                    ax[0, i].scatter(
                        bar_x_labels[3],
                        None,
                        s=150,
                        c="w",
                        alpha=1,
                    )

                    # then by size
                    ax[0, i].scatter(
                        bar_x_labels[4:],
                        pair_taskwy(df=arch_df, val_col="last_value")[3:],
                        s=150,
                        linewidth=1.2,
                        marker=ARCH_SCATTER_STYLE_DICT[arch],
                        c=bright_3,
                        alpha=0.8,
                    )

                # Add onehot cross board
                ax[0, i].axhline(
                    y=onehot_val, color="#666666", alpha=0.8, linestyle="dotted", linewidth=1.2
                )

                # Add a vertical dotted line in the middle of the plot
                middle_x = (ax[0, i].get_xlim()[1] - ax[0, i].get_xlim()[0]) / 2 - 0.28
                ax[0, i].axvline(
                    x=middle_x, color="gray", linestyle="dotted", linewidth=0.5, alpha=0.5
                )

                # ax[0, i].set_ylim([0, 1])
                ax[0, i].set_title(task)
                ax[0, i].tick_params(axis="x", rotation=30)

                # Hide the middle tick
                ticks = ax[0, i].xaxis.get_major_ticks()
                ticks[3].tick1line.set_visible(False)  # Hide the bottom tick line
                
                # do lbl
                for arch in ARCH_TYPE:

                    if add_sm:

                        for emb_size, emb_dict in enumerate([selected_emb_s, selected_emb_m, selected_emb]):

                            arch_marker = {"carp": {"linestyle": "solid"}, 
                                            "esm": {"linestyle": "dashed"}}

                            # emb
                            ax[1, i].plot(
                                df1[(df1["model"] == emb_dict[arch]) & (df1["ablation"] == "emb")][
                                    "value"
                                ].to_numpy()[0],
                                linewidth=1.2,
                                alpha=0.8,
                                color=bright_3[emb_size],
                                **arch_marker[arch]
                            )
                    else:

                        arch_marker = {"carp": {"color": bright_3[-1]}, 
                                            "esm": {"color": deep_3[-1]}}
                        
                        # emb
                        ax[1, i].plot(
                            df1[(df1["model"] == selected_emb[arch]) & (df1["ablation"] == "emb")][
                                "value"
                            ].to_numpy()[0],
                            linewidth=1.2,
                            alpha=0.8,
                            **arch_marker[arch]
                        )


                ax[1, i].set_xlabel("Layer number")
                # ax[1, i].set_ylim([0, 1])

                # overlay onehot
                ax[1, i].axhline(
                    y=onehot_val, color="gray", alpha=0.8, linestyle="dotted", linewidth=1.2
                )

                carp_df = (
                    df2[df2["ablation"] == "emb"]
                    .sort_values(["model", "ptp"], ascending=[True, False])
                    .copy()
                )

                # adjust to match bar alpha
                carp_color2match = {
                    n: c for n, c in zip(["carp_38M", "carp_76M", "carp_640M"], bright_3)
                }
                # carp_alaphs = np.linspace(1, 0.4, 4)

                for model in carp_df["model"].unique():
                    carp_model_df = carp_df[carp_df["model"] == model].copy()

                    xs = [CARP_CHECKPOINT_LOSSES[model][float(x)] for x in carp_model_df["ptp"]]
                    c = carp_color2match[model]

                    # for the line
                    ax[2, i].plot(
                        xs,
                        carp_model_df["last_value"],
                        linestyle="solid",
                        color=c,
                    )

                    # for the dots
                    ax[2, i].scatter(
                        xs,
                        carp_model_df["last_value"],
                        marker="o",
                        s=150,
                        linewidth=1.2,
                        color=c,
                        edgecolor=c,
                        # alpha=carp_alaphs,
                        alpha=0.8
                    )
                    ax[2, i].set_xlabel("Pretrain loss")

                    # make y with closest .1
                    if "loss" not in metric:
                        # Determine the y-axis limits
                        ymin, ymax = ax[2, i].get_ylim()

                        # Generate ticks at one decimal place within the current y-axis limits
                        ticks = np.arange(np.floor(ymin*10)/10, np.ceil(ymax*10)/10 + 0.1, 0.1)

                        # Set custom ticks
                        ax[2, i].yaxis.set_major_locator(ticker.FixedLocator(ticks))
                        ax[2, i].yaxis.set_major_formatter(ticker.FormatStrFormatter('%.1f'))

            # add alpha legend for carp sizes
            model_size_descript = [
                "Small", # : CARP-38M, ESM-43M
                "Medium", # : CARP-76M, ESM-85M
                "Large", # : CARP-640M, ESM-650M
            ]

            arch_legend_list = [None] * len(model_size_descript)

            for i, (n, c) in enumerate(zip(model_size_descript, bright_3)):

                arch_legend_list[i] = Line2D(
                    [0],
                    [0],
                    marker="o",
                    color=c,
                    linestyle="none",
                    label=n,
                    alpha=0.8,
                    markersize=12,
                    markerfacecolor=c,
                    markeredgecolor=c,
                )

            ax[2, 0].add_artist(
                ax[2, 0].legend(
                    handles=arch_legend_list,
                    ncol=3, loc='upper center', bbox_to_anchor=(0.5, -0.2),
                    markerscale=0.6,
                    columnspacing=0.08, handletextpad=0.02, handlelength=1,
                    title="Size",
                )
            )

            # Manually create arch type elements
            arch_legend_list = []

            for a in ARCH_TYPE:
                # dot 
                arch_legend_list.append(Line2D(
                    [0],
                    [0],
                    marker=ARCH_SCATTER_STYLE_DICT[a],
                    label=a.upper(),
                    alpha=0.8,
                    markersize=12,
                    markerfacecolor="gray",
                    markeredgecolor="none",
                    linestyle="none",
                ))
                # line
                arch_legend_list.append(Line2D(
                    [0],
                    [0],
                    alpha=0.8,
                    markersize=12,
                    color=bright_3[-1],
                    **arch_marker[a]
                ))
            
            arch_legend_list.append(
                Line2D(
                    [0],
                    [0],
                    linestyle="dotted",
                    markersize=10,
                    marker="s",
                    color="#666666",
                    alpha=0.8,
                    label="One-hot",
                )
            )
            # just the line now
            arch_legend_list.append(
                Line2D(
                    [0],
                    [0],
                    linestyle="dotted",
                    color="#666666",
                    alpha=0.8,
                )
            )

            ax[2, 1].add_artist(
                ax[2, 1].legend(
                    handles=arch_legend_list,
                    ncol=3, loc='upper center', bbox_to_anchor=(0.5, -0.2),
                    markerscale=0.6,
                    columnspacing=0.08, handletextpad=0.02, handlelength=1,
                    title="Model",
                )
            )

            # make ablation legend
            ab_legend_list = []
            ab_legend_list.append(
                Line2D(
                    [0],
                    [0],
                    marker="o",
                    alpha=0.4,
                    linestyle="none",
                    markersize=12,
                    markerfacecolor="#666666",
                    markeredgecolor=bright_3[-1],
                    label="Random init",
                )
            )

            ab_legend_list.append(
                Line2D(
                    [0],
                    [0],
                    marker="o",
                    alpha=0.8,
                    linestyle="none",
                    markersize=12,
                    markerfacecolor=dark_3[-1],
                    markeredgecolor=bright_3[-1],
                    label="Stat transfer",
                )
            )

            ax[2, 2].add_artist(
                ax[2, 2].legend(
                    handles=ab_legend_list,
                    ncol=3, loc='upper center', bbox_to_anchor=(0.5, -0.2),
                    markerscale=0.6,
                    columnspacing=0.08, handletextpad=0.02, handlelength=1,
                    title="Ablation",
                )
            )

            # Adjust layout and white space
            plt.subplots_adjust(wspace=0.125)  # Adjust the white space between columns

            # Manually set subplot positions
            for i, a in enumerate(ax.flatten()):

                if i // numb_task >= 1:
                    # Get the bounding box of the second row's subplot in figure coordinates
                    bbox = a.get_position()

                    # Set the new position for the subplot
                    a.set_position([bbox.x0, bbox.y0 - 0.025, bbox.width, bbox.height ])

            # Add a super title for the figure with padding
            # fig.suptitle(f"Summary by {cluster}", fontsize="xx-large", x=0.5125, y=0.96)

            save_plt(
                fig, plot_title=f"Summary for {cluster}", 
                path2folder=checkNgen_folder(os.path.join("results/summary/bytask", metric))
            )

            # clean up the df
            # Convert nested dictionary to DataFrame
            all_task_summary_df = pd.DataFrame.from_dict(all_task_summary, orient="index")
            # Set the name of the index
            all_task_summary_df.index.name = "Tasks"
            all_task_summary_df = all_task_summary_df.reindex(
                list(itertools.chain(*task_cluster.values()))
            ).rename(columns={})
            # reorder
            selected_col_df = all_task_summary_df[[
                "Transfer > One-hot",
                "Transfer > Random init",
                "Transfer > Stat transfer",
                "Scale with PLM sizes",
                "Scale with layer depths",
                "Scale with pretrain losses",
            ]].copy().T

            # Generate custom annotations based on the conditions
            annotations = np.where(
                selected_col_df.astype(float) > 0.5,
                "✓",
                np.where(selected_col_df.astype(float) < 0.5, "✗", "~"),
            )

            # Plotting the heatmap
            fig, ax = plt.subplots(figsize=(5, 2))

            sns.heatmap(
                selected_col_df.astype(float),
                ax=ax,
                annot=annotations, fmt="",
                cmap=ListedColormap("white"),
                cbar=False,
                linewidths=0.5,
                linecolor="none",
            )
            
            # recolor# Iterate over text annotations in the Axes
            for text in ax.texts:
                value = text.get_text()
                # Set custom colors based on the annotation text
                if value == "✓":
                    text.set_color(bright_3[-1])
                    text.set_fontsize(14)  # Make the font larger
                    text.set_weight('bold')  # Make the text bolder
                elif value == "✗":
                    text.set_color('#666666')
                elif value == "~":
                    text.set_color(dark_3[-1])

            # Hide the middle tick
            ax.tick_params(axis="x", length=0)
            ax.tick_params(axis="y", length=0)
            ax.xaxis.tick_top()
            ax.set_xticklabels(ax.get_xticklabels(), rotation=45, ha="left")

            # remove x axis task titel
            ax.set_xlabel("")
            ax.set_ylabel("")

            save_plt(
                fig, plot_title=f"Summary checkbox for tasks", 
                path2folder=checkNgen_folder(os.path.join("results/summary/bytask", metric))
            )

    @property
    def result_df_path(self) -> str:
        """Return full summary result csv path"""
        df_path = os.path.join(
            os.path.normpath(self._sum_folder), self._sum_df_name + ".csv"
        )

        assert os.path.exists(df_path), f"{df_path} does not exist"

        return df_path

    @property
    def result_df(self) -> pd.DataFrame:

        """Return full result df with value cleaned up"""

        result_df = pd.read_csv(self.result_df_path)

        # check column name existance
        for c in ["metric", "ablation", "arch", "value", "task", "model", "ptp"]:
            assert c in result_df.columns, f"{c} not in df from {self.result_df_path}"

        # Convert the string of lists to NumPy arrays
        result_df["value"] = result_df["value"].apply(ast.literal_eval).apply(np.array)

        # make ptp float
        result_df["ptp"] = result_df["ptp"].astype(float)

        return result_df

    @property
    def prepped_df(self) -> pd.DataFrame:

        """Return a more plotting compatible df"""

        prepped_df = self.result_df.copy()

        # add task type and model size details for plotting legends
        prepped_df["task_type"] = prepped_df["task"].str.split("_").str[0]
        prepped_df["model_size"] = prepped_df["model"].map(MODEL_SIZE)
        prepped_df["model_layer"] = prepped_df["model"].map(MODEL_LAYER)

        # get rid of pooling details
        prepped_df["task"] = prepped_df["task"].str.replace("_mean", "")
        prepped_df["task"] = prepped_df["task"].str.replace("_noflatten", "")

        # sort based on given task order for plot legend
        prepped_df["task"] = pd.Categorical(
            prepped_df["task"], categories=ORDERED_TASK_LIST, ordered=True
        ).map(TASK_LEGEND_MAP)
        prepped_df = prepped_df.sort_values(["task", "ptp"], ascending=[True, False])

        return prepped_df


def get_taskdf2plot(
    df: pd.DataFrame,
    task: str,
    metric: str,
    large_esm: str == "esm1b_t33_650M_UR50S",
    concise: bool = True,
):
    """
    A function define the bars to be plotted based on each task

    Args:
    - df: pd.DataFrame,
    - task: str,
    - large_esm: str == "esm1b_t33_650M_UR50S",
    - concise: bool = True
    """
    #  & (prepped_df["ptp"].isin([0, 1]))].copy()
    if "last_value" not in df.columns:
        df["last_value"] = df["value"].apply(
            lambda x: x[-1] if len(x) > 0 else None
        ).copy()

    task_df = df[(df["metric"] == metric) & (df["task"] == task)].copy()

    slice_df = task_df[
        (
            ((task_df["ablation"] == "emb"))  # get emb
            | (task_df["ablation"] == "onehot")  # get onehot
            | (
                (task_df["ablation"].isin(["rand", "stat"]))
                & (task_df["model"].isin([large_esm, "carp_640M"]))
            )
        )
    ].copy()

    # if further slice based on more concise
    select_model = slice_df["model"].isin(EMB4TASK)
    select_ptp = slice_df["ptp"].isin([0, 1])
    select_cp = (slice_df["arch"] != "esm") & (
        slice_df["ablation"].isin(["emb", "onehot"])
    )

    if concise:
        return (
            slice_df[select_model & select_ptp].copy(),
            slice_df[select_model & select_cp].copy(),
        )
    else:
        return (
            slice_df[select_ptp].copy(),
            slice_df[select_cp].copy(),
        )
    
def pair_taskwy(df: pd.DataFrame, val_col: str == "last_value") -> np.array:

    """A function to pair x for task based plots with y"""

    baseline_df = df[df["ablation"] != "emb"].copy()
    emb_df = df[df["ablation"] == "emb"].copy()

    baseline_df["ablation"] = pd.Categorical(
        baseline_df["ablation"],
        categories=list(BASELINE_NAME_DICT.keys()),
        ordered=True,
    )

    return list(baseline_df.sort_values("ablation")[val_col].values) + list(
        emb_df.sort_values("model_size")[val_col].values
    )

def eval_emb_vs_ab(
    df1: pd.DataFrame,
    randstat: str,
    error_margin: float = 0.1,
    large_esm: str = "esm1b_t33_650M_UR50S",
) -> float:
    """
    A function that counts largest emb vs ablation

    Args:
    - delta: float = 0.1, 5% increase at least
    """

    emb_vs_ab = 0
    # emb > rand * 1.05 & emb > stat * 1.05
    for largest in ["carp_640M", large_esm]:
        emb_perf = df1[(df1["model"] == largest) & (df1["ablation"] == "emb")][
            "last_value"
        ].values
        emb_ab = df1[(df1["model"] == largest) & (df1["ablation"] == randstat)][
            "last_value"
        ].values

        for ab in emb_ab:
            if emb_perf > ab * (1 + error_margin):
                emb_vs_ab += 1

    return emb_vs_ab / 2


# Check if the array is monotonically increasing within a 5% error margin
def is_monotonically_increasing_with_error(
    vals: np.array, error_margin: float = 0.1, tolerance: float = 0.2
) -> bool:
    """
    A function to see if a given array is monotonically increasing

    Args:
    - error_margin: float = 0.1, allowing each data point 5% error
    - tolerance: float = 0.2, allowing such 5% happen 20% out of the all data points
    """

    cap = len(vals) * tolerance
    exception_count = 0

    for i in range(1, len(vals)):
        if exception_count > cap:
            return False
        # Calculate the minimum value the next element should have to be considered as increasing
        min_value = vals[i - 1] * (1 - error_margin)
        if vals[i] < min_value:
            return False
        elif vals[i] < vals[i - 1]:
            exception_count += 1

    return True


def task_summary(
    task: str,
    df1: pd.DataFrame,
    df2: pd.DataFrame,
    large_esm: str = "esm1b_t33_650M_UR50S",
    error_margin: float = 0.1,
    rho_cutoff: float = 0.9,
    tolerance: float = 0.2,
    window_size: int = 6,
) -> dict:

    """
    A function summarize a given task in terms of:

    Emb > OH
    Emb > Ab
    Scale with model size
    Scale with layer
    Scale with pretrain

    Args:
    - error_margin: float = 0.1, allowing each data point 5% error
    - rho_cutoff: float = 0.9, to what degree we call scale
    - tolerance: float = 0.2, allowing such 5% happen 20% out of the all data points
    - window_size: int = 5, moving average for the layer performance
    """

    summary_task_dict = {}

    # get onehot for the task
    onehot_val = df1[(df1["task"] == task) & (df1["model"] == "onehot")][
        "last_value"
    ].values[0]

    emb_df = df1[df1["ablation"] == "emb"].copy()

    # emb > onehot *1.05
    summary_task_dict["Transfer > One-hot"] = (
        sum(emb_df["last_value"] > onehot_val * (1 + error_margin)) / 6
    )
    summary_task_dict["Transfer > Random init"] = eval_emb_vs_ab(
        df1, "rand", error_margin
    )
    summary_task_dict["Transfer > Stat transfer"] = eval_emb_vs_ab(
        df1, "stat", error_margin
    )

    model_scale = 0
    layer_scale = 0

    for arch, large in zip(ARCH_TYPE, ["carp_640M", large_esm]):

        # do strickly greater than
        arch_model_scale = is_monotonically_increasing_with_error(
            emb_df[emb_df["arch"] == arch]
            .sort_values(by=["model_size"])["last_value"]
            .values,
            error_margin=0,
            tolerance=0,
        )
        summary_task_dict[f"Scale w {arch} size"] = arch_model_scale
        model_scale += arch_model_scale

        # make the layers smoother
        layer_perf = emb_df[(emb_df["model"] == large)]["value"].values[0]

        (
            layer_rho,
            summary_task_dict[f"{large} layer p"],
        ) = spearmanr(range(0, len(layer_perf)), layer_perf)

        if layer_rho >= rho_cutoff:
            arch_layer_scale = True
        else:
            arch_layer_scale = False

        summary_task_dict[f"{large} layer rho"] = layer_rho
        summary_task_dict[f"Scale w {large} layer"] = arch_layer_scale
        layer_scale += arch_layer_scale

    summary_task_dict["Scale with PLM sizes"] = model_scale / 2
    summary_task_dict["Scale with layer depths"] = layer_scale / 2

    pretrain_perf = (
        df2[df2["ablation"] == "emb"]
        .sort_values(by=["model_size", "ptp"])["last_value"]
        .values
    )

    (
        summary_task_dict["pretrain losses rho"],
        summary_task_dict["pretrain losses p"],
    ) = spearmanr(range(0, len(pretrain_perf)), pretrain_perf)

    if summary_task_dict["pretrain losses rho"] >= rho_cutoff:
        summary_task_dict["Scale with pretrain losses"] = True
    else:
        summary_task_dict["Scale with pretrain losses"] = False

    return summary_task_dict

def simplify_test_metric(metric: str) -> str:

    """
    A function to unify metric for plotting
    """

    for t in ["test_performance_1", "test_performance_2", "test_loss"]:
        metric = metric.replace(t, "test performance")

    return metric
```
