## Supplemental data and code for "Feature Reuse and Scaling: Understanding Transfer Learning with Protein Language Models": vis_utils.py

```
"""For general plotting functions"""

from __future__ import annotations

import os

import matplotlib.pyplot as plt

import bokeh
from bokeh.io import show, export_svg, export_png
from bokeh.plotting import show
from bokeh.models.annotations import Title

bokeh.io.output_notebook()

import holoviews as hv

hv.extension("bokeh")

from scr.params.vis import PLOT_EXTS
from scr.utils import checkNgen_folder


def save_plt(fig, plot_title: str, path2folder: str):

    """
    A helper function for saving plt plots
    """

    for ext in PLOT_EXTS:
        plot_title_no_space = plot_title.replace(" ", "_")
        plt.savefig(
            os.path.join(checkNgen_folder(path2folder), f"{plot_title_no_space}{ext}"),
            bbox_inches="tight",
            dpi=300
        )
    
    plt.close()


class BokehSave:
    """Export Bokeh plots"""

    def __init__(
        self,
        bokeh_plot: bokeh.plotting.figure,
        path2folder: str = "results/dataset_vis",
        plot_name: str = "plot",
        plot_exts: list = PLOT_EXTS,
        plot_height: int = 400,
        plot_width: int = 600,
        axis_font_size: str = "10pt",
        title_font_size: str = "10pt",
        x_name: str = "",
        y_name: str = "",
        gridoff: bool = True,
        showplot: bool = True
    ) -> None:
    
        self.bokeh_plot = bokeh_plot

        self.bokeh_plot.height = plot_height
        self.bokeh_plot.width = plot_width
        self.bokeh_plot.width_policy = "fixed"
        self.bokeh_plotheight_policy = "fixed"

        # change the axis title
        if x_name != "":
            self.bokeh_plot.xaxis.axis_label = x_name
        if y_name != "":
            self.bokeh_plot.yaxis.axis_label = y_name

        # Hide the grid
        if gridoff:
            self.bokeh_plot.xgrid.grid_line_color = None
            self.bokeh_plot.ygrid.grid_line_color = None

        # The subfolder for each landscape
        self.path2folder = path2folder

        # Check if the folder exists and create one if not
        os.makedirs(self.path2folder, exist_ok=True)

        # The actual name of the plot
        self.plot_name = plot_name

        # Add title to the plot itself
        t = Title()
        t.text = self.plot_name.replace("-", " ").replace("_", " ")
        self.bokeh_plot.title = t
        # self.bokeh_plot.title.text = self.plot_name.replace("-", " ").replace("_", " ")

        # adjust title and axis
        self.bokeh_plot.title.text_font_size = title_font_size
        self.bokeh_plot.xaxis.axis_label_text_font_size = axis_font_size
        self.bokeh_plot.xaxis.major_label_text_font_size = axis_font_size
        self.bokeh_plot.yaxis.axis_label_text_font_size = axis_font_size
        self.bokeh_plot.yaxis.major_label_text_font_size = axis_font_size

        # done formatting
        if showplot:
            show(self.bokeh_plot)

        # A list of the extensions
        self.plot_exts = plot_exts

        assert isinstance(
            self.plot_exts, (list, str)
        ), "plot_exts should be a string or a list of strings"
        # Convert to a list if it is a string
        if isinstance(self.plot_exts, str):
            self.plot_exts = [self.plot_exts]

        # Check if the extension has the period and add one if not
        self.plot_exts = ["." + ext if ext[0] != "." else ext for ext in self.plot_exts]

        assert (
            len([ext for ext in self.plot_exts if ext not in PLOT_EXTS]) == 0
        ), "More extension than supported"

        # Call the function to save the paths and return the paths
        self._plotpaths = self.get_path_export_plots()

        # Actually save plots
        self.export_plots()

    def get_path_export_plots(self):
        """Check the extension and join the parts into a path"""
        self._plotpaths = [os.path.join(self.path2folder, self.plot_name)] * len(
            self.plot_exts
        )

        for p, plotext in enumerate(self.plot_exts):

            # Create full plot path
            self._plotpaths[p] = self._plotpaths[p] + plotext
            # self.export_plots(plotext, self._plotpaths[p])

        return self._plotpaths

    def export_plots(
        self,
    ):
        """Export based on the plot extension"""

        for (plotext, plotpath) in zip(self.plot_exts, self._plotpaths):

            if plotext == ".svg":
                export_svg(self.bokeh_plot, filename=plotpath, timeout=30000)
            elif plotext == ".png":
                # Need to remove the tool bar and logos
                self.bokeh_plot.toolbar.logo = None
                self.bokeh_plot.toolbar_location = None
                export_png(self.bokeh_plot, filename=plotpath, timeout=30000)

    @property
    def plotpaths(self):
        return self._plotpaths
```
